# Elucidation and refinement of synthetic receptor mechanisms

**DOI:** 10.1101/2020.04.16.045039

**Authors:** Hailey I. Edelstein, Patrick S. Donahue, Joseph J. Muldoon, Anthony K. Kang, Taylor B. Dolberg, Lauren M. Battaglia, Everett R. Allchin, Mihe Hong, Joshua N. Leonard

## Abstract

Synthetic receptors are powerful tools for engineering mammalian cell-based devices. These biosensors enable cell-based therapies to perform complex tasks such as regulating therapeutic gene expression in response to sensing physiological cues. Although multiple synthetic receptor systems now exist, many aspects of receptor performance are poorly understood. In general, it would be useful to understand how receptor design choices influence performance characteristics. In this study, we examined the modular extracellular sensor architecture (MESA) and systematically evaluated previously unexamined design choices, yielding substantially improved receptors. A key finding that might extend to other receptor systems is that the choice of transmembrane domain (TMD) is important for generating high-performing receptors. To provide mechanistic insights, we adopted and employed a Förster resonance energy transfer (FRET)-based assay to elucidate how TMDs affect receptor complex formation and connected these observations to functional performance. To build further insight into these phenomena, we developed a library of new MESA receptors that sense an expanded set of ligands. Based upon these explorations, we conclude that TMDs affect signaling primarily by modulating intracellular domain geometry. Finally, to guide the design of future receptors, we propose general principles for linking design choices to biophysical mechanisms and performance characteristics.

## INTRODUCTION

Engineered cell-based therapies are a promising strategy for the targeted treatment of many diseases (1-4). Central to this approach is the use of genetically encoded sense-and-response programs, which cause the cell to enact a therapeutic function upon detection of specified cues. Designing and implementing a customized functional program generally requires integrating native and engineered cellular components, including receptors, signal transduction pathways, and genetic regulators. Developing principles and tools for doing so is an active frontier in the field of synthetic biology. This study focuses on elucidating principles that serve the broad goal of building, refining, and utilizing synthetic receptor systems.

Synthetic receptor systems can be designed either to interact with or to be independent of endogenous signaling (5, 6). One strategy is to couple synthetic receptor-mediated signaling to endogenous pathways such as those involving JAK/STAT, MAPK/ERK, PLCG, PI3K/AKT, NFAT, and mediators downstream of GPCRs (7-10). When native signal mediators are paired with downstream engineered promoters, these signals can be redirected to new transcriptional outputs (11, 12) or cell behaviors (13, 14). A second strategy involves using engineered components to redirect the signaling outputs from native receptors via modified phosphorylation (15-18), dimerization (19), or protein recruitment events (20). A third strategy is to employ *orthogonal* systems with mechanisms that are essentially independent of endogenous signaling and regulation. Examples of this approach include synthetic Notch (synNotch) receptors, which sense surface-bound ligands (21-24), and the modular extracellular sensor architecture (MESA), which detects soluble ligands (25-28). Both of these receptor systems can regulate either endogenous or exogenous genes directly through ligand binding-induced release of a transcription factor (TF). Orthogonal receptor systems have several potential advantages, including evasion of inadvertent activation or repression by cellular factors, the potential for use in diverse cell types, and the potential to multiplex receptors to implement sophisticated functions. As a result, orthogonal systems are of great interest for building cell-based devices, and their intrinsic modularity should facilitate extensions to new ligand inputs and functional outputs (29). Notably, these systems have not been tuned through evolution nor studied deeply in the biological literature, yet their modular structure renders them uniquely suited to iterative improvement. Therefore, there exist unique opportunities for elucidating and improving the performance characteristics of orthogonal synthetic receptors.

This study focuses on exploring and improving MESA receptors—-a class of self-contained synthetic receptors that signal in a manner that is independent of endogenous cellular pathways (25-28). Each MESA receptor comprises two transmembrane chains that dimerize upon ligand binding, triggering an intracellular proteolytic *trans*-cleavage reaction that releases an initially sequestered TF (**Fig. 1a**). Across these studies, we demonstrated that obtaining desirable performance characteristics—low ligand-independent (background) signaling and high fold induction of signaling upon ligand addition—required tuning both the absolute and relative levels at which each receptor chain is expressed. While this phenomenon is not entirely different from what one observes with native receptors and other systems, it would be desirable to minimize this sensitivity to receptor expression level, for example to facilitate translational applications. Moreover, our computational analysis (27) indicated that if design changes could improve the two performance characteristics noted above, this could render biosensor function robust to variation in receptor expression level. These observations motivate this investigation into refining the MESA design.

**Fig. 1.**
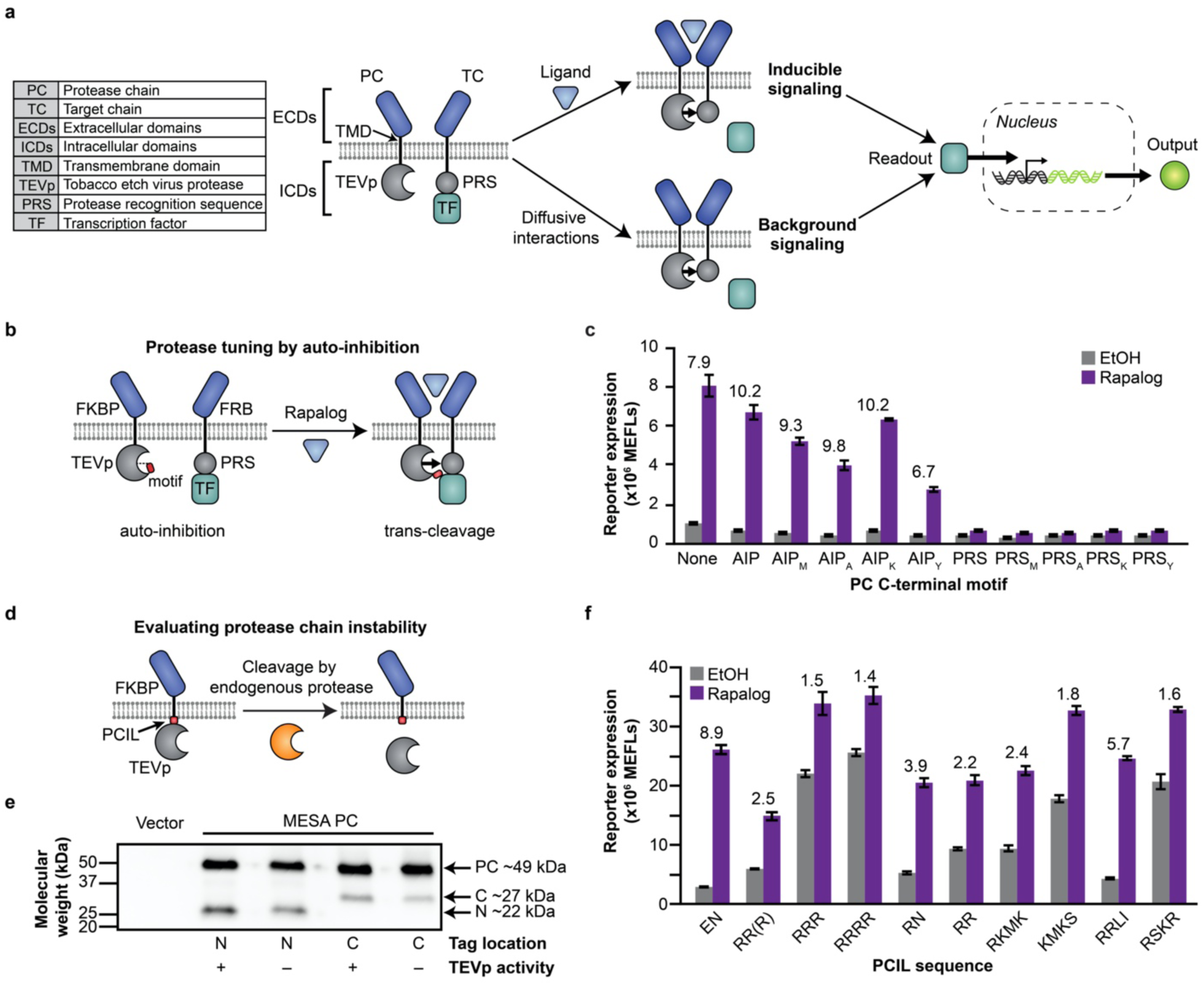
Protease chain tuning to improve MESA receptor performance. **a** This schematic depicts the MESA signaling mechanism. Ligand-induced receptor dimerization results in TEVp-mediated *trans*-cleavage to release a TF, which then enters the nucleus and induces target gene expression. Ligand-independent (background) receptor interactions can also result in TF release. **b**,**c** This proposed TEVp auto-inhibition strategy (**b**) was explored by functional evaluation (**c**) of MESA receptor variants in which a peptide—either a modified auto-inhibitory peptide (AIP: ELVYSQX) or protease recognition sequence (PRS: ENLYFQX), where X is a variable amino acid (e.g., AIP_M_: ELVYSQM)—has been appended onto the C-terminus of the protease chain (PC) TEVp. The leftmost condition is the base case (no peptide appended). Each condition uses a target chain (TC) with M at the P1’ site of the PRS. Numbers above bars indicate fold difference in reporter signal between samples treated with rapalog (dissolved in EtOH) vs. EtOH (vehicle-only control). Fold difference values are shown for samples in which ligand treatment induced significant signal above background (two-way ANOVA, *p* < 0.05). **d**,**e** Juxtamembrane cleavage of the PC (**d**) is suggested by Western blot analysis (**e**) of PCs tagged with 3x-FLAG on either the N-terminus or C-terminus; the PC appears to be cleaved into two fragments having sizes consistent with cleavage near the transmembrane domain (N = N-terminal product, C = C-terminal product). PCs with the TEVp^D81N^ mutation, which renders TEVp catalytically inactive (41), were cleaved similarly to the catalytically active PCs; thus we conclude that the observed cleavage can be attributed to endogenous cellular processes. **f** PCIL substitution generally led to decreased receptor performance through increased background and/or decreased induced signal (two-way ANOVA, *p* < 0.05). Fold difference values are shown above bars for samples in which ligand treatment induced significant signal above background (two-way ANOVA, *p* < 0.05). Bars depict the mean of three biological replicates, and error bars represent the S.E.M. Outcomes from ANOVA and Tukey’s HSD tests for **c**,**f** are in **Supplementary Notes 1 and 2**. In this experiment and subsequent experiments, we employ a rapamycin analog (rapalog) as a model ligand, which induces higher ligand-induced reporter expression than does rapamycin (**Supplementary Fig. 4a**).

In this investigation, we build mechanistic understanding and improve the functional performance of MESA receptors by systematically exploring variations upon MESA receptor design that we hypothesized might overcome the aforementioned limitations. We identify several features that may be rationally modified to tune performance (i.e., design handles), and we employ this knowledge to generate new high-performing receptors, expand the MESA repertoire to sense new ligands, and enhance a recently developed MESA variant that employs a distinct mechanism (26). Finally, we synthesize these observations to propose a framework of biophysically motivated design considerations for building novel synthetic receptors.

## MATERIALS AND METHODS

### General DNA assembly

Plasmid cloning was performed primarily using standard PCR and restriction enzyme cloning with Vent DNA Polymerase (New England Biolabs (NEB)), *Taq* DNA Polymerase (NEB), Phusion DNA Polymerase (NEB), restriction enzymes (NEB; Thermo Fisher), T4 DNA Ligase (NEB), Antarctic Phosphatase (NEB), and T4 PNK (NEB). Golden gate assembly and Gibson assembly were also utilized. The pBI-EYFP reporter was described previously (Addgene #58855) (25). GBP2-containing and GBP7-containing source plasmids were a generous gift from Constance Cepko (30). Plasmids were transformed into chemically competent TOP10 *E. coli* (Thermo Fisher), and cells were grown at 37°C.

### Cloning MESA receptors

MESA receptors were cloned into pcDNA backbones to confer high expression in HEK293FT cells. These plasmid backbones are versions of the pcDNA3.1/Hygro(+) Mammalian Expression Vector (Thermo Fisher #V87020), modified by our laboratory in previous work (Addgene #138749) (31). In general, restriction sites were chosen to facilitate modular swapping of parts via restriction enzyme cloning. A complete list of plasmids used in this study is provided in **Supplementary Data 1**. Plasmid maps are included as GenBank files in **Supplementary Data 2**.

### Plasmid preparation

TOP10 *E. coli* were grown overnight in 100 mL LB medium with an appropriate selective antibiotic. The following morning, cells were pelleted at 3,000×g for 10 min and then resuspended in 4 mL of a solution of 25 mM Tris pH 8.0, 10 mM EDTA, 15% sucrose, and 5 mg/mL lysozyme (Fisher Scientific #AAJ6070114). Cells were lysed for 15 min by addition of 8 mL of a solution of 0.2 M NaOH and 1% SDS, followed by neutralization with 5 mL 3 M sodium acetate (pH 5.2). The precipitate was pelleted by centrifugation at 9,000×g for 20 min. Supernatant was decanted and treated with RNase A for 1 h at 37°C. 5 mL phenol chloroform was added, and the solution was mixed and then centrifuged at 7,500×g for 20 min. The aqueous layer was removed and subjected to another round of phenol chloroform extraction with 7 mL phenol chloroform (Fisher Scientific #BP1752I). The aqueous layer was then decanted and subjected to an isopropanol precipitation (41% final volume isopropanol, 10 min at room temperature—approximately 22°C, 9,000×g for 20 min), and the pellet was briefly dried and resuspended in 420 μL water. The DNA mixture was incubated on ice for at least 12 h in a solution of 6.5% PEG 20,000 and 0.4 M NaCl (1 mL final volume). DNA was precipitated by centrifugation at 21,000×g for 20 min. The pellet was washed once with ethanol, dried for several h at 37°C, and resuspended for several h in TE buffer (10 mM Tris, 1 mM EDTA, pH 8.0). DNA purity and concentration were confirmed using a Nanodrop 2000 (Thermo Fisher).

### Cell culture

The HEK293FT cell line was purchased from Thermo Fisher/Life Technologies (RRID: CVCL_6911 [https://web.expasy.org/cellosaurus/CVCL_6911]) and was not further authenticated. Cells were cultured in DMEM (Gibco #31600-091) with 4.5 g/L glucose (1 g/L, Gibco #31600-091; 3.5 g/L additional, Sigma #G7021), 3.7 g/L sodium bicarbonate (Fisher Scientific #S233), 10% FBS (Gibco #16140-071), 6 mM L-glutamine (2 mM, Gibco #31600-091; 4 mM additional, Gibco #25030-081), penicillin (100 U/μL), and streptomycin (100 μg/mL) (Gibco #15140122), in a 37°C incubator with 5% CO_2_. Cells were subcultured at a 1:5 to 1:10 ratio every 2–3 d using Trypsin-EDTA (Gibco #25300-054). The HEK293FT cell line tested negative for mycoplasma with the MycoAlert Mycoplasma Detection Kit (Lonza #LT07-318).

### Transfection

Transient transfection of HEK293FT cells was conducted using the calcium phosphate method. Cells were plated at a minimum density of 1.5×10^5^ cells per well in a 24-well plate in 0.5 mL DMEM, supplemented as described above. For surface staining experiments, cells were plated at a minimum density of 3.0×10^5^ cells per well in a 12-well plate in 1 mL DMEM. For microscopy experiments, glass coverslips placed in 6-well plates were coated in a 0.1 mg/mL solution of poly-L-lysine hydrobromide (Sigma #P6282) for 5 min and left to dry overnight before plating 6×10^5^ cells per well in 2 mL DMEM. After at least 6 h, by which time the cells had adhered to the plate, the cells were transfected. For transfection, plasmids were mixed in H_2_O, and 2 M CaCl_2_ was added to a final concentration of 0.3 M CaCl_2_. The exact DNA amounts added to the mix per well and plasmid details for each experiment are listed in **Supplementary Data 3**. This mixture was added dropwise to an equal-volume solution of 2× HEPES-Buffered Saline (280 mM NaCl, 0.5 M HEPES, 1.5 mM Na_2_HPO_4_) and gently pipetted up and down four times. After 2.5–4 min, the solution was mixed vigorously by pipetting ten times. 100 μL of this mixture was added dropwise to the plated cells in 24-well plates, 200 μL was added to the plated cells in 12-well plates, or 400 μL was added to the plated cells in 6-well plates, and the plates were gently swirled. The next morning, the medium was aspirated and replaced with fresh medium. In some assays, fresh medium contained ligand and/or vehicle as described in **Supplementary Table 1** and indicated in figure legends.

Typically at 36–48 h post-transfection and at least 24 h post-media change, cells were harvested. As noted in figure captions, some experiments involved treatment with ligand or vehicle at later time points. In these cases, medium was still replaced the morning after transfection, and ligand diluted in serum-free DMEM was added as indicated. Cells were harvested for flow cytometry using FACS buffer (PBS pH 7.4, 2–5 mM EDTA, 0.1% BSA) or using 0.05% Trypsin-EDTA (Thermo Fisher Scientific #25300120) for 5 min followed by quenching with medium. The resulting cell solution was added to at least 2 volumes of FACS buffer. Cells were spun at 150×g for 5 min, supernatant was decanted, and fresh FACS buffer was added.

### Luciferase assays

Some functional assays used a luciferase readout (Dual-Glo, Promega #E2940) measured by a microplate reader (BioTek Synergy H1). Cells were transfected in biological triplicate with MESA receptor chains, an inducible Firefly luciferase, a constitutive *Renilla* luciferase, an inducible EYFP, a constitutive EBFP2, and an empty vector (as needed to maintain equal total plasmid mass across conditions). The day after transfection, vehicle and ligand treatments were applied during the medium change. Two days after transfection, EBFP2 and EYFP served as confirmatory microscopy readouts, and the two luciferase signals were quantified. Dual-Glo kit reagents were stored and prepared per the manufacturer-supplied instructions and equilibrated to room temperature; all steps were carried out at room temperature. At the time of assaying, medium was aspirated from cell cultures, and cells were washed with PBS. Passive lysis buffer stock (5X, Promega #E1941) was diluted in water, and the diluted buffer (120 μL, 1X) was added to each well. Plates were placed on a rocker for 15 min, after which the lysates were transferred into a 1.5 mL Eppendorf tube. Each tube was vortexed (15 s), and tubes were centrifuged (15,000×g, 30 s). 30 μL of supernatant was pipetted from each tube into a well of a 96-well plate (Thermo Fisher Scientific #655906). Luciferase reagent (30 μL, a volume equal to that of cell lysate) was pipetted into each well and mixed. Plates were incubated by rocking in the dark for 15 min.

Data were collected using the microplate reader’s luminescence fiber. For Firefly luciferase signal acquisition, autogain was set to scale the brightest wells to a value of 5,000,000 RLU. Integration time was set to 1 s and three technical replicate measurements per well were obtained. Stop & Glo reagent (30 μL, equal to that of the previous reagent) was added to each well, and plates were incubated by rocking in the dark for 15 min. For *Renilla* luciferase signal, the gain was set to 200, the integration time was set to 5 s, and three technical replicate measurements per well were obtained. For each well, the mean of the three technical replicates was calculated for each of the two luciferase readouts. Subsequently, the autoluminescence signal (calculated from the mean of the three vector-only transfected wells) was subtracted from each condition to background-normalize for each of the two luciferase readouts, respectively. Then for each biological replicate, Firefly luciferase signal was divided by *Renilla* luciferase signal. Quotients were linearly scaled such that the mean of the quotients for a condition transfected with only the reporter was equal to 1 a.u. Error was propagated accordingly.

### Western blotting

For Western blotting, HEK293FT cells were plated at 7.5×10^5^ cells per well in 2 mL DMEM in 6-well plates and transfected as above, using 400 μL transfection reagent per well (the reaction scales with the volume of medium). At 36–48 h after transfection, cells were lysed with 500 μL RIPA (150 mM NaCl, 50 mM Tris-HCl pH 8.0, 1% Triton X-100, 0.5% sodium deoxycholate, 0.1% sodium dodecyl sulfate) with protease inhibitor cocktail (Pierce/Thermo Fisher #A32953) and incubated on ice for 30 min. Lysate was cleared by centrifugation at 14,000×g for 20 min at 4°C, and supernatant was harvested. A BCA assay was performed to determine protein concentration, and after a 10 min incubation in Lamelli buffer (final concentration 60 mM Tris-HCl pH 6.8, 10% glycerol, 2% sodium dodecyl sulfate, 100 mM dithiothreitol, and 0.01% bromophenol blue) at 70°C (or 100°C for experiments that involved multiple co-transfected MESA receptors), protein (0.5 μg for experiments that were imaged with film, 10 to 25 μg for experiments that were imaged digitally) was loaded onto a 4–15% Mini-PROTEAN TGX Precast Protein Gel (Bio-Rad) and run either at 50 V for 10 min followed by 100 V for at least 1 h, or at 100 V for at least 1 h. Wet transfer was performed onto an Immuno-Blot PVDF membrane (Bio-Rad) for 45 min at 100 V. Ponceau-S staining was used to confirm protein transfer. Membranes were blocked for 30 min with 3% milk in Tris-buffered saline pH 8.0 (TBS pH 8.0: 50 mM Tris, 138 mM NaCl, 2.7 mM KCl, HCl to pH 8.0), washed once with TBS pH 8.0 for 5 min, and incubated for 1 h at room temperature or overnight at 4°C in primary antibody (Mouse-anti-FLAG M2 [Sigma #F1804, RRID: AB_262044 [http://antibodyregistry.org/AB_262044]), diluted 1:1000 in 3% milk in TBS pH 8.0. Primary antibody solution was decanted, and the membrane was washed once with TBS pH 8.0 and then twice with TBS pH 8.0 with 0.05% Tween, for 5 min each. Secondary antibody (HRP-anti-Mouse [CST 7076, RRID: AB_330924 [http://antibodyregistry.org/AB_330924]), diluted 1:3000 in 5% milk in TBST pH 7.6 (TBST pH 7.6: 50 mM Tris, 150 mM NaCl, HCl to pH 7.6, 0.1% Tween), was applied for 1 h at room temperature, and the membrane was washed three times for 5 min each time with TBST pH 7.6. The membrane was incubated with Clarity Western ECL Substrate (Bio-Rad) for 5 min, and then either exposed to film, which was developed and scanned, or digitally captured using an Azure c280 (Azure Biosystems). Images were cropped with Photoshop CC (Adobe). No other image processing was employed. Original images will be provided in **Source Data 2 upon final submission in a digital repository**.

### Expression normalization of MESA chains

Scanned Western blot images were imported into ImageJ and analyzed using the analyze gel feature. The intensity for each MESA chain band was quantified and reported as the percent of the total signal from all MESA bands on the blot; the same calculation was performed for all of the NanoLuciferase bands. This analysis was repeated for multiple images captured for each blot (including a range of exposure times to minimize bias) with non-detectable and saturated bands excluded from the analysis. The calculated intensity was averaged across all exposure times, and then MESA intensity (expression level) was divided by the NanoLuciferase intensity (expression level). This value was compared to the intensity calculated for the CD28-TMD rapamycin-binding TC sample, which was included on each blot as an internal cross-comparison control. This calculated ratio was used to adjust the doses of plasmids used for transfections in subsequent rounds of experiments, which were again evaluated by Western blots. The set of plasmid doses used in each round is in **Supplementary Table 2**.

### Immunohistochemistry

For surface staining, HEK293FT cells were plated at 3×10^5^ cells per well in 1 mL DMEM in 12-well plates and transfected as described above, using 200 μL transfection reagent per well. At 36–48 h after transfection, cells were harvested with 500 μL FACS buffer and spun at 150×g at 4°C for 5 min. For the experiment in which multiple harvest methods were compared (**Supplementary Fig. 14**), some samples were harvested using 0.05% Trypsin-EDTA (3 min or 10 min incubation, 37°C), which were then quenched with medium and added to two volumes of FACS buffer. Supernatant was decanted, and 50 μL fresh FACS buffer and 10 μL human IgG (Human IgG Isotype Control, ThermoFisher Scientific #02-7102, RRID: AB_2532958, stock concentration 1 mg/mL) was added. Cells were incubated in this mixture at 4°C for 5 min. 5 μL FLAG tag antibody (Anti-DDDDK-PE, Abcam ab72469, RRID: AB_1268475, or Anti-DDDDK-APC, Abcam ab72569, RRID: AB_1310127) was added at a concentration of 0.5 μg per sample and cells incubated at 4°C for 30 min. Following incubation, 1 mL FACS buffer was added, cells were spun at 150×g at 4°C for 5 min, and supernatant was decanted. This wash step was repeated two more times to total three washes. After decanting supernatant in the final wash, 1–3 drops of FACS buffer were added.

### Analytical flow cytometry

Flow cytometry was run on a BD LSR Fortessa Special Order Research Product (Robert H. Lurie Cancer Center Flow Cytometry Core). Lasers and filter sets used for data acquisition are listed in **Supplementary Table 3** (for experiments involving reporter expression), **Supplementary Table 4** (for experiments quantifying receptor expression on the cell surface), and **Table 1** (for experiments involving FRET). Approximately 2,000–3,000 single transfected cells were analyzed per sample, using a single transfection control or, when available, multiple transfected fluorophores for gating (e.g., mCerulean+/mVenus+ cells were classified as the subset of transfected cells of interest for FRET experiments).

**Table 1.**
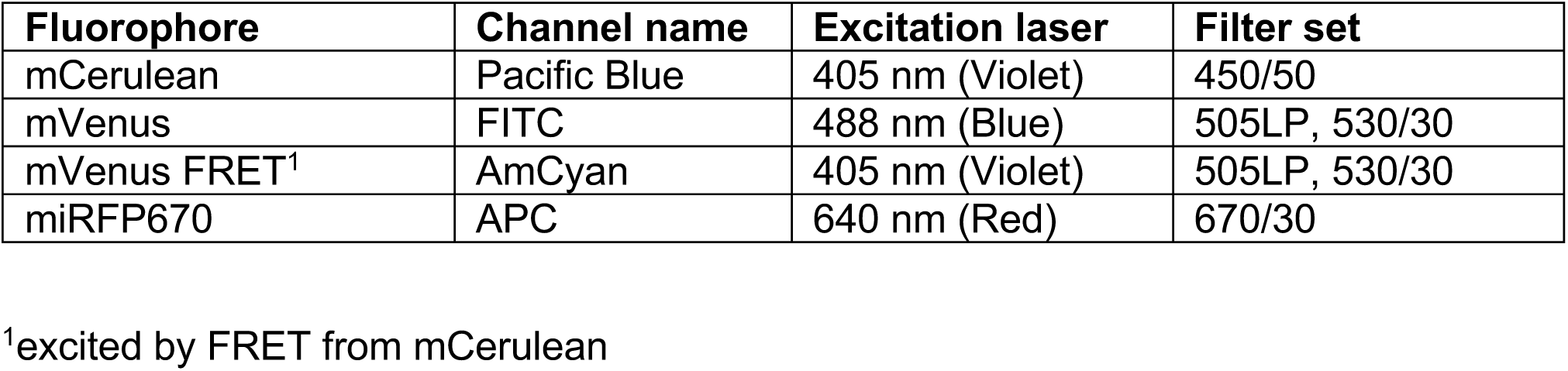
Instrument specifications for using analytical flow cytometry to quantify FRET

Samples were analyzed using FlowJo v10 software (FlowJo, LLC). Fluorescence data were compensated for spectral bleed-through. Additionally, spectral bleed-through compensation in FRET experiments included compensation of the fluorescence of either mCerulean or mVenus out of the AmCyan channel. As shown in **Supplementary Fig. 1**, the HEK293FT cell population was identified by SSC-A vs. FSC-A gating, and singlets were identified by FSC-A vs. FSC-H gating. To distinguish transfected from non-transfected cells, a control sample of cells was generated by transfecting cells with a mass of pcDNA (empty vector) equivalent to the mass of DNA used in other samples in the experiment. For the single-cell subpopulation of the pcDNA-only sample, a gate was made to identify cells that were positive for the constitutive fluorescent protein used as a transfection control in other samples, such that the gate included no more than 1% of the non-fluorescent cells.

### Quantification of reporter output

MESA signaling was quantified by measuring the expression of a downstream fluorescent reporter protein (**Fig. 1a**). For most experiments, the reporter protein was EYFP. For experiments in which the ligand was sGFP, DsRed2 was used as the reporter (instead of EYFP) to avoid spectral overlap with sGFP. The mean fluorescence intensity (MFI) for each relevant channel (as defined in **Supplementary Table 3**) of the single-cell transfected population was calculated and exported for further analysis. To calculate reporter expression, MFI in the FITC channel (for EYFP reporter) or PE-Texas Red channel (for DsRed2 reporter) was averaged across three biological replicates. From this number, cell autofluorescence was subtracted. To calculate cell autofluorescence, in each experiment, a control group of cells transfected with DNA encoding the fluorescent protein transfection control and pcDNA was used. The background-subtracted MFI was converted to Molecules of Equivalent Fluorescein (MEFLs) or Molecules of Equivalent PE-Texas Red (MEPTRs); as shown in **Supplementary Fig. 2**, to determine conversion factors for MFI to MEFLs and for MFI to MEPTRs, Rainbow Calibration Particles (Spherotech #RCP-30-5) or UltraRainbow Calibration Particles (Spherotech #URCP-100-2H) were run with each flow cytometry experiment. These reagents contain six (RCP) or nine (URCP) subpopulations of beads, each with a known number of various fluorophores. The total bead population was identified by FSC-A vs. SSC-A gating, and bead subpopulations were identified through two fluorescent channels. MEFL and MEPTR values corresponding to each subpopulation were supplied by the manufacturer. A calibration curve was generated for the experimentally determined MFI vs. the manufacturer-supplied MEFLs or MEPTRs, and a linear regression was performed with the constraint that 0 MFI equals 0 MEFLs or MEPTRs. The slope from the regression was used as the conversion factor, and error was propagated. Fold differences were calculated by dividing reporter expression with ligand treatment by the reporter expression without ligand treatment (vehicle). Standard error was propagated through all calculations.

### Quantification of FRET by flow cytometry

Detailed laser and filter setups for FRET data collection are listed in **Table 1**. Briefly, the donor fluorescence intensity was quantified using the Pacific Blue channel (λ_ex_ = 405nm, λ_em_ = 450/50nm), the acceptor fluorescence intensity was quantified using the FITC channel (λ_ex_ = 488nm, λ_em_ = 530/30nm), and the FRET fluorescence intensity was quantified using the AmCyan channel (λ_ex_ = 405nm, λ_em_ = 530/30nm). The mCerulean+/mVenus+ population was distinguished from samples transfected with the transfection control and pcDNA only, mCerulean only, and mVenus only. This gate was drawn such that less than 1% of the listed single-color samples were included. The normalized FRET (NFRET) parameter (32) was defined in the FlowJo workspace by dividing the compensated fluorescence intensity (FI) of a cell in the AmCyan channel by the square root of the product of the compensated FI of that cell in the Pacific Blue and FITC channels, as described by equation (1):

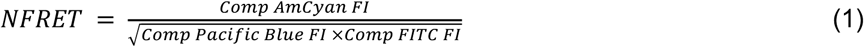

Average NFRET metrics were calculated for controls included in each experiment, including a negative control (cytosolic mCerulean and mVenus co-transfected) and positive control (membrane-tethered mCerulean-mVenus fusion protein). A calibrated NFRET parameter was defined based on the negative and positive FRET controls such that the NFRET of all samples is scaled linearly between these controls, as described by equation (2):

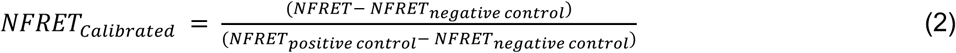

The calibrated NFRET metrics were exported, along with each mean FI in the relevant channels of the single-cell, transfected, mCerulean+/mVenus+ population for further analysis. In this study, all plotted NFRET metrics are calibrated values, and all FRET FI are compensated. Fold differences were calculated by dividing NFRET in the presence of ligand by NFRET in the absence of ligand (presence of vehicle only). Standard error was propagated through all calculations.

### Confocal microscopy

Confocal microscopy was performed on a Leica SP5 II laser scanning confocal microscope (Northwestern Chemistry of Life Processes Institute, Biological Imaging Facility) with a 100× (1.44 NA) oil-immersion objective. Coverslips were removed from media and mounted on a glass slide immediately before imaging. Ten fields of view were captured for each sample. Fields of view were selected using the brightfield channel to identify areas with adherent cells, and the z-axis was adjusted to focus on the central plane of most cells within the field of view before exposing the cells to lasers. All images were captured at a 512×512 image resolution with a scanning speed of 400 Hz. Excitation and emission settings were selected depending on which fluorophore (donor (D) or acceptor (A)) was excited and which emission was captured. The excitation and emission setup is described in **Supplementary Table 5**. Hybrid detector voltage settings were held constant for all samples within each experiment.

### Quantification of FRET by image processing

Images were exported as stacks and separated into single-channel images using Fiji software. A custom image processing script was produced in MATLAB to import single-channel images into matrices (**Supplementary Software**). To describe channels throughout this section, DD indicates that the donor was excited and emission from the donor was captured, AA indicates that the acceptor was excited and emission from the acceptor was captured, and DA indicates that the donor was excited and emission from the acceptor was captured. Empty vector-only (pcDNA) transfected samples were used to identify the upper limit of autofluorescence in each channel (DD, AA, DA), and thresholds were defined as the 99.9^th^ percentile of fluorescence of pixels in the vector-only samples in each channel. Pixels below this threshold were set to an intensity of zero. Saturated pixels were also removed from the matrices. Next, respective donor-only and acceptor-only controls (single-receptor transfections) were used to calculate spectral bleed-through parameters. These parameters, as defined in equations (3–6), were calculated by averaging across all pixels in ten fields of view, excluding pixels with infinite or undefined values. Intensities (I) are subscripted with DD, AA, or DA to indicate the excitation and emission conditions, and with (D) or (A) to indicate that this parameter was calculated using donor-only or acceptor-only samples, respectively.

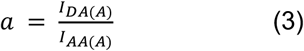

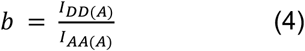

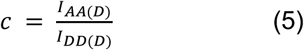

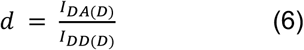

The parameters calculated for each pair of receptors were used to subtract the contribution of donor and acceptor fluorescence from the FRET channel fluorescence on a pixel-by-pixel basis as described in equation (7) and as reported previously (33). This step produced a corrected FRET fluorescence intensity (F_c_):

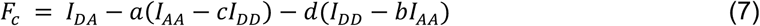

NFRET is calculated by normalizing F_c_ to the square root of the product of the donor and acceptor fluorescence intensities on a pixel-by-pixel basis as described in equation (8) and as reported previously (32).

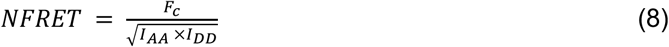

NFRET values were exported across pixels within each field of view and across all fields of view, then averaged to generate a mean NFRET metric for the entire sample. Corrected FRET intensity and NFRET matrices were plotted and visualized as processed images.

### Ectodomain distance estimations

Published crystal structures for ligand-bound ectodomains (ECDs) were analyzed using Chimera (FKBP/FRB PDB: 3FAP (34), GID1/DELLA PDB: 2ZSH (35), ABI1/PYL1 PDB: 3KDJ (36), G6-311 VEGF scFv PDB: 2FJG (37), B20-4.1 VEGF scFv PDB: 2FJH (37)). Residues that correspond to the C-termini used in our experimental receptor system were identified, and inter-termini distances were measured.

### Statistical analyses

Unless otherwise stated, three independent biological replicates were evaluated for each condition. The data shown reflect the mean across these biological replicates of the MFI of approximately 2,000–3,000 single, transfected cells or the mean NFRET or NFRET fold difference of approximately 2,000–3,000 single, transfected, mCerulean+/mVenus+ cells. Error bars represent the standard error of the mean (S.E.M.).

ANOVA tests and Tukey’s HSD tests were performed using RStudio. Tukey’s HSD tests were performed with *α* = 0.05. Pairwise comparisons were made using a two-tailed Welch’s *t*-test, which is a version of Student’s *t*-test in which the variance between samples is treated as not necessarily equal. Two-tailed Welch’s *t*-tests were performed in GraphPad. To decrease the false discovery rate, the Benjamini-Hochberg procedure was applied to each set of tests per figure panel; in all tests, after the Benjamini-Hochberg procedure, the null hypothesis was rejected for *p*-values < 0.05. The outcomes for each statistical test are provided in the figure captions, and additional details for some panels are in referenced supplementary notes. **Supplementary Note 1** includes the outcomes for one-way ANOVA tests followed by Tukey’s HSD tests for **Fig. 1c** and **Supplementary Fig. 3e. Supplementary Note 2** includes the outcomes for two-way ANOVA tests followed by Tukey’s HSD tests for **Fig. 4a,b** and **Fig. 6a–f. Supplementary Note 3** includes the outcomes for three-way ANOVA tests followed by Tukey’s HSD tests for **Fig. 4c, Supplementary Fig. 11, Fig. 6a–f**, and **Supplementary Fig. 28b. Supplementary Note 4** includes the outcomes for two-tailed Welch’s *t*-tests followed by the Benjamini-Hochberg procedure for **Supplementary Fig. 13b**.

## RESULTS

### Protease tuning reduces background

We initially focused on the goal of decreasing MESA receptor background signaling by investigating two strategies that we hypothesized could modulate the kinetics of proteolytic *trans*-cleavage. Our first strategy was motivated by extending a published observation. In our initial development of MESA (25), we used prior biochemical analyses of the tobacco etch virus protease (TEVp) (38-41) to vary cleavage kinetics by mutating the amino acid residue that immediately follows the cleavage site (the P1’ position) within the protease recognition sequence (PRS: ENLYFQX, where X is the P1’ position) on the target chain (TC). This search enabled the identification of a kinetic regime in which dimerization-inducible signaling was feasible. However, in the initial study, we did not explore all kinetic regimes that are accessible using known TEVp biochemistry. Here, we investigated whether TEVp kinetics could be subtly tuned to preferentially reduce background signaling more so than ligand-induced signaling. We introduced mutations to the TEVp on the protease chain (PC) that reduce cleavage kinetics to varying degrees (38-41) (**Supplementary Fig. 3a**), and we subsequently combined these with mutations to the P1’ position of the PRS on the TC (**Supplementary Fig. 3b**). We found many combinations of TC and PC variants that exhibited lower background signaling compared to the base case, as desired. However, the fold difference in reporter expression (the ratio of reporter expressed with vs. without ligand) did not improve; these receptor variants exhibited a decrease in ligand-induced signaling that was proportional to or greater than the decrease in background signaling, and therefore we chose not to pursue the strategy of tuning TEVp kinetics (alone) to improve receptor performance.

We next explored a second strategy motivated by the observation that TEVp naturally includes a C-terminal auto-inhibitory peptide (AIP) that is often omitted from biochemical tools that use TEVp (including MESA). Crystallographic evidence suggests that the AIP can reside in the TEVp active site (40), and since the PRS and AIP are similar in sequence (**Supplementary Fig. 3c**), it is possible that these peptides compete for binding to the active site of TEVp. We hypothesized that by placing variants of the PRS or AIP on the TEVp C-terminus, these peptides might reduce background signaling by reversibly occupying the TEVp active site such that TC-PC *trans*-cleavage is inhibited during transient diffusive encounters (in the absence of ligand), but cleavage would eventually occur during sustained ligand binding-induced chain dimerization (**Fig. 1b**). Appending the full AIP to TEVp decreased both background and ligand-induced signaling, and a similar outcome was observed when this effect was combined with the previously evaluated mutations in TEVp or the PRS (**Supplementary Fig. 3d**). Appending most AIP and PRS variants also produced this pattern (**Fig. 1c, Supplementary Fig. 3e**), but four appended peptides (ELVYSQ, ELVYSQM, ELVYSQA, ELVYSQK) slightly increased fold difference compared to the base case. Altogether, we conclude that adding active site-occupying peptides can modestly reduce background and improve fold inducibility, but other strategies are needed to more substantially improve receptor performance.

### Protease chain expression can be stabilized by linker selection

In prior work, we observed that PC surface expression was often lower than TC surface expression (25, 28), and we hypothesized that some aspects of PC design might render this chain less stable. Western blot analysis showed expression of the full-length PC as well as a smaller fragment, the size of which was consistent with juxtamembrane cleavage (**Fig. 1d,e**). Since this pattern was observed even when TEVp was mutated to be catalytically inactive via D81N substitution (41) (**Fig. 1d,e**), the observed cleavage can be attributed to endogenous cellular processes, rather than the catalytic activity of TEVp. Cleavage of the PC is potentially problematic because the residual membrane-tethered ectodomain (ECD) could function as a competitive inhibitor of intact receptors, and the TEVp released into the cytosol could contribute to background signaling. To explore alterations that might prevent cleavage of the PC, we first varied the sequence of the PC inner linker (PCIL) that connects the TEVp and transmembrane domain (TMD) by introducing positively charged residues and sequences from native receptors. These substitutions improved protein stability, reducing the appearance of the originally observed juxtamembrane cleavage product (**Supplementary Fig. 5a**). However, none of these substitutions improved receptor performance, and some substitutions diminished functional performance by increasing background signaling and/or decreasing induced signaling and therefore decreasing the ligand-induced fold difference (**Fig. 1f**). This effect could not be overcome by decreasing the PC plasmid dose (to compensate for the increased levels of intact PC) without diminishing ligand-induced signaling (**Supplementary Fig. 5b**,**c**), and additionally these substitutions reduced surface expression of the PC (**Supplementary Fig. 5d**). Although it is not clear why each substitution conferred these undesirable effects, it was clear that PCIL substitution alone did not address PC stability or background signaling via a useful mechanism, and so we turned to other modifications as alternative approaches.

### Transmembrane domain substitution can improve receptor performance and stabilizes the protease chain

TMD choice is an as-of-yet unexplored and potentially important aspect of MESA design. Previous MESA receptor designs (25, 28) employ a form of the CD28-TMD commonly used in chimeric antigen receptors (CARs) that differs somewhat from the native CD28-TMD sequence. When engineering CARs, TMD choice has proven to be a useful handle for tuning interactions between receptor chains and modulating the strength of target antigen binding-induced receptor signaling (42, 43). Since the native CD28 receptor clusters as a member of the immunological synapse formed between a T cell and an antigen-presenting cell (44, 45), we hypothesized that when the CD28-TMD is employed in a synthetic receptor system such as MESA, some residual clustering (mediated by the TMD alone) could lead to ligand-independent signaling. We speculated that using an alternative TMD might avoid this problem. In natural receptor tyrosine kinases (RTKs), the TMDs regulate diverse aspects of receptor signaling mechanisms, including dimerization propensity and geometry, ligand binding-induced rotational conformation changes, and clustering (46). Therefore, we decided to investigate whether replacing the CD28-TMD in MESA with other TMD variants might improve receptor performance.

We selected a panel of seven natural TMDs from RTKs, sampling a range of reported dimerization propensities (48), and two synthetic TMDs (47), and we substituted these for the CD28-TMD (**Fig. 2a**). TMD substitution had a substantial effect on receptor expression, and in some cases this effect differed for the TC and PC (**Supplementary Fig. 6a**). Since MESA performance depends upon both chain expression level and the ratio of expression of the two chains (27), we sought to isolate the mechanistic effects of TMD choice from the effects upon expression levels. Therefore, we normalized protein expression by iteratively varying plasmid dose **(Methods, Supplementary Fig. 6**). In this context, we observed that all of the TMD substitutions except for the FGFR1-TMD stabilized the PC (i.e., resolved the juxtamembrane cleavage observed with the CD28-TMD) (**Fig. 2b, Supplementary Fig. 5)**, and all substitutions altered surface expression (**Supplementary Fig. 7a**). In subsequent functional evaluations, using TC:PC protein expression ratios of approximately 1:1, TMD substitution conferred substantial, expression level-independent effects on performance **(Fig. 2c, Supplementary Fig. 7b**). Notably, employing the TMDs from GpA and FGFR4 increased the ligand-induced signaling compared to the CD28-TMD without increasing background signaling, leading to high-performing systems. Conversely, utilizing the TMD from FGFR1 increased both ligand-induced and background signaling, and the TMDs from FGFR2, FGFR3, EphA4, and VEGFR1 did not result in receptors that were capable of signaling.

**Fig. 2.**
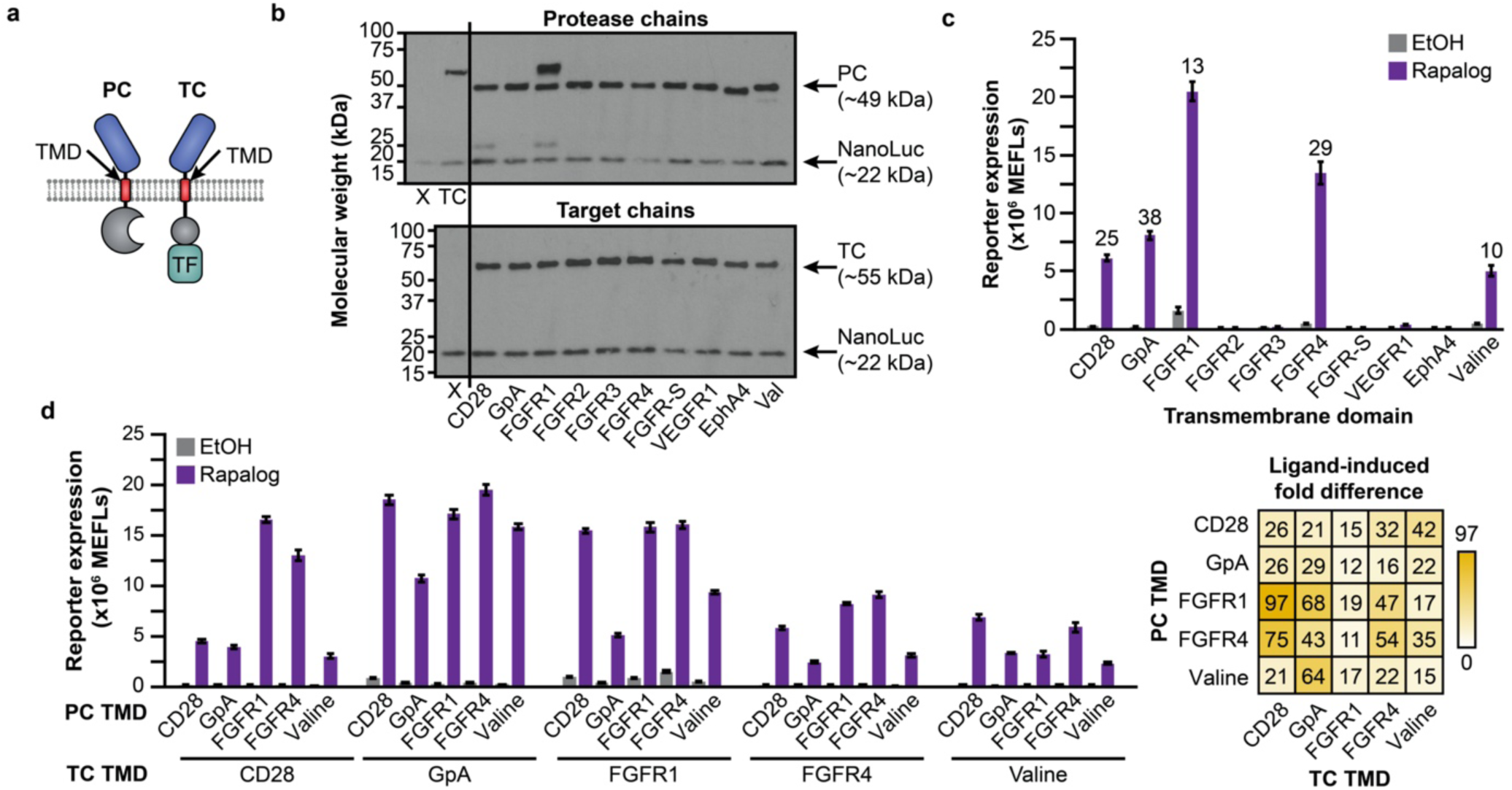
TMD contributions to MESA receptor signaling. **a** This schematic identifies the design choice examined here—the TMD sequence. **b** Effect of TMD choice on the expression of expected bands (PC, TC, co-transfected NanoLuc loading control) versus cleavage products (CD28 and FGFR1 cases). For this experiment, chain expression levels were first normalized to that of CD28-TMD TC expression (M, upper panel) by varying transfected plasmid dose through iterative Western blot analyses (**Supplementary Fig. 6**). The X denotes a vector-only negative control (including NanoLuc); TC denotes a CD28-TMD TC. **c** Paired TMD substitution conferred varying effects on receptor performance. Labels indicate the TMD that was used on both the TC and PC. Numbers above bars indicate fold difference when the ligand induced a significant signal above background (two-way ANOVA, *p* < 0.05). **d** Combinatorial TMD substitution further improved receptor performance. Fold difference is reported in the heatmap at right. All combinations exhibit a significant increase in reporter expression upon ligand treatment (three-way ANOVA, *p* < 0.001). Bars depict the mean of three biological replicates, and error bars represent the S.E.M. Outcomes from ANOVAs and Tukey’s HSD tests for **c**,**d** are in **Supplementary Notes 1, 2, and 3**.

To determine whether using different TMDs on the TC and PC could yield further improvements, we evaluated the 100 (10×10) pairwise TC-PC combinations (**Fig. 2d, Supplementary Fig. 8a**,**b**). In our initial screen, most TCs that conferred little or no detectable ligand-induced signaling when paired with a PC bearing the same TMD (matched pairs) also showed little or no ligand-induced signaling when paired with a different TMD PC (mixed pairs) (**Supplementary Fig. 8a**,**b**). An exception to this trend is the VEGFR1-TMD, which showed some ligand-induced signaling when paired with a PC containing the CD28-, GpA-, FGFR1-, FGFR3-, FGFR4-, or Valine-TMD. Notably, many other mixed TMD pairs also showed substantially improved performance compared to the matched CD28-TMD base case, including reduced background signal and/or increased ligand-induced signal, leading to fold differences as high as 97 (**Fig. 2d, Supplementary Fig. 8c**). Together, these results suggest that TMD choice is a key determinant of receptor performance and a rich target for tuning.

### Transmembrane domain choice does not substantially impact receptor dimerization propensity

Given the promising results obtained with certain TMD choices, we next sought mechanistic insight into the roles of these domains in MESA signaling. For native receptors, TMDs can affect both localization and function (46), but how this choice affects synthetic receptor function is unexplored. Since some TMD sequence motifs mediate receptor homodimerization, we hypothesized that TMD choice might affect MESA receptor performance by modulating the propensity for chains to associate. We first evaluated TMD association computationally using TMDOCK, a tool that uses amino acid sequence to predict TMD association by simulating alpha helix packing arrangements and conducting local energy minimization (48) (**Supplementary Fig. 9a–c**). This analysis predicted differences in matched TMD interactions, although the predicted trends only partially agreed with our experimental observations. For example, TMDOCK predicted the FGFR1-TMD to exhibit a high propensity to homodimerize, which is consistent with the observed high background signal (**Fig. 2c**). In contrast, TMDOCK also predicted the GpA-TMD to homodimerize (with more stability than the CD28-TMD), which is consistent with prior biochemical analyses (49, 50), yet we observed no evidence of enhanced homodimerization (i.e., in the form of high background) in functional assays. One possible explanation for this discrepancy is that TMDOCK evaluates TMD interactions in isolation, omitting any effects that might be conferred by the intracellular or extracellular receptor domains, and thus this tool alone is insufficient to explain the mechanistic consequences of TMD choice. Therefore, we next sought to directly investigate how TMD choice affects MESA receptor association by experimental characterization of full-length proteins.

To develop an assay for experimentally quantifying MESA inter-chain association, we employed a Förster resonance energy transfer (FRET)-based method (32, 51) (**Fig. 3a**). We replaced the TC and PC intracellular domains (ICDs) with mVenus or mCerulean—fluorescent proteins that exhibit FRET in a manner dependent on spatial co-localization (FRET occurs within a donor-acceptor distance of approximately 10 nm (52, 53)). Using this assay, ligand-independent association and ligand-dependent association are quantified by measuring acceptor (mVenus) fluorescence upon donor (mCerulean) excitation. To establish a high-throughput workflow that yields single-cell resolution data, we first adapted a reported approach to quantify FRET by flow cytometry (51). Importantly, this workflow includes normalization of FRET signal to fluorophore levels on a single-cell basis, which is necessary to account for heterogeneity in protein expression. In this approach, single-cell mCerulean donor fluorescence, mVenus acceptor fluorescence, mVenus FRET fluorescence, and fluorescence from constitutively expressed miRFP670 are quantified by flow cytometry (**Table 1**). Single-color samples are analyzed to apply post-hoc linear compensation for spectral bleed-through across fluorophore and FRET channels using an expression range that encompasses the receptor samples (**Fig. 3b, Supplementary Fig. 10a**); only transfected cells (miRFP670+) that are both mCerulean+ and mVenus+ are analyzed (**Supplementary Fig. 1a, Supplementary Fig. 10b**).

**Fig. 3.**
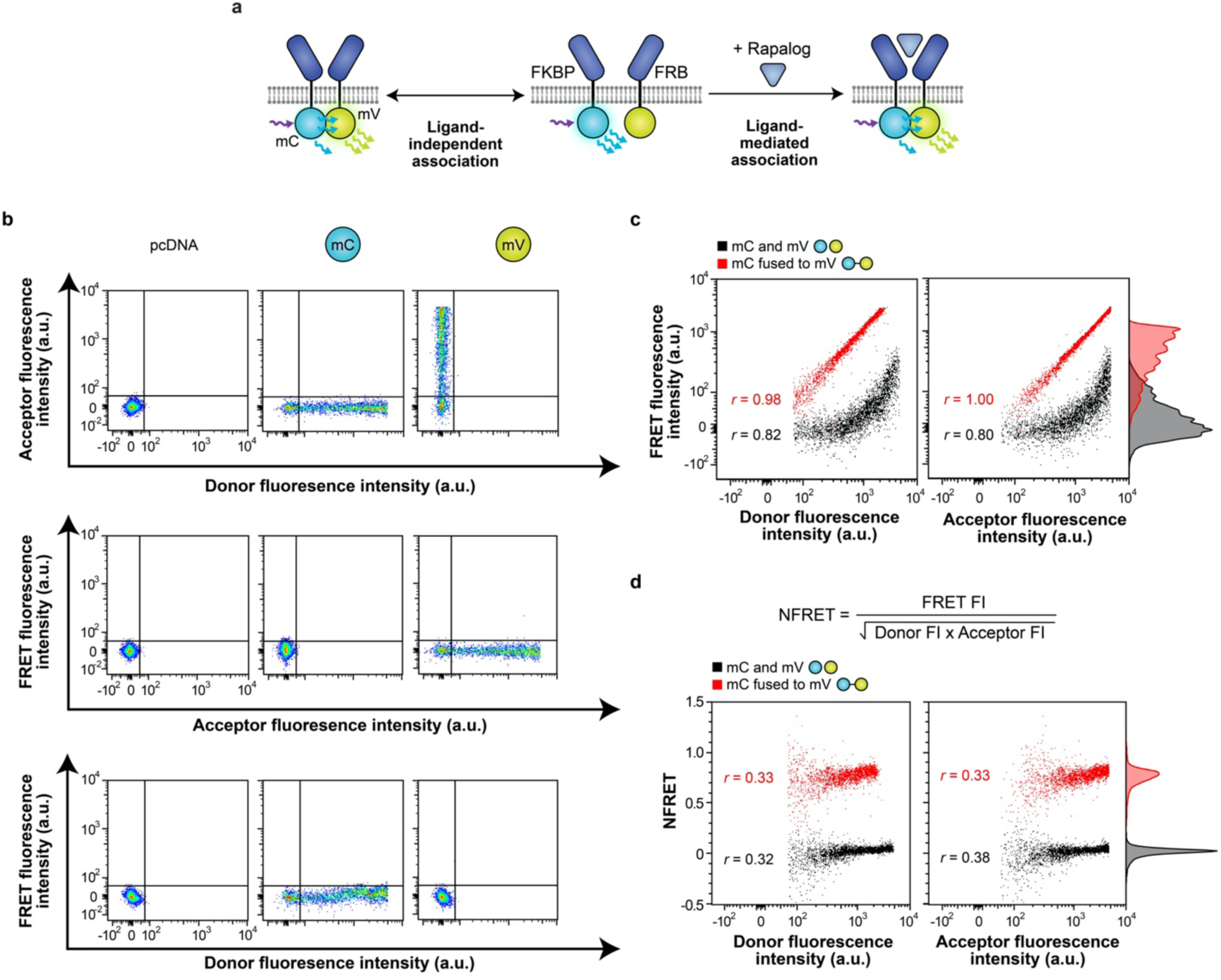
Development of a flow cytometric FRET approach to probe receptor chain association. **a** This schematic illustrates our strategy for quantifying ligand-independent (left) and ligand-mediated (right) receptor associations using Förster resonance energy transfer (FRET). Rapamycin-sensing MESA receptor ICDs were replaced with mCerulean (donor) and mVenus (acceptor) fluorophores. **b** Single-fluorophore samples were used to linearly compensate bleed-through from individual fluorophores into both the other fluorophore channel and the FRET channel. These plots also illustrate the gating used to identify cells expressing both the donor and acceptor fluorophores (mCerulean+/mVenus+). Abbreviations: mC, mCerulean; mV, mVenus. **c** Cytosolically expressed control constructs that are expected to display low FRET (separate soluble donor and acceptor proteins) or high FRET (donor-acceptor fusion protein) differ by a vertical shift in fluorescence in the FRET channel. Fluorescence in the FRET channel is linearly correlated with donor and acceptor fluorescence, respectively. The cells shown are singlets that are transfected (miRFP670+) and that express the donor and acceptor (mCerulean+/mVenus+). **d** When FRET fluorescence is normalized to donor and acceptor fluorescence intensities by the calculated NFRET metric (equation shown), the cytosolic controls still display a vertical shift in NFRET, but NFRET only has a low correlation with donor and acceptor fluorescence, respectively; NFRET is more independent of expression differences observed across the cell population (compared to FRET fluorescence intensity in **c**). The NFRET metric better distinguishes low and high FRET controls than does unprocessed FRET fluorescence. The cells shown are singlets that are transfected (miRFP670+) and that express the donor and acceptor (mCerulean+/mVenus+). Experiments were conducted in biological triplicate, and individual representative samples are shown. Adjunct histograms represent probability density and are scaled to unit area. Data were analyzed as described in **Methods**.

To validate our FRET assay, we first used a model system: cells expressed either a positive control (an mVenus-mCerulean fusion protein that is expected to exhibit strong FRET (54)) or a negative control (mVenus and mCerulean expressed as separate proteins that are expected to exhibit minimal FRET). We evaluated two possible metrics to quantify FRET efficiency. The first metric, FRET fluorescence intensity, was quantified using unprocessed fluorescence intensity in the FRET channel. As expected, this metric correlated linearly with donor fluorescence and acceptor fluorescence (proxies for the expression level of each fluorescent protein), respectively, and as expected, FRET fluorescence intensity was higher for the fusion protein than for the non-fused control (**Fig. 3c**). The second metric, normalized FRET (NFRET) (32, 55), was calculated by normalizing the FRET fluorescence intensity of each cell to that cell’s donor and acceptor fluorophore expression levels to provide a whole-cell FRET metric that accounts for fluorescent protein expression level (**Fig. 3d**). Like FRET fluorescence intensity, NFRET was higher for the fusion protein than for the non-fused control. However, NFRET demonstrated the anticipated low correlation with fluorophore expression level and enabled comparisons across two orders of magnitude in donor and acceptor fluorescence (**Fig. 3d, Supplementary Fig. 10c**). Although each of these metrics can report *whether* FRET occurs in this model system, NFRET provides better quantitative resolution separating the two control populations (**Fig. 3c,d**), and since it inherently controls for variation in protein expression, we utilized this metric exclusively going forward. As a final validation of our analytical pipeline, and to enable comparison of our methods with classic microscopy-based FRET analyses, we adopted our method to a confocal microcytometry workflow (**Methods**), which confirmed the patterns observed by flow cytometry (**Supplementary Fig. 11**).

Having validated this FRET assay, we next employed it to interrogate the contribution of TMD choice to MESA inter-chain interactions. To investigate the mechanistic questions and apparent contradictions discussed above, we selected three TMDs for investigation: the CD28-TMD was selected for its predicted large number of energetically favorable modes of association (**Supplementary Fig. 9**), propensity to cluster (50), and use in previous MESA receptors (25, 28); the GpA-TMD was selected for its documented high propensity to homodimerize (49, 50), and surprising lack of elevated background signaling (**Fig. 2c**); and the FGFR4-TMD was selected for its documented (50) and predicted low propensity to dimerize (**Supplementary Fig. 9**) and high performance in functional assays (**Fig. 2c**). We hypothesized that if TMD choice impacts receptor function *primarily* through modulating inter-chain affinity, then functional characteristics (e.g., low vs. high background signaling) would correlate with FRET trends (e.g., low vs. high basal FRET signal, respectively). We started by evaluating MESA with matched TMD pairs and found that background NFRET was slightly (but significantly) lower for the GpA-TMD and FGFR4-TMD based MESA than for the CD28-TMD based MESA, although ligand-induced increases in NFRET (i.e., fold difference) were significant and similar across all three TMD choices (**Fig. 4a**). These receptors also exhibited similar ligand dose responses (**Fig. 4b, Supplementary Fig. 12**). We next examined MESA receptors with mixed TMD pairs and again observed that constructs exhibited similar and significant ligand-induced NFRET (**Fig. 4c**). This pattern held upon swapping the fluorophore domains (**Supplementary Fig. 13**), testing different cell harvest methods (**Supplementary Fig. 14**), and measuring NFRET by confocal microscopy (**Supplementary Fig. 15**). To investigate whether TMD choice impacts association kinetics, we performed a time course assay (**Fig. 4d, Supplementary Fig. 16**), which revealed that MESA receptors containing the CD28-TMD and/or the FGFR4-TMD exhibited similar NFRET induction kineticsonset was generally observed within 15 min of ligand addition (**Fig. 4d, Supplementary Fig. 17**), indicating that this signal is attributable to the ligand-induced association of existing chains, rather than potential ligand-induced stabilization of newly synthesized chains. Altogether, these evaluations revealed no substantial TMD-dependent effects, which argues against the hypothesis that TMD choice impacts MESA function primarily through modulating inter-chain affinity.

**Fig. 4.**
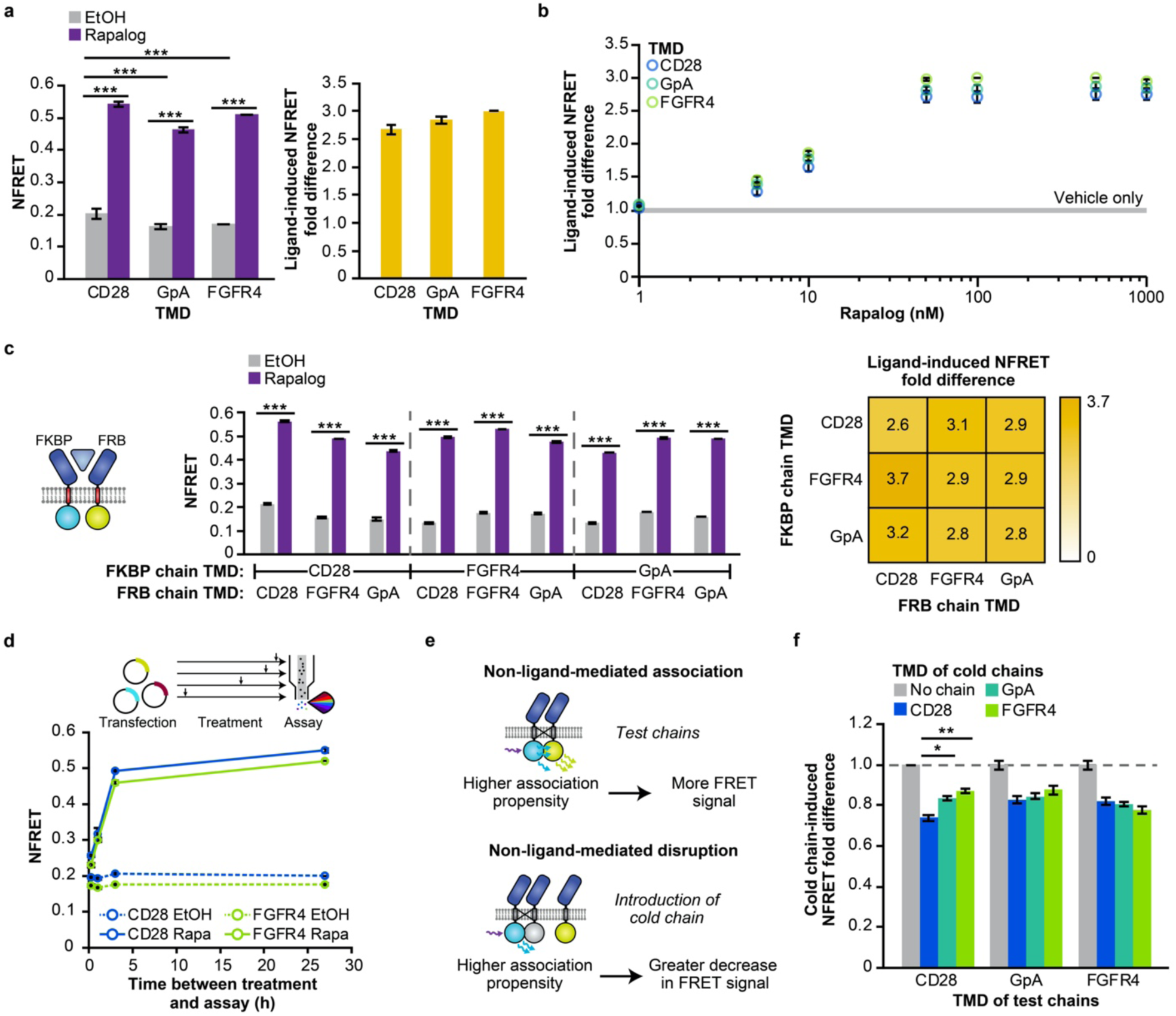
Effect of TMD choice on receptor chain association. **a** Receptor pairs with matched TMDs exhibit a ligand-induced increase in NFRET (27 h incubation, 100 nM rapalog) (two-way ANOVA, *** *p* < 0.001) (left). The CD28-TMD matched receptor pair exhibits slightly higher NFRET in the absence of ligand compared to the GpA-TMD pair and FGFR4-TMD pair (two-way ANOVA, *** *p* < 0.001). Fractional change in NFRET upon ligand treatment (ligand-induced NFRET fold difference) is comparable across TMDs (two-tailed Welch’s *t*-test, all *p* > 0.05) (right). In all cases, the donor fluorophore is on the FKBP chain and the acceptor fluorophore is on the FRB chain. **b** NFRET induction varies with rapalog dose (measurement at 27 h incubation). **c** Pairs of receptors with mixed and matched TMDs exhibit a significant ligand-induced increase in NFRET (27 h incubation, 100 nM rapalog) (three-way ANOVA, *p* < 0.001). The ligand-induced NFRET increase is comparable across mixed and matched TMD pairs. **d** Dynamics of NFRET response to ligand. By 3 h post-ligand treatment, the NFRET increase is nearly maximal (87% relative to the NFRET at 27 h). Abbreviation: Rapa, rapalog. **e**,**f** In a cold (non-fluorescent) chain competition assay with matched TMD fluorescent receptors, CD28-TMD exhibited a slightly higher propensity to associate: the NFRET decrease conferred by introduction of a CD28-TMD cold chain was greater in the CD28-TMD matched case than in other cases (two-tailed Welch’s *t*-test, all *p* > 0.05 except for comparison between CD28-mediated and GpA-mediated disruption of CD28 FRET and CD28-mediated vs. FGFR4-mediated disruption of CD28 FRET, * *p* < 0.05, ** *p* < 0.01). All chains contain the same ECD (FRB). Experiments were conducted in biological triplicate, and data were analyzed as described in **Methods**. Error bars represent the S.E.M. Outcomes from ANOVAs and Tukey’s HSD tests for **a–c** are in **Supplementary Notes 2 and 3**.

Given the surprising finding that TMD choice did not appear to impact inter-chain association, we next used a separate and potentially more sensitive FRET assay to seek confirmation of this finding. For this evaluation, we performed a cold chain competition (**Fig. 4e**), in which fluorescently tagged MESA chains (called test chains) compete with MESA chains employing a non-fluorescent mVenus mutant (called cold chains) for TMD-mediated interactions (**Supplementary Fig. 18a**). We expect that introduction of a cold chain will lead to a decrease in FRET between test chains, and that the magnitude of this decrease will trend with the affinity with which test and cold chains associate in a ligand-independent manner. As expected, we observed that the introduction of cold chains reduced NFRET between all test chains, and notably this reduction was approximately 20% in all but one case (**Fig. 4f, Supplementary Fig. 18b**): CD28-TMD–containing test chains showed a slightly but significantly greater decrease in NFRET when co-expressed with CD28-TMD cold chains rather than with any other cold chains. This result indicates that MESA with the CD28-TMD have a slightly higher association propensity than do MESA with the GpA-TMD or FGFR4-TMD, which is consistent with the previously observed slight elevation in background NFRET for MESA with the CD28-TMD, compared to MESA with other TMD choices (**Fig. 4a**). Together, these results indicate a slightly higher propensity for CD28-TMD MESA to self-associate compared to MESA with the FGFR4-TMD and GpA-TMD. Interestingly, the increased propensity for CD28-TMD MESA to self-associate did not manifest in elevated background signal compared to MESA with other TMDs, and these results are insufficient to explain the observed TMD-dependent differences observed with GpA-TMD and FGFR4-TMD MESA in functional assays (**Fig. 2**). Therefore, we conclude that TMD choice primarily affects MESA receptor performance by modulating a property other than TMD-dependent inter-chain association propensity.

### Transmembrane domain choice governs receptor dimerization geometry and *trans*-cleavage efficiency

Considering the preceding observations, we hypothesized that TMD choice might impact the geometry of receptor dimerization in a manner that affects *trans*-cleavage efficiency without affecting the propensity for the chains to associate. To test this hypothesis, we used a panel of synthetic TMDs that were designed to dimerize with varying geometries and were previously used to study geometric constraints on juxtamembrane regions of RTKs that stemmed from the TMD (56). In this panel, changes in TMD sequence are used to systematically vary the orientation of dimerization (**Fig. 5a**). TMDOCK analysis predicted that this panel of TMDs can homodimerize in configurations with distances between TMD C-termini (at the inner leaflet of the membrane) ranging from 5.4–16.0 Å and in a variety of rotational orientations (**Supplementary Fig. 19**). Notably, TMD-dependent differences in inter-chain orientation on this scale would not necessarily generate TMD-specific FRET signals. We hypothesized that if TMD interaction geometry does indeed impact *trans*-cleavage efficiency, then MESA receptors built using this panel of synthetic TMDs would exhibit differential signaling in a functional assay. We found that ligand-induced signaling indeed varied across the panel and decreased substantially as the residues mediating dimerization were moved away from the inner leaflet of the membrane (**Fig. 5b**). Notably, some synthetic TMD choices yielded very little ligand-induced signal, similar to what was observed with several native TMDs (**Fig. 2c**). The aforementioned effects of TMD choice could not be explained by differences in expression level of the receptors—the chains with the lowest magnitude of ligand-induced signaling were the most highly expressed (**Supplementary Fig. 20a**)—nor by differences in ligand-induced MESA protein accumulation, which was modest and similar across the panel (**Supplementary Fig. 20b**). Altogether, these data support the conclusion that TMD choice substantially influences the efficiency of MESA *trans*-cleavage via a mechanism that involves the intracellular geometry in which MESA chains dimerize upon ligand-binding.

**Fig. 5.**
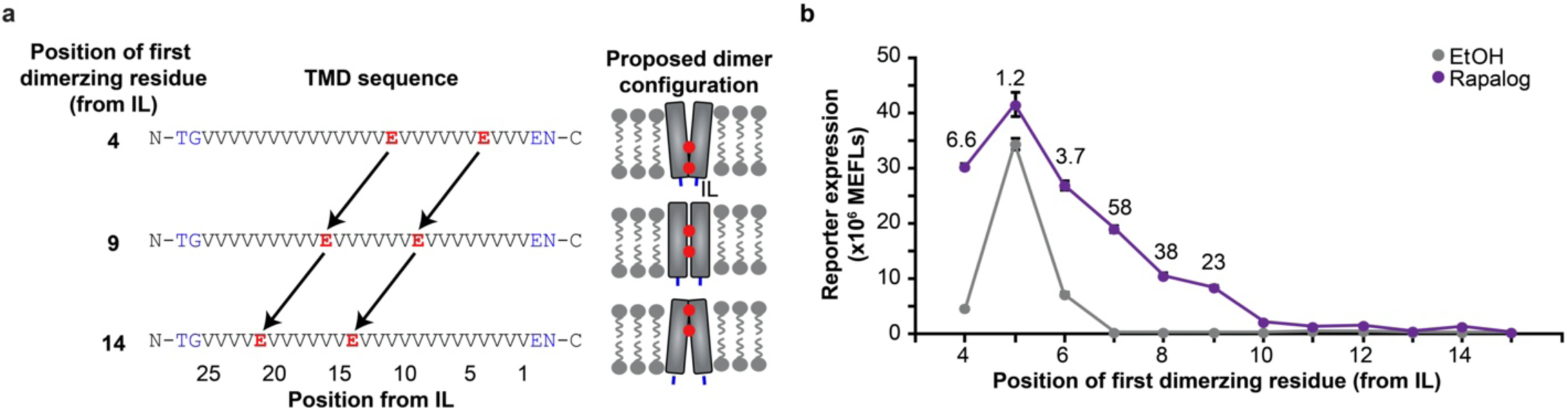
Effects of TMD dimerization geometry on receptor signaling. **a** The schematic depicts the design of synthetic TMDs used to constrain receptor dimerization geometry. Dimerization of the valine-rich alpha helices occurs at the hydrophilic glutamic acid residues. Juxtamembrane N-terminal outer linker and C-terminal inner linker (IL) residues are shown in blue. **b** Positioning the dimerizing residues at different locations in the alpha helix conferred highly varied effects on background and ligand-induced signaling. Moving the position of the first dimerizing residue from the IL from position 4 to 5, 5 to 6, 6 to 7, 7 to 8, and 9 to 10 resulted in a significant difference in ligand-induced reporter expression (two-way ANOVA, *p* < 0.01). Fold difference values are shown above points where ligand addition induced reporter expression that was significantly above background. Experiments were conducted in biological triplicate, and data were analyzed as described in **Methods**. Error bars represent the S.E.M. Outcomes from ANOVAs and Tukey’s HSD tests for **b** are in **Supplementary Note 2**.

### Design rules extend to the development of new synthetic receptors

Given our new understanding of the role of TMD choice, we next investigated whether the observed trends would extend to MESA systems with different ECDs and ligands. To test this hypothesis, we first built two new MESA receptors to sense small molecules—gibberellin (GA3-AM is a cell-permeable analog) and abscisic acid (ABA) (**Supplementary Fig. 21–26**)—and a third new MESA receptor to sense green fluorescent protein (GFP; here we use co-expressed secreted GFP, sGFP, as an expedient testing system), and explored design considerations that are typically expected to be ECD-specific, such as how linker length affects expression and cell-surface localization (57, 58). Interestingly, functional assays revealed TMD-associated trends that were largely consistent across ECDs (**Fig. 6a–d, Supplementary Fig. 22, 24, 27, 28**). The TMD choice for each chain significantly affected background signaling and induced signaling, and the interaction between TMD choices was also significant, indicating that the choice of TC TMD or PC TMD alone does not fully explain the trends (**Supplementary Note 3**). Additionally, the TMD choices together account for most of the variance in background and induced signaling observed (**Supplementary Note 2**). These observations indicate that satisfying any one design objective (e.g., maximize fold difference, minimize background, etc.) requires choosing a pair of TMDs suited to that design goal. Additionally, some general trends held across the rapamycin MESA receptors and these three new receptors, suggesting that these might represent generalizable principles guiding the design of future MESA receptors. For example, high background and modest induced signaling were observed for pairs including FGFR1-TMD TC, resulting in generally low fold difference. Conversely, FGFR4-TMD–containing pairs often exhibited low background signaling and high fold difference (**Fig. 6a–d**). Overall, we conclude that some effects of TMD choice extend across multiple MESA receptors, and that a limited experimental evaluation of these few TMD choices enables one to generate multiple new high-performing receptors.

**Fig. 6.**
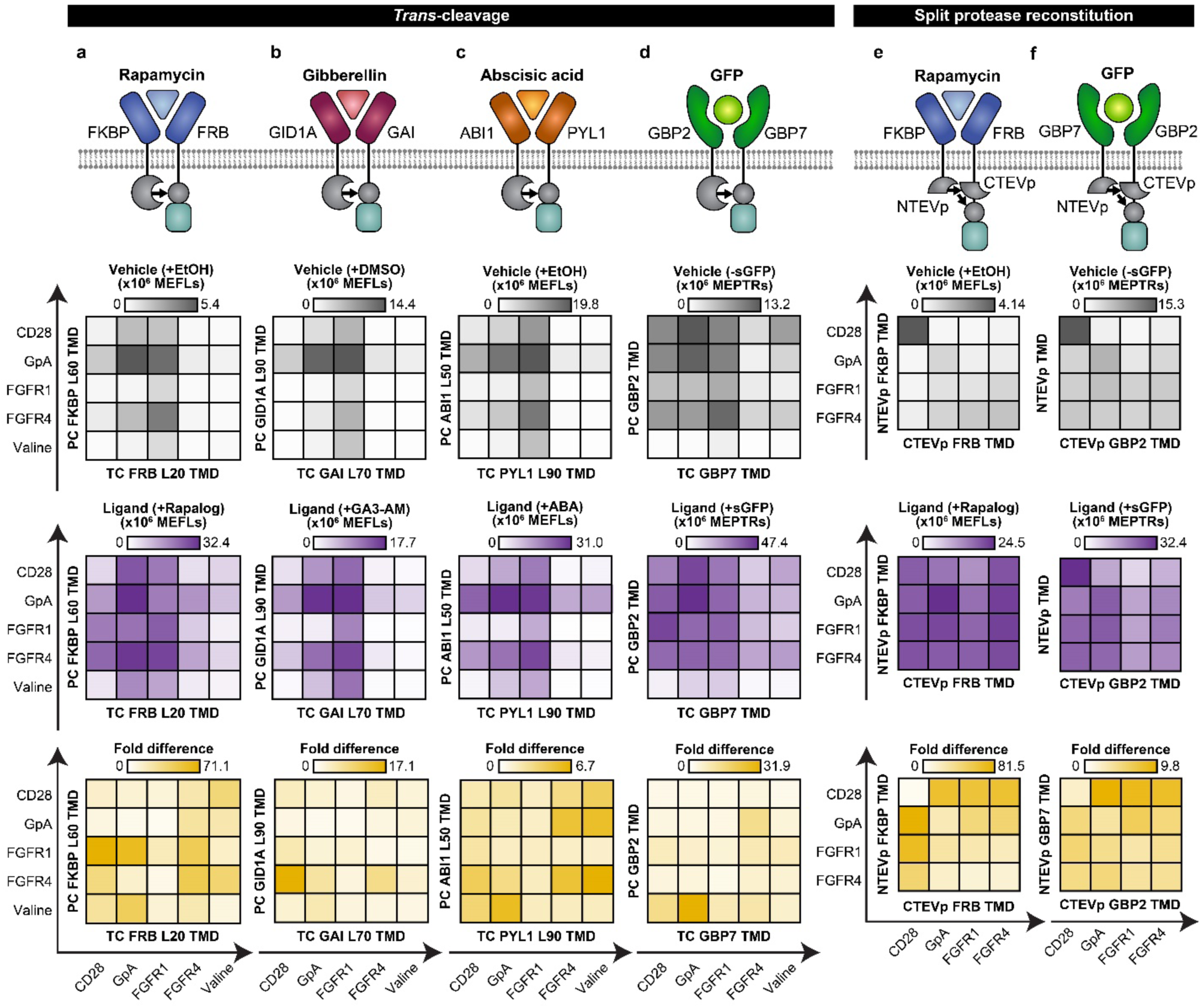
Tuning an expanded panel of MESA receptor systems. **a–d** Functional assays for MESA receptors for sensing rapamycin, gibberellin, abscisic acid, and sGFP were constructed using the full TEVp-based *trans*-cleavage mechanism. Axes shown on the perimeter of the heatmaps in **a–d** apply to all heatmaps in **a–d. e**,**f** Functional assays for MESA receptors for sensing rapamycin and sGFP were constructed using a revised mechanism, including previously reported H75S/L190K mutations for tuning split TEVp reconstitution propensity (26). ECDs and extracellular linker lengths are unique to each set of ligand-binding domains (**Supplementary Figs. 21, 23**). Axes shown on the perimeter of the heatmaps in **e** and **f** apply to all heatmaps in **e** and **f**. Heatmaps display the mean from three biological replicates of reporter expression with vehicle only (gray), reporter expression with ligand (purple), and ligand-induced fold difference (gold). Within each system, a consistent plasmid dose was used across conditions. Each panel (column) is an independent experiment, and each heatmap is internally scaled by the system. Corresponding bar graphs are in **Supplementary Figs. 22, 24, 27–30**). Data were analyzed as described in **Methods**. Outcomes from ANOVAs and Tukey’s HSD tests are in **Supplementary Notes 2 and 3**.

We next investigated whether the observed effects of TMD choice might extend to synthetic receptors that operate by a distinct mechanism. To test this, we employed a recently reported MESA system in which ligand binding induces reconstitution of a mutant split TEVp (26). The reconstituted protease then releases a TF from the receptor, enabling the TF to enter the nucleus and drive reporter expression (**Fig. 6e**). We replaced the CD28-TMD used in the reported rapamycin-sensing base case (26) with other TMDs and also built GFP-sensing variants of these receptors. In functional assays, we again found that the TMD proved to be a useful handle for tuning performance (**Fig. 6e,f, Supplementary Fig. 29, 30**). Background and ligand-induced signaling were again significantly affected by TMD choice for each chain and the interaction between the TMD choices (**Supplementary Note 3**). Strikingly, across all mixed and matched TMD pairs, the ligand-induced signal was relatively consistent, which differs from the trends observed for the *trans*-cleavage mechanism. Moreover, for these receptors, using the CD28-TMD on only one chain resulted in very low background signaling, but using the CD28-TMD on both chains (as previously reported (26)) resulted in the highest background of all combinations tested, with the interaction between TMD choices accounting for most of the variance in background signaling (**Supplementary Note 2**). When considered together with the FRET experiments (**Fig. 4a,f**), these results again suggest that CD28-TMD MESA exhibits increased ligand-independent association propensity compared to MESA employing other TMDs. This effect can be problematic if both chains include this TMD, yet this same property can be useful if only one chain bears this TMD. Comparing these two distinct MESA receptor mechanisms suggests that whereas subtle geometric effects conferred by specific TMD pairs have a substantial impact on MESA receptors employing the original *trans*-cleavage mechanism (**Fig. 6a–d**), these nuances can have less impact on the performance of receptors employing the distinct split-TEVp reconstitution mechanism (**Fig. 6e,f**). The observation that the effects conferred by TMD choice are dependent on signaling mechanism is also consistent with our hypothesis that TMD choice is most important for determining the geometry of dimerized receptors. If, conversely, the main effect of TMD choice were modulating inter-chain association propensity, then we might expect TMD choice to yield similar effects when used with either mechanism, yet we see this consistency only in CD28-TMD-mediated elevation of background signaling. A final important finding is that for both MESA receptor mechanisms examined, our initial model systems generated insights enabling the design of novel receptors.

Finally, given these new insights into the important connection between MESA receptor geometry and functional performance, we sought to utilize these tools to guide the selection of new ligands and ligand-binding domains. For MESA receptors engineered to sense rapamycin, gibberellin, abscisic acid, or in previous work, vascular endothelial growth factor (VEGF) (5), there exist crystal structures enabling us to estimate the displacement between the C-termini of the ECDs when they are dimerized in the ligand-bound state (34-37). To determine whether this spatial variation is sufficient to impact intracellular receptor geometry, we again employed FRET analysis (employing the well-behaved FGFR4-TMD) to best decouple our investigation of this question from the effects of receptor geometry on *trans*-cleavage (**Fig. 7a, Supplementary Fig. 31**). Interestingly, ligand-induced NFRET fold difference showed a strong negative correlation with the ECD C-terminal distance, suggesting that one can predict some aspects of receptor structure and function from prior knowledge of the ECDs alone—an attractive feature for a modular design strategy. We confirmed that these effects were indeed attributable to receptor complex formation by performing a ligand-ECD orthogonality analysis (**Fig. 7b**). Notably, the VEGF-binding ECDs—which correspond to the largest ECD C-terminal distances evaluated—showed a *diminishment* in NFRET upon ligand treatment in both experiments, suggesting that ligand-induced dimerization might diminish, on average, transient inter-chain ICD interactions compared to the ligand-free state. For a given TMD choice, the ligand-induced fold difference values exhibit a positive correlation (in rank order of effects) between NFRET and reporter expression. Comparing all FGFR4-TMD receptors used in both FRET and functional assays, the highest values for both metrics were observed for the rapamycin-sensing system, with lower but similar values observed for the gibberellin and abscisic acid-sensing systems (**Fig. 6a–c, Fig. 7a**). These observations could help to explain why MESA receptors with shorter ECD C-terminal distances (e.g., for rapamycin-sensing) generally exhibit strong signaling. They also suggest that functional sensing of gibberellin, abscisic acid, and VEGF (including receptor chain *trans*-cleavage) can be attributed to inter-chain interactions that occur with lower frequency than those that are required to confer FRET. Altogether, the analyses presented in this study provide powerful new insights into the connections among tunable protein design choices, biophysical consequences, and impacts on the performance of synthetic receptors (**Fig. 7c**).

**Fig. 7.**
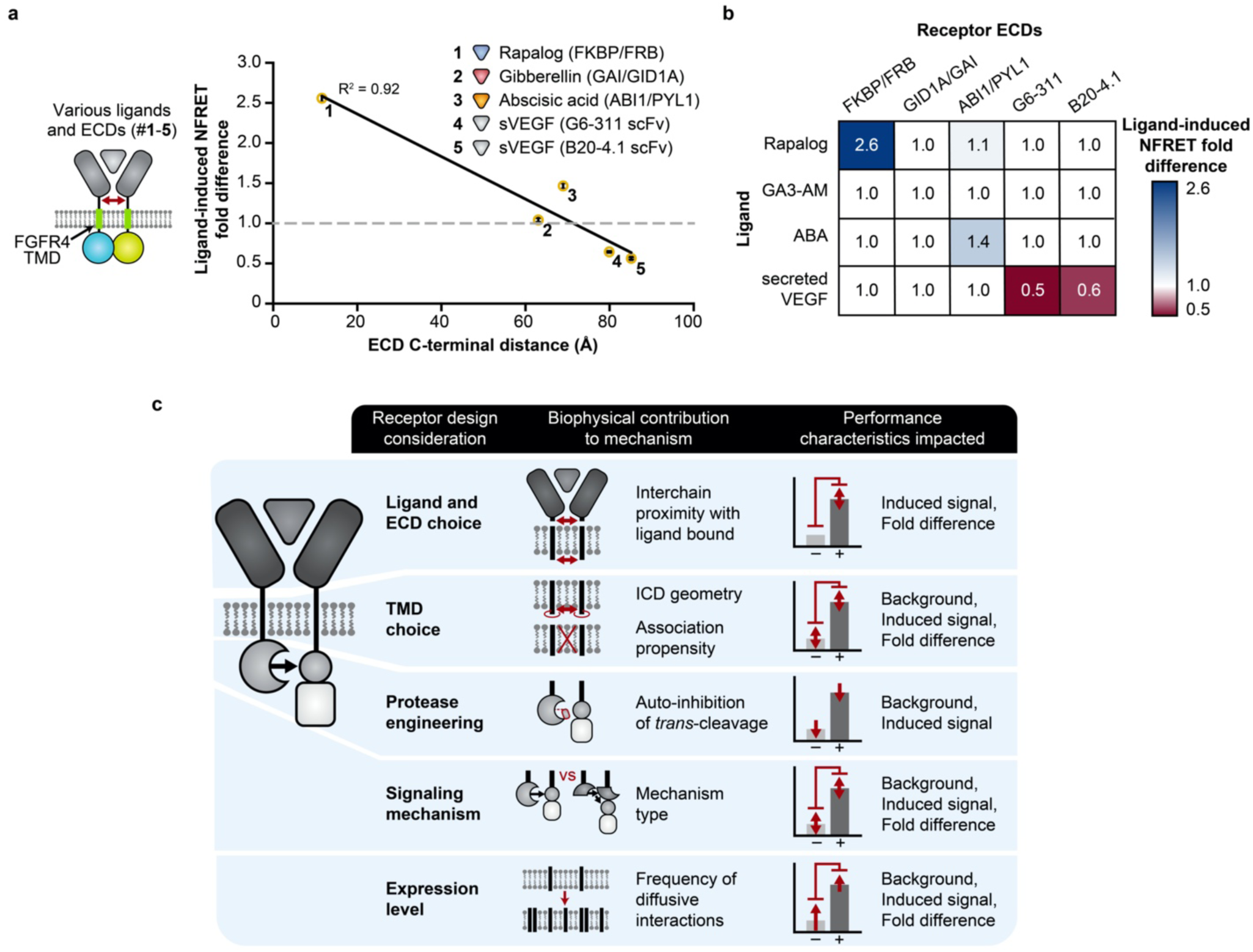
Generalizing principles for receptor engineering. **a** Ligand-induced inter-chain association (NFRET fold difference) varies with ECD C-terminal distance, with a negative linear relationship (*y* = – 0.027*x* + 2.9, where *y* is NFRET fold difference and *x* is C-terminal distance in Å, R^2^ = 0.92, two-tailed Student’s *t*-test, *p* = 0.01). The ligands are rapalog, GA3-AM, ABA, and secreted VEGF. The red arrow indicates the ECD C-terminal distance—the spatial displacement between C-termini of the ligand-binding domains. **b** Validation of receptor-ligand orthogonality: substantial fold differences in NFRET were observed only when each ligand was paired with its cognate ligand-binding domains. Experiments were conducted in biological triplicate, and data were analyzed as described in **Methods**. Error bars represent the S.E.M. **c** This schematic synthesizes the findings of this investigation with prior MESA receptor development (26, 28) by relating receptor design choices to the proposed biophysical consequences and performance characteristics affected.

## DISCUSSION

A key finding of this study is that systematic re-evaluation of the MESA synthetic receptor system enabled identification of design modifications that substantially improve receptor performance. While some modifications had modest effects, TMD substitution proved to be a particularly useful handle for receptor tuning and optimization (**Fig. 2**). Some TMD combinations greatly decreased background signaling while maintaining or increasing ligand-induced signaling, yielding high-performing receptors with fold differences on the order of 100. By this metric, these rank among the best synthetic receptors reported to date (12, 18, 23). Moreover, evaluating a relatively small library of TMD variants enabled us to generate high-performing MESA receptors for three new ligands (**Fig. 6**). It is notable that even for the fully synthetic mechanism employed by MESA receptors, it was not clear ahead of time which design modification strategies would be most fruitful. These improvements were achieved largely by investigating MESA receptor biophysics using the approaches and strategies typically employed to study natural receptors. There likely exists substantial room for improving synthetic receptor systems—a recent report evidences the utility of applying this approach to improve synNotch (24)—and powerful tools developed by the receptor biophysics community comprise substantial yet underutilized potential.

An integral part of this investigation was the use of flow cytometric FRET. This technique enabled us to link disparate observations and build understanding (**Fig. 4**). FRET has generally proven useful for elucidating native receptor mechanisms, and these experiments typically employ confocal microscopy (59-61). We employed a substantially more high-throughput assay that provides single-cell resolution (thousands of individual cells per sample) by adapting a reported flow cytometry workflow (51) to this investigation (**Fig. 3**). A key utility of the NFRET metric is that it enables comparisons across a heterogeneous population of cells by expression-normalizing FRET. This feature of the NFRET metric makes it useful for evaluating receptor dimerization when receptors contain new protein domain choices (such as ECD), as it decouples the effects of these new domains on expression level from their effects on dimerization (with or without ligand). To our knowledge, this is the first application of flow-FRET to characterize synthetic receptors, and this approach may be useful for studying and optimizing systems beyond MESA. A consideration for future use is that our assay used the mCerulean/mVenus fluorescent protein pair and standard flow cytometry filters, which sufficed for our analysis, but it is possible that FRET signal could be increased by selecting alternative fluorophores and/or bespoke filters.

Altogether, this investigation provides new mechanistic insights into how TMD choice impacts MESA receptor performance. Only the CD28-TMD used in previous MESA receptors (25, 26, 28) showed some modest propensity to cluster in the absence of ligand (**Fig. 4a,f**), which is consistent with roles played by CD28 in T cell signaling at the immunological synapse (45). In general, TMD choice conferred little effect (CD28-TMD) or no effect (all other tested TMDs) on the propensity of MESA chains to associate in the presence or absence of ligand. Thus, there must exist an alternative explanation as to why TMD choice profoundly impacts background signaling, ligand-induced signaling, and even PRS cleavability. We propose that TMDs primarily contribute to receptor performance by influencing the geometry in which intracellular domains interact. Several lines of evidence support this hypothesis. Systematically varying the geometry with which TMDs associate either facilitates or constrains MESA receptor signaling (**Fig. 5**), and some TMDs can render a TC resistant to cleavage by a PC (**Fig. 2c, Fig. 5b, Supplementary Fig. 8**). TMD-associated trends also held to some extent across various choices of ligand and ligand-binding domains (**Fig. 6a–d**). Although it is not yet clear *why* each TMD pair impacts the geometry with which intracellular domains associate, experimentally screening the TMD library reported here enabled the improvement of existing MESA receptors based upon each of the two distinct mechanisms (**Fig. 6**) (25, 26, 28). It remains to be seen whether further TMD screening or engineering may represent an opportunity for further enhancement of MESA receptors, and whether TMD substitution will improve other synthetic receptors.

Ultimately, we hope that this investigation will serve as a guide for building and tuning synthetic receptors by connecting design choices to biophysical consequences and performance characteristics of interest (**Fig. 7c**). When designing new MESA receptors, some choices must be made at the outset. First, the geometry with which ligand-binding domains are separated in physical space when bound to ligand fundamentally limits inter-chain interactions (**Fig. 7a**,**b**), so minimizing the apposition of these domains may benefit MESA receptor performance (**Figs. 6,7**). Since the magnitude of NFRET induction upon ligand addition roughly correlates with the fold difference in ligand-induced signaling (**Fig. 6a–c, Fig. 7a**), our FRET assay may be a useful tool for selecting candidate ligand-binding domains when a crystal structure cannot be used as a guide; since FRET phenomena depend weakly on TMD choice, such an initial evaluation need not include a combinatorial TMD screen. Second, since TMD choice substantially influences MESA performance, we recommend that functional evaluations of new receptors include a limited combinatorial sampling of TMDs (e.g., using a panel based upon those included in **Fig. 6**). Finally, expression tuning also provides a fairly facile handle for optimizing (empirically, if needed) specific receptor performance characteristics, although this might be less important when using MESA that employ the split TEVp mechanism, which is less sensitive to receptor expression level (26). The methods and insights developed here should facilitate the construction of novel, high-performing receptors for diverse ligands, including both MESA receptors and potentially synthetic receptor systems in general.

## Supporting information

Supplementary Software

## DATA AVAILABILITY

Plasmid maps are provided as annotated GenBank files in **Supplementary Data 2**. Plasmids and complete annotated GenBank files will be deposited with Addgene. This study uses Addgene plasmids #58855, #138749 (described in **Materials and Methods**). All reported experimental data will be provided in **Source Data 1 upon final submission in a digital repository**.

## ACKNOWLEDGEMENTS

The authors would like to thank Stephen Lander and Simrita Deol for assistance with some of the experiments and data analysis, Cameron McDonald for assistance with the PC mutagenesis study, the Neha Kamat lab for use of an Azure c280 imager, and members of the Leonard lab for useful discussions.

## AUTHOR CONTRIBUTIONS

H.I.E., P.S.D., J.J.M., T.B.D., and J.N.L. conceived and designed this study. H.I.E., P.S.D., J.J.M., A.K.K., T.B.D., L.M.B., E.R.A, and M.H. generated reagents. H.I.E., P.S.D., J.J.M., A.K.K., T.B.D., L.M.B., and E.R.A. designed and performed experiments and analyzed the data. H.I.E., P.S.D., J.J.M., and J.N.L. wrote the manuscript, and all authors edited and approved the final manuscript.

## REFERENCES

1. Muldoon, J.J., Donahue, P.S., Dolberg, T.B., Leonard, J.N. (2017) Building with intent: technologies and principles for engineering mammalian cell-based therapies to sense and respond. Curr Opin Biomed Eng, 4, 127–133.

2. Guedan, S., Ruella, M., June, C.H. (2019) Emerging Cellular Therapies for Cancer. Annu Rev Immunol, 37, 145–171.

3. Roybal, K.T., Lim, W.A. (2017) Synthetic Immunology: Hacking Immune Cells to Expand Their Therapeutic Capabilities. Annu Rev Immunol, 35, 229–253.

4. Kojima, R., Aubel, D., Fussenegger, M. (2020) Building sophisticated sensors of extracellular cues that enable mammalian cells to work as “doctors” in the body. Cell Mol Life Sci.

5. Schwarz, K.A., Leonard, J.N. (2016) Engineering cell-based therapies to interface robustly with host physiology. Adv Drug Deliv Rev, 105, 55–65.

6. Chen, L.C., Chen, Y.Y. (2019) Outsmarting and outmuscling cancer cells with synthetic and systems immunology. Curr Opin Biotechnol, 60, 111–118.

7. Ishizuka, S. et al. (2018) Designing Motif-Engineered Receptors To Elucidate Signaling Molecules Important for Proliferation of Hematopoietic Stem Cells. ACS Synth Biol, 7, 1709–1714.

8. Engelowski, E. et al. (2018) Synthetic cytokine receptors transmit biological signals using artificial ligands. Nat Commun, 9, 2034.

9. Qudrat, A., Truong, K. (2017) Engineering Synthetic Proteins to Generate Ca(2+) Signals in Mammalian Cells. ACS Synth Biol, 6, 582–590.

10. Conklin, B.R. et al. (2008) Engineering GPCR signaling pathways with RASSLs. Nat Methods, 5, 673–678.

11. Kojima, R., Scheller, L., Fussenegger, M. (2018) Nonimmune cells equipped with T-cell-receptor-like signaling for cancer cell ablation. Nat Chem Biol, 14, 42–49.

12. Scheller, L., Strittmatter, T., Fuchs, D., Bojar, D., Fussenegger, M. (2018) Generalized extracellular molecule sensor platform for programming cellular behavior. Nat Chem Biol, 14, 723–729.

13. Qudrat, A., Mosabbir, A.A., Truong, K. (2017) Engineered Proteins Program Mammalian Cells to Target Inflammatory Disease Sites. Cell Chem Biol, 24, 703–711 e702.

14. Qudrat, A., Truong, K. (2017) Autonomous Cell Migration to CSF1 Sources via a Synthetic Protein-Based System. ACS Synth Biol, 6, 1563–1571.

15. Chung, H.K. et al. (2019) A compact synthetic pathway rewires cancer signaling to therapeutic effector release. Science, 364.

16. Barnea, G. et al. (2008) The genetic design of signaling cascades to record receptor activation. Proc Natl Acad Sci U S A, 105, 64–69.

17. Kipniss, N.H. et al. (2017) Engineering cell sensing and responses using a GPCR-coupled CRISPR-Cas system. Nat Commun, 8, 2212.

18. Maze, A., Benenson, Y. (2020) Artificial signaling in mammalian cells enabled by prokaryotic two-component system. Nat Chem Biol, 16, 179–187.

19. Baeumler, T.A., Ahmed, A.A., Fulga, T.A. (2017) Engineering Synthetic Signaling Pathways with Programmable dCas9-Based Chimeric Receptors. Cell Rep, 20, 2639–2653.

20. Krawczyk, K., Scheller, L., Kim, H., Fussenegger, M. (2020) Rewiring of endogenous signaling pathways to genomic targets for therapeutic cell reprogramming. Nature Communications, 11.

21. Morsut, L. et al. (2016) Engineering Customized Cell Sensing and Response Behaviors Using Synthetic Notch Receptors. Cell, 164, 780–791.

22. Roybal, K.T. et al. (2016) Precision Tumor Recognition by T Cells With Combinatorial Antigen-Sensing Circuits. Cell, 164, 770–779.

23. Roybal, K.T. et al. (2016) Engineering T Cells with Customized Therapeutic Response Programs Using Synthetic Notch Receptors. Cell, 167, 419–432 e416.

24. Yang, Z., Yu, Z., Cai, Y., Du, R., Cai, L. (2020) Engineering of an enhanced synthetic Notch receptor by reducing ligand-independent activation. Communications Biology, 3, 116.

25. Daringer, N.M., Dudek, R.M., Schwarz, K.A., Leonard, J.N. (2014) Modular extracellular sensor architecture for engineering mammalian cell-based devices. ACS Synth Biol, 3, 892–902.

26. Dolberg, T.B. et al. (2019) Computation-guided optimization of split protein systems. bioRxiv.

27. Hartfield, R.M., Schwarz, K.A., Muldoon, J.J., Bagheri, N., Leonard, J.N. (2017) Multiplexing Engineered Receptors for Multiparametric Evaluation of Environmental Ligands. ACS Synth Biol, 6, 2042–2055.

28. Schwarz, K.A., Daringer, N.M., Dolberg, T.B., Leonard, J.N. (2017) Rewiring human cellular input-output using modular extracellular sensors. Nat Chem Biol, 13, 202–209.

29. Brenner, M., Cho, J.H., Wong, W.W. (2017) Synthetic biology: Sensing with modular receptors. Nat Chem Biol, 13, 131–132.

30. Tang, J.C. et al. (2013) A nanobody-based system using fluorescent proteins as scaffolds for cell-specific gene manipulation. Cell, 154, 928–939.

31. Donahue, P.S. et al. (2020) The COMET toolkit for composing customizable genetic programs in mammalian cells. Nat Commun, 11, 779.

32. Xia, Z., Liu, Y. (2001) Reliable and global measurement of fluorescence resonance energy transfer using fluorescence microscopes. Biophys J, 81, 2395–2402.

33. Zal, T., Gascoigne, N.R. (2004) Photobleaching-corrected FRET efficiency imaging of live cells. Biophys J, 86, 3923–3939.

34. Liang, J., Choi, J., Clardy, J. (1999) Refined structure of the FKBP12-rapamycin-FRB ternary complex at 2.2 A resolution. Acta Crystallogr D Biol Crystallogr, 55, 736–744.

35. Murase, K., Hirano, Y., Sun, T.P., Hakoshima, T. (2008) Gibberellin-induced DELLA recognition by the gibberellin receptor GID1. Nature, 456, 459–463.

36. Yin, P. et al. (2009) Structural insights into the mechanism of abscisic acid signaling by PYL proteins. Nat Struct Mol Biol, 16, 1230–1236.

37. Fuh, G. et al. (2006) Structure-function studies of two synthetic anti-vascular endothelial growth factor Fabs and comparison with the Avastin Fab. J Biol Chem, 281, 6625–6631.

38. Dougherty, W., Cary, S.M., Parks, T.D. (1989) Molecular genetic analysis of a plant virus polyprotein cleavage sequence: a model. Virology, 171, 356–364.

39. Kapust, R.B., Tozser, J., Copeland, T.D., Waugh, D.S. (2002) The P1’ specificity of tobacco etch virus protease. Biochem Biophys Res Commun, 294, 949–955.

40. Nunn, C.M. et al. (2005) Crystal structure of tobacco etch virus protease shows the protein C terminus bound within the active site. J Mol Biol, 350, 145–155.

41. Kapust, R.B. et al. (2001) Tobacco etch virus protease: mechanism of autolysis and rational design of stable mutants with wild-type catalytic proficiency. Protein Eng, 14, 993–1000.

42. Alabanza, L. et al. (2017) Function of Novel Anti-CD19 Chimeric Antigen Receptors with Human Variable Regions Is Affected by Hinge and Transmembrane Domains. Mol Ther, 25, 2452–2465.

43. Bridgeman, J.S. et al. (2010) The optimal antigen response of chimeric antigen receptors harboring the CD3zeta transmembrane domain is dependent upon incorporation of the receptor into the endogenous TCR/CD3 complex. J Immunol, 184, 6938–6949.

44. Biggs, M.J., Milone, M.C., Santos, L.C., Gondarenko, A., Wind, S.J. (2011) High-resolution imaging of the immunological synapse and T-cell receptor microclustering through microfabricated substrates. J R Soc Interface, 8, 1462–1471.

45. Yokosuka, T., Saito, T. (2009) Dynamic regulation of T-cell costimulation through TCR-CD28 microclusters. Immunol Rev, 229, 27–40.

46. Westerfield, J.M., Barrera, F.N. (2020) Membrane receptor activation mechanisms and transmembrane peptide tools to elucidate them. J Biol Chem, 295, 1792–1814.

47. Chen, L.I., Webster, M.K., Meyer, A.N., Donoghue, D.J. (1997) Transmembrane domain sequence requirements for activation of the p185c-neu receptor tyrosine kinase. J Cell Biol, 137, 619–631.

48. Lomize, A.L., Pogozheva, I.D. (2017) TMDOCK: An Energy-Based Method for Modeling alpha-Helical Dimers in Membranes. J Mol Biol, 429, 390–398.

49. Doura, A.K., Fleming, K.G. (2004) Complex interactions at the helix-helix interface stabilize the glycophorin A transmembrane dimer. J Mol Biol, 343, 1487–1497.

50. Finger, C., Escher, C., Schneider, D. (2009) The single transmembrane domains of human receptor tyrosine kinases encode self-interactions. Sci Signal, 2, ra56.

51. Banning, C. et al. (2010) A flow cytometry-based FRET assay to identify and analyse protein-protein interactions in living cells. PLoS One, 5, e9344.

52. Goedhart, J. et al. (2010) Bright cyan fluorescent protein variants identified by fluorescence lifetime screening. Nat Methods, 7, 137–139.

53. Day, R.N., Davidson, M.W. (2009) The fluorescent protein palette: tools for cellular imaging. Chem Soc Rev, 38, 2887–2921.

54. Koushik, S.V., Chen, H., Thaler, C., Puhl, H.L., 3rd, Vogel, S.S. (2006) Cerulean, Venus, and VenusY67C FRET reference standards. Biophys J, 91, L99–L101.

55. Berney, C., Danuser, G. (2003) FRET or no FRET: a quantitative comparison. Biophys J, 84, 3992–4010.

56. Bell, C.A. et al. (2000) Rotational coupling of the transmembrane and kinase domains of the Neu receptor tyrosine kinase. Mol Biol Cell, 11, 3589–3599.

57. Kirchhofer, A. et al. (2010) Modulation of protein properties in living cells using nanobodies. Nat Struct Mol Biol, 17, 133–138.

58. Gao, Y. et al. (2016) Complex transcriptional modulation with orthogonal and inducible dCas9 regulators. Nat Methods, 13, 1043–1049.

59. Algar, W.R., Hildebrandt, N., Vogel, S.S., Medintz, I.L. (2019) FRET as a biomolecular research tool - understanding its potential while avoiding pitfalls. Nat Methods, 16, 815–829.

60. Chan, F.K. et al. (2000) A domain in TNF receptors that mediates ligand-independent receptor assembly and signaling. Science, 288, 2351–2354.

61. Hochreiter, B., Garcia, A.P., Schmid, J.A. (2015) Fluorescent proteins as genetically encoded FRET biosensors in life sciences. Sensors (Basel), 15, 26281–26314.

