## Supplementary Software for "Elucidation and refinement of synthetic receptor mechanisms": FRET_SE_Correct.m

```
%Edelstein HI, Donahue PS, Muldoon JJ, Kang AK, Dolberg TB, Battaglia LM,
%Allchin ER, Hong M, Leonard JN.
%Elucidation and refinement of synthetic receptor mechanisms.

function [pair_etoh_bulkNFRET, pair_etoh_bulkNFRET_SEM ...
    pair_rapa_bulkNFRET, pair_rapa_bulkNFRET_SEM] = ...
    FRET_SE_Correct

% Experimental and filename setup

% In the provided images, files are labeled as follows:
% [sample letter]_[field of view]_[channel]
% where sample letter = A, B, C, D, or E;
%       field of view = 1, 2, 3, 4, 5, 6, 7, 8, 9, or 10;
%       channel = DD (donor ex and em), AA (acceptor ex and em),
%       or DA (donor excitation and acceptor em).

% Specify number of images (fields of view) collected for each transfection
% (receptor pair treated with EtOH, receptor pair treated with rapalog,
% receptor tagged with donor alone, receptor tagged with acceptor alone,
% vector only).
n_pair_etoh    = 10;
n_pair_rapa    = 10;
n_donoronly    = 10;
n_acceptoronly = 10;
n_vectoronly   = 10;

% Specify sample filename labeling
vector    = 'A';
donor     = 'B';
acceptor  = 'C';
pair_rapa = 'D';
pair_etoh = 'E';

% Specify folder name for saving processed plots
sampleFolder = 'NFRET_images';
folderName = strcat(pwd, '/', sampleFolder);
if ~exist(folderName, 'dir')
    mkdir 'NFRET_images';
end
folderName = [folderName, '/'];


% Image processing
%**** Part 1: Load fields of view (images) into matrices.
% Each set of images in a channel are loaded into a 512x512xn matrix,
% where n is the corresponding number for each sample.
% In this example, each sample (A-E) will produce three
% 512x512x10 matrices (one for DD, AA, DA channels).

disp('Reading acceptor chain images into matrices');
[acceptor_DD, acceptor_DA, acceptor_AA] = deal(zeros(512, 512, n_acceptoronly));
for i = 1:n_acceptoronly
    acceptor_DD(:,:,i) = double(imread(strcat(acceptor, '_', int2str(i), '_DD.tif')));
    acceptor_DA(:,:,i) = double(imread(strcat(acceptor, '_', int2str(i), '_DA.tif')));
    acceptor_AA(:,:,i) = double(imread(strcat(acceptor, '_', int2str(i), '_AA.tif')));
end

disp('Reading donor chain images into matrices');
[donor_DD, donor_DA, donor_AA] = deal(zeros(512, 512, n_donoronly));
for i = 1:n_donoronly
    donor_DD(:,:,i) = double(imread(strcat(donor, '_', int2str(i), '_DD.tif')));
    donor_DA(:,:,i) = double(imread(strcat(donor, '_', int2str(i), '_DA.tif')));
    donor_AA(:,:,i) = double(imread(strcat(donor, '_', int2str(i), '_AA.tif')));
end

disp('Reading vector control images into matrices');
[vector_DD, vector_DA, vector_AA] = deal(zeros(512, 512, n_vectoronly));
for i = 1:n_vectoronly
    vector_DD(:,:,i) = double(imread(strcat(vector, '_', int2str(i), '_DD.tif')));
    vector_DA(:,:,i) = double(imread(strcat(vector, '_', int2str(i), '_DA.tif')));
    vector_AA(:,:,i) = double(imread(strcat(vector, '_', int2str(i), '_AA.tif')));
end

disp('Reading paired chain images into matrices');
[pair_etoh_DD, pair_etoh_DA, pair_etoh_AA] = deal(zeros(512, 512, n_pair_etoh));
for i = 1:n_pair_etoh
    pair_etoh_DD(:,:,i) = double(imread(strcat(pair_etoh, '_', int2str(i), '_DD.tif')));
    pair_etoh_DA(:,:,i) = double(imread(strcat(pair_etoh, '_', int2str(i), '_DA.tif')));
    pair_etoh_AA(:,:,i) = double(imread(strcat(pair_etoh, '_', int2str(i), '_AA.tif')));
end
[pair_rapa_DD, pair_rapa_DA, pair_rapa_AA] = deal(zeros(512, 512, n_pair_rapa));
for i = 1:n_pair_rapa
    pair_rapa_DD(:,:,i) = double(imread(strcat(pair_rapa, '_', int2str(i), '_DD.tif')));
    pair_rapa_DA(:,:,i) = double(imread(strcat(pair_rapa, '_', int2str(i), '_DA.tif')));
    pair_rapa_AA(:,:,i) = double(imread(strcat(pair_rapa, '_', int2str(i), '_AA.tif')));
end

disp('All images loaded!');


%**** Part 2: Identify pixel intensities below background and saturated.
% A threshold is calculated for each channel from vector only samples,
% and all pixel intensities at or below the respective channel
% threshold are identified as background and set to 0.
% Pixels in each channel at intensity 255 (the maximum recorded) are
% considered to be saturated and removed.

% Calculate 99.9th perentile of autofluorescence in each channel.
Background_DD = prctile(vector_DD,99.9,'all');
Background_DA = prctile(vector_DA,99.9,'all');
Background_AA = prctile(vector_AA,99.9,'all');

% Set pixels with intensities below background threshold to zero
% for acceptor only, donor only, and paired samples.
acceptor_DD(find(acceptor_DD<Background_DD))=0;
acceptor_DA(find(acceptor_DA<Background_DA))=0;
acceptor_AA(find(acceptor_AA<Background_AA))=0;
donor_DD(find(donor_DD<Background_DD))=0;
donor_DA(find(donor_DA<Background_DA))=0;
donor_AA(find(donor_AA<Background_AA))=0;
pair_rapa_DD(find(pair_rapa_DD<Background_DD))=0;
pair_rapa_DA(find(pair_rapa_DA<Background_DA))=0;
pair_rapa_AA(find(pair_rapa_AA<Background_AA))=0;
pair_etoh_DD(find(pair_etoh_DD<Background_DD))=0;
pair_etoh_DA(find(pair_etoh_DA<Background_DA))=0;
pair_etoh_AA(find(pair_etoh_AA<Background_AA))=0;

% Set saturated pixel values to NaN
% for acceptor only, donor only, and paired samples.
acceptor_DD(find(acceptor_DD==255))=NaN;
acceptor_DA(find(acceptor_DA==255))=NaN;
acceptor_AA(find(acceptor_AA==255))=NaN;
donor_DD(find(donor_DD==255))=NaN;
donor_DA(find(donor_DA==255))=NaN;
donor_AA(find(donor_AA==255))=NaN;
pair_rapa_DD(find(pair_rapa_DD==255))=NaN;
pair_rapa_DA(find(pair_rapa_DA==255))=NaN;
pair_rapa_AA(find(pair_rapa_AA==255))=NaN;
pair_etoh_DD(find(pair_etoh_DD==255))=NaN;
pair_etoh_DA(find(pair_etoh_DA==255))=NaN;
pair_etoh_AA(find(pair_etoh_AA==255))=NaN;

disp('Background and saturated pixels have been adjusted!');


%**** Part 3: Correct for donor and acceptor spectral bleedthrough.
% Spectral bleedthrough ratios are calculated on a pixel-by-pixel
% basis from all fields of view in donor only and acceptor only
% samples and then averaged across all fields of view to arrive at
% constant value coefficients.
% Coefficients for spectral bleedthrough are based on the correction
% strategy by: Zal T & Gascoigne NRJ (2004) Biophys J 86:3923-3939.

% Calculate pixel-based ratios, remove infinite ratios, and calculate
% average coefficient values.
a_matrix = acceptor_DA./acceptor_AA;
a_matrix(find(a_matrix==Inf))=NaN;
a = nanmean(a_matrix,'all');
b_matrix = acceptor_DD./acceptor_AA;
b_matrix(find(b_matrix==Inf))=NaN;
b = nanmean(b_matrix,'all');
c_matrix = donor_AA./donor_DD;
c_matrix(find(c_matrix==Inf))=NaN;
c = nanmean(c_matrix,'all');
d_matrix = donor_DA./donor_DD;
d_matrix(find(d_matrix==Inf))=NaN;
d = nanmean(d_matrix,'all');

% Perform spectral bleedthrough correction from FRET channel matrices,
% setting intensities that become negative to 0.
pair_rapa_cDA = pair_rapa_DA-(a*(pair_rapa_AA-(c*pair_rapa_DD)))-(d*(pair_rapa_DD-(b*pair_rapa_AA)));
pair_rapa_cDA(find(pair_rapa_cDA<0))=0;
pair_etoh_cDA = pair_etoh_DA-(a*(pair_etoh_AA-(c*pair_etoh_DD)))-(d*(pair_etoh_DD-(b*pair_etoh_AA)));
pair_etoh_cDA(find(pair_etoh_cDA<0))=0;

% Remove pixels where corrected FRET (cDA) intensity exceeds the
% intensity in the AA channel.
pair_rapa_cDA(find(pair_rapa_cDA>pair_rapa_AA))=NaN;
pair_etoh_cDA(find(pair_etoh_cDA>pair_etoh_AA))=NaN;

disp('Bleedthrough has been corrected!')


%**** Part 4: Calculate NFRET
% Normalize corrected FRET intensity to the square root of the product
% of donor and acceptor intensities on a pixel-by-pixel basis.
% This normalized metric is termed NFRET, originally coined by
% Xia Z, Liu Y. Biophys J. (2001).

% Calculate NFRET on a pixel-by-pixel basis, removing pixels with
% infinite values.
pair_rapa_NFRET = pair_rapa_cDA./(sqrt(pair_rapa_DD).*sqrt(pair_rapa_AA));
pair_rapa_NFRET(find(pair_rapa_NFRET==inf))=NaN;
pair_etoh_NFRET = pair_etoh_cDA./(sqrt(pair_etoh_DD).*sqrt(pair_etoh_AA));
pair_etoh_NFRET(find(pair_etoh_NFRET==inf))=NaN;

% Calculate NFRET mean within each field of view
pair_rapa_avgNFRET = zeros(1,n_pair_rapa);
pair_etoh_avgNFRET = zeros(1,n_pair_rapa);
for i = 1:n_pair_rapa
    pair_rapa_avgNFRET(i) = nanmean(pair_rapa_NFRET(:,:,i),[1 2]);
end
for i = 1:n_pair_etoh
    pair_etoh_avgNFRET(i) = nanmean(pair_etoh_NFRET(:,:,i),[1,2]);
end

% Calculate bulk NFRET mean and SEM across all fields of view
pair_rapa_bulkNFRET = nanmean(pair_rapa_avgNFRET,'all');
pair_etoh_bulkNFRET = nanmean(pair_etoh_avgNFRET,'all');
pair_rapa_bulkNFRET_SEM = std(pair_rapa_avgNFRET)/sqrt(length(pair_rapa_avgNFRET));
pair_etoh_bulkNFRET_SEM = std(pair_etoh_avgNFRET)/sqrt(length(pair_etoh_avgNFRET));

disp('NFRET pixel values and averages have been calculated!');


%**** Part 5: Plot and export images depicting NFRET

% Generate colormap matrix (fire)
fire =  [
    0         0         0
    0         0    0.0275
    0         0    0.0588
    0         0    0.0863
    0         0    0.1176
    0         0    0.1490
    0         0    0.1765
    0         0    0.2078
    0         0    0.2392
    0         0    0.2549
    0         0    0.2706
    0         0    0.2902
    0         0    0.3059
    0         0    0.3216
    0         0    0.3412
    0         0    0.3569
    0.0039         0    0.3765
    0.0157         0    0.3922
    0.0275         0    0.4078
    0.0392         0    0.4235
    0.0510         0    0.4431
    0.0627         0    0.4588
    0.0745         0    0.4745
    0.0863         0    0.4902
    0.0980         0    0.5098
    0.1098         0    0.5255
    0.1216         0    0.5412
    0.1333         0    0.5608
    0.1451         0    0.5765
    0.1569         0    0.5922
    0.1686         0    0.6118
    0.1804         0    0.6275
    0.1922         0    0.6471
    0.2039         0    0.6588
    0.2157         0    0.6706
    0.2275         0    0.6863
    0.2392         0    0.6980
    0.2510         0    0.7098
    0.2627         0    0.7255
    0.2745         0    0.7373
    0.2863         0    0.7529
    0.2980         0    0.7647
    0.3098         0    0.7804
    0.3216         0    0.7922
    0.3333         0    0.8078
    0.3451         0    0.8196
    0.3569         0    0.8353
    0.3686         0    0.8471
    0.3843         0    0.8627
    0.3961         0    0.8627
    0.4078         0    0.8667
    0.4196         0    0.8706
    0.4314         0    0.8745
    0.4431         0    0.8784
    0.4549         0    0.8824
    0.4667         0    0.8863
    0.4784         0    0.8902
    0.4902         0    0.8784
    0.5020         0    0.8706
    0.5137         0    0.8627
    0.5255         0    0.8549
    0.5373         0    0.8471
    0.5490         0    0.8392
    0.5608         0    0.8314
    0.5725         0    0.8235
    0.5804         0    0.8078
    0.5882         0    0.7922
    0.5961         0    0.7804
    0.6039         0    0.7647
    0.6118         0    0.7490
    0.6196         0    0.7373
    0.6275         0    0.7216
    0.6353         0    0.7098
    0.6392         0    0.6941
    0.6431         0    0.6784
    0.6510         0    0.6627
    0.6549         0    0.6510
    0.6588         0    0.6353
    0.6667         0    0.6196
    0.6706         0    0.6039
    0.6784         0    0.5922
    0.6824         0    0.5765
    0.6863         0    0.5608
    0.6941         0    0.5490
    0.6980         0    0.5333
    0.7020         0    0.5176
    0.7098         0    0.5059
    0.7137         0    0.4902
    0.7216         0    0.4784
    0.7255         0    0.4627
    0.7294         0    0.4471
    0.7373         0    0.4353
    0.7412         0    0.4196
    0.7451         0    0.4039
    0.7529         0    0.3922
    0.7569         0    0.3765
    0.7647         0    0.3647
    0.7686    0.0039    0.3490
    0.7765    0.0118    0.3333
    0.7804    0.0196    0.3216
    0.7882    0.0275    0.3059
    0.7922    0.0314    0.2902
    0.8000    0.0392    0.2784
    0.8039    0.0471    0.2627
    0.8118    0.0549    0.2510
    0.8157    0.0627    0.2353
    0.8196    0.0745    0.2196
    0.8235    0.0824    0.2078
    0.8314    0.0941    0.1922
    0.8353    0.1059    0.1765
    0.8392    0.1137    0.1647
    0.8431    0.1255    0.1490
    0.8510    0.1373    0.1373
    0.8549    0.1451    0.1216
    0.8627    0.1569    0.1059
    0.8667    0.1686    0.0902
    0.8745    0.1804    0.0784
    0.8784    0.1882    0.0627
    0.8863    0.2000    0.0471
    0.8902    0.2118    0.0314
    0.8980    0.2235    0.0196
    0.9020    0.2314    0.0157
    0.9059    0.2431    0.0118
    0.9137    0.2549    0.0118
    0.9176    0.2667    0.0078
    0.9216    0.2745    0.0039
    0.9294    0.2863    0.0039
    0.9333    0.2980         0
    0.9412    0.3098         0
    0.9451    0.3176         0
    0.9529    0.3294         0
    0.9569    0.3412         0
    0.9647    0.3529         0
    0.9686    0.3608         0
    0.9765    0.3725         0
    0.9804    0.3843         0
    0.9882    0.3961         0
    0.9882    0.4039         0
    0.9882    0.4118         0
    0.9922    0.4196         0
    0.9922    0.4275         0
    0.9922    0.4353         0
    0.9961    0.4431         0
    0.9961    0.4510         0
    1.0000    0.4588         0
    1.0000    0.4667         0
    1.0000    0.4745         0
    1.0000    0.4824         0
    1.0000    0.4902         0
    1.0000    0.4980         0
    1.0000    0.5059         0
    1.0000    0.5137         0
    1.0000    0.5216         0
    1.0000    0.5255         0
    1.0000    0.5333         0
    1.0000    0.5412         0
    1.0000    0.5490         0
    1.0000    0.5529         0
    1.0000    0.5608         0
    1.0000    0.5686         0
    1.0000    0.5765         0
    1.0000    0.5804         0
    1.0000    0.5882         0
    1.0000    0.5961         0
    1.0000    0.6039         0
    1.0000    0.6078         0
    1.0000    0.6157         0
    1.0000    0.6235         0
    1.0000    0.6314         0
    1.0000    0.6353         0
    1.0000    0.6431         0
    1.0000    0.6510         0
    1.0000    0.6588         0
    1.0000    0.6627         0
    1.0000    0.6706         0
    1.0000    0.6784         0
    1.0000    0.6863         0
    1.0000    0.6902         0
    1.0000    0.6980         0
    1.0000    0.7059         0
    1.0000    0.7137         0
    1.0000    0.7216         0
    1.0000    0.7294         0
    1.0000    0.7373         0
    1.0000    0.7451         0
    1.0000    0.7490         0
    1.0000    0.7569         0
    1.0000    0.7647         0
    1.0000    0.7725         0
    1.0000    0.7804         0
    1.0000    0.7882         0
    1.0000    0.7961         0
    1.0000    0.8039         0
    1.0000    0.8078         0
    1.0000    0.8157         0
    1.0000    0.8235         0
    1.0000    0.8314         0
    1.0000    0.8353         0
    1.0000    0.8431         0
    1.0000    0.8510         0
    1.0000    0.8588         0
    1.0000    0.8627         0
    1.0000    0.8706         0
    1.0000    0.8784         0
    1.0000    0.8863         0
    1.0000    0.8941         0
    1.0000    0.9020         0
    1.0000    0.9098         0
    1.0000    0.9176         0
    1.0000    0.9216    0.0157
    1.0000    0.9294    0.0314
    1.0000    0.9373    0.0510
    1.0000    0.9451    0.0667
    1.0000    0.9490    0.0824
    1.0000    0.9569    0.1020
    1.0000    0.9647    0.1176
    1.0000    0.9725    0.1373
    1.0000    0.9725    0.1647
    1.0000    0.9765    0.1961
    1.0000    0.9804    0.2275
    1.0000    0.9843    0.2588
    1.0000    0.9882    0.2902
    1.0000    0.9922    0.3216
    1.0000    0.9961    0.3529
    1.0000    1.0000    0.3843
    1.0000    1.0000    0.4118
    1.0000    1.0000    0.4431
    1.0000    1.0000    0.4745
    1.0000    1.0000    0.5059
    1.0000    1.0000    0.5333
    1.0000    1.0000    0.5647
    1.0000    1.0000    0.5961
    1.0000    1.0000    0.6275
    1.0000    1.0000    0.6549
    1.0000    1.0000    0.6863
    1.0000    1.0000    0.7176
    1.0000    1.0000    0.7490
    1.0000    1.0000    0.7804
    1.0000    1.0000    0.8118
    1.0000    1.0000    0.8431
    1.0000    1.0000    0.8745
    1.0000    1.0000    0.8902
    1.0000    1.0000    0.9059
    1.0000    1.0000    0.9216
    1.0000    1.0000    0.9373
    1.0000    1.0000    0.9529
    1.0000    1.0000    0.9686
    1.0000    1.0000    0.9843
    1.0000    1.0000    1.0000
    1.0000    1.0000    1.0000
    1.0000    1.0000    1.0000
    1.0000    1.0000    1.0000
    1.0000    1.0000    1.0000
    1.0000    1.0000    1.0000
    1.0000    1.0000    1.0000
    1.0000    1.0000    1.0000];


% Plot and export images depicting NFRET for each field of view
% for both the rapalog-treated and EtOH-treated samples (20 total images).
for i = 1:n_pair_rapa
    imAlpha = ones(size(pair_rapa_NFRET(:,:,i)));
    imAlpha(isnan(pair_rapa_NFRET(:,:,i)))=0;
    figure; imagesc(pair_rapa_NFRET(:,:,i),'AlphaData',imAlpha);
    truesize;
    set(gca,'color',0*[1,1,1]);
    colormap(gca,fire);
    caxis([0 1]);
    title([strcat('NFRET_', pair_rapa, '\_'),num2str(i)]);
end
for i = 1:n_pair_etoh
    imAlpha = ones(size(pair_etoh_NFRET(:,:,i)));
    imAlpha(isnan(pair_etoh_NFRET(:,:,i)))=0;
    figure; imagesc(pair_etoh_NFRET(:,:,i),'AlphaData',imAlpha);
    truesize;
    set(gca,'color',0*[1,1,1]);
    colormap(gca,fire);
    caxis([0 1]);
    title([strcat('NFRET_', pair_etoh, '\_'),num2str(i)]);
end
disp('All plots open, now saving...');

% Find all open figures.
figList = findobj(allchild(0), 'flat', 'Type', 'figure');

% Save and close all open plots to specified folder.
for i = 1:length(figList)
    fig = figList(i);
    figTitle = get(gca, 'Title');
    imageName = get(figTitle, 'String');
    imageName = erase(imageName, '\');
    figFrame = getframe(gca);
    imageName = strcat(folderName, imageName, '.png');
    imwrite(figFrame.cdata, imageName);
    close(fig);
end

disp('All NFRET plots have been closed and saved to specified folder.');
disp('Script complete!');


end
```
