## Supplementary Software for "Elucidation and refinement of synthetic receptor mechanisms": LICENSE.rtf

Permission is hereby granted, free of charge, to any person obtaining a copy of this software and associated documentation 
files (the "Software"), to deal in the Software without restriction, including without limitation the rights to use, copy, 
modify, merge, publish, distribute, sublicense, and/or sell copies of the Software, and to permit persons to whom the Software 
is furnished to do so, subject to the following conditions:

The above copyright notice and this permission notice shall be included in all copies or substantial portions of the Software.

THE SOFTWARE IS PROVIDED "AS IS", WITHOUT WARRANTY OF ANY KIND, EXPRESS OR IMPLIED, INCLUDING BUT NOT LIMITED TO THE WARRANTIES 
OF MERCHANTABILITY, FITNESS FOR A PARTICULAR PURPOSE AND NONINFRINGEMENT. IN NO EVENT SHALL THE AUTHORS OR COPYRIGHT HOLDERS BE 
LIABLE FOR ANY CLAIM, DAMAGES OR OTHER LIABILITY, WHETHER IN AN ACTION OF CONTRACT, TORT OR OTHERWISE, ARISING FROM, OUT OF OR 
IN CONNECTION WITH THE SOFTWARE OR THE USE OR OTHER DEALINGS IN THE SOFTWARE.
