## Supplementary Software for "Elucidation and refinement of synthetic receptor mechanisms": README.rtf


Description of files: 

FRET_SE_Correct.m: Script for processing confocal microscopy images for FRET analysis using sensitized emission. This workflow performs background subtraction, corrects for spectral bleed-through, and calculates both a sensitized FRET emission metric and a normalized FRET (NFRET) metric. 


1. System requirements

The code can be run using Matlab (https://www.mathworks.com/products/matlab.html) and run on an operating system that supports Matlab. Code was developed and tested using Matlab version 2019a on macOS Catalina.


2. Installation guide

No other specific installations are required.


3. Instructions for use

This script is designed to serve as an example of the image processing workflow used in this study. It is tailored to a set of provided sample images but can be used as a template and adjusted for a user-collected set of images. Input images must be separated channel images (as opposed to stacks) that are each 512x512 pixels. These images should be stored in the same folder with the script. A set of 50 sample images is provided to evaluate FRET for the 3xFLAG-FKBP-CD28-mCerulean and 3xFLAG-FRB-CD28-mVenus rapamycin-sensing MESA receptor pair. The set includes 10 fields of view for samples transfected with: 
	A. Vector-only (modified pcDNA)
	B. Donor (3xFLAG-FKBP-CD28-mCerulean alone)
	C. Acceptor (3xFLAG-FRB-CD28-mVenus alone)
	D. Donor and acceptor (treated with EtOH)
	E. Donor and acceptor (treated with rapalog). 

Specifications are first made for number of fields of view for each sample type, file naming system, and processed image output folder. The image processing and export then occurs in parts 1 through 5. Output images are saved as separate .png files, 512x512 pixels. 

The expected run time for the script is 10-15 seconds on a standard desktop computer.
