## Supplementary figures and images for "Elucidation and refinement of synthetic receptor mechanisms"

### D_3_DA.tif

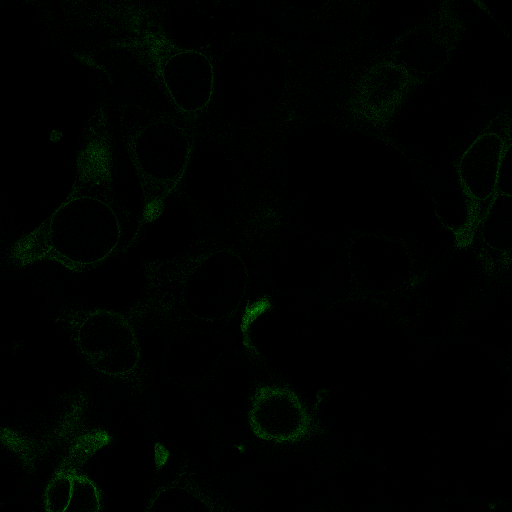

### D_3_DD.tif

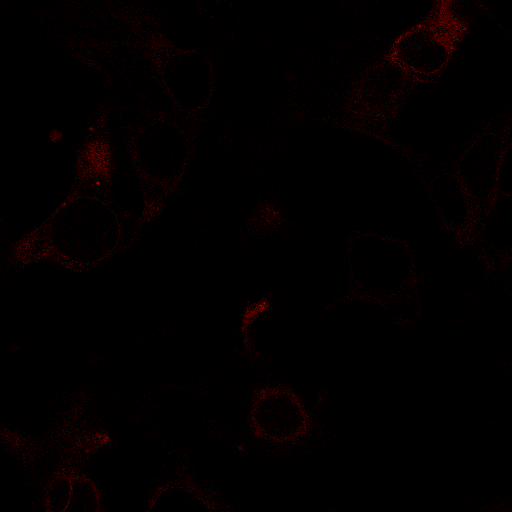

### D_4_AA.tif

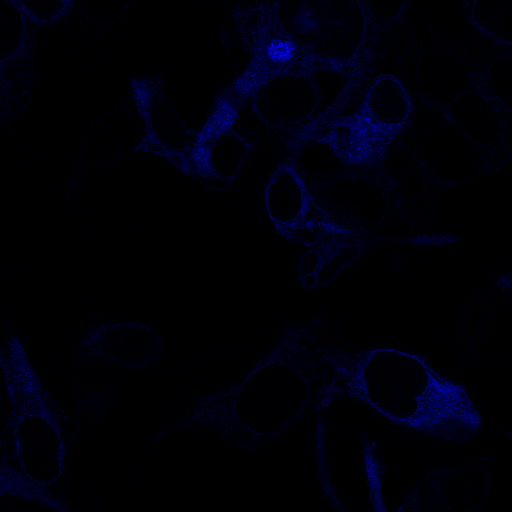

### D_4_DA.tif

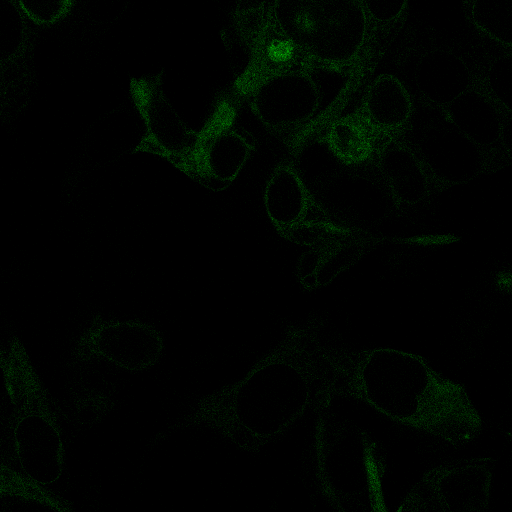

### D_4_DD.tif

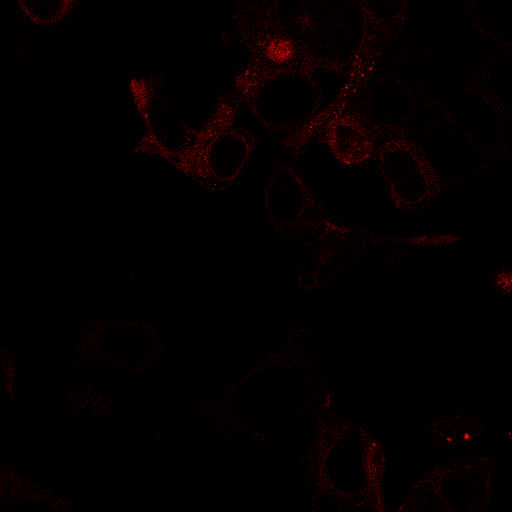

### D_5_AA.tif

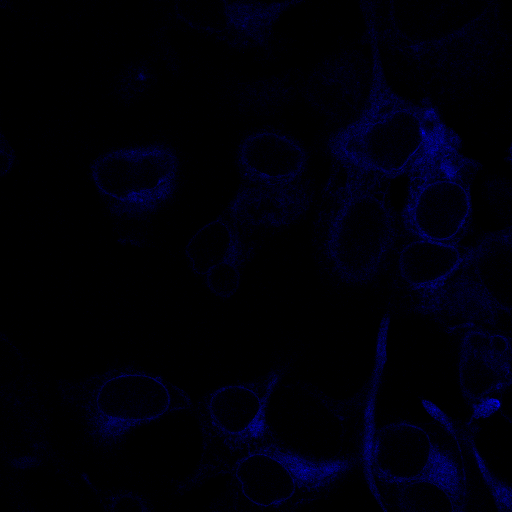

### D_5_DA.tif

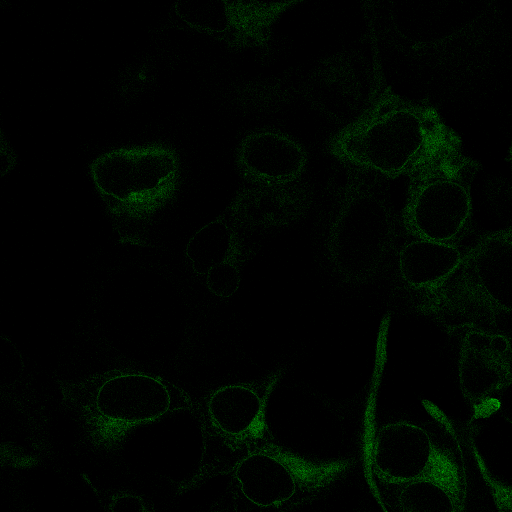

### D_5_DD.tif

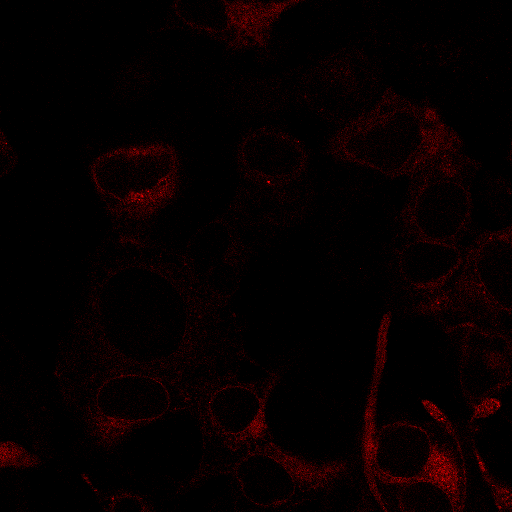

### D_6_AA.tif

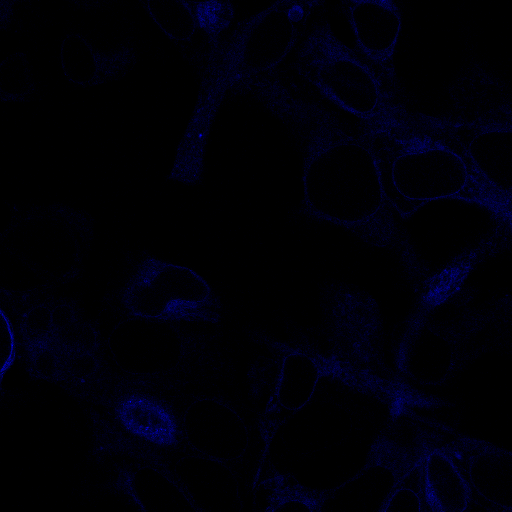

### D_6_DA.tif

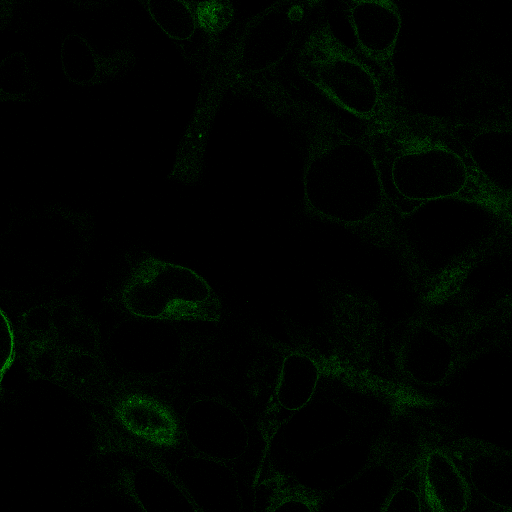

### D_6_DD.tif

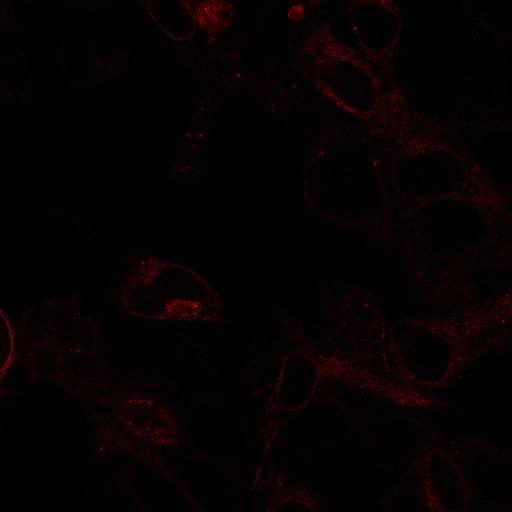

### D_7_AA.tif

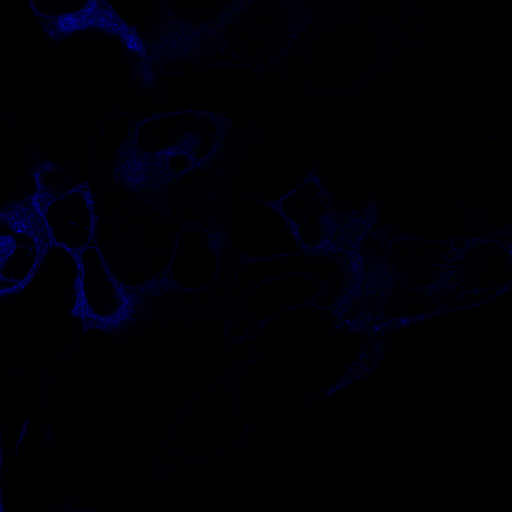

### D_7_DA.tif

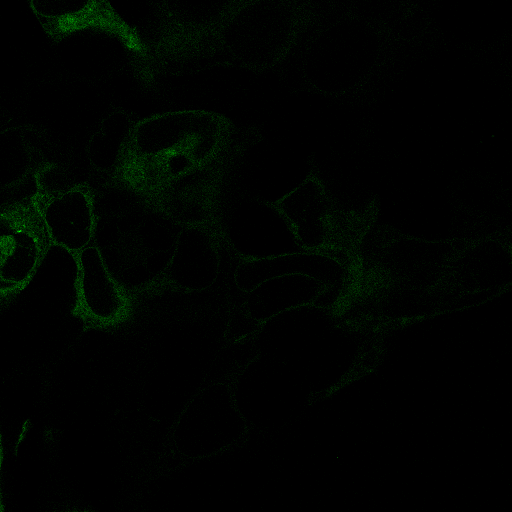

### D_7_DD.tif

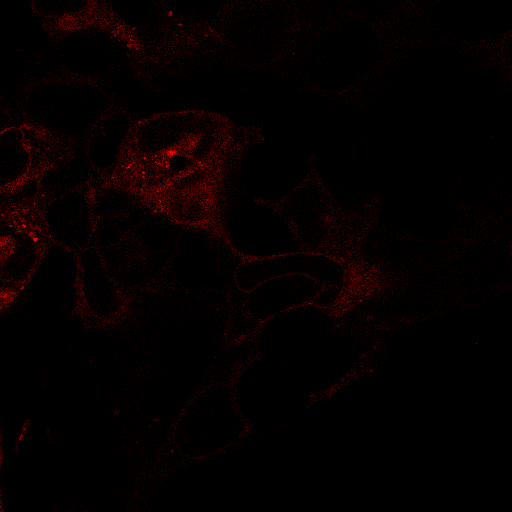

### D_8_AA.tif

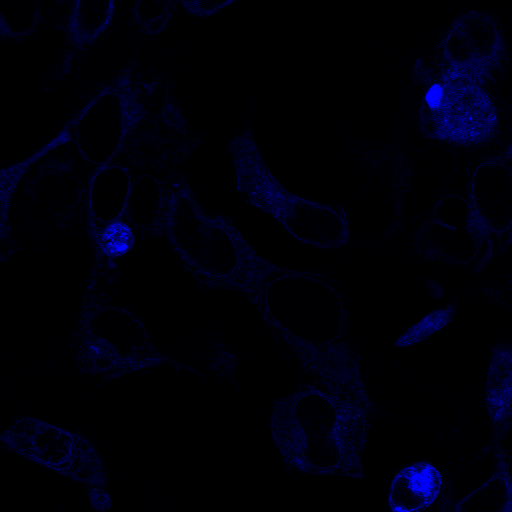

### D_8_DA.tif

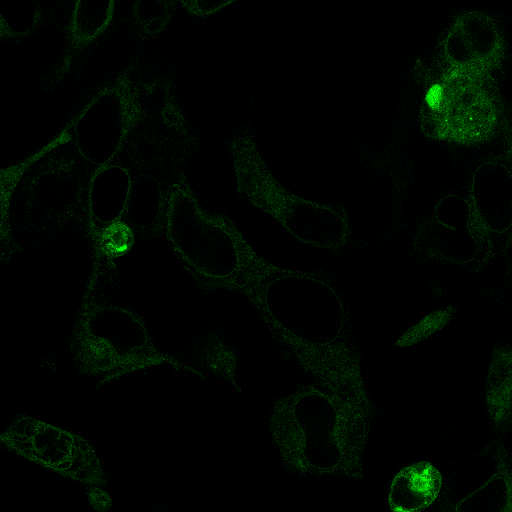

### D_8_DD.tif

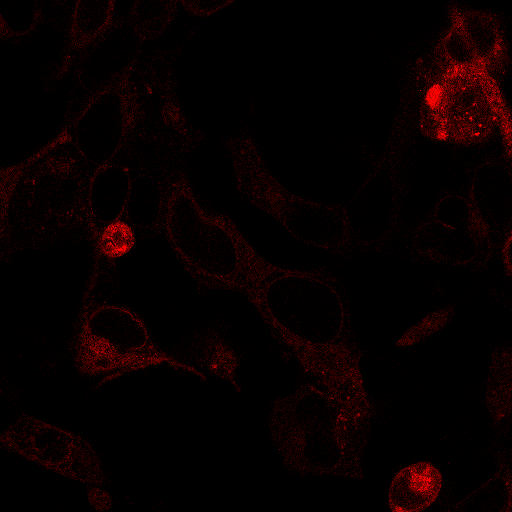

### D_9_AA.tif

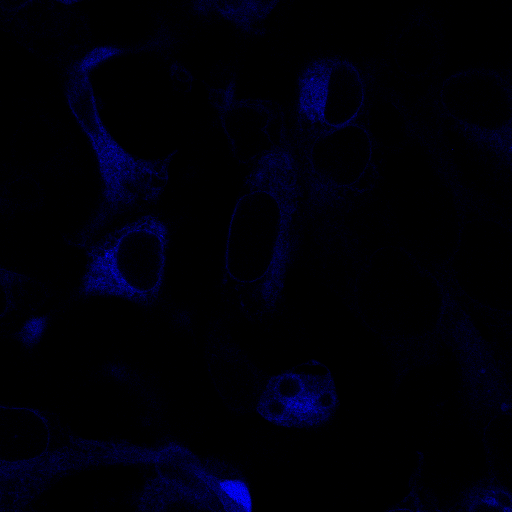

### D_9_DA.tif

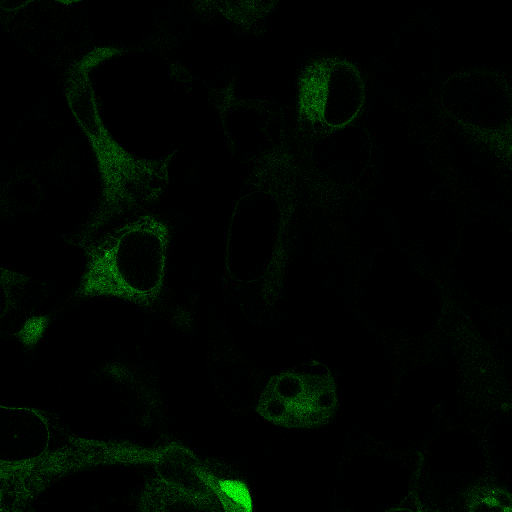

### D_9_DD.tif

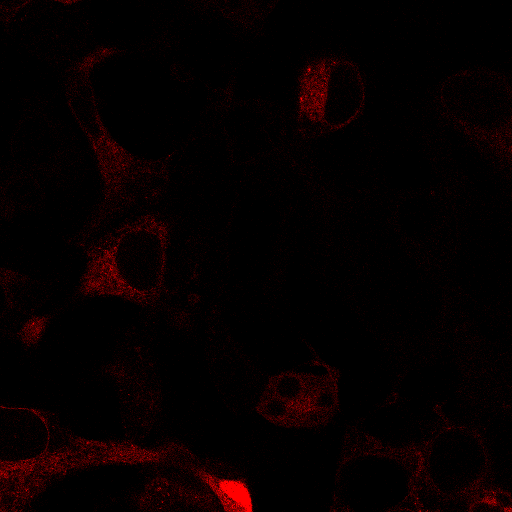

### E_1_AA.tif

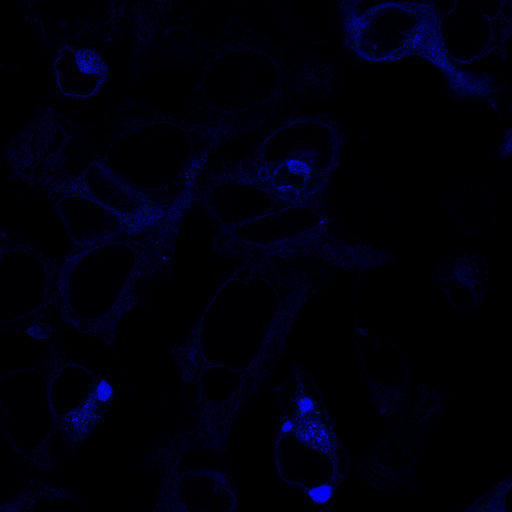

### E_1_DA.tif

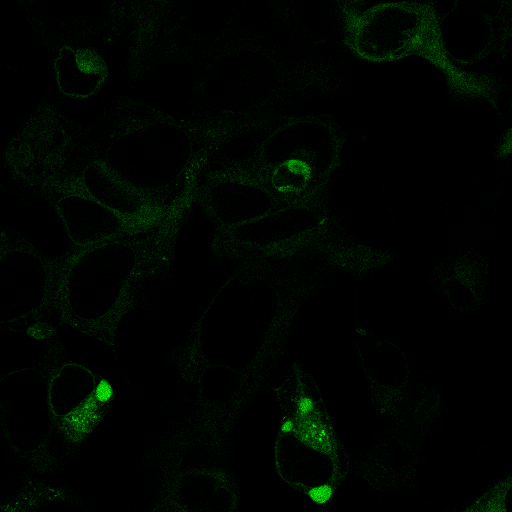

### E_1_DD.tif

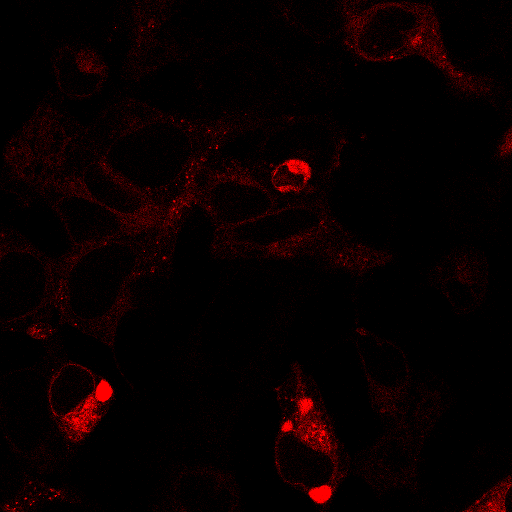

### E_2_AA.tif

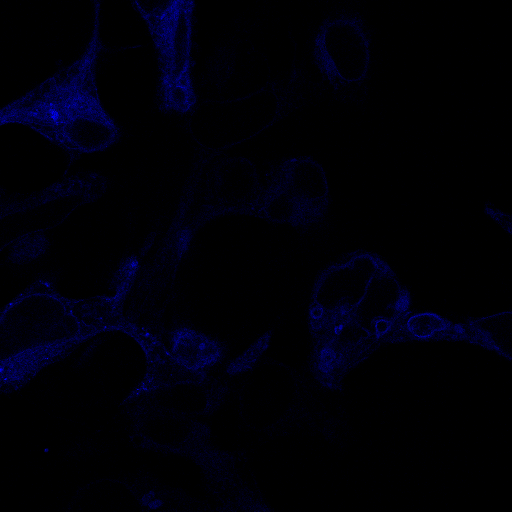

### E_2_DA.tif

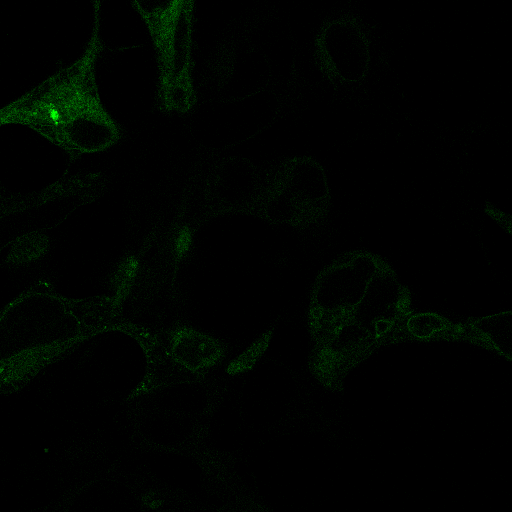

### E_2_DD.tif

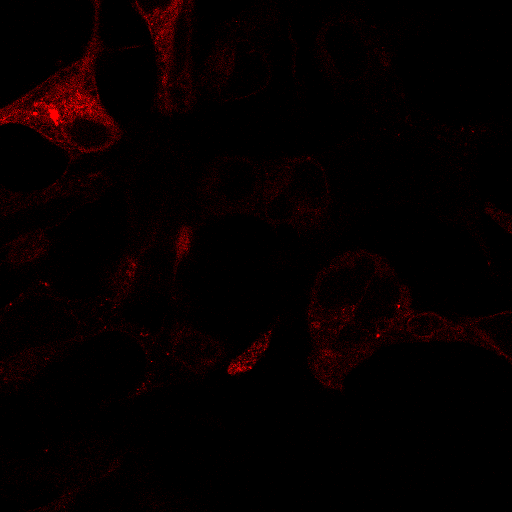

### E_3_AA.tif

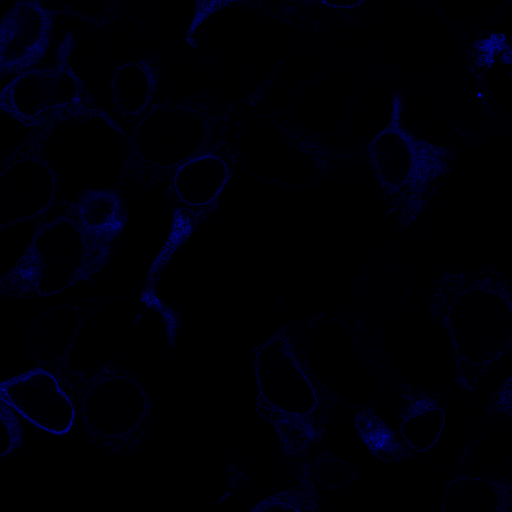

### E_3_DA.tif

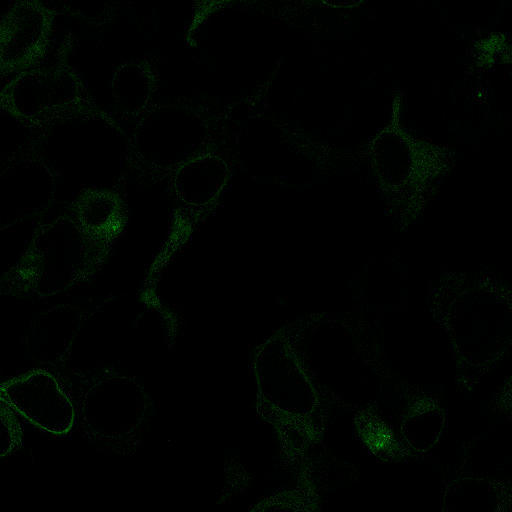

### E_3_DD.tif

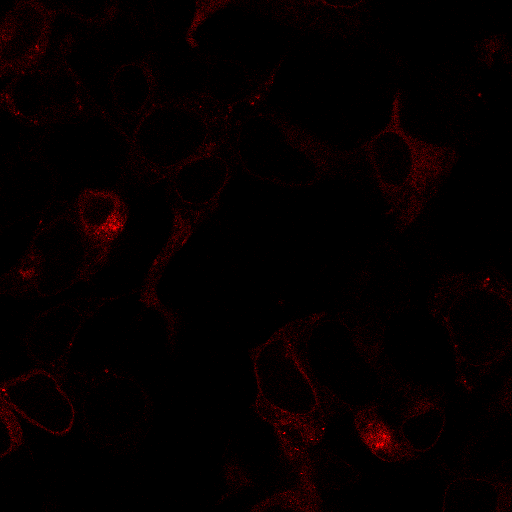

### E_4_AA.tif

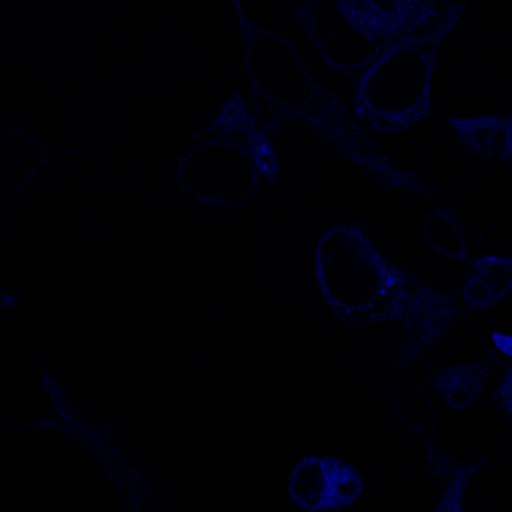

### E_4_DA.tif

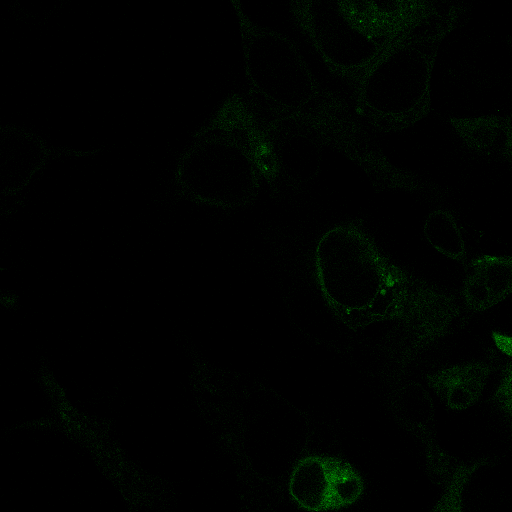

### E_4_DD.tif

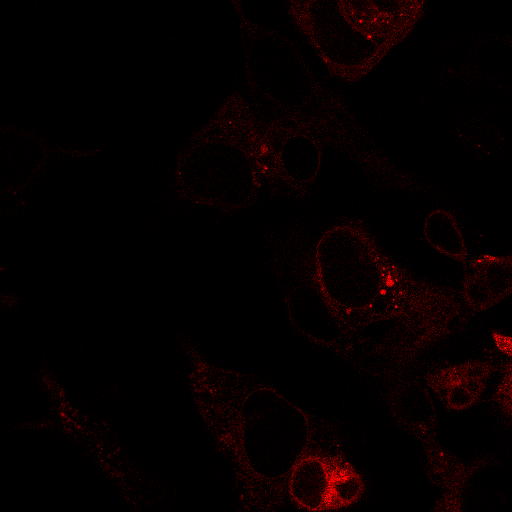

### E_5_AA.tif

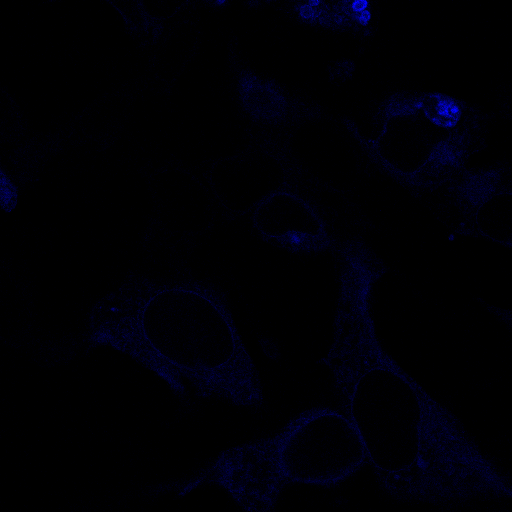

### E_5_DA.tif

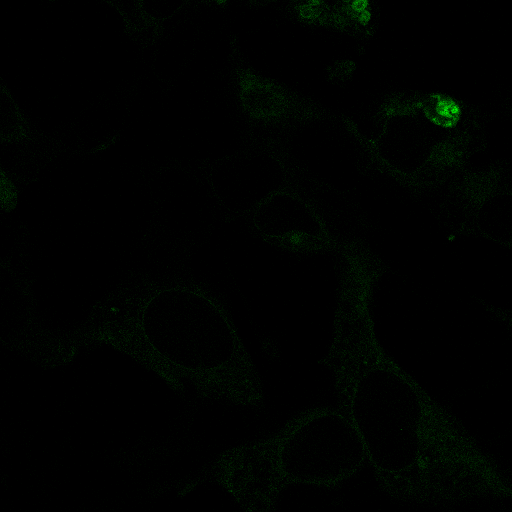

### E_5_DD.tif

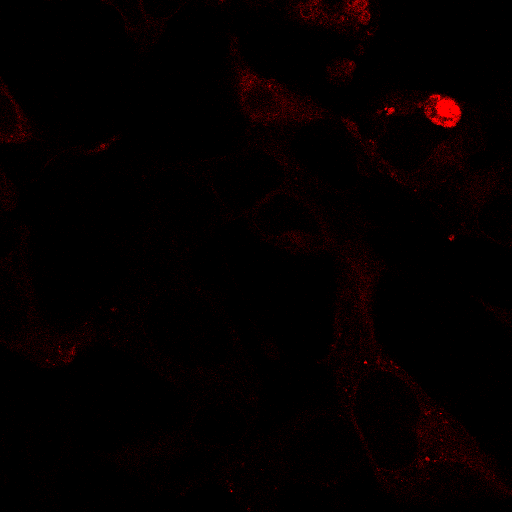

### E_6_AA.tif

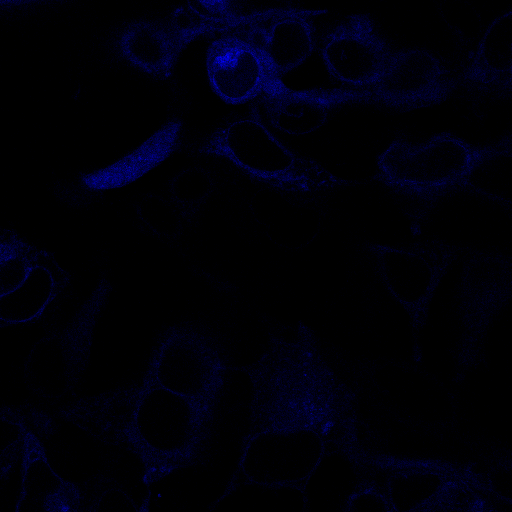

### E_6_DA.tif

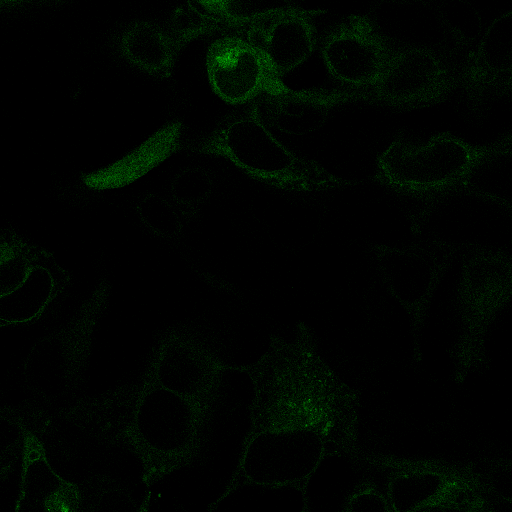

### E_6_DD.tif

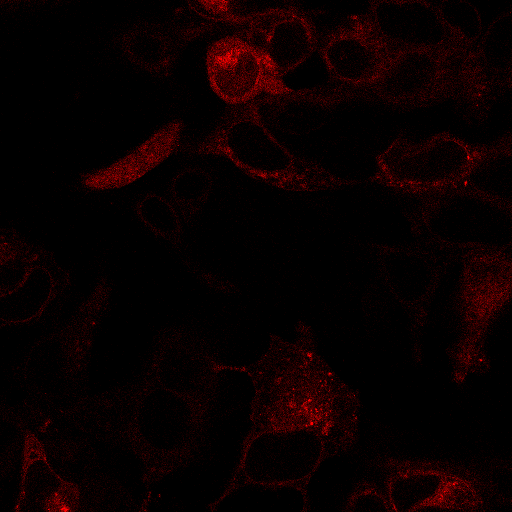

### E_7_AA.tif

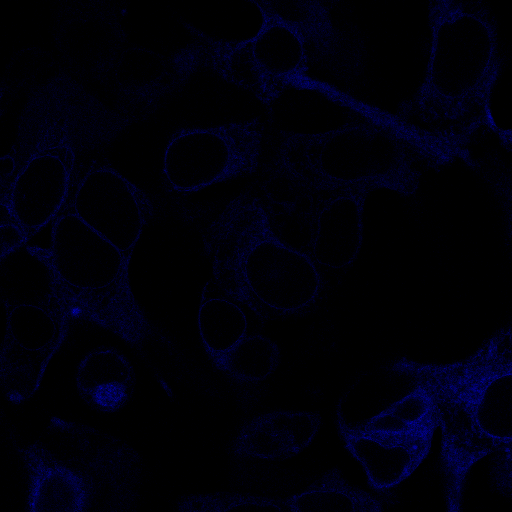

### E_7_DA.tif

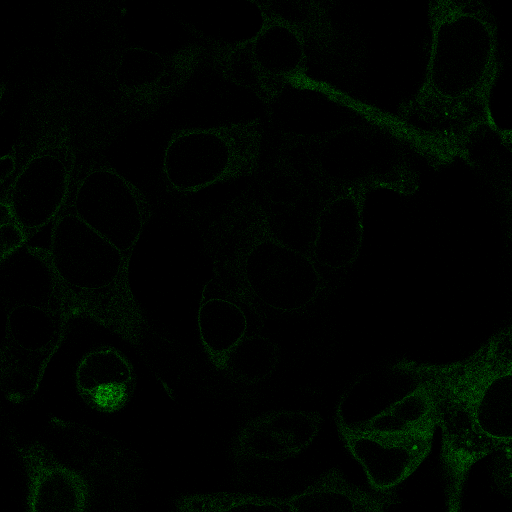

### E_7_DD.tif

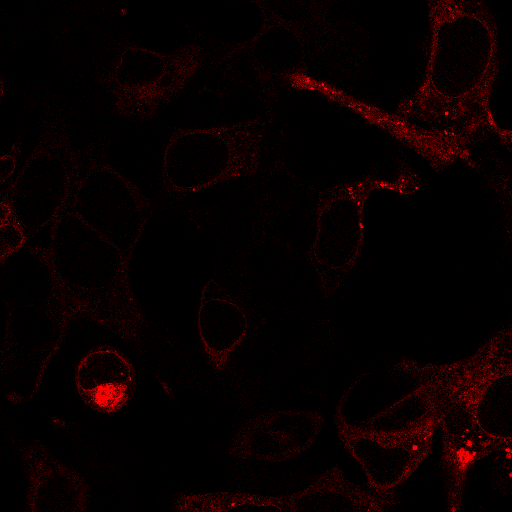

### E_8_AA.tif

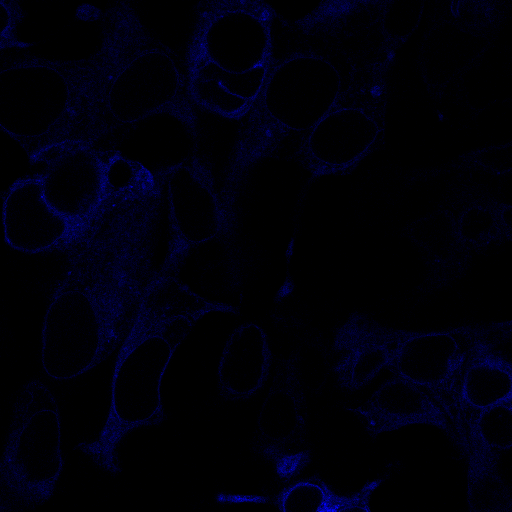

### E_8_DA.tif

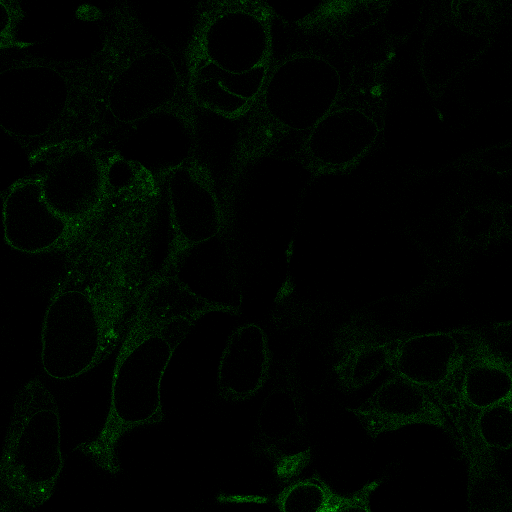

### E_8_DD.tif

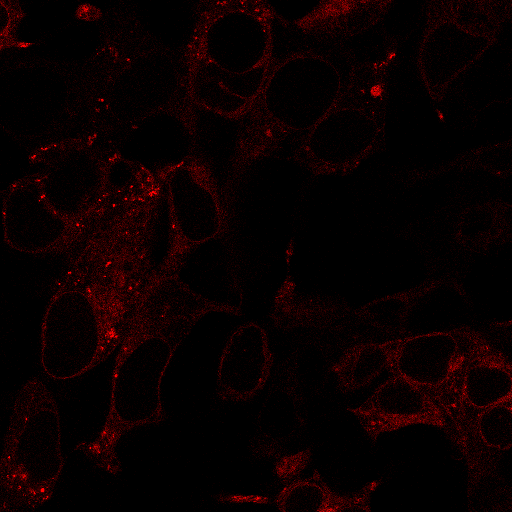

### E_9_AA.tif

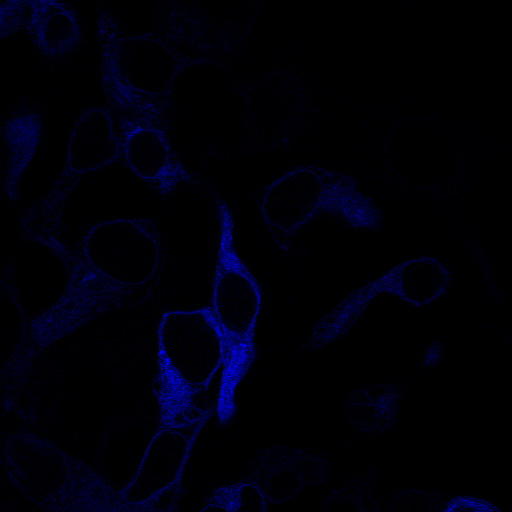

### E_9_DA.tif

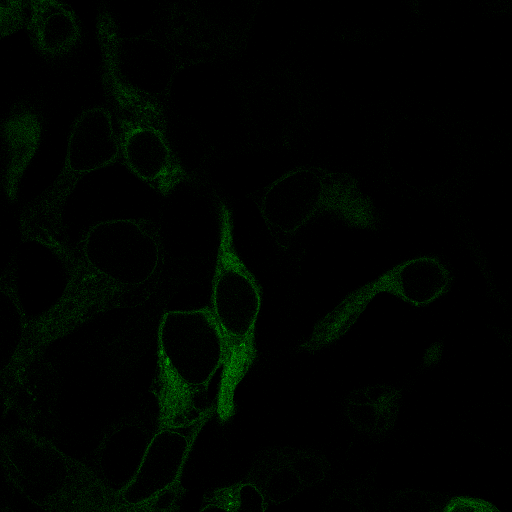

### E_9_DD.tif

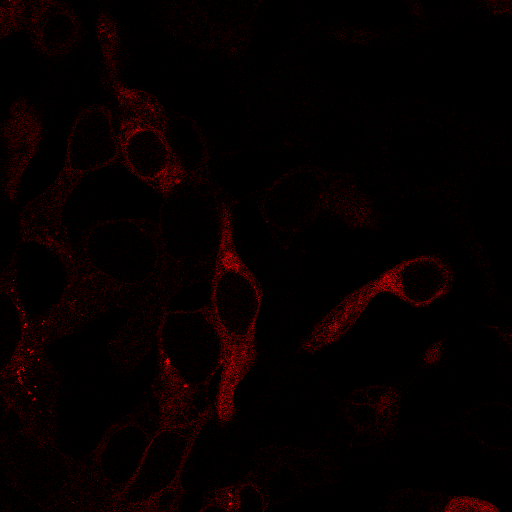

### E_10_AA.tif

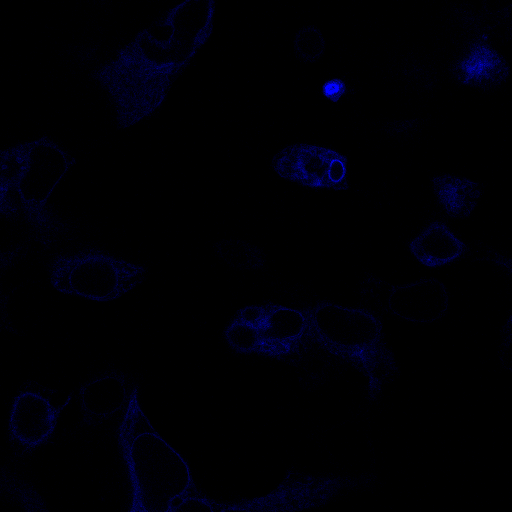

### E_10_DA.tif

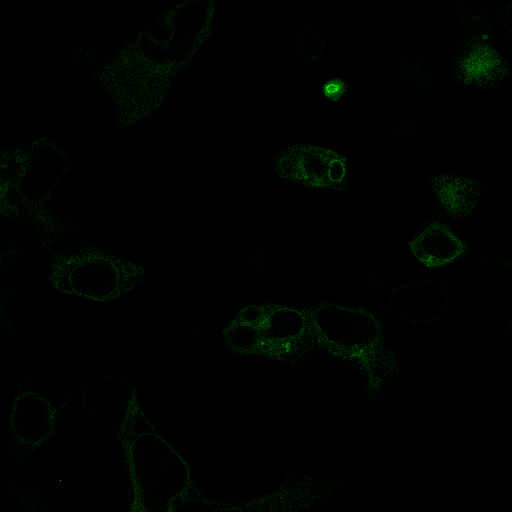

### E_10_DD.tif

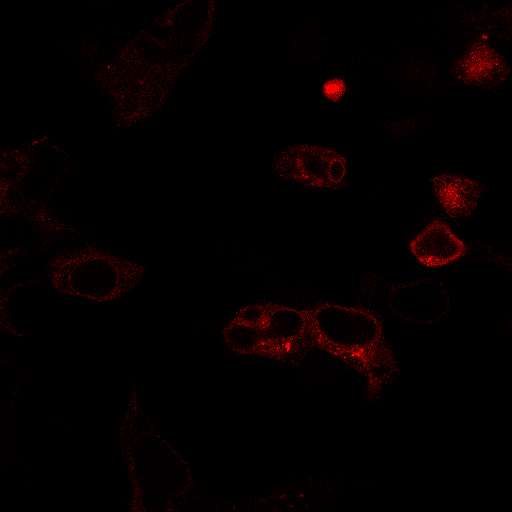
