## Supplementary figures and images for "Elucidation and refinement of synthetic receptor mechanisms"

### A_1_AA.tif

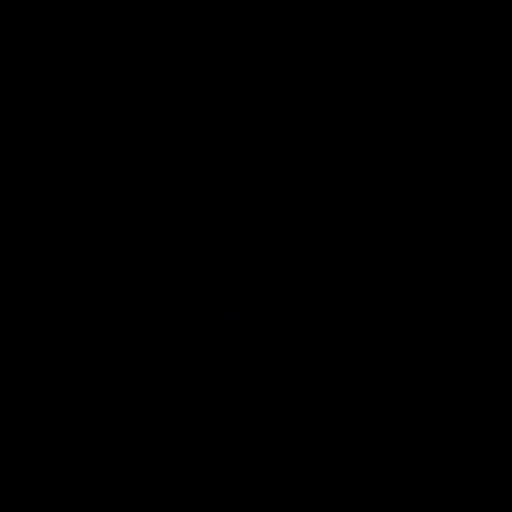

### A_1_DA.tif

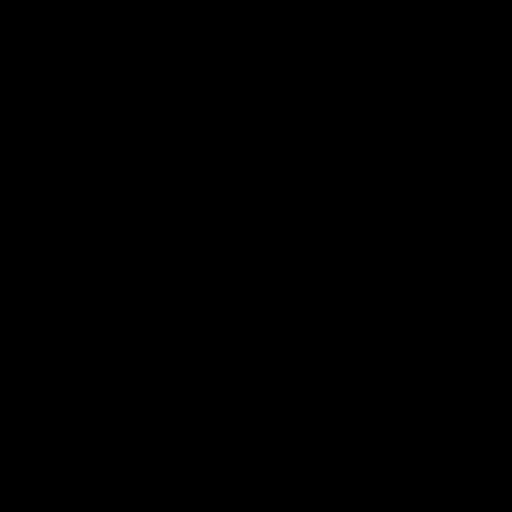

### A_1_DD.tif

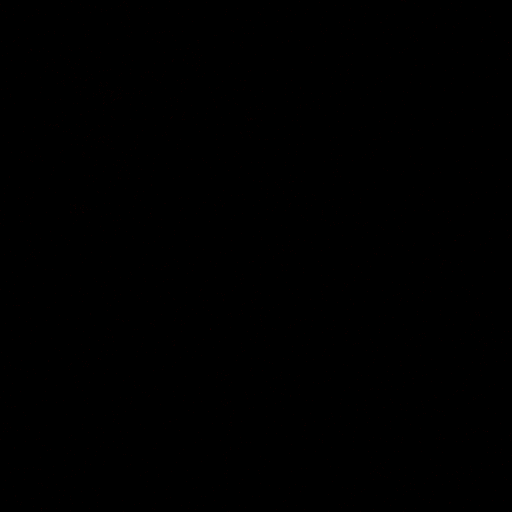

### A_2_AA.tif

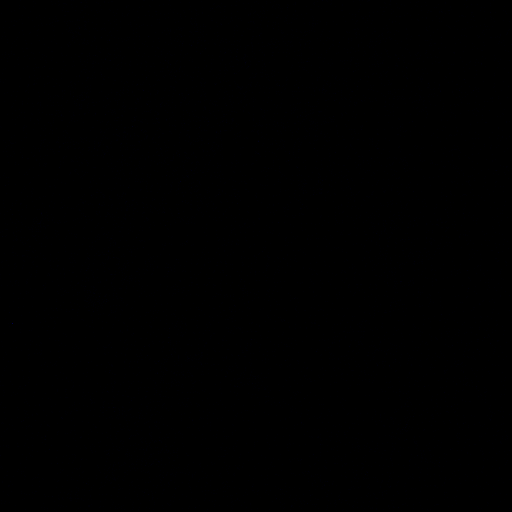

### A_2_DA.tif

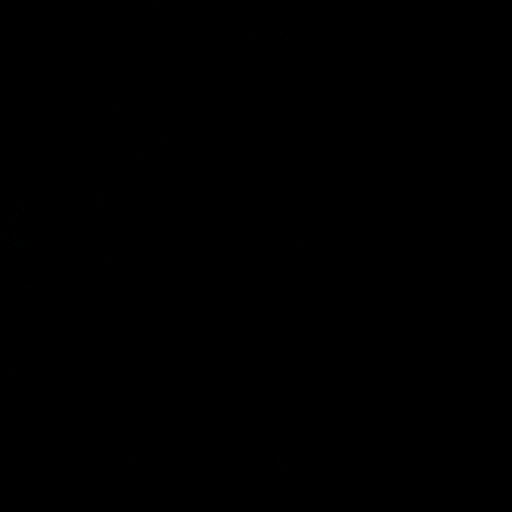

### A_2_DD.tif

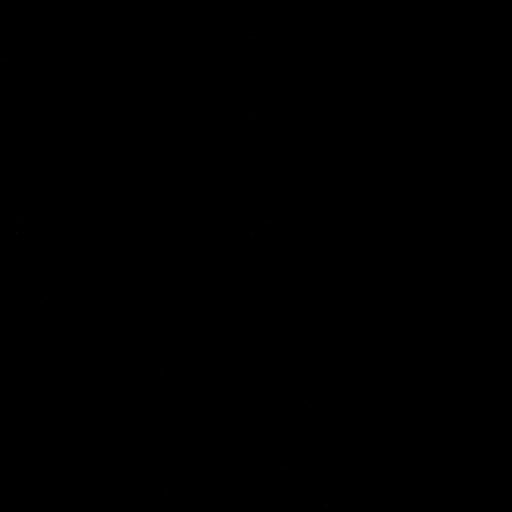

### A_3_AA.tif

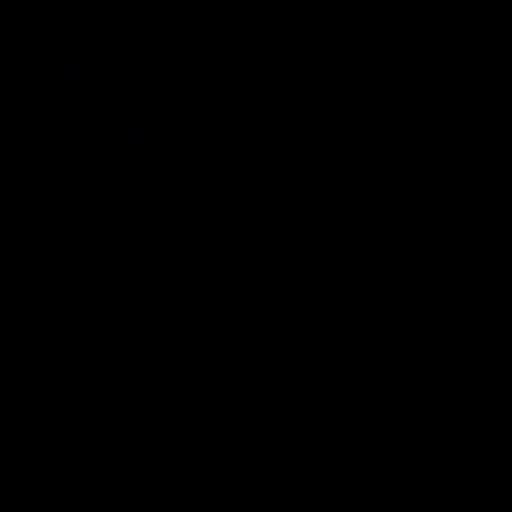

### A_3_DA.tif

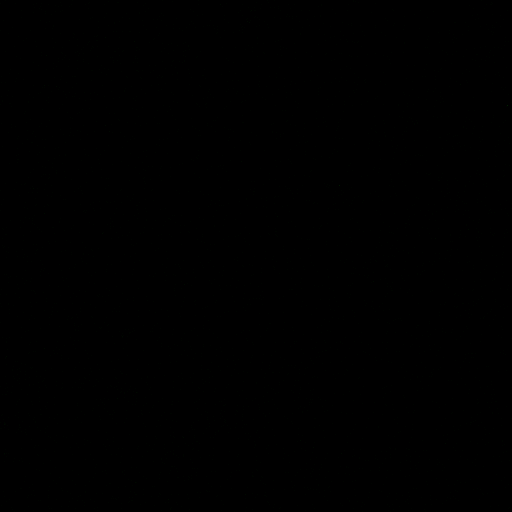

### A_3_DD.tif

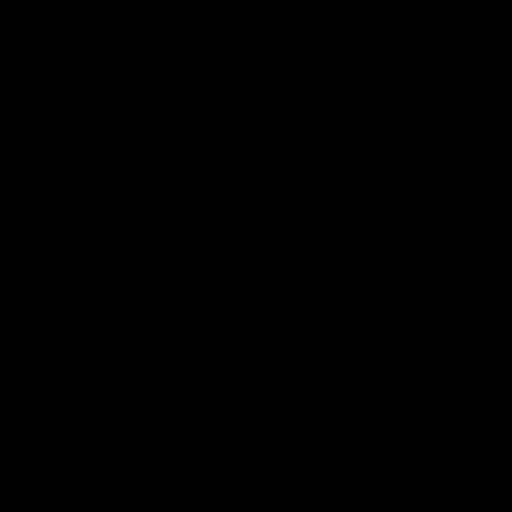

### A_4_AA.tif

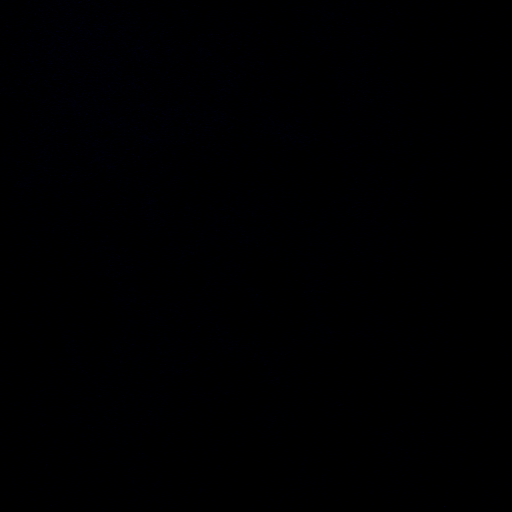

### A_4_DA.tif

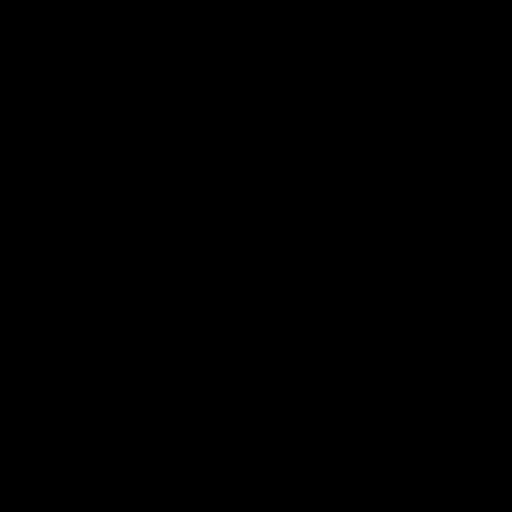

### A_4_DD.tif

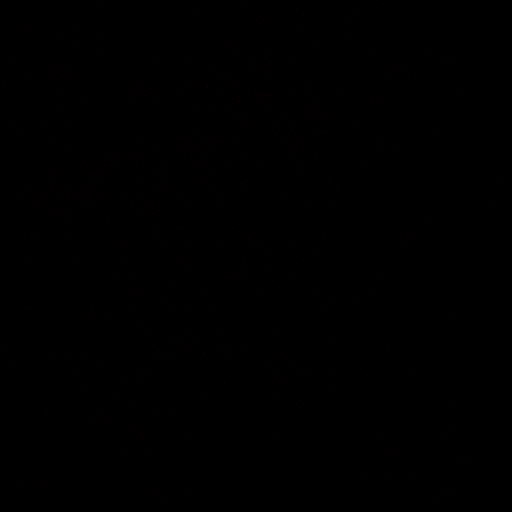

### A_5_AA.tif

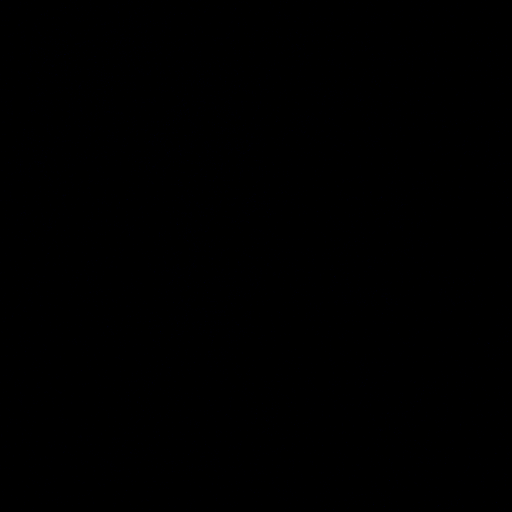

### A_5_DA.tif

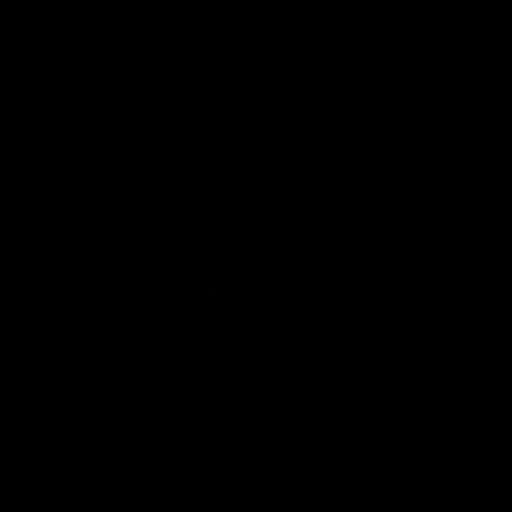

### A_5_DD.tif

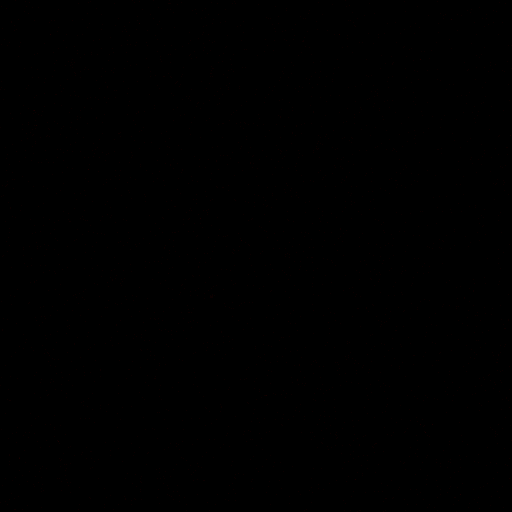

### A_6_AA.tif

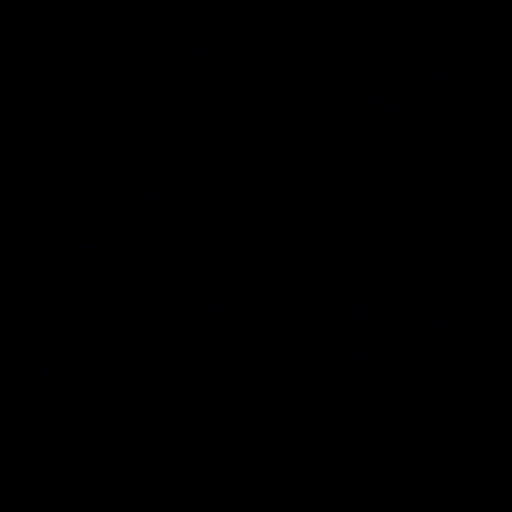

### A_6_DA.tif

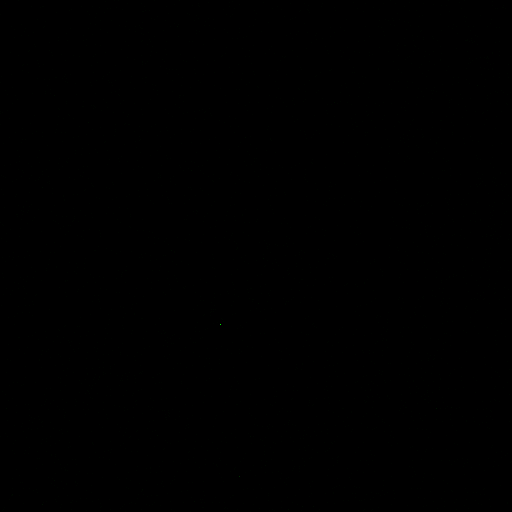

### A_6_DD.tif

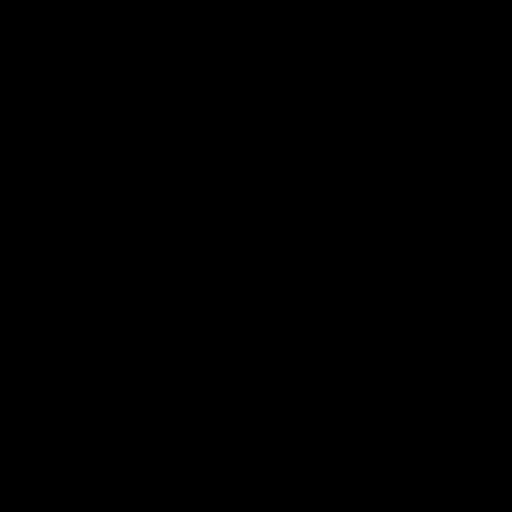

### A_7_AA.tif

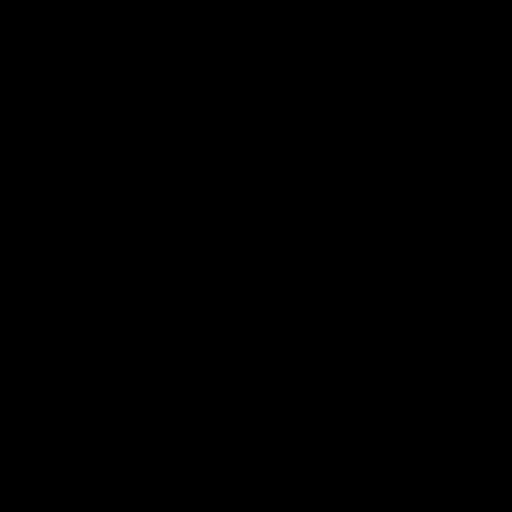

### A_7_DA.tif

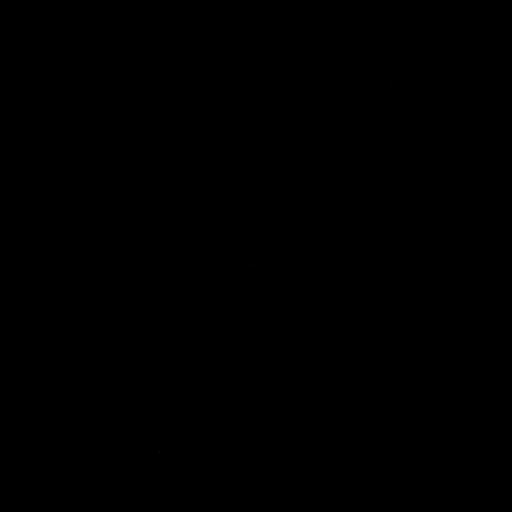

### A_7_DD.tif

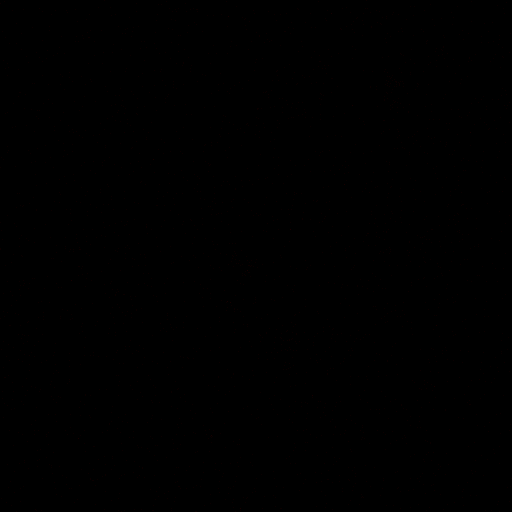

### A_8_AA.tif

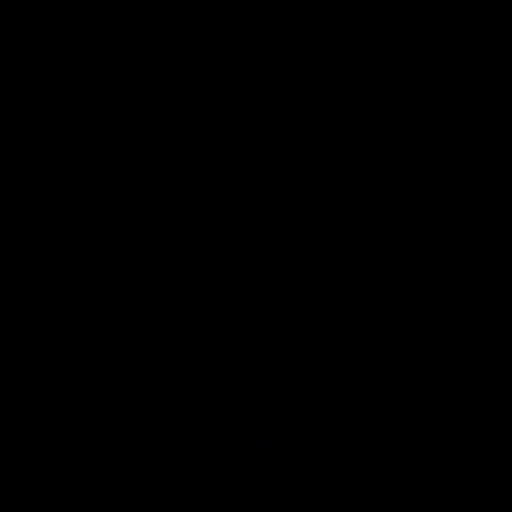

### A_8_DA.tif

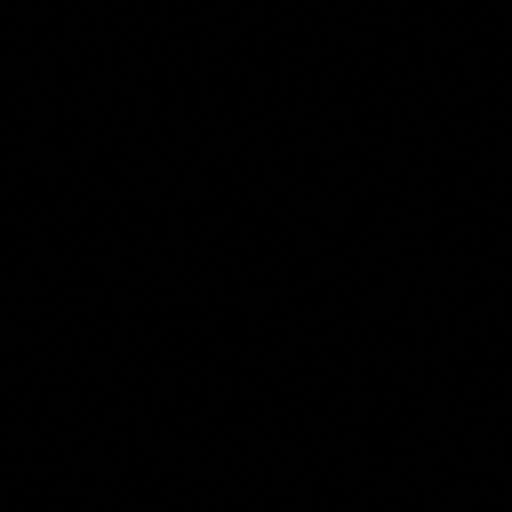

### A_8_DD.tif

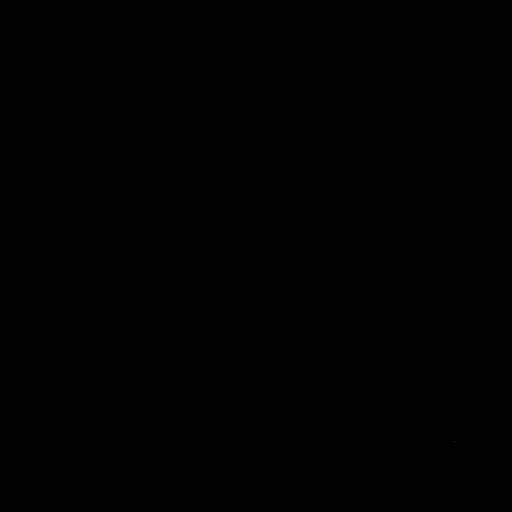

### A_9_AA.tif

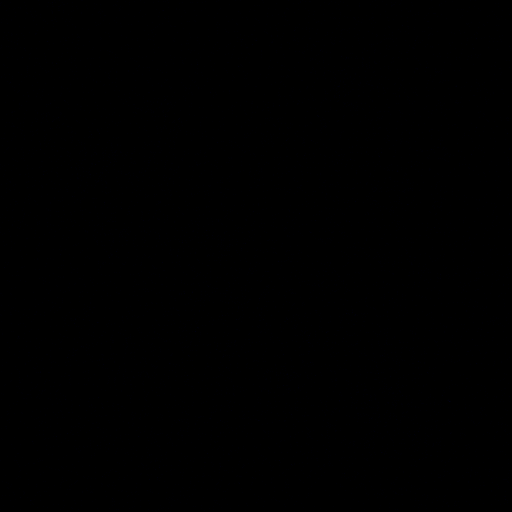

### A_9_DA.tif

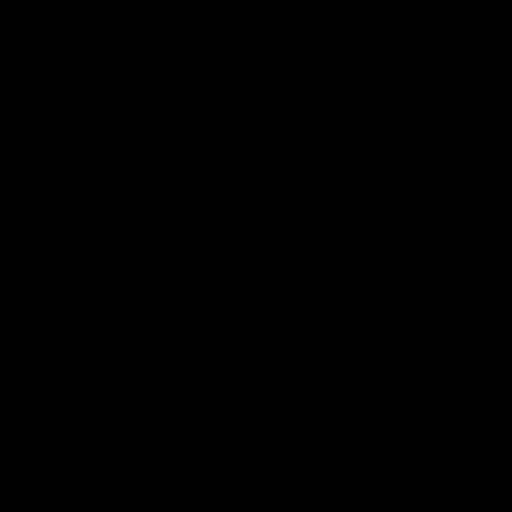

### A_9_DD.tif

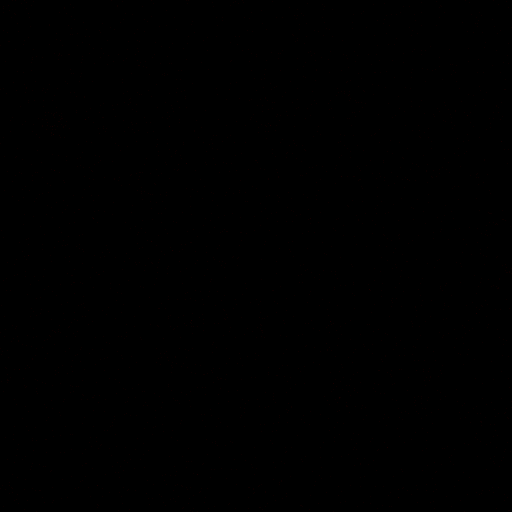

### A_10_AA.tif

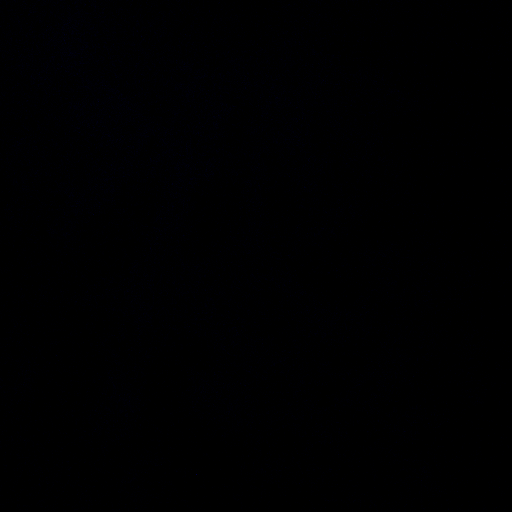

### A_10_DA.tif

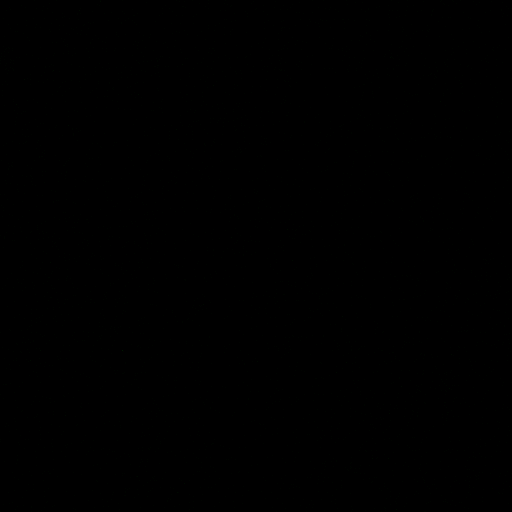

### A_10_DD.tif

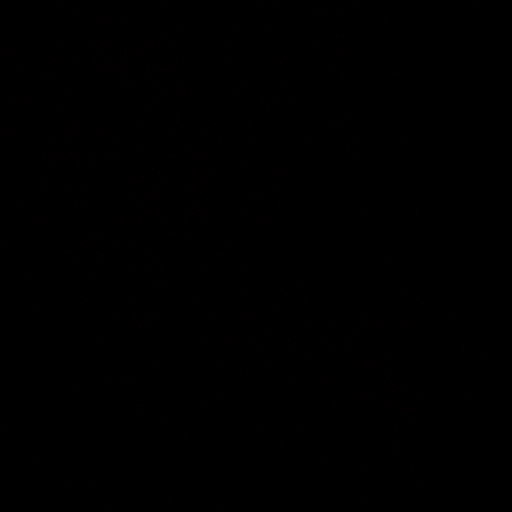

### B_1_AA.tif

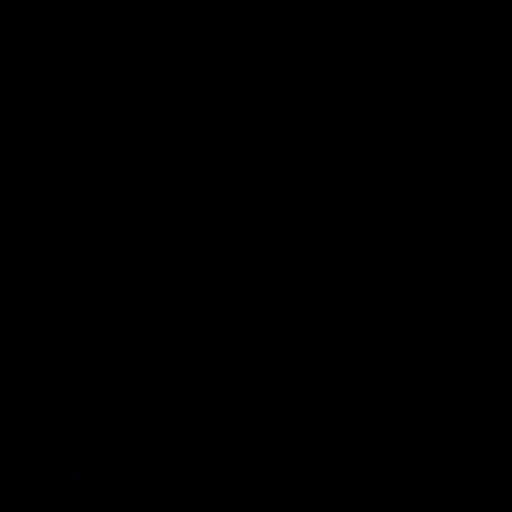

### B_1_DA.tif

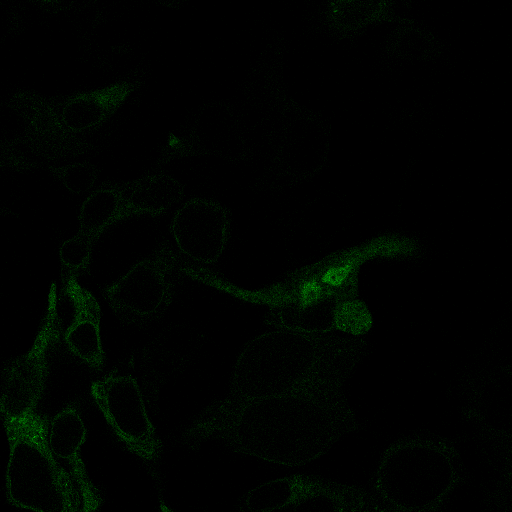

### B_1_DD.tif

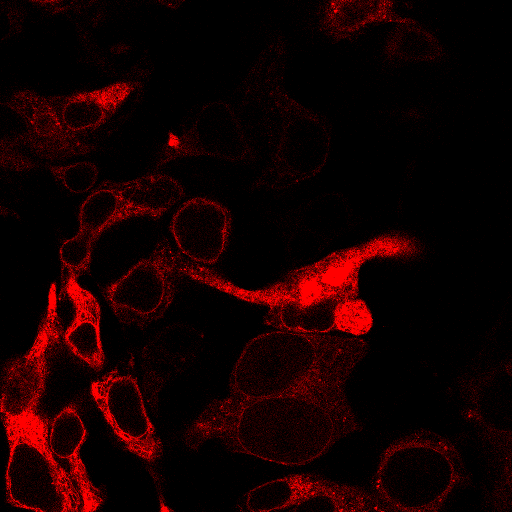

### B_2_AA.tif

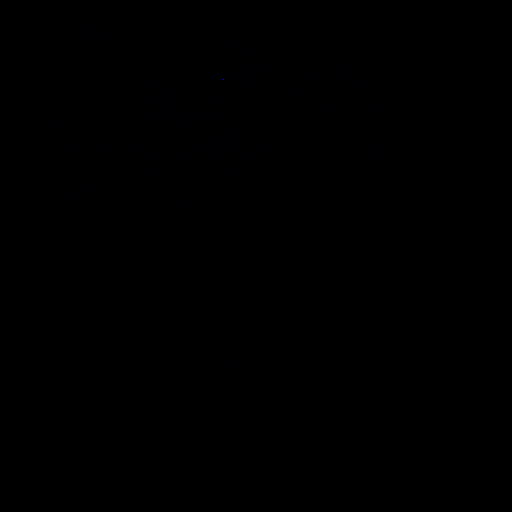

### B_2_DA.tif

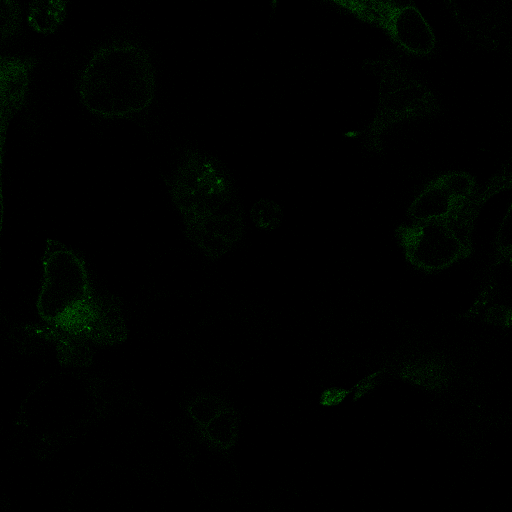

### B_2_DD.tif

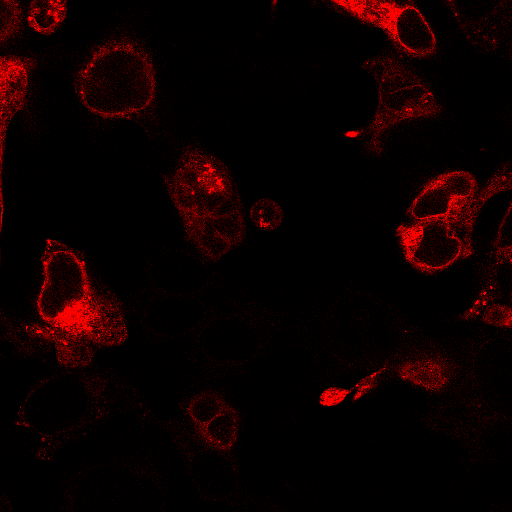

### B_3_AA.tif

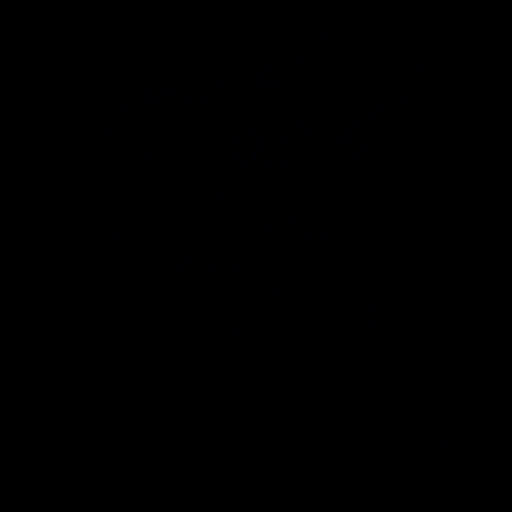

### B_3_DA.tif

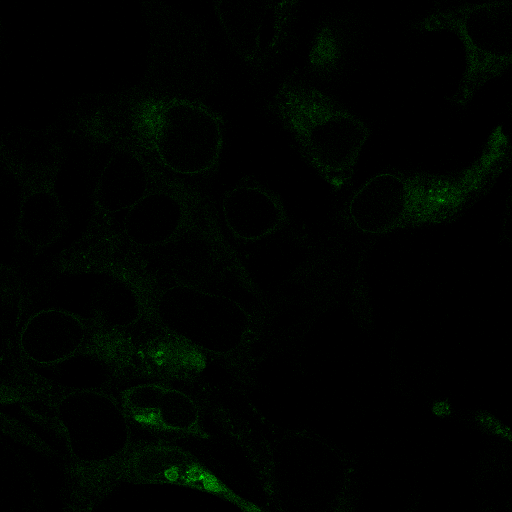

### B_3_DD.tif

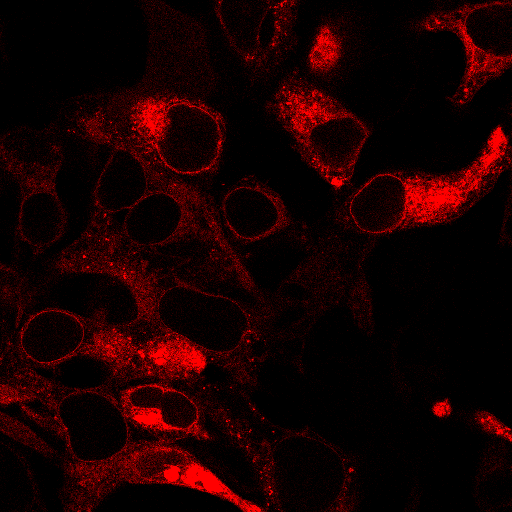

### B_4_AA.tif

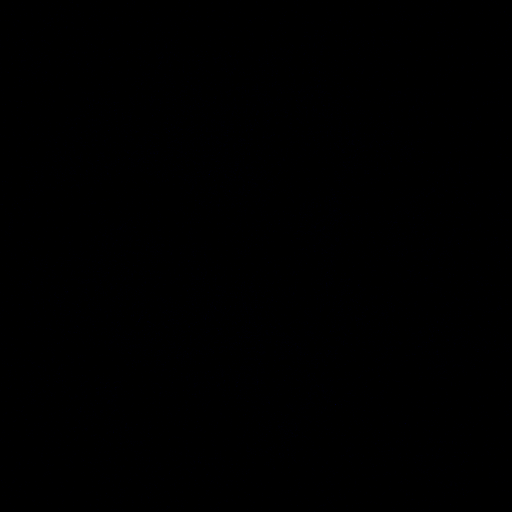

### B_4_DA.tif

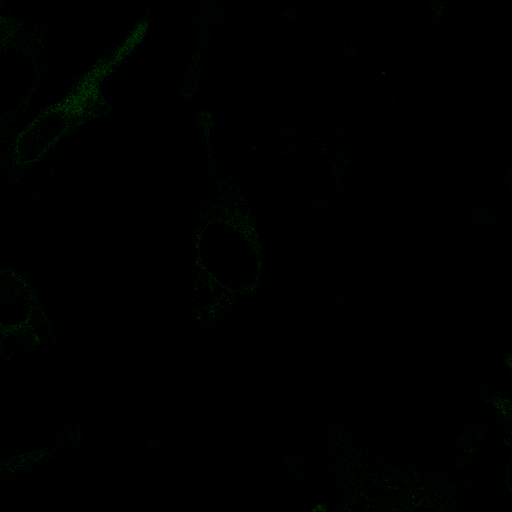

### B_4_DD.tif

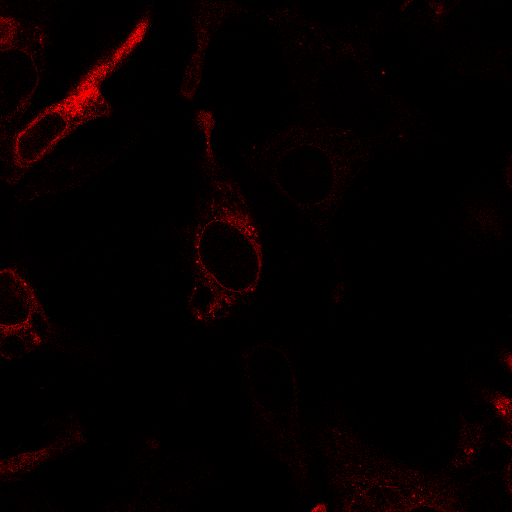

### B_5_AA.tif

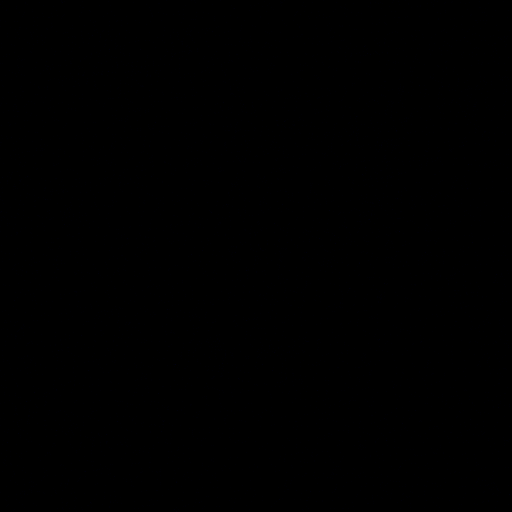

### B_5_DA.tif

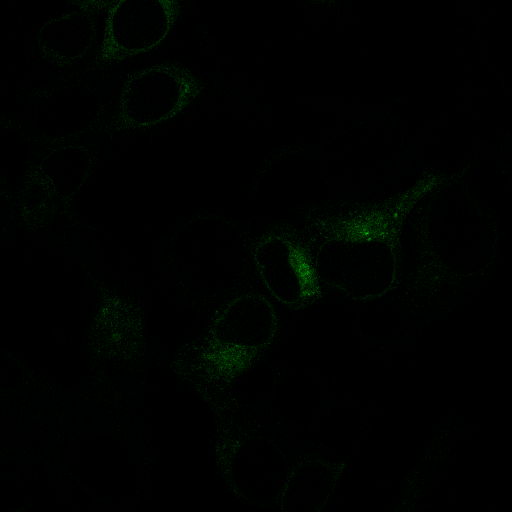

### B_5_DD.tif

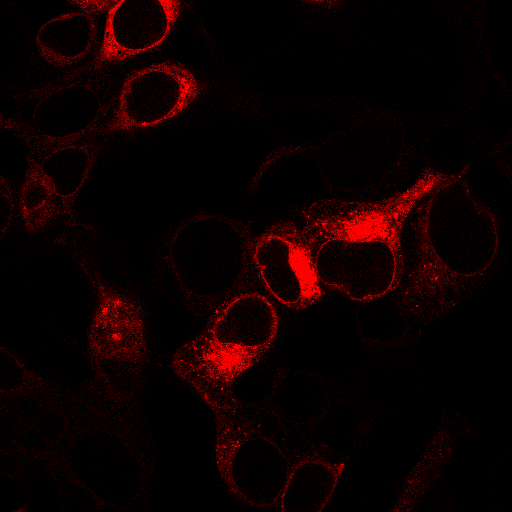

### B_6_AA.tif

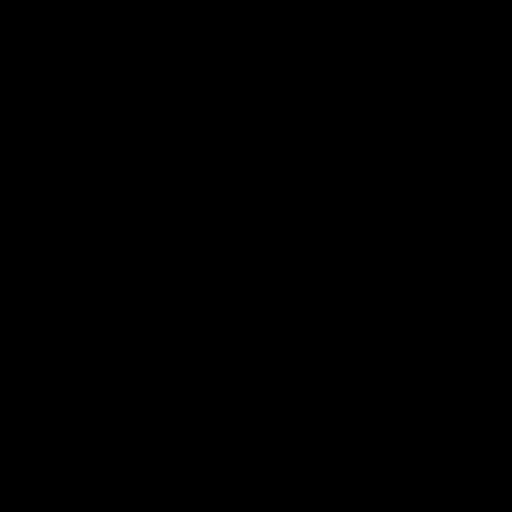

### B_6_DA.tif

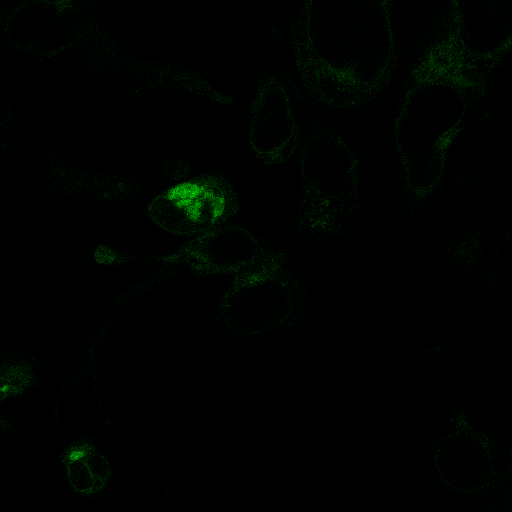

### B_6_DD.tif

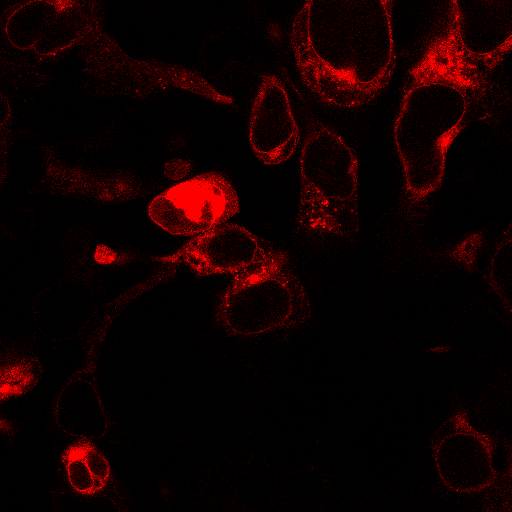

### B_7_AA.tif

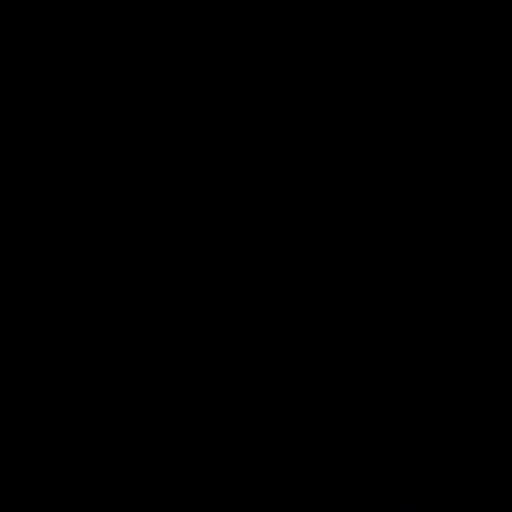

### B_7_DA.tif

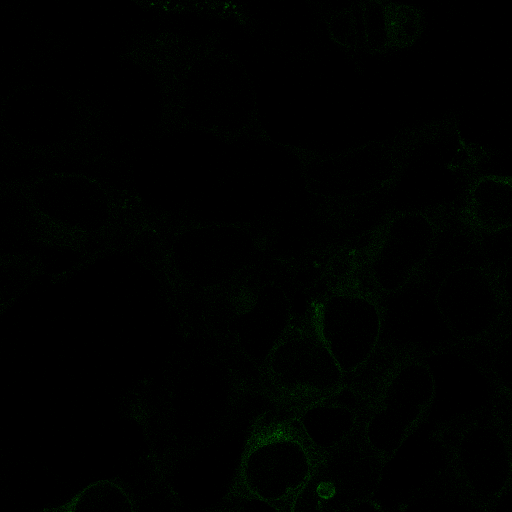

### B_7_DD.tif

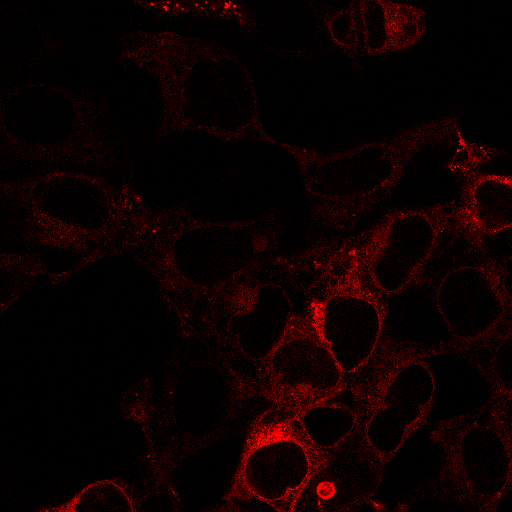

### B_8_AA.tif

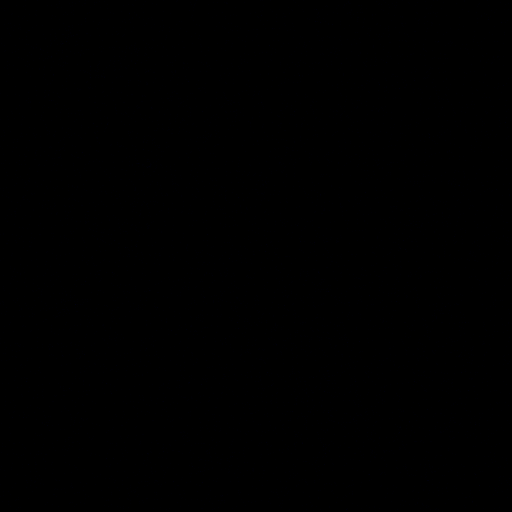

### B_8_DA.tif

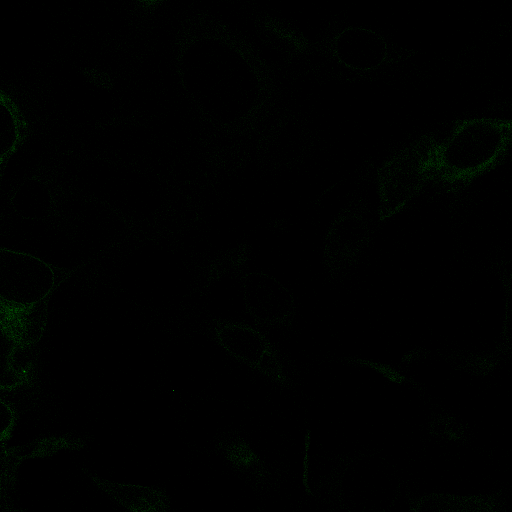

### B_8_DD.tif

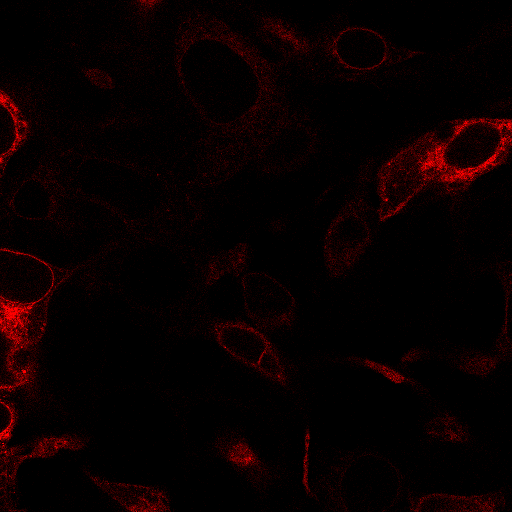

### B_9_AA.tif

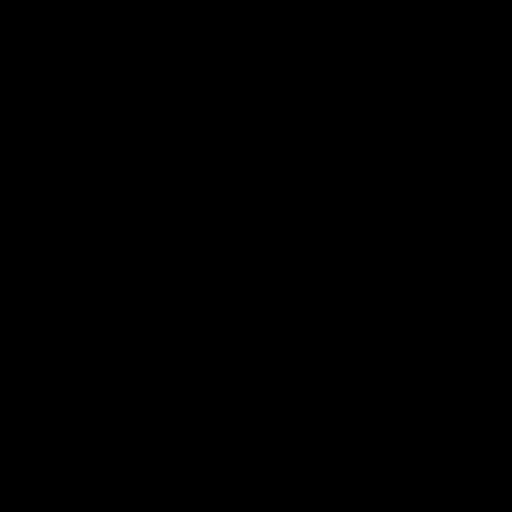

### B_9_DA.tif

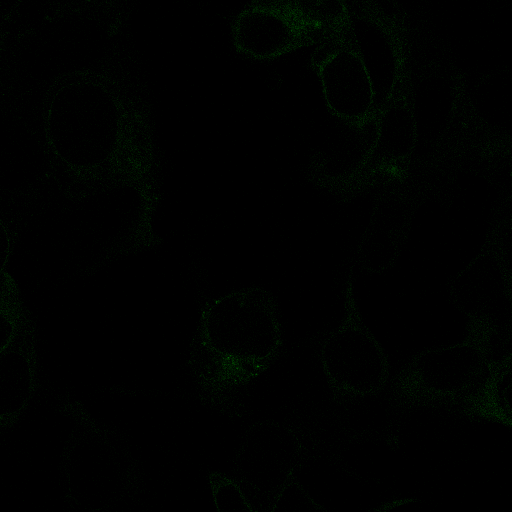

### B_9_DD.tif

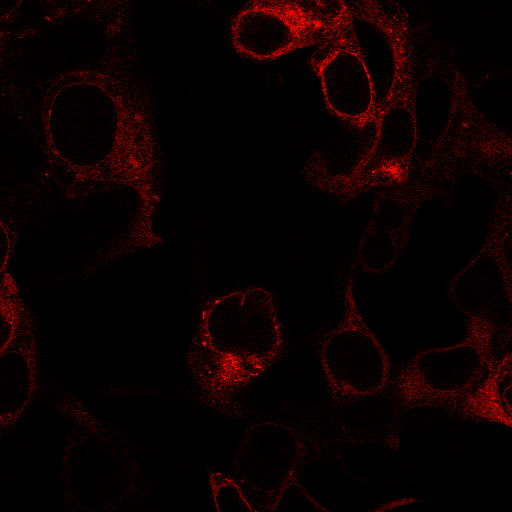

### B_10_AA.tif

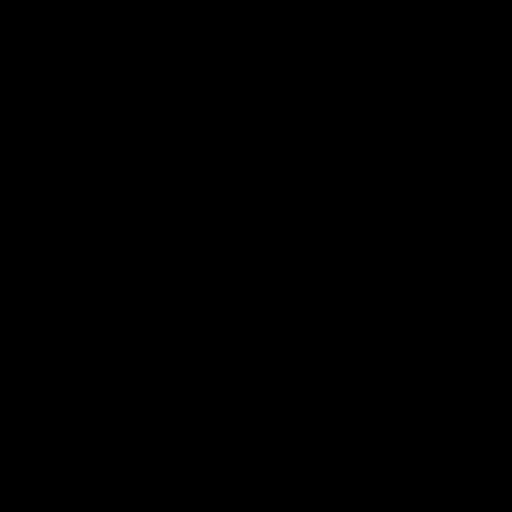

### B_10_DA.tif

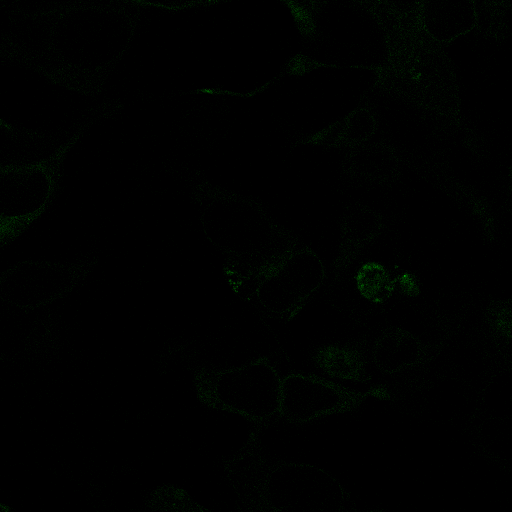

### B_10_DD.tif

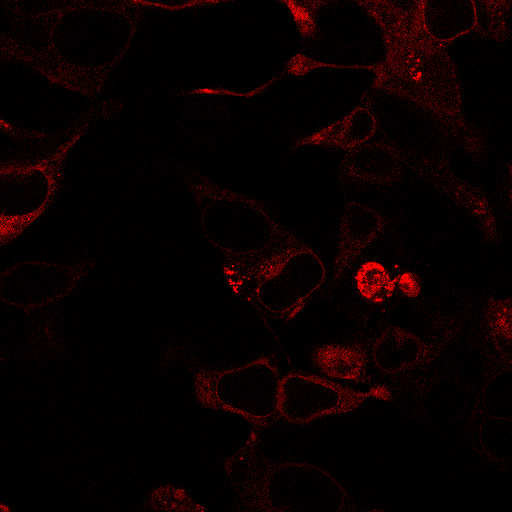

### C_1_AA.tif

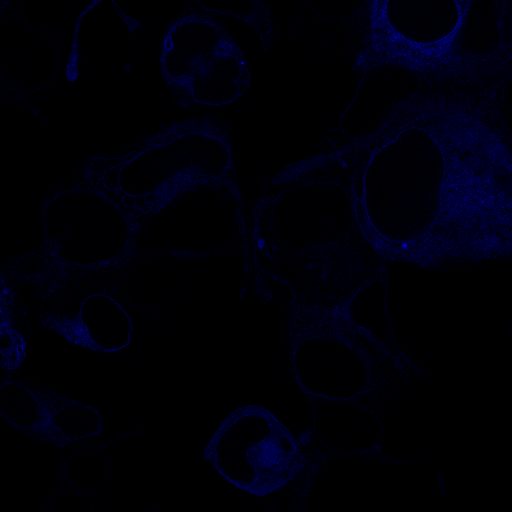

### C_1_DA.tif

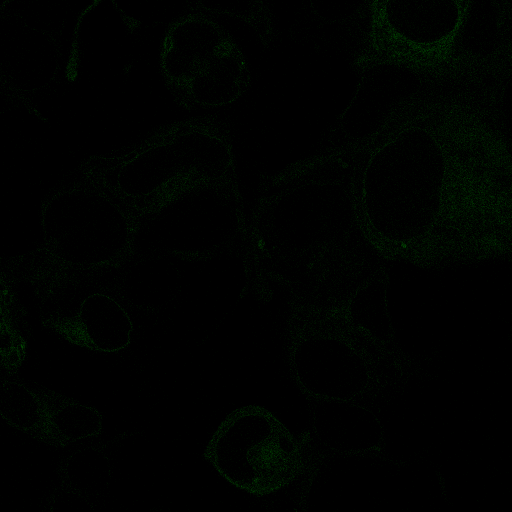

### C_1_DD.tif

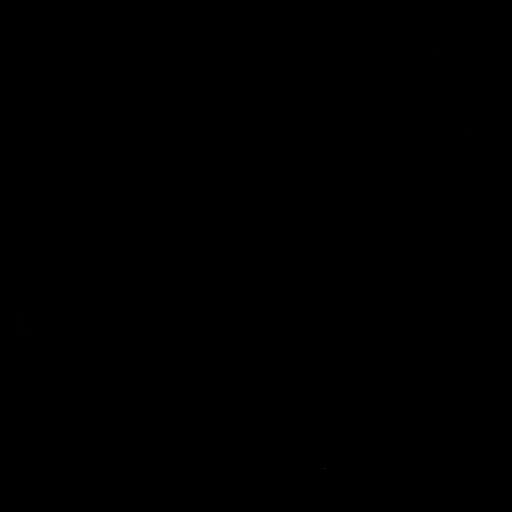

### C_2_AA.tif

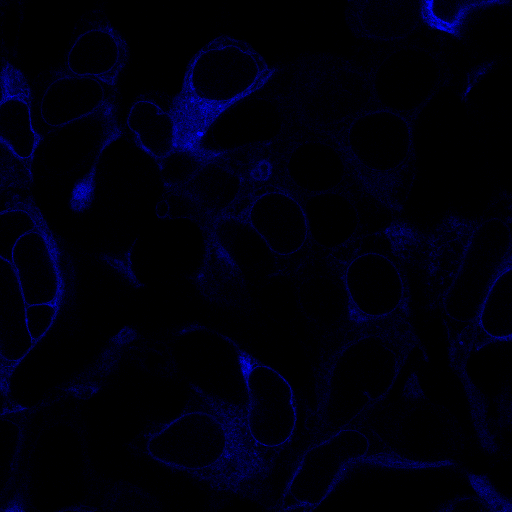

### C_2_DA.tif

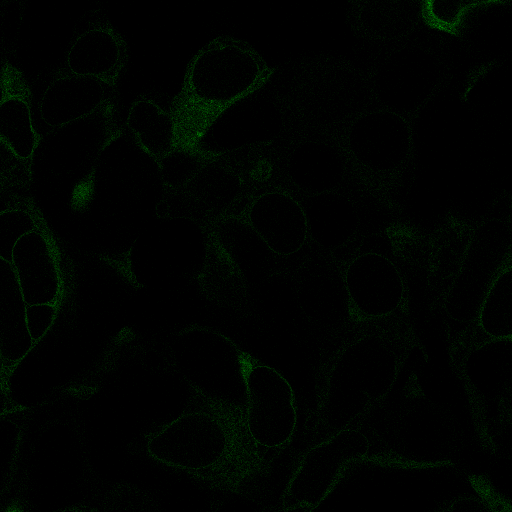

### C_2_DD.tif

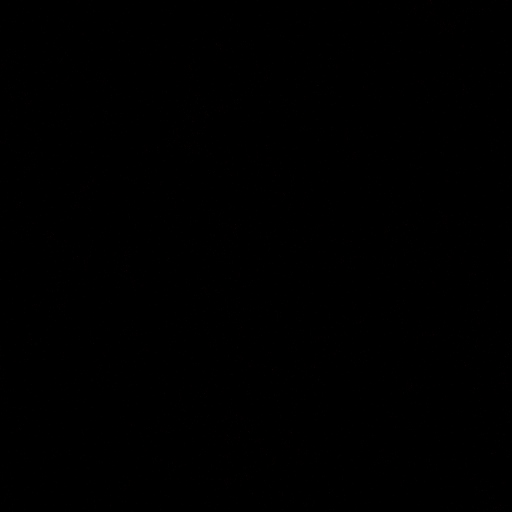

### C_3_AA.tif

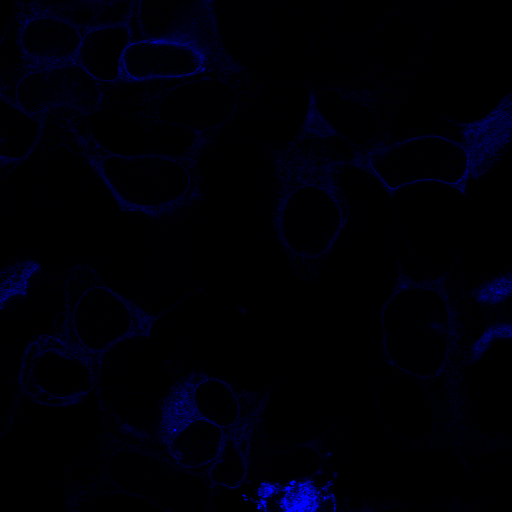

### C_3_DA.tif

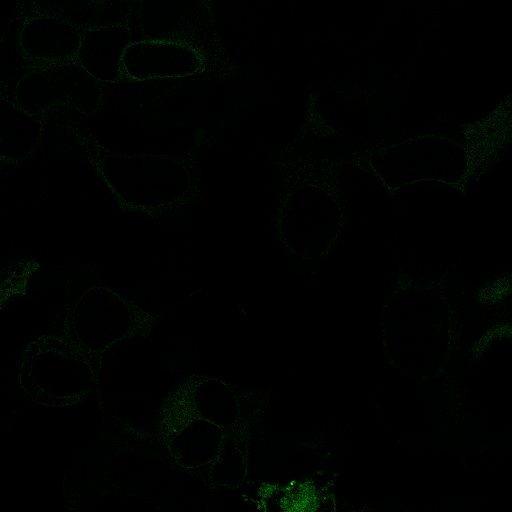

### C_3_DD.tif

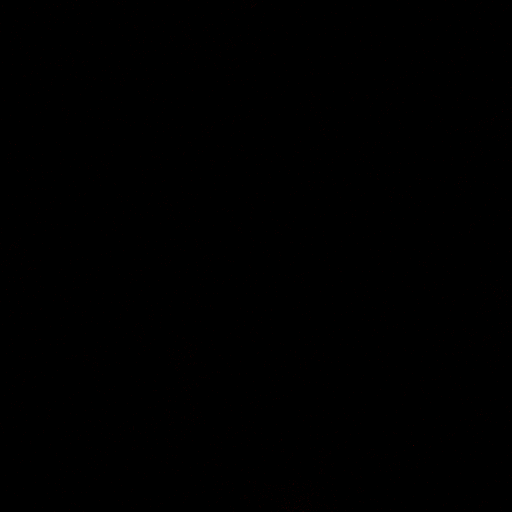

### C_4_AA.tif

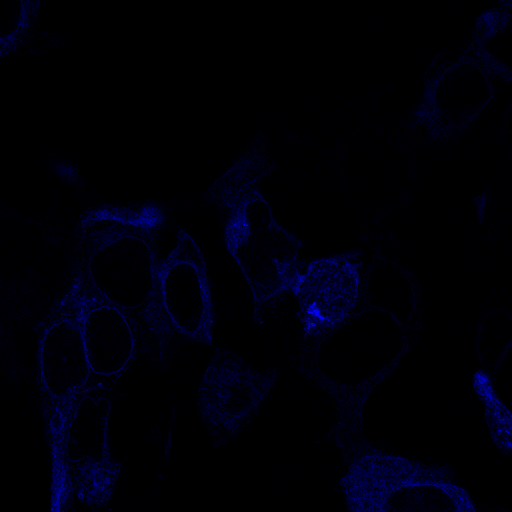

### C_4_DA.tif

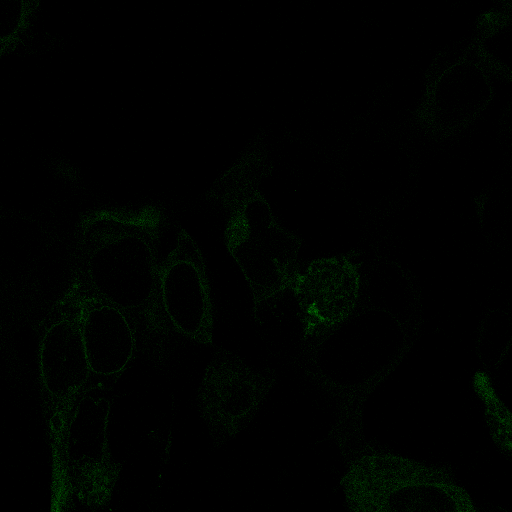

### C_4_DD.tif

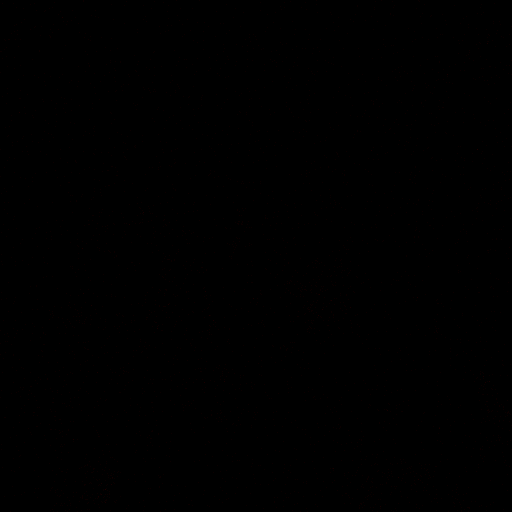

### C_5_AA.tif

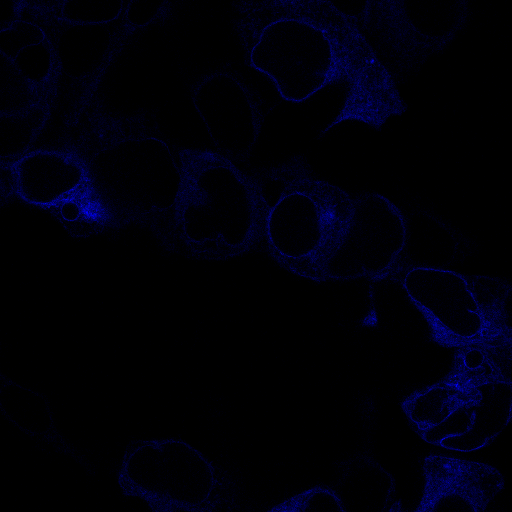

### C_5_DA.tif

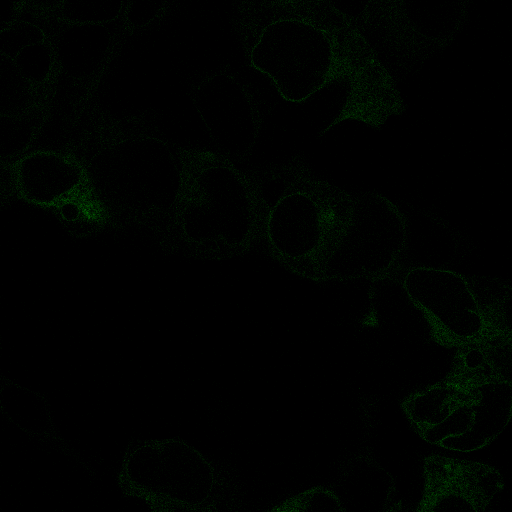

### C_5_DD.tif

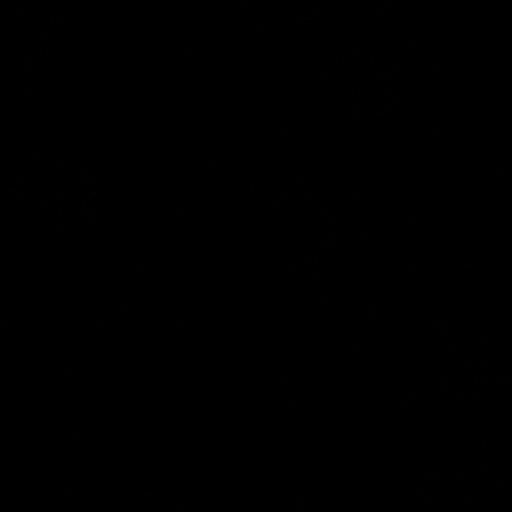

### C_6_AA.tif

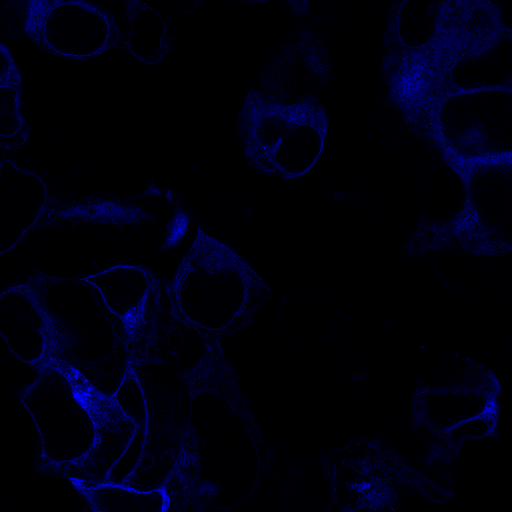

### C_6_DA.tif

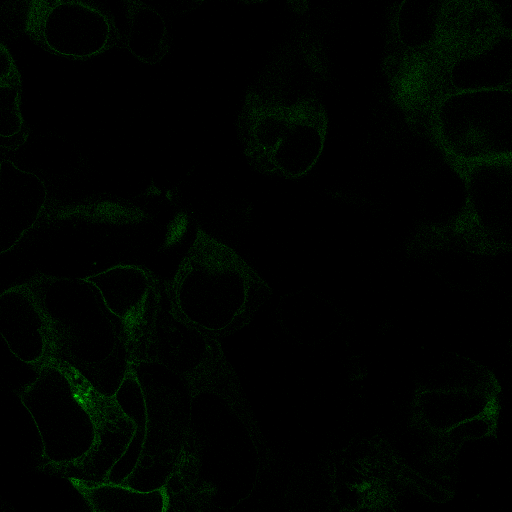

### C_6_DD.tif

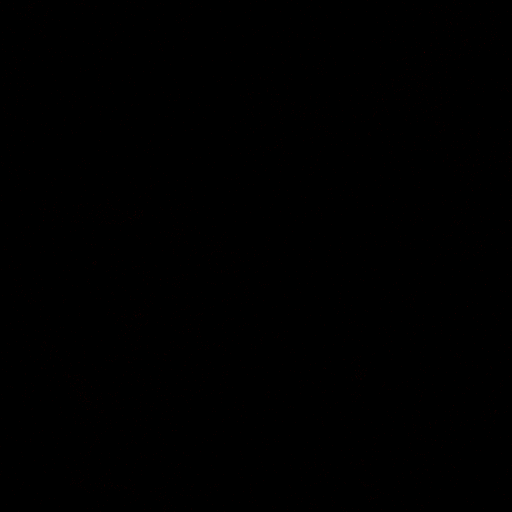

### C_7_AA.tif

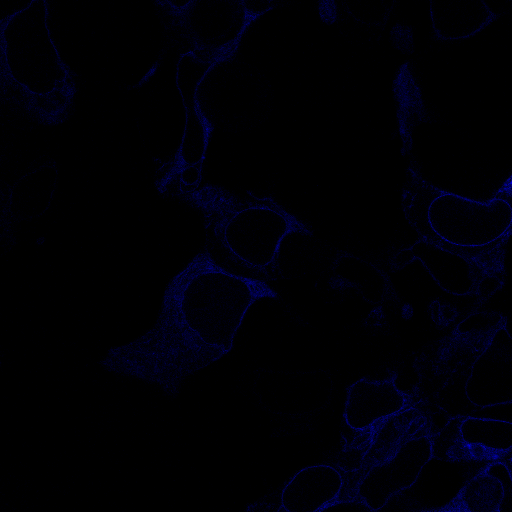

### C_7_DA.tif

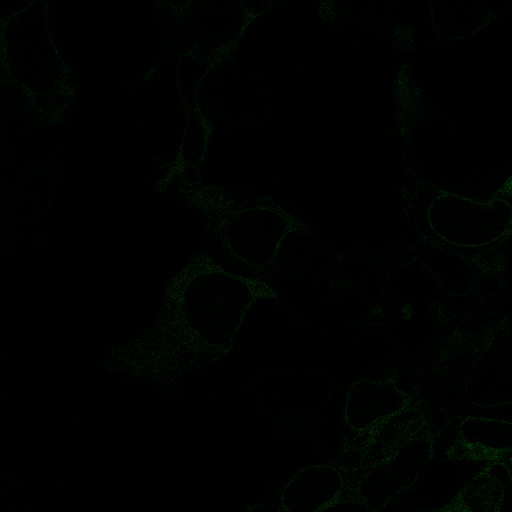

### C_7_DD.tif

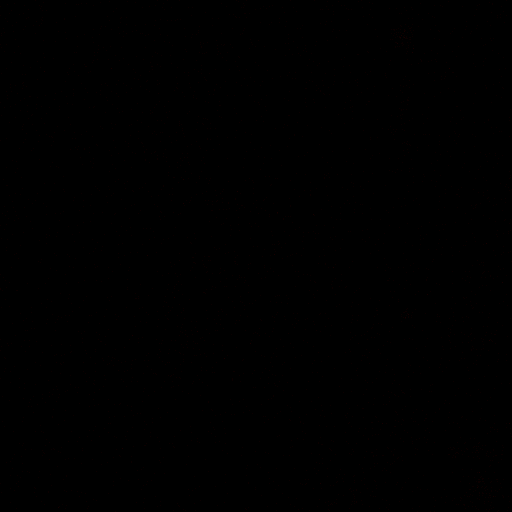

### C_8_AA.tif

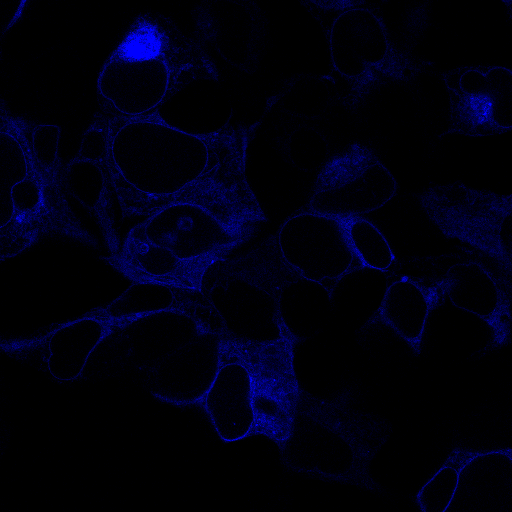

### C_8_DA.tif

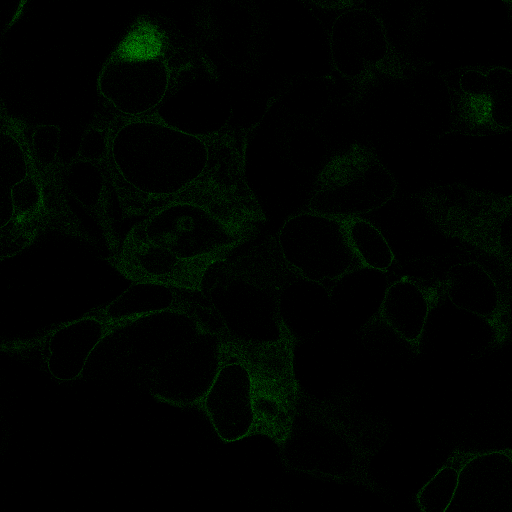

### C_8_DD.tif

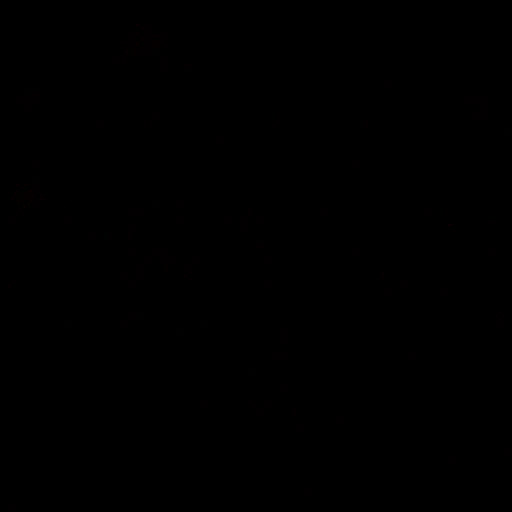

### C_9_AA.tif

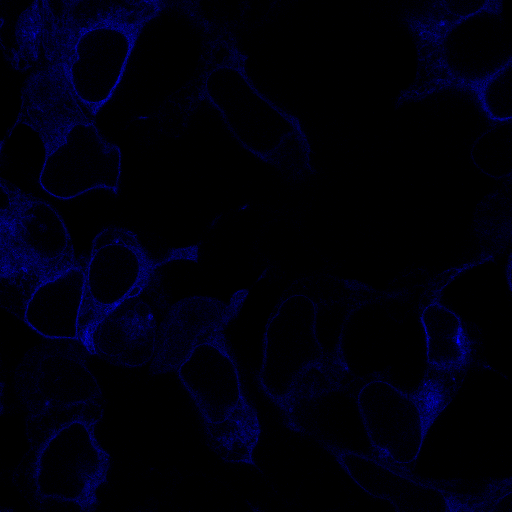

### C_9_DA.tif

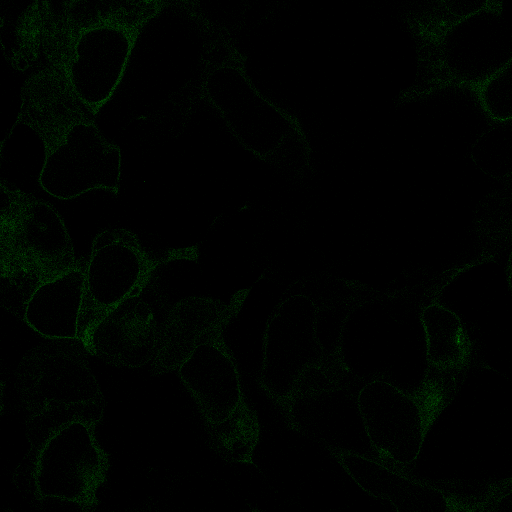

### C_9_DD.tif

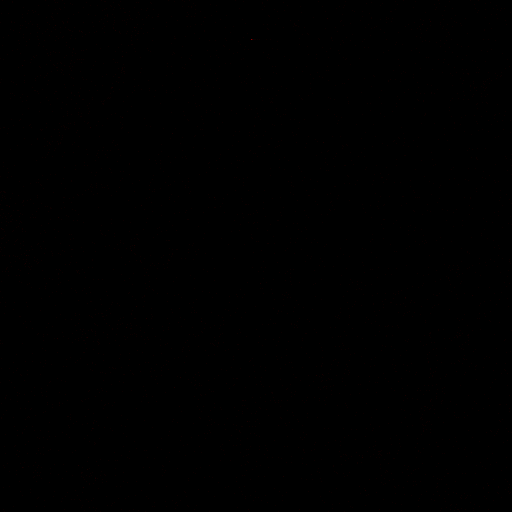

### C_10_AA.tif

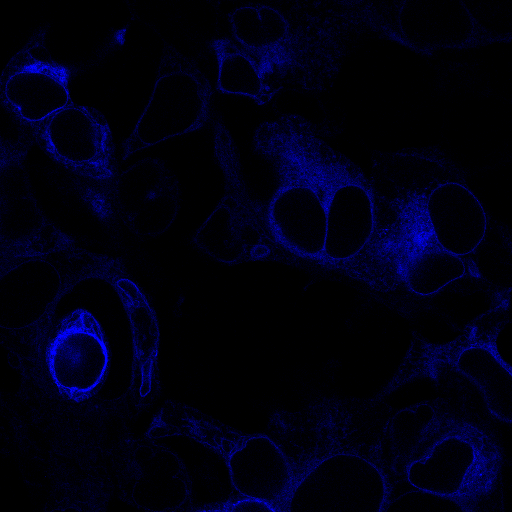

### C_10_DA.tif

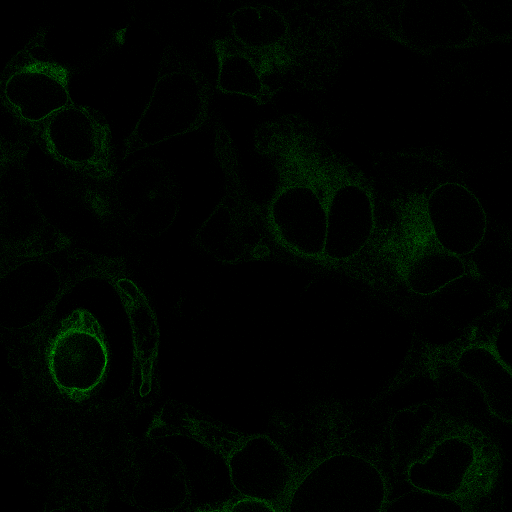

### C_10_DD.tif

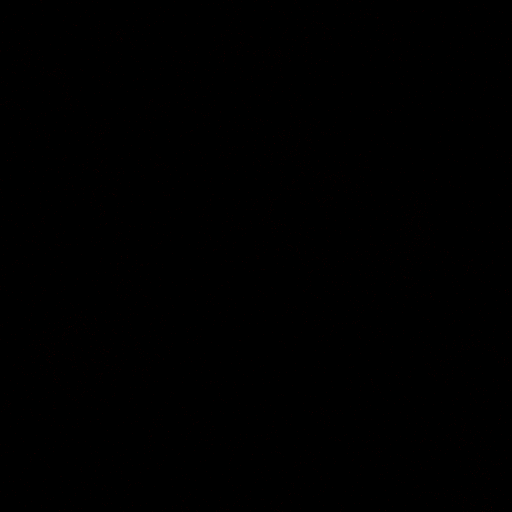

### D_1_AA.tif

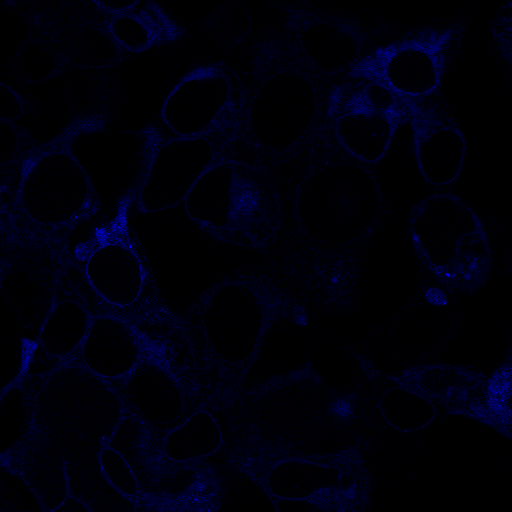

### D_1_DA.tif

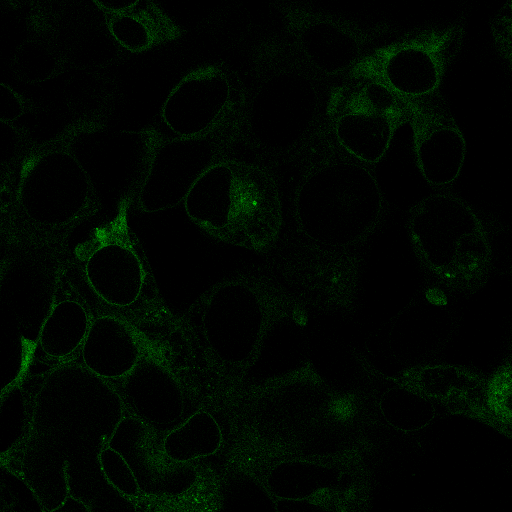

### D_1_DD.tif

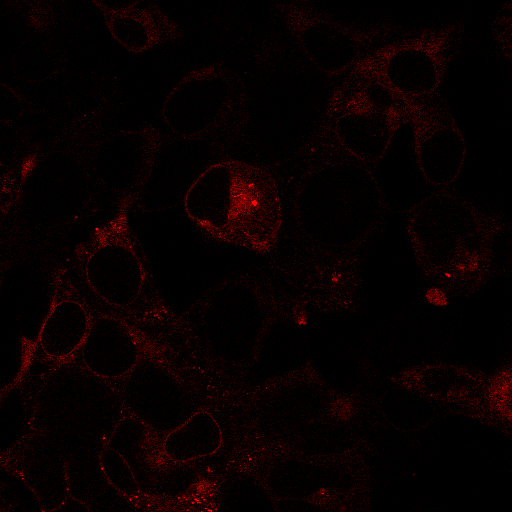

### D_2_AA.tif

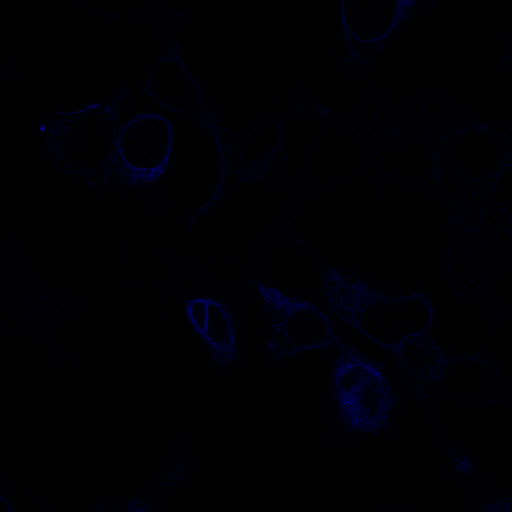

### D_2_DA.tif

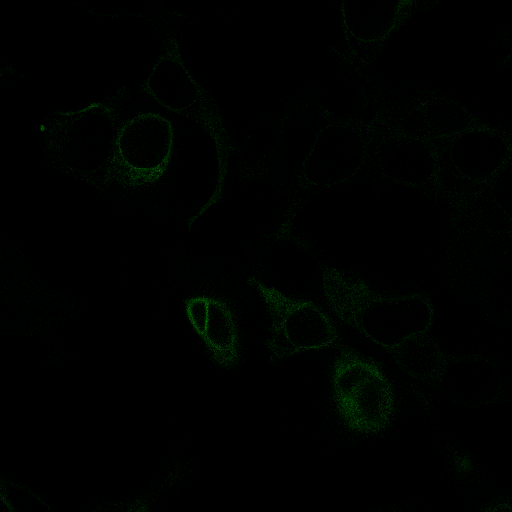

### D_2_DD.tif

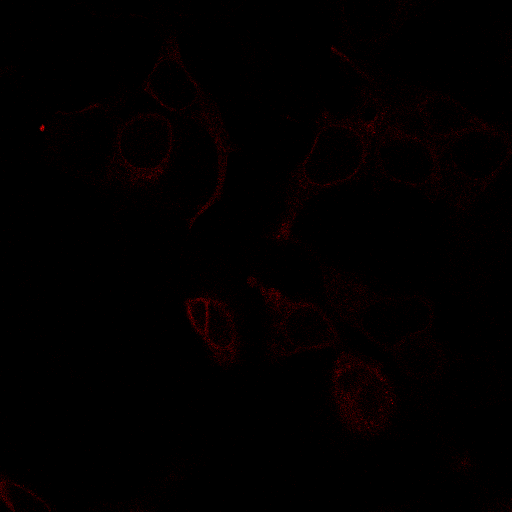

### D_3_AA.tif

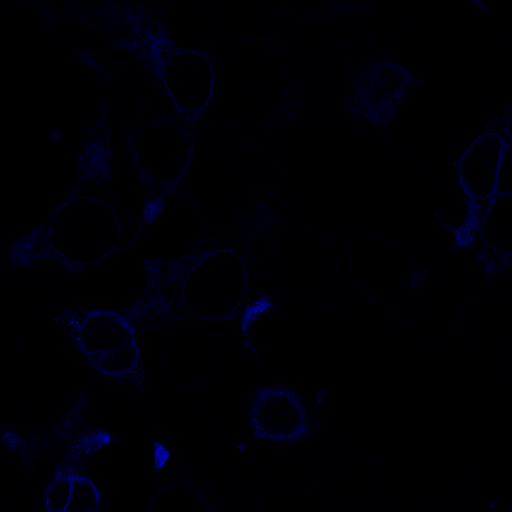

### D_10_AA.tif

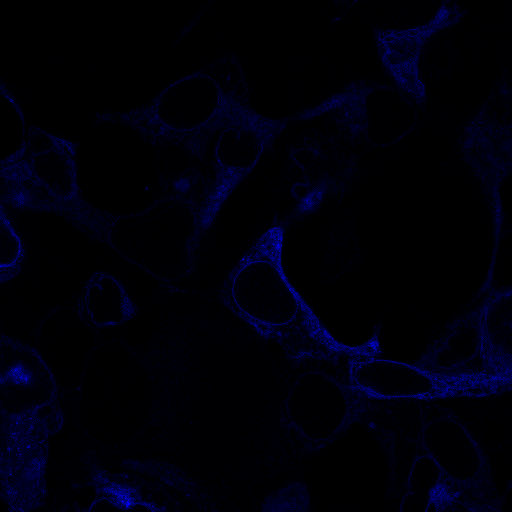

### D_10_DA.tif

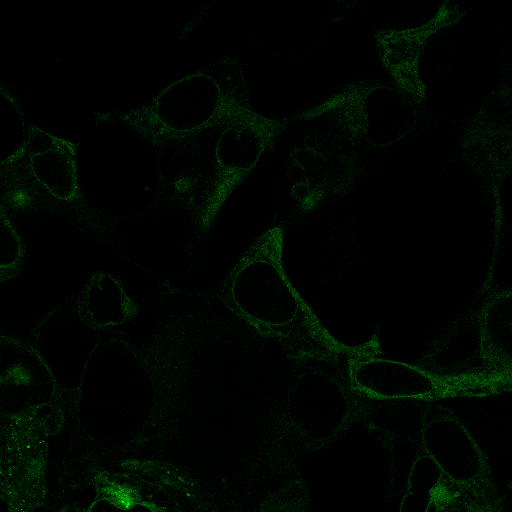

### D_10_DD.tif

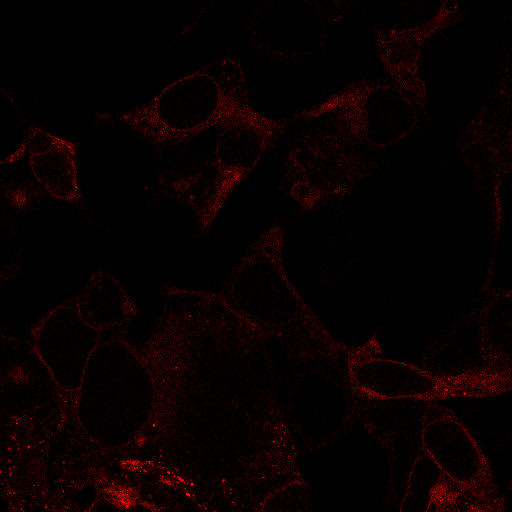
