## Supplementary Information for "Elucidation and refinement of synthetic receptor mechanisms"

**Contents:**

Supplementary Figures 1–31

Supplementary Tables 1–5

Supplementary Notes 1–5

References cited in this document

### SUPPLEMENTARY FIGURES

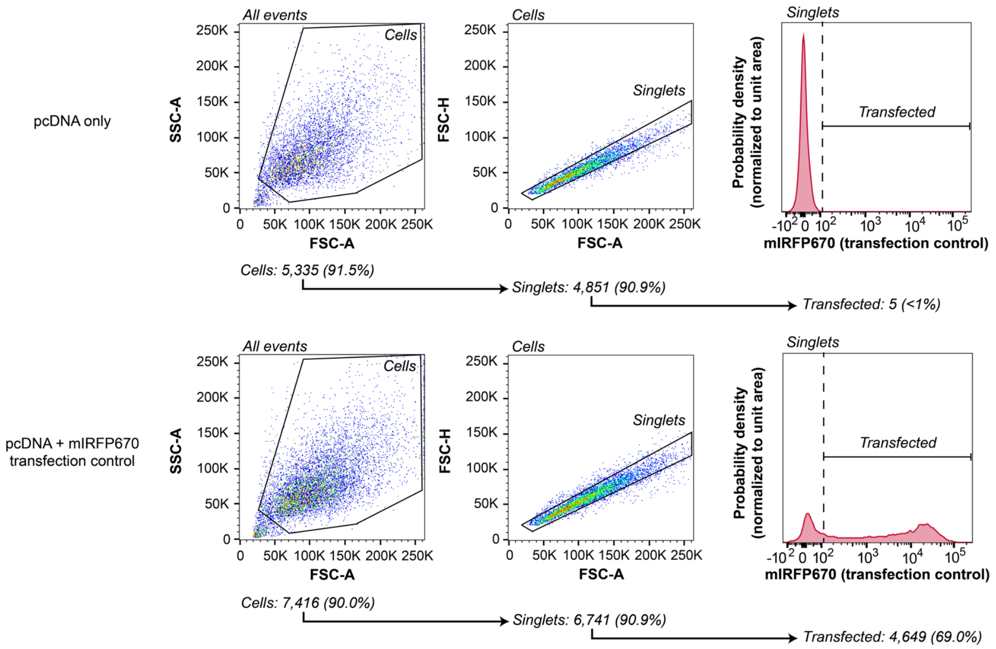


**Supplementary Fig. 1 Flow cytometry gating for transfected cells for FRET.** (**a**) The plots illustrate the flow cytometry gating strategy used to identify single transfected cells for a sample of cells transfected with pcDNA vector only, or pcDNA vector combined with 200 ng of a transfection control plasmid (here, constitutively expressed miRFP670). For both samples, the total mass of DNA transfected was held at 400 ng. In this gating procedure, HEK293FT cells were identified based on the FSC-A vs. SSC-A profile. From this population, singlets were identified based on the FSC-A vs. FSC-H profile. The transfected population was defined as all single cells with miRFP670 fluorescence intensity greater than the sample of single cells transfected with pcDNA only. The transfection gate was set such that it does not encompass more than 1% of the pcDNA only-transfected (non-fluorescent) singlet population. miRFP670 was used as a transfection control in the majority of FRET experiments in this study. For functional signaling assays, EBFP2 was used as a transfection control and a similar gating strategy was employed.

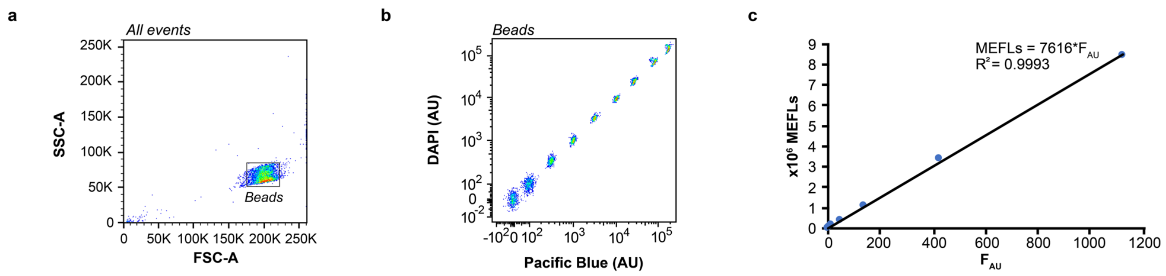


**Supplementary Fig. 2 Calibration of fluorescence intensities to absolute units.** UltraRainbow Calibration Particles (URCP) have nine fluorescent bead populations. (**a**) Beads were identified based on the FSC-A vs. SSC-A profile. (**b**) For each experiment, two fluorescent channels were used to identify the nine bead populations. (**c**) The mean fluorescence intensity (MFI) of each population in the FITC channel (in which EYFP was also recorded) in arbitrary units ($F_{AU}$) was recorded and plotted against manufacturer-provided values for fluorophores per bead for each population (Molecules of Equivalent Fluorescein, MEFLs). To generate the calibration curve, a linear regression was performed with the constraint that the *y*-intercept equals zero. This calibration is done for each experiment, and then exported MFI values (which have arbitrary fluorescence units) are converted to absolute units using the multiplier obtained from the regression. An analogous process is used for experiments with an mCherry-based reporter (AU in PE-Texas Red channel is converted to Molecules of Equivalent PE-Texas Red, MEPTRs).

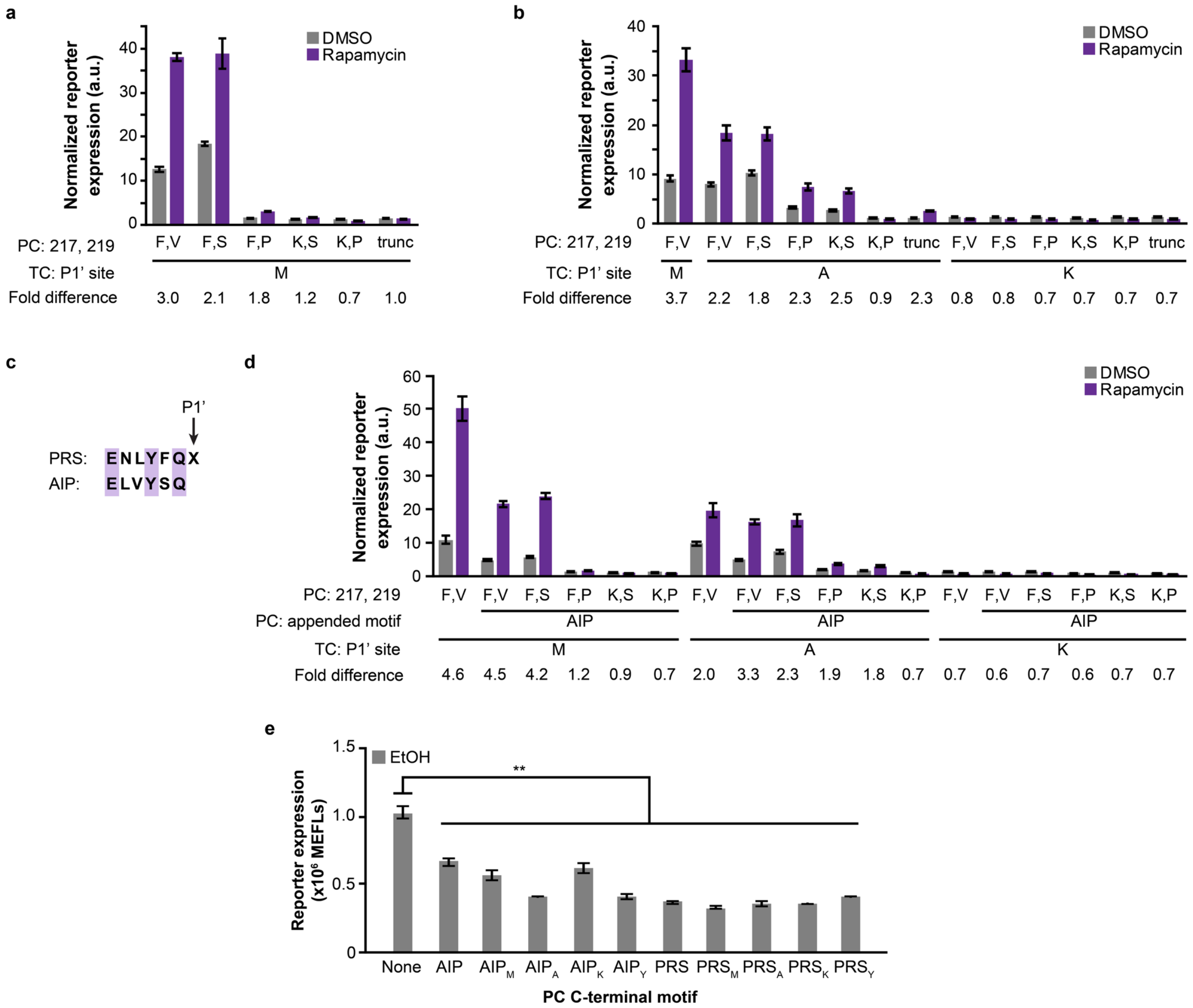


**Supplementary Fig. 3 Investigation of variants for the PRS P1' site, protease active site, and protease C-terminus.** (**a–c**) Combinations of mutant variants for the P1' site of the protease recognition sequence (PRS: ENLYFQX; X is a variable amino acid) on the TC and for positions 217 and 219 of the protease on the PC, with and without the autoinhibitory peptide (AIP: ELVYSQ) on the protease C-terminus. "Trunc" indicates a truncation of the protease after V219. The base case condition has TC with M at the P1' site, PC with F and V at the protease residues 217 and 219, respectively, and treatment with DMSO vehicle. These functional assays utilized a luciferase reporter assay. All conditions are CD28-TMD rapamycin-sensing MESA receptor variants. Reporter expression was calculated from the ratio of inducible Firefly luciferase signal to constitutive *Renilla* luciferase signal for each biological replicate. Quotients were linearly scaled such that the mean of the quotients for a condition transfected with reporter only was equal to 1 a.u. Bars are the means of three biological replicates, and error bars depict S.E.M. Numbers indicate fold difference in reporter expression between samples treated with rapamycin and DMSO. (**d**) This schematic shows the sequence similarity between the PRS and AIP and the location of the P1' position. (**e**) Reporter expression for the samples treated with vehicle (EtOH) shown in **Fig. 1c**. Data are reproduced here with a y-axis scaled to highlight differences between background reporter expression conferred by receptors including the original PC versus PC variants with appended C-terminal AIP and PRS peptides. All variants confer significantly less background reporter expression than receptors with the original PC (one-way ANOVA, ** $p < 0.01$). Statistical analysis is in **Supplementary Note 1**. Bars are the means of three biological replicates, and error bars depict S.E.M.

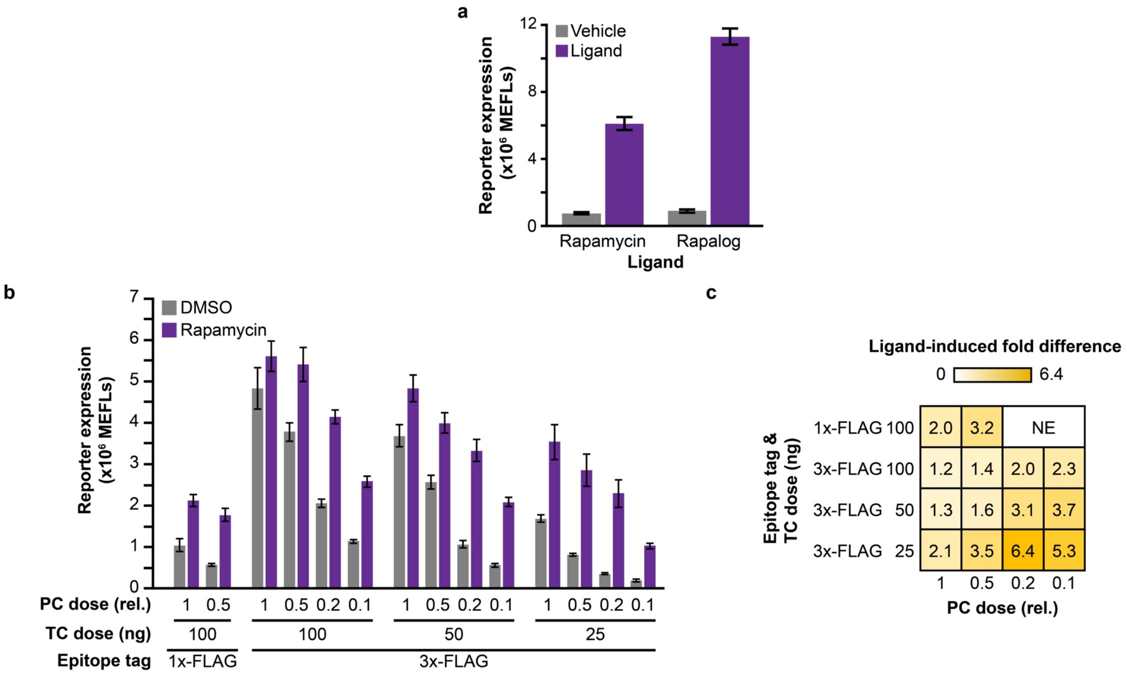


**Supplementary Fig. 4 Comparisons across MESA assays**. (**a**) Comparison between rapalog and rapamycin ligands. The rapalog AP21967 is a synthetic rapamycin analog that binds only to FRB containing the T2098L mutation (which are contained in all reported rapamycin-sensing MESA receptors). Rapalog is less immunosuppressive than rapamycin, due to diminished interference with the endogenous mTOR pathway (1). Rapalog produces higher ligand-induced reporter expression than does rapamycin. We noted some cytotoxicity by microscopy for cells treated with rapamycin, which is consistent with this outcome. The vehicle for rapamycin is 50% DMSO/50% water. The vehicle for rapalog is 100% EtOH. Stock and final working ligand concentrations used in cell culture are listed in **Supplementary Table 1**. (**b,c**) To increase the sensitivity of detection on Western blots and via flow cytometry, we replaced the 1x-FLAG epitope tag on the N-terminus of each MESA chain with a 3x-FLAG tag. The use of 100 ng of plasmid encoding each MESA chain led to higher levels of background and ligand-induced signaling for the 3x-FLAG constructs than for the 1x-FLAG constructs. As this phenomenon is similar to what has been observed when MESA receptors are overexpressed, we performed a combinatorial dose scanning experiment, reducing the plasmid doses of TC and PC (labels show the PC dose relative to the TC dose, where a value of unity denotes equal amounts of each plasmid). The use of 25 ng of 3x-FLAG tagged TC plasmid yielded similar fold differences compared to 1x-FLAG tagged constructs (**c**), though both the background and ligand-induced signaling remained higher with the 3x-FLAG constructs. Abbreviation: NE, not evaluated. Bars are the means of three biological replicates, and error bars depict S.E.M.

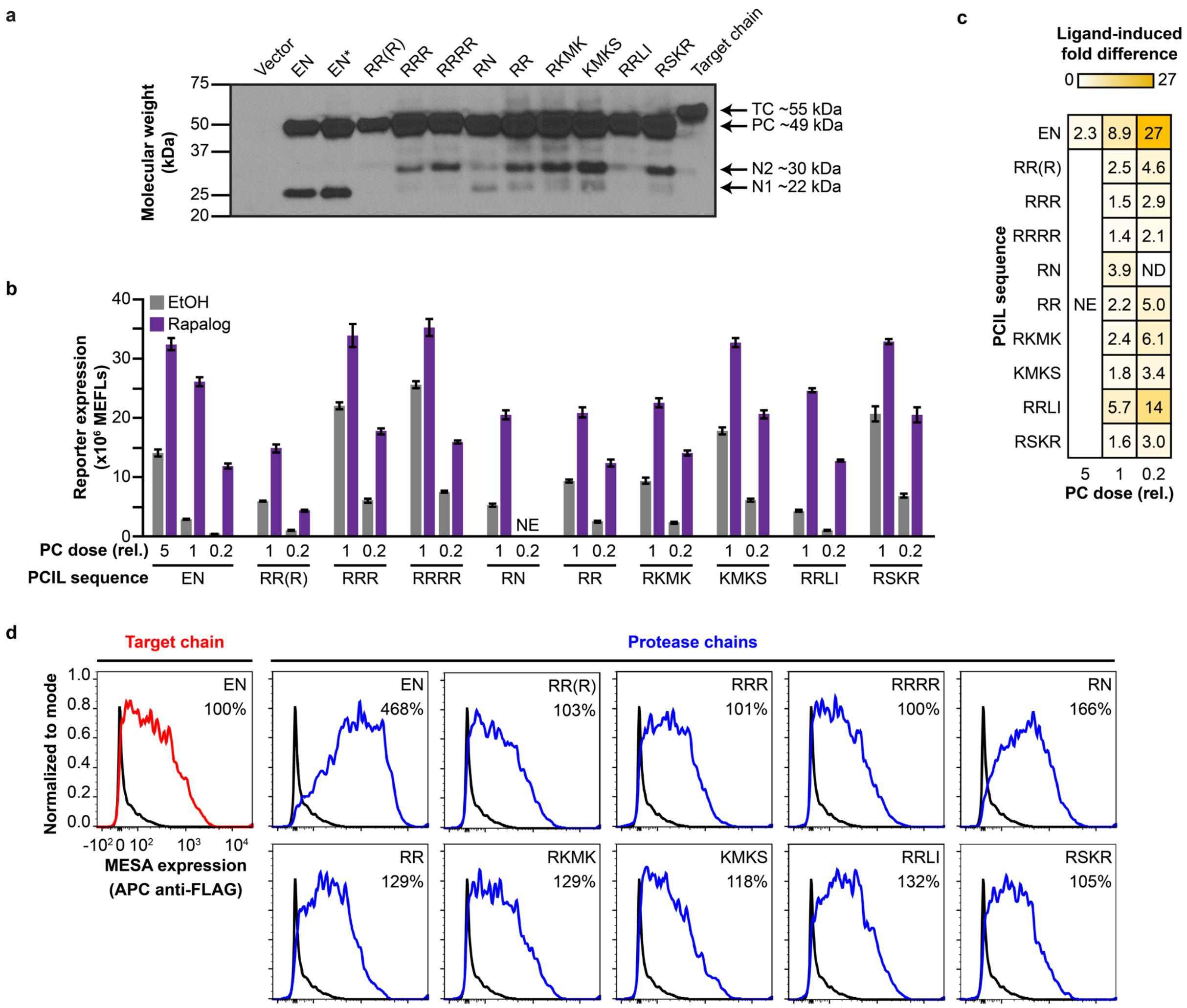


**Supplementary Fig. 5 Investigating the role of PCIL on PC stability**. (**a**) Based on a prior observation that many native juxtamembrane domains are positively charged, we substituted the PCIL with linkers comprising varying numbers of arginine residues or sequences from native receptors (EN is the original PCIL; EN* contains the original PCIL and the D81N mutation in the TEVp; the parenthetical R in the fourth lane indicates that this R replaces the first amino acid TEVp coding sequence). While in many cases a diminishment of the originally observed cleavage product (N1) was observed, we also observed new cleavage products (N2) with many of the constructs. (**b,c**) In a functional assay, with 25 ng of each 3x-FLAG tagged MESA chain plasmid, PCIL substitution led to decreased receptor performance through increased background. As this outcome was consistent with what would be expected if the number of intact PCs per cell were increased (far left condition), we decreased the PC plasmid dose by 5-fold (to 5 ng) while holding the TC plasmid dose constant (at 25 ng). Though this change did result in lower background and ligand-induced signaling, only one case (RRLI) exhibited fold differences greater than that of the original construct (EN); however, the new outcome was not ideal as it came at a cost of decreased ligand-induced signaling. The data shown here include the data reported in **Fig. 1f** in addition to samples with adjusted DNA doses. Bars are the means of three biological replicates, and error bars depict S.E.M. Abbreviation: NE, not evaluated. (**d**) Surface staining profiles of cells expressing 1x-FLAG tagged MESA PCs with various PCIL sequences. All PCIL sequences investigated led to decreased expression relative to the original construct (EN). Percentages indicate PC expression relative to TC expression (the left plot). A stained, vector-only (no MESA chain) sample is shown in each plot as a black-lined histogram.

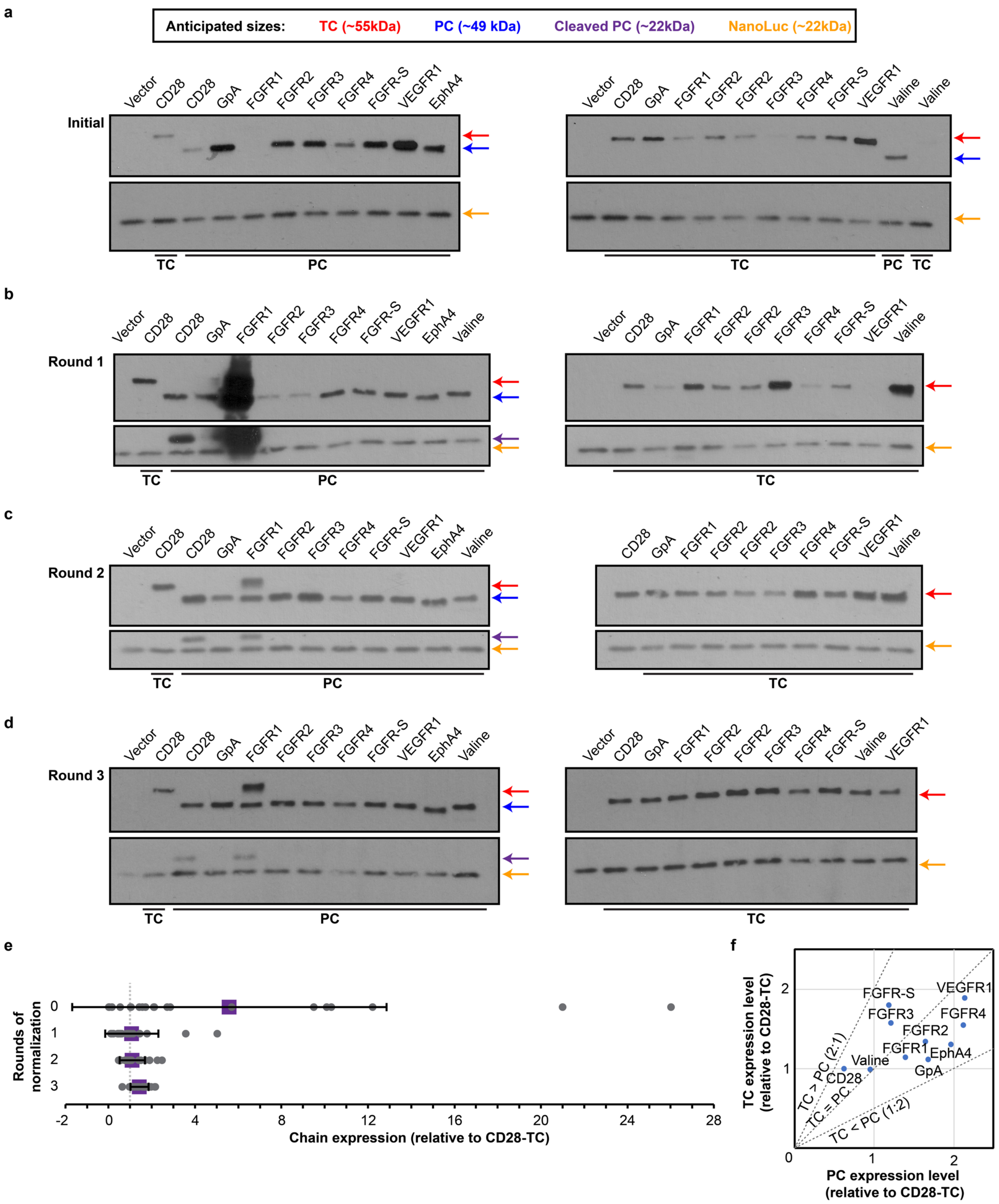
a
Anticipated sizes: TC (~55kDa) PC (~49 kDa) Cleaved PC (~22kDa) NanoLuc (~22kDa)
Initial
Vector CD28 CD28 GpA FGFR1 FGFR2 FGFR3 FGFR4 FGFR-S VEGFR1 EphA4
TC PC
Vector CD28 GpA FGFR1 FGFR2 FGFR2 FGFR3 FGFR4 FGFR-S VEGFR1 Valine Valine
TC PC TC
b
Round 1
Vector CD28 CD28 GpA FGFR1 FGFR2 FGFR3 FGFR4 FGFR-S VEGFR1 EphA4 Valine
TC PC
Vector CD28 GpA FGFR1 FGFR2 FGFR2 FGFR3 FGFR4 FGFR-S VEGFR1 Valine
TC
c
Round 2
Vector CD28 CD28 GpA FGFR1 FGFR2 FGFR3 FGFR4 FGFR-S VEGFR1 EphA4 Valine
TC PC
CD28 GpA FGFR1 FGFR2 FGFR2 FGFR3 FGFR4 FGFR-S VEGFR1 Valine
TC
d
Round 3
Vector CD28 CD28 GpA FGFR1 FGFR2 FGFR3 FGFR4 FGFR-S VEGFR1 EphA4 Valine
TC PC
Vector CD28 GpA FGFR1 FGFR2 FGFR2 FGFR3 FGFR4 FGFR-S Valine VEGFR1
TC
e
Rounds of normalization
0
1
2
3
-2 0 4 8 12 16 20 24 28
Chain expression (relative to CD28-TC)
f
TC expression level (relative to CD28-TC)
PC expression level (relative to CD28-TC)
0 1 2
1 2
FGFR-S
VEGFR1
FGFR3
FGFR4
FGFR2
FGFR1
EphA4
Valine
GpA
CD28
TC > PC (2:1)
TC = PC
TC < PC (1:2)

**Supplementary Fig. 6 Normalization of protein expression for MESA containing various TMDs.** (**a**) Western blot of MESA containing various TMDs, using the same plasmid dose for each construct. Successive rounds of normalization were performed by quantifying the expression of each chain (normalizing first to the intensity from the co-transfected 3x-FLAG tagged NanoLuciferase in each sample to control for loading and transfection efficiency, and then normalizing to the normalized intensity from the CD28-TMD TC as an internal control) and varying the plasmid dose in each successive round. Details are provided in **Methods**. Anticipated sizes for applicable bands are indicated by the colored arrows, as described by the legend, which also applies to **b–d**. Western blots after (**b**) one round, (**c**) two rounds, and (**d**) three rounds of normalization. The Western blot in **d** is the same as in **Fig. 2b**. Plasmid doses in **d** were used in **Fig. 2c,d**. (**e**) The sequential normalization procedure resulted in more similar protein expression levels across with each round of quantification. Each chain's relative expression level is represented by a grey dot. The mean relative expression level of all chains is depicted by the purple square. Error bars represent the S.D. The vertical dotted grey line at 1 relative unit corresponds to the expression level of the CD28-TMD TC base case. (**f**) Ratio of TC:PC expression for each chain, using the final plasmid doses. All constructs were expressed within a 1:2 and 2:1 TC:PC ratio.

**a**

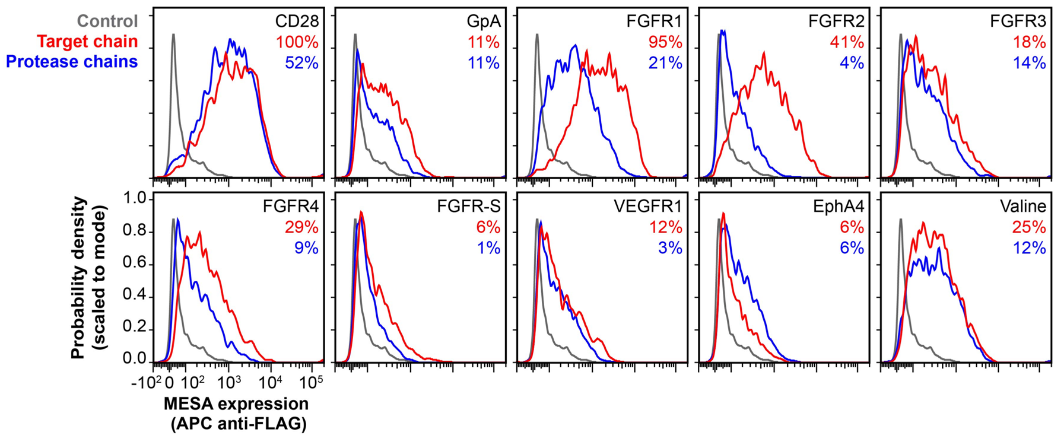


**b**

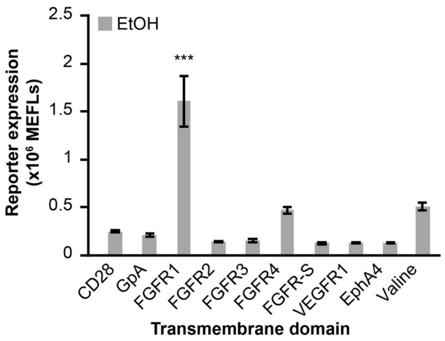


**Supplementary Fig. 7 Surface stain and background signal of MESA containing various TMDs.** (**a**) Surface stain of MESA chains containing various TMDs, using the plasmid doses that yielded approximately equal whole-cell protein expression in **Supplementary Fig. 6d**. Percentages are the receptor surface expression relative to the CD28-TMD TC. The control sample in all plots is a stained, vector-only (no MESA) sample. (**b**) Reporter expression for the samples treated with vehicle (EtOH) shown in **Fig. 2c**. Data are reproduced here with a y-axis scaled to highlight differences between background reporter expression conferred by pairs of receptors with different matched TMDs. Comparing all vehicle-treated TMDs, only FGFR1 exhibits significantly greater background signal than the others (one-way ANOVA, *** $p < 0.001$). Statistical analysis is in **Supplementary Note 1**. Bars are the means of three biological replicates, and error bars depict S.E.M.

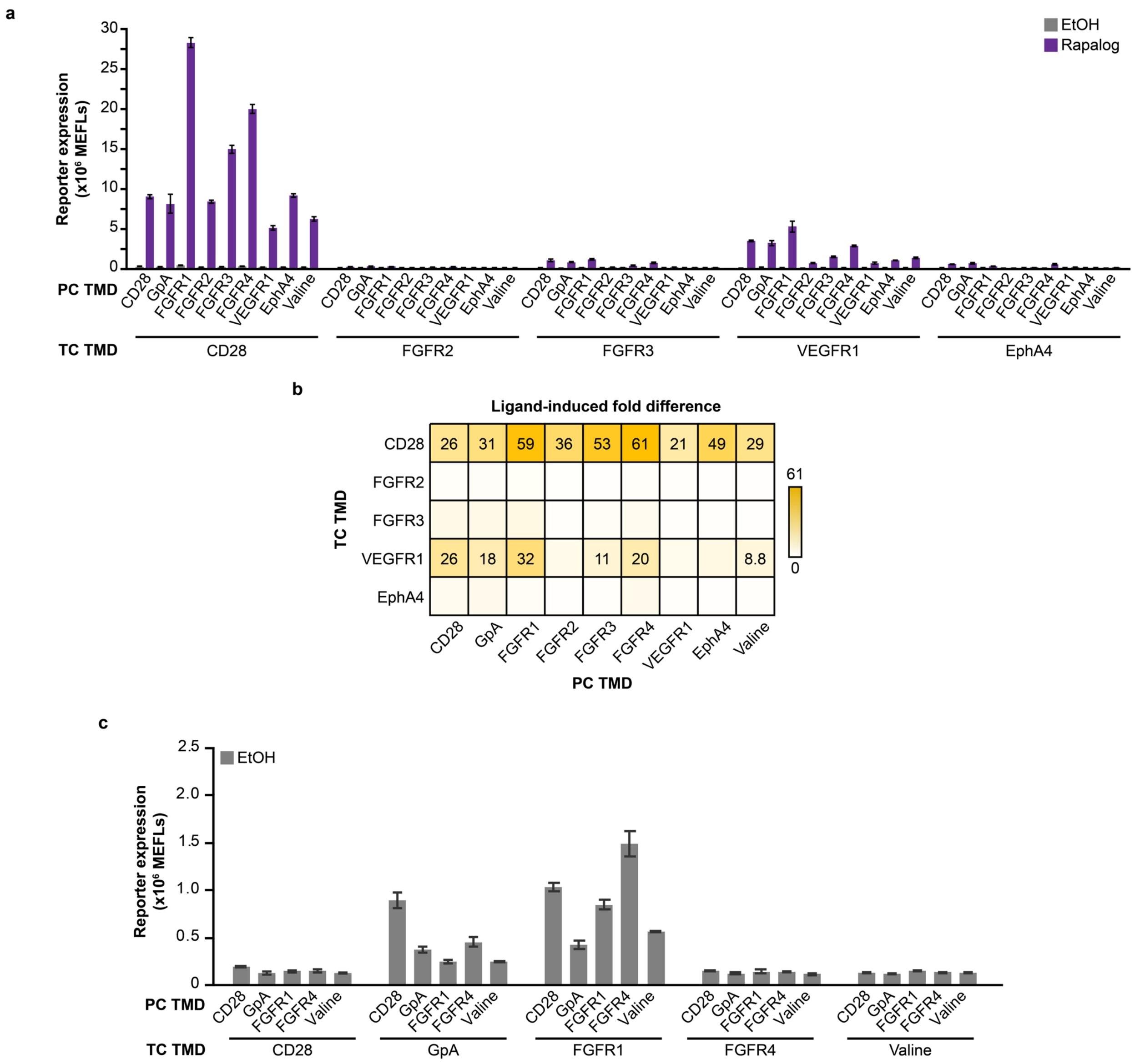


**Supplementary Fig. 8 Evaluating TCs that showed low ligand-induced signaling with TMD-matched PC.** (**a**) TCs that produced little or no ligand-induced signaling in **Fig. 2c** were tested with PCs containing matched or mixed TMDs. CD28-TMD TC was used as a positive control, as this chain cleaved well with the matched CD28-TMD PC. Most combinations produced little or no ligand-induced signaling, expect for those with the VEGFR1-TMD containing TC. However, even in these cases, ligand-induced signaling remained below those seen with the CD28-TMD. (**b**) Corresponding ligand-induced fold difference values are shown for conditions that yielded a significant difference between the vehicle and ligand-treated conditions. Statistical analysis is in **Supplementary Note 3**. (**c**) Reporter expression for the samples treated with vehicle (EtOH) shown in **Fig. 2d**. Data are reproduced here with a y-axis scaled to highlight differences between background reporter expression conferred by pairs of receptors with different mixed TMDs. Background signal varies with choice of TC TMD and PC TMD and the interaction between these variables, but TC choice explains most of this variance. Statistical analysis (two-way ANOVA) is in **Supplementary Note 2**. Throughout all panels, bars are the means of three biological replicates, and error bars depict S.E.M.

**a**

TMDOCK analysis

| TMD | Models generated | Highest ranked model $\Delta G_{asc}$ | Minimum $\Delta G_{asc}$ |
|---|---|---|---|
| CD28 | 6 | -7.4 | -7.4 |
| EphA4 | 7 | -7.9 | -7.9 |
| FGFR1 | 2 | -16.1 | -16.1 |
| FGFR2 | 7 | -2.5 | -2.5 |
| FGFR3 | 6 | -11.9 | -11.9 |
| FGFR4 | 3 | -3.4 | -3.4 |
| GpA | 3 | -11.9 | -11.9 |
| VEGFR1 | 7 | -7.8 | -7.8 |
| Valine | 5 | -1.4 | -1.5 |

Color scales: Models generated 7 to 2; Highest ranked model $\Delta G_{asc}$ -1.4 to -16.1; Minimum $\Delta G_{asc}$ -1.5 to -16.1

**b**

TMDOCK-identified transmembrane sequences

*(Blue residues indicate parts of intracellular domains)*

| | |
|---|---|
| CD28 | STGLVVVAGVLFCYGLLVTVALCVENE |
| EphA4 | VLLVSVSGSVVLVVILIAAFVIE |
| FGFR1 | TGIIIYCTGAFLISCMVGSVIV |
| FGFR2 | GIAIYCIGVFLIACMVVTVILE |
| FGFR3 | GILSYGVGFFLFILVVAAVTL |
| FGFR4 | TGIILYASGSLALAVLLLLAGL |
| GpA | TGITLIIFGVMAGVIGTILLIS |
| VEGFR1 | ITLTCTCVAATLFWLLLTLFIE |
| Valine | TGVVVVVVVVVVVVVVVVVVVVVVVVVE |

**c**

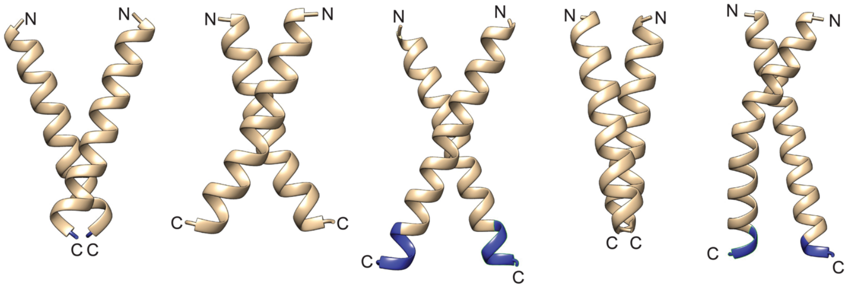


| TMD | FGFR1 | GpA | CD28 | FGFR4 | Valine |
|---|---|---|---|---|---|
| $\Delta G_{asc}$ (kcal/mol) | -16.1 | -11.9 | -7.4 | -3.4 | -1.4 |
| Predicted C-terminal distance (CTD) between dimerized TMDs (Å) | 4.3 | 22.0 | 16.4 | 4.6 | 13.0 |

**Supplementary Fig. 9 TMDOCK analysis** (**a**) TMDOCK (2) analysis of MESA receptors containing a panel of TMDs generated a set of stable dimerization models. The number of models is a proxy for the flexibility of association within each system, and the models are ranked based on free energies of association ($\Delta G_{asc}$) and other parameters. Values of $\Delta G_{asc}$ are shown for the highest ranked and most energetically stable predicted models. Larger absolute values for negative free energies of association indicate more stable association. The $\Delta G_{asc}$ predictions suggest that the panel of TMDs spans a range of association propensities. The number of models generated for each TMD suggests that the panel also spans a range of flexibility with respect to association modes. (**b**) TMD sequences identified by TMDOCK from full receptor protein sequences. Blue coloring denotes residues that are part of the ICD. (**c**) The highest ranked model prediction for each of the CD28, GpA, FGFR1, FGFR4, and Valine TMDs, listed in order from most to least stable $\Delta G_{asc}$. The range of predicted C-terminal distances across dimerized TMDs suggests that ICD geometries vary with TMD choice.

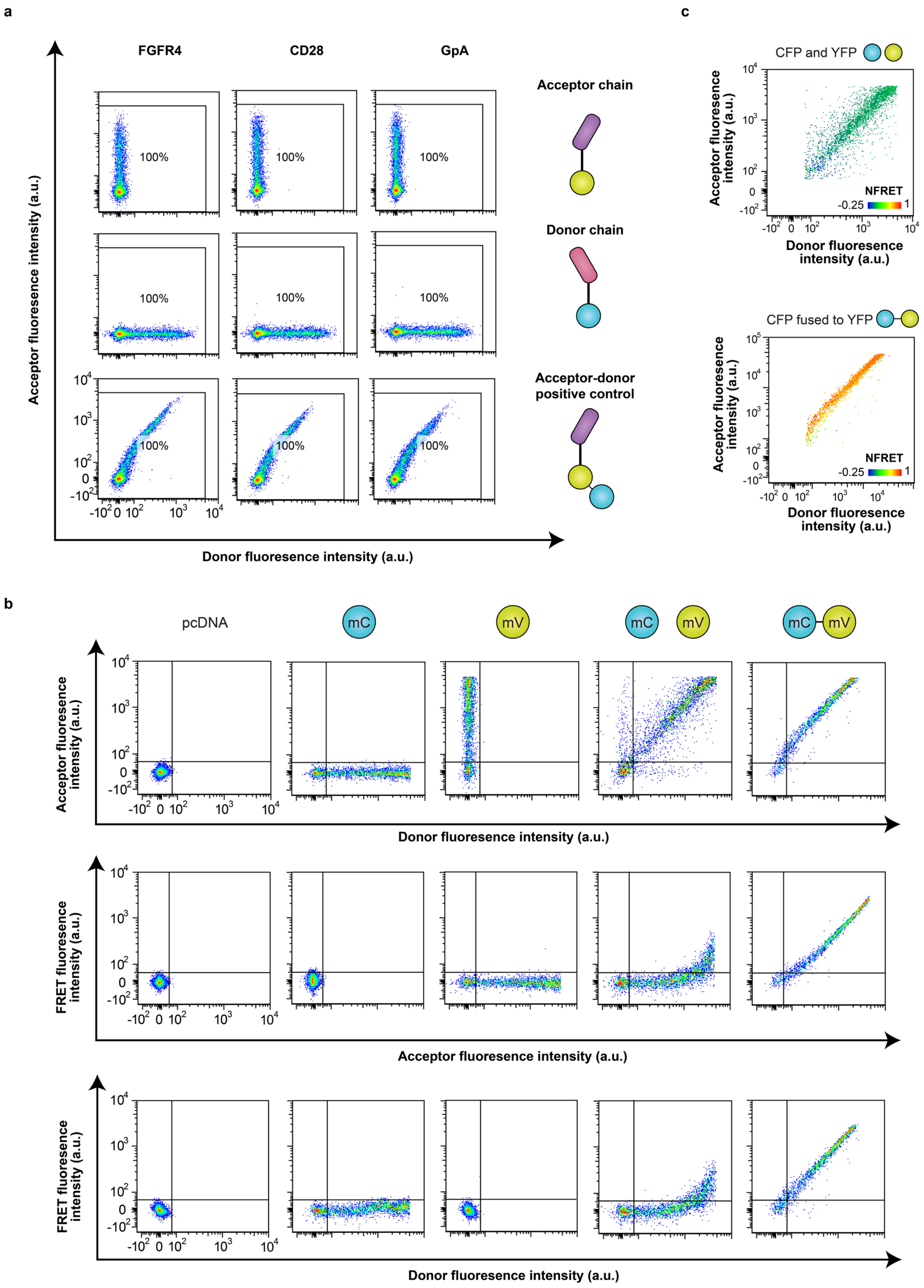
a
FGFR4
CD28
GpA
Acceptor chain
Donor chain
Acceptor-donor positive control
100%
Acceptor fluoresence intensity (a.u.)
Donor fluoresence intensity (a.u.)
c
CFP and YFP
CFP fused to YFP
NFRET
-0.25
1
Acceptor fluoresence intensity (a.u.)
Donor fluoresence intensity (a.u.)
b
pcDNA
mC
mV
Acceptor fluoresence intensity (a.u.)
Donor fluoresence intensity (a.u.)
FRET fluoresence intensity (a.u.)
Acceptor fluoresence intensity (a.u.)
FRET fluoresence intensity (a.u.)
Donor fluoresence intensity (a.u.)

**Supplementary Fig. 10 Gating for analytical flow cytometry to quantify FRET.** (**a**) Representative flow cytometry plots for samples that have a single donor chain or acceptor chain or a membrane-tethered positive FRET control containing CD28-TMD, GpA-TMD, or FGFR4-TMD. The populations shown include all single cells. The rectangular gate denotes the range of donor and acceptor expression over which spectral bleed-through correction was performed. This gate encompasses all single cells. (**b**) Representative flow cytometry plots for cytosolic control samples illustrate fluorescence intensities in the donor, acceptor, and FRET channels after compensation. The populations shown include all single transfected cells within the range designated in **a**. In **a** and **b**, dot color represents probability density. Cells expressing high levels of both donor and acceptor fluorophores exhibit some non-zero signal in the FRET channel, which appears to represent a true FRET signal arising under these conditions of high fluorophore concentration. (**c**) Representative flow cytometry plots for donor and acceptor expression of cytosolic negative and positive FRET controls. In **c**, dot color represents NFRET value, and homogeneity of dot color across a cell population indicates effective expression normalization.

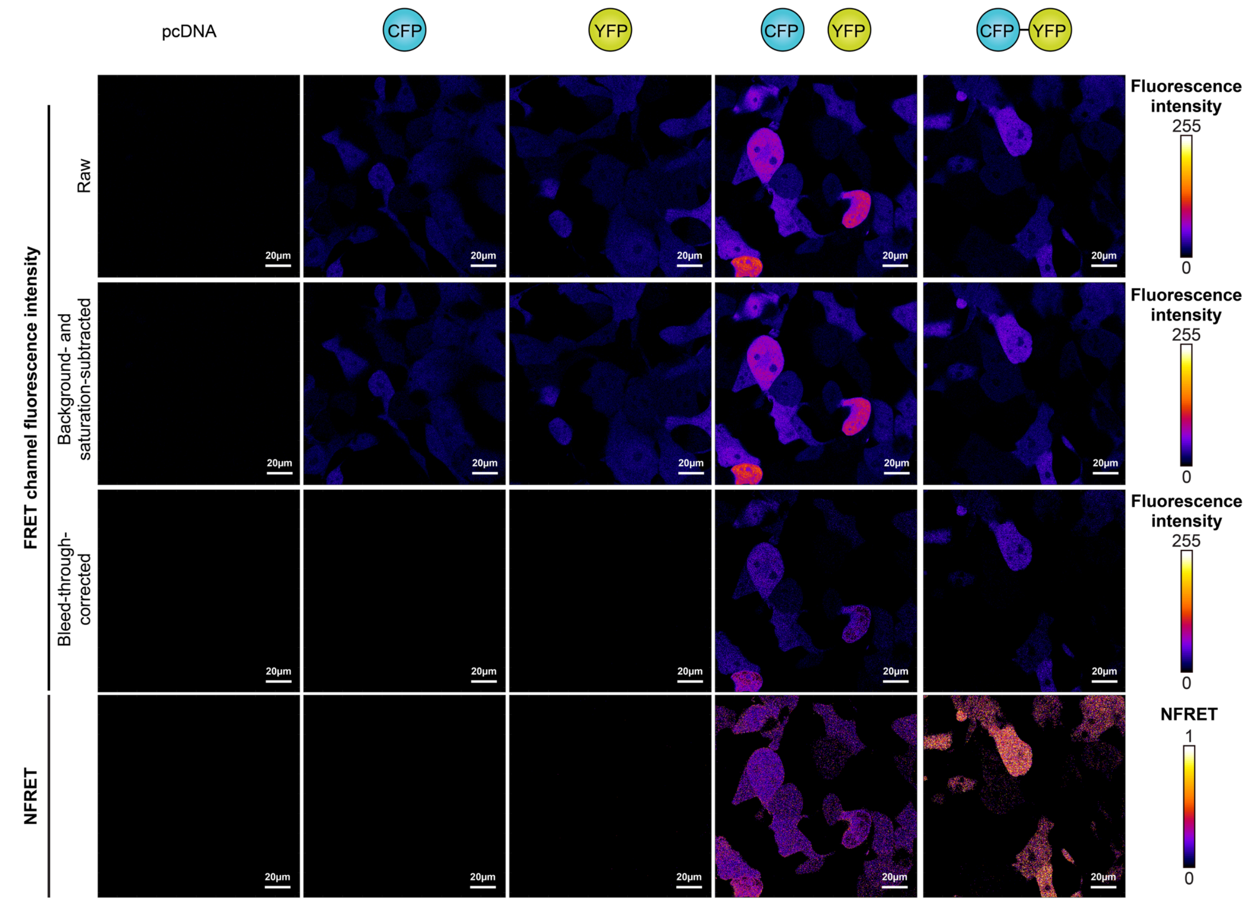


**Supplementary Fig. 11 Workflow for confocal microscopy image processing.** Representative confocal microscopy images of cytosolic control samples validate FRET observed by flow cytometry. The upper three rows show raw, background-subtracted, and bleed-through–corrected images of fluorescence intensity in the FRET channel. The bottom row shows NFRET across samples. The correction workflow is detailed in **Methods**. Cells co-expressing unlinked donor and acceptor fluorophores appear to exhibit low, non-zero FRET signal; we suspect that this is an artifact due to the approach that we employed to correct for spectral bleed-through across cells with a wide range of protein expression levels. Our bleed-through correction (subtraction of FRET signal contributed by the presence of either the donor or acceptor alone) used a maximally unbiased approach that employs mean background values calculated across a combined dataset (i.e., ten fields of view were analyzed together). While this method is conservative and necessary when analyzing a relatively small number of samples, fluorescence values vary widely between cells and the brightest cells may be under-corrected (i.e., some signal attributed to FRET could be uncorrected bleed-through of the acceptor and/or donor fluorophore into the FRET channel).

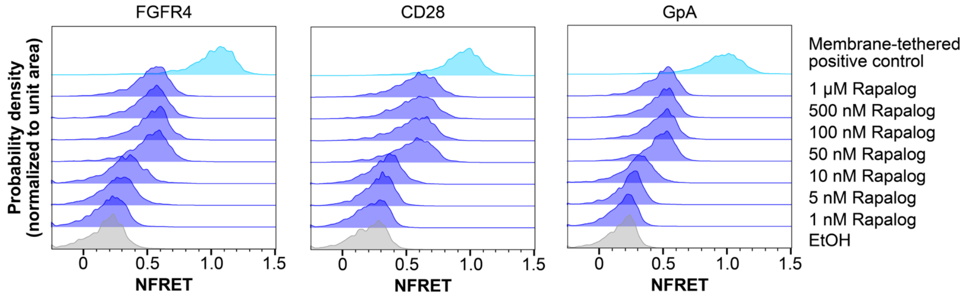


**Supplementary Fig. 12 Characterization of mixed and matched TMD receptor FRET by flow cytometry.** Flow cytometry histograms correspond to mean data in **Fig. 4b**. The populations shown are single, transfected, mCerulean+/mVenus+ cells. The top row of each plot shows the NFRET histogram for a membrane-tethered mCerulean-10aa-mVenus fusion protein containing the respective TMD.

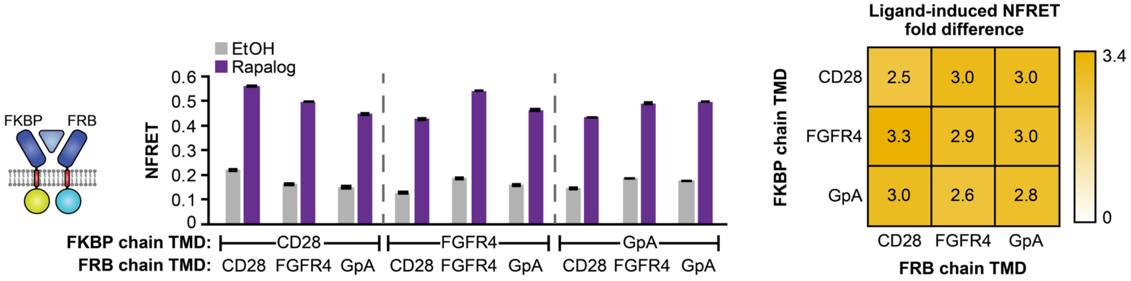


**Supplementary Fig. 13 Characterization of mixed and matched TMD receptor FRET by flow cytometry.** Complementary configuration of ECDs and fluorophores to **Fig. 4c**. Pairs of receptors in this orientation exhibit significant ligand-induced increase in NFRET after 27 h incubation with 100 nM rapalog or vehicle (three-way ANOVA, $p < 0.001$). This panel produces ligand-independent and ligand-induced NFRET (left) and ligand-induced NFRET fold differences (right) that are comparable to those observed for fluorophores attached in the opposite orientation (**Fig. 4c**). Bars are the means of three biological replicates, and error bars depict S.E.M. Outcomes from ANOVAs and Tukey's HSD tests are in **Supplementary Note 3**.

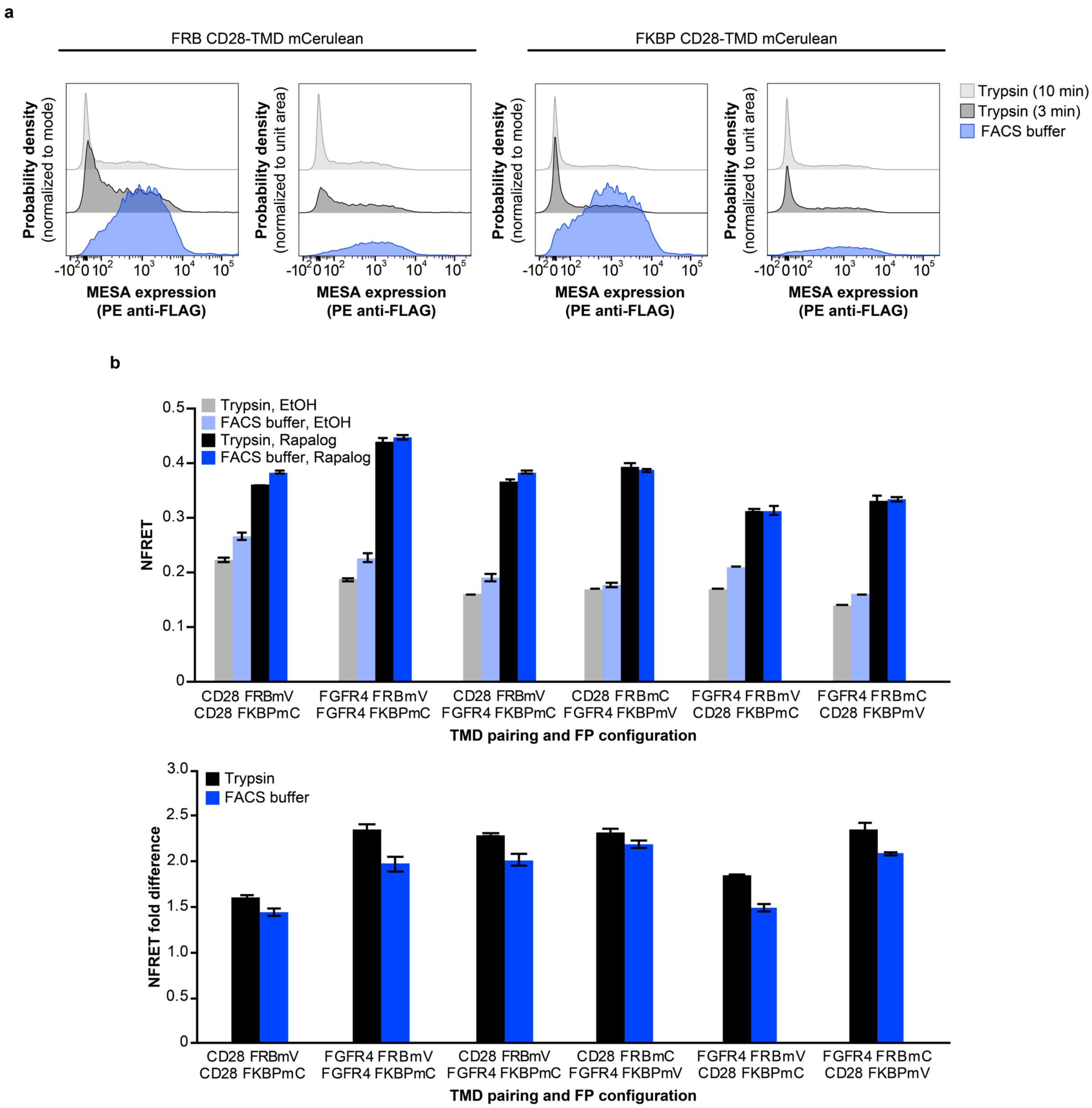


**Supplementary Fig. 14 Characterization of the effect of different harvest methods on receptor surface expression and NFRET.** (**a**) Comparison of surface expression measurement for CD28-TMD rapamycin-sensing MESA receptors after short (3 min) or long (10 min) trypsinizations vs. harvest with FACS Buffer (1× PBS, 0.1% BSA, 5mM EDTA). The short trypsinization, which is used in all flow cytometric FRET experiments, reduces some but not all surface expression. (**b**) Comparison of NFRET (upper) and comparison of ligand-induced NFRET fold difference (lower) for samples harvested with short trypsinization vs. FACs Buffer. A consistent decrease in NFRET fold difference is observed with FACS Buffer compared to with trypsin, due to proportionally larger NFRET values upon vehicle treatment with FACS Buffer than with trypsin. This decrease does not affect the conclusions made about significant differences between samples within each harvest method (two-tailed Welch's *t*-tests produced consistent conclusions for significant differences across TMD pairs, **Supplementary Note 4**). Bars are the means of three biological replicates, and error bars depict S.E.M.

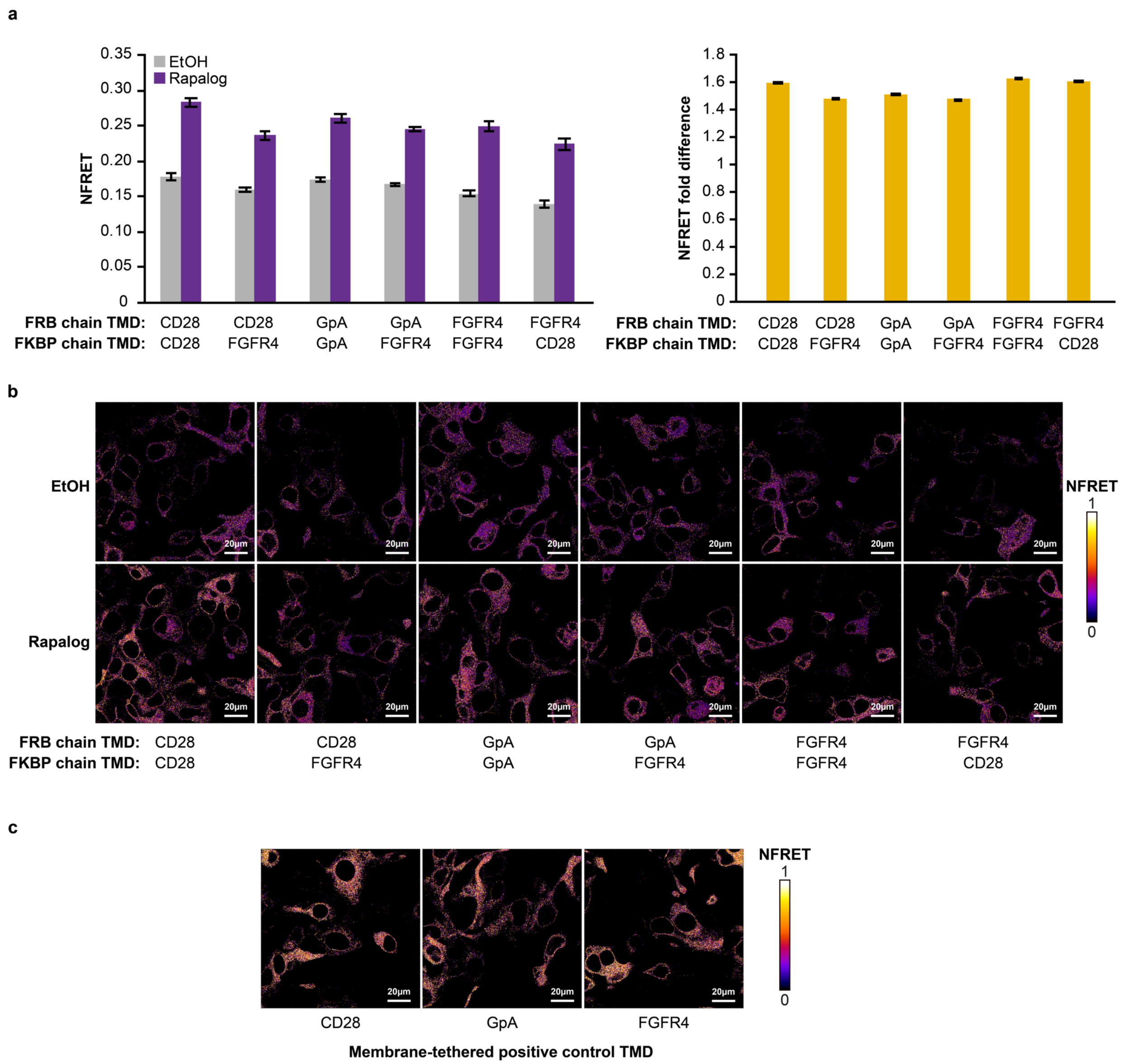


**Supplementary Fig. 15 Characterization of mixed and matched TMD receptor FRET by confocal microscopy.** (**a**) FRET assay of selected receptor TMD pairs using confocal microscopy. Left: NFRET was averaged across pixels in ten fields of view (as described in **Methods**) and plotted. Right: ligand-induced NFRET fold difference was calculated by dividing NFRET with ligand treatment by NFRET with vehicle-only treatment. Fold differences across pairs are not significantly different (two-tailed Welch's *t*-test, all $p > 0.05$). (**b**) Representative, processed confocal images of NFRET for receptor pairs treated with EtOH vehicle only (upper) or with rapalog (lower). (**c**) Representative confocal images of NFRET for membrane-tethered mCerulean-10aa-mVenus fusion proteins containing different TMDs (positive controls).

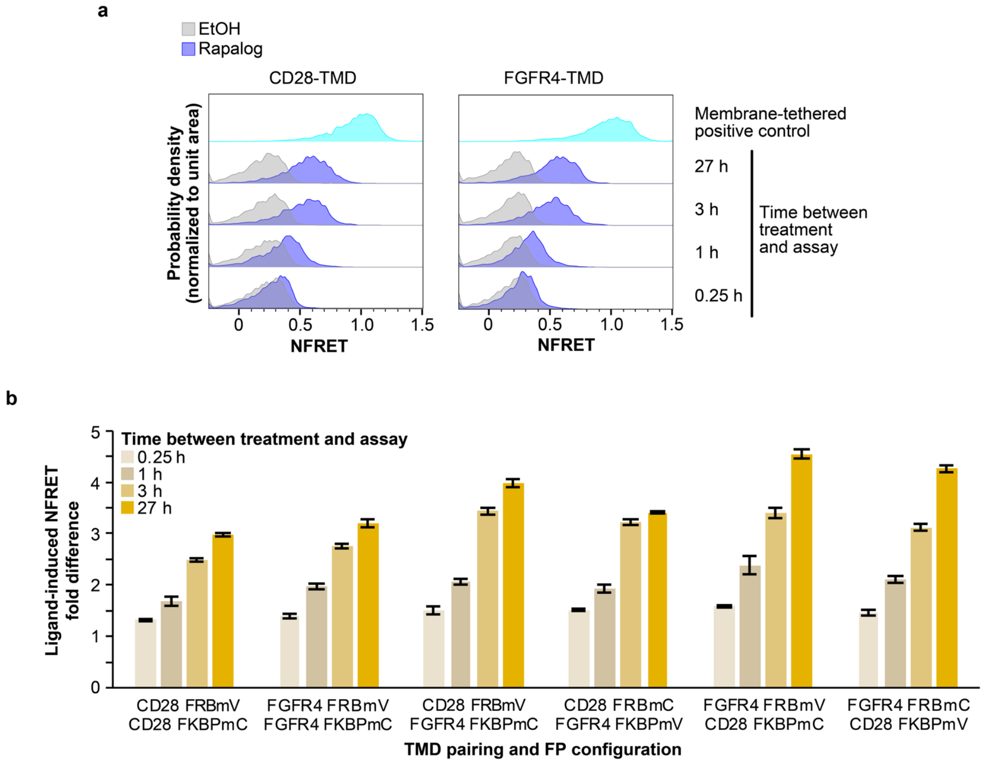


**Supplementary Fig. 16 Quantification of receptor FRET over time.** (**a**) Flow cytometry histograms corresponding to mean data in **Fig. 4d** show increasing NFRET in rapalog-treated samples over time. The populations shown are single transfected mCerulean+/mVenus+ cells. The top row of each plot shows the NFRET histogram for a membrane-tethered mCerulean-10aa-mVenus fusion protein containing the respective TMD. (**b**) Ligand-induced NFRET fold difference is comparable for each combination of CD28-TMD and FGFR4-TMD receptors for different treatment durations. Bars are the means of three biological replicates, and error bars depict S.E.M.

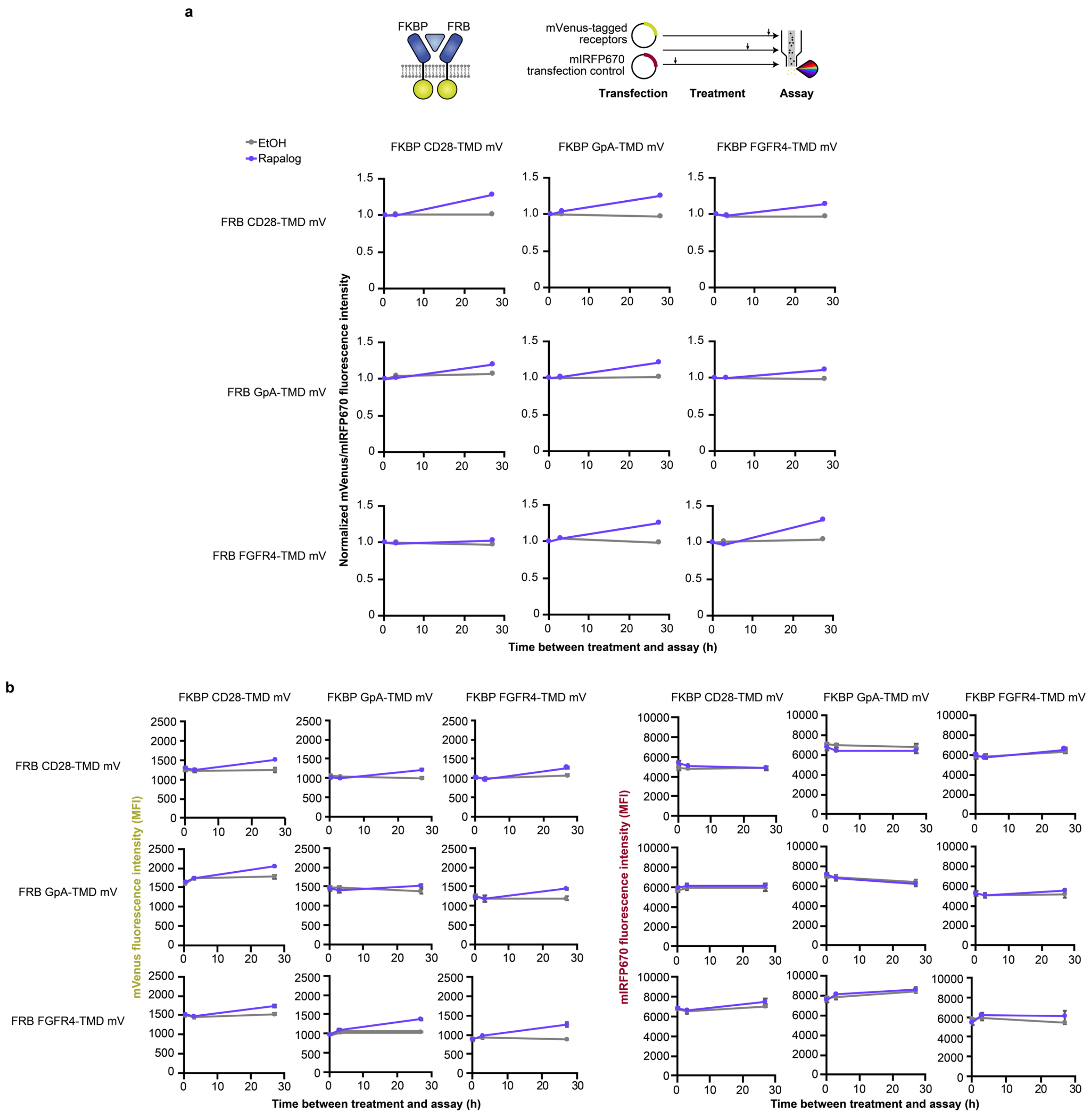


**Supplementary Fig. 17 Quantification of ligand-mediated effects on receptor expression.** Cells were transfected with mVenus-tagged receptor pairs (that have mixed and matched TMDs) and with an miRFP670 transfection control plasmid. Cells were treated with rapalog or with EtOH vehicle only for 15 min, 3 h, or 27 h prior to the assay. (**a**) To account for differences in transfection efficiency across samples, mVenus MFI was divided by miRFP670 MFI, and this ratio was normalized to the mVenus/miRFP670 ratio measured after 15 min of treatment with vehicle only or ligand. Across TMD pairs, an increase in mVenus fluorescence intensity was observed—up to ~5% between 0.25–3 h and up to ~30% between 3–27 h of ligand treatment. (**b**) Non-normalized mVenus MFI (left) and miRFP670 MFI (right) indicate that the increase is largely due to increased mVenus MFI; miRFP670 MFI is less affected. Although ligand treatment modestly increased the accumulation of individual chains over a long interval (e.g., 27 h), this modest increase would not be expected to substantially impact NFRET, particularly because this metric is already scaled to account for difference in protein expression level (**Fig. 3d**). Plotted points are the means of three biological replicates, and error bars depict S.E.M.

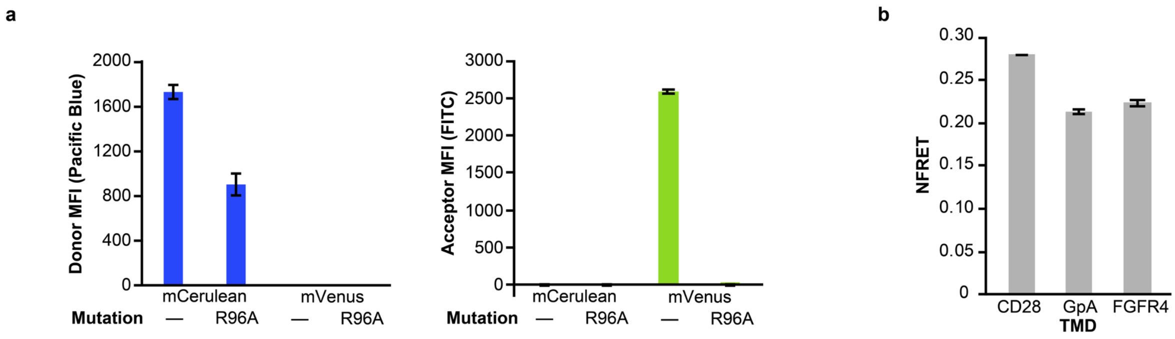


**Supplementary Fig. 18 Characterization of FP mutant variants.** (**a**) To construct dead chains that do not fluoresce and are not capable of participating in FRET, we mutated both mCerulean and mVenus to disrupt chromophore maturation (3). Plots show MFI after compensation of mCerulean, mCerulean R96A, mVenus, and mVenus R96A in the donor channel (Pacific Blue) and acceptor channel (FITC). While the R96A mutation caused a decrease in mCerulean fluorescence in the donor channel, the mutation actually ablated detectable fluorescence of mVenus in the acceptor channel. Based on this finding, the R96A mVenus mutant was used in cold chain competition assays. (**b**) NFRET for matched TMD pairs of FP-tagged chains containing FRB ECDs; these data correspond to the fold difference data in the gray bars in **Fig. 4f.** Bars are the means of three biological replicates, and error bars depict S.E.M.

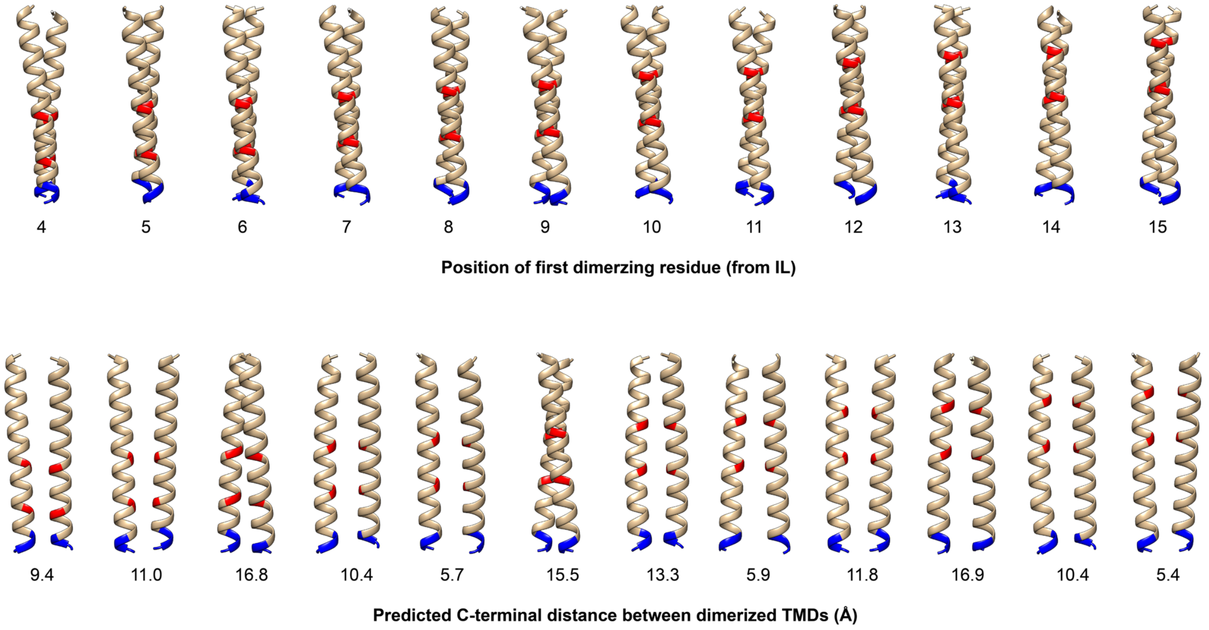


**Supplementary Fig. 19 Syn Rot TMD C-terminal distances**. TMDOCK (2) analysis for MESA receptors containing synthetic TMDs that have dimerizing residues positioned at different locations (positions are with respect to the inner linker (IL), described in **Fig. 5**). The models shown are those predicted to be the most energetically stable (i.e., with the lowest free energy of association, $\Delta G_{asc}$) for each TMD. Blue coloring indicates intracellular residues, and red coloring indicates interacting glutamic acid residues. In the top row, model representations are aligned so that interacting residues are overlapping in the plane of view, and in the bottom row they are aligned to show the predicted C-terminal distances (CTDs), which are listed below each pair. These predictions do not account for potential geometric constraints imposed by other domains of MESA receptors.

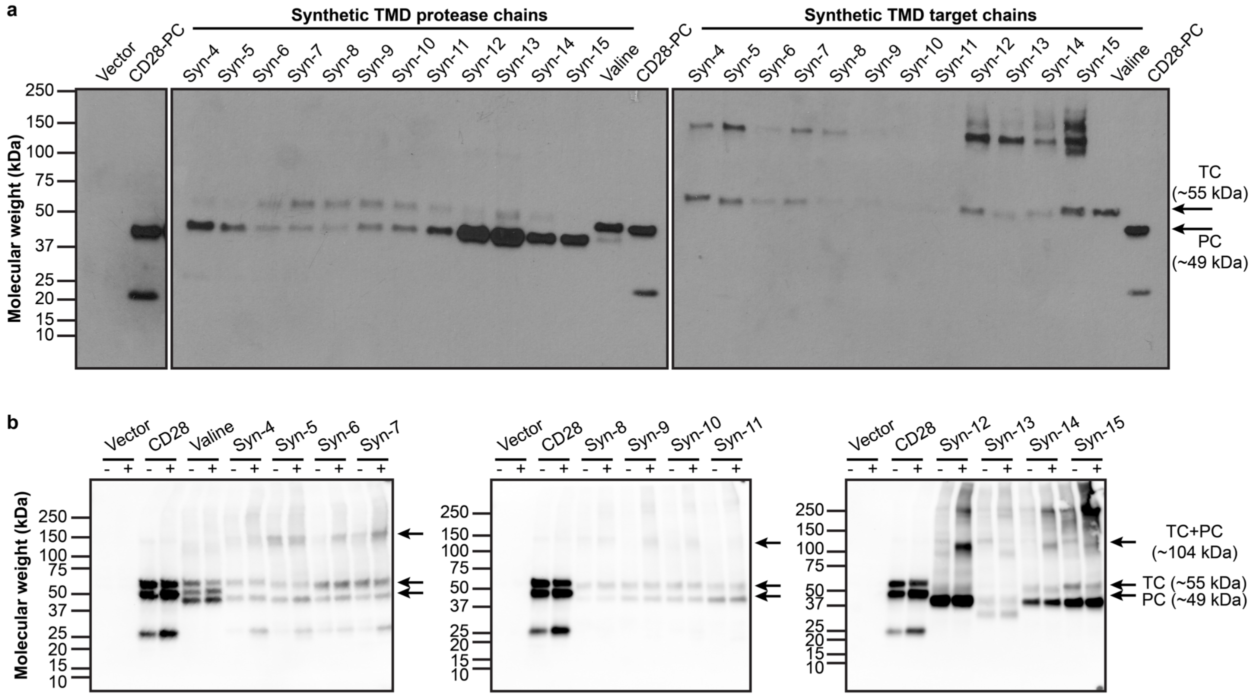


**Supplementary Fig. 20. Investigating ligand-induced accumulation of MESA receptors containing synthetic TMDs.** (**a**) Western blots of individual MESA chains containing synthetic TMDs show a range of expression, with higher expression for cases in which dimerizing motifs are located near the outer leaflet of the membrane. (**b**) Some co-transfected MESA chains containing matched, synthetic TMDs and a catalytically inactive TEVp on the PC (D81N mutation (4)) show modestly higher expression with rapalog treatment (+) than with vehicle-only treatment (-). The expression level trends observed across the vehicle-only conditions are similar to the expression level trends observed across the rapalog conditions. Therefore, we conclude that chain expression alone does not explain the trends in **Fig. 5b**.

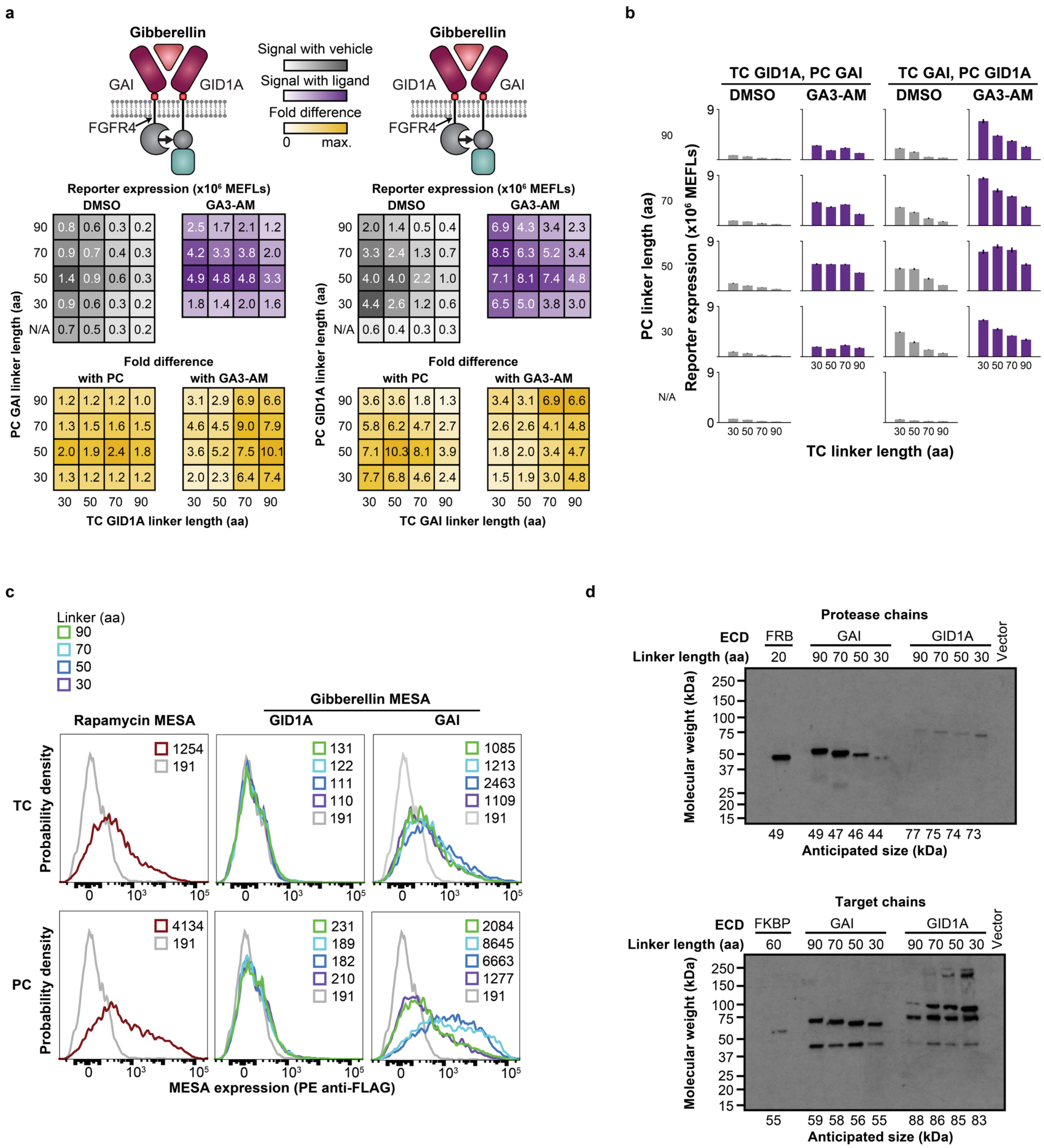
a
Gibberellin
GAI
GID1A
FGFR4
Signal with vehicle
Signal with ligand
Fold difference
0
max.
Reporter expression (x10⁶ MEFLs)
DMSO
GA3-AM
PC GAI linker length (aa)
PC GID1A linker length (aa)
Fold difference
with PC
with GA3-AM
TC GID1A linker length (aa)
TC GAI linker length (aa)
b
TC GID1A, PC GAI
TC GAI, PC GID1A
DMSO
GA3-AM
PC linker length (aa)
Reporter expression (x10⁶ MEFLs)
TC linker length (aa)
N/A
c
Linker (aa)
90
70
50
30
Rapamycin MESA
Gibberellin MESA
GID1A
GAI
TC
PC
Probability density
MESA expression (PE anti-FLAG)
d
Protease chains
ECD
FRB
GAI
GID1A
Linker length (aa)
Vector
Molecular weight (kDa)
Anticipated size (kDa)
Target chains
FKBP

**Supplementary Fig. 21 Selection of an extracellular linker for the gibberellin-sensing receptor system.** Gibberellin-sensing MESA receptor variants with different ligand binding domain (LBD)-intracellular domain (ICD) configurations (fusion of GID1 and GAI to the PC or TC) and extracellular linker lengths (30, 50, 70, 90 amino acids) were built and characterized. These receptors all contain FGFR4-TMD. (**a,b**) Functional assay for receptor configurations. Receptor plasmids were transfected at a consistent plasmid ratio (2:1 TC plasmid:PC plasmid) along with a reporter plasmid and an EBFP2 transfection control plasmid, holding total plasmid dose constant at 700 ng DNA per well. Heatmaps and bar graphs depict the mean of three biological replicates of transfected (EBFP2+) cells treated with ligand (GA3-AM) or vehicle only (DMSO). Fold difference was calculated by dividing reporter signal with ligand treatment by reporter signal with vehicle-only treatment. Error bars depict S.E.M. (**c**) Surface staining with anti-FLAG-PE for each single chain configuration to assay for surface expression. Numbers indicate mean PE signal for each sample. A sample of EBFP2-transfected cells (without any FLAG tag-containing component) is the negative control, and rapamycin-sensing MESA receptor containing FGFR4-TMD is a positive control. (**d**) Western blots for each single chain configuration to assay whole-cell expression.

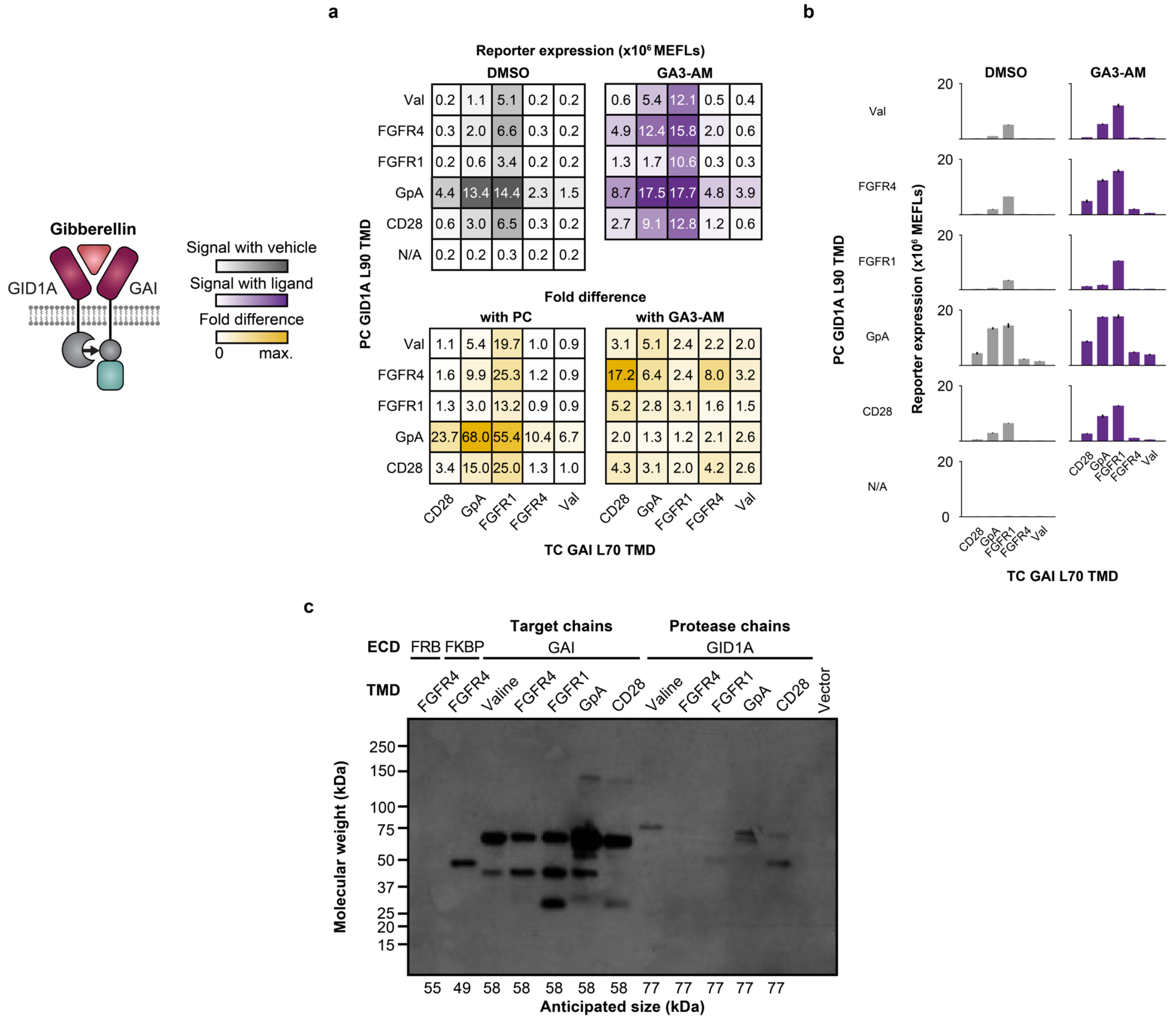


**Supplementary Fig. 22 Characterization of TMDs in the gibberellin-sensing receptor system.** Gibberellin-sensing MESA receptor variants with different TMDs (CD28, GpA, FGFR1, FGFR4, and Valine) were built and characterized. (**a**,**b**) Functional assay for receptor configurations. Receptor plasmids were transfected at a consistent plasmid ratio (2:1 TC plasmid:PC plasmid) along with a reporter plasmid and an EBFP2 transfection control plasmid, holding total plasmid dose constant at 700 ng DNA per well. Some of the data shown here are also shown in **Fig. 6b**. Heatmaps and bar graphs depict the mean of three biological replicates of transfected (EBFP2+) cells treated with ligand (GA3-AM) or vehicle only (DMSO). Fold difference was calculated by dividing reporter signal with ligand treatment by reporter signal with vehicle-only treatment. Error bars depict S.E.M. Outcomes from ANOVAs and Tukey's HSD tests are in **Supplementary Notes 2 and 3**. (**c**) Western blot for each single chain to assay whole-cell expression.

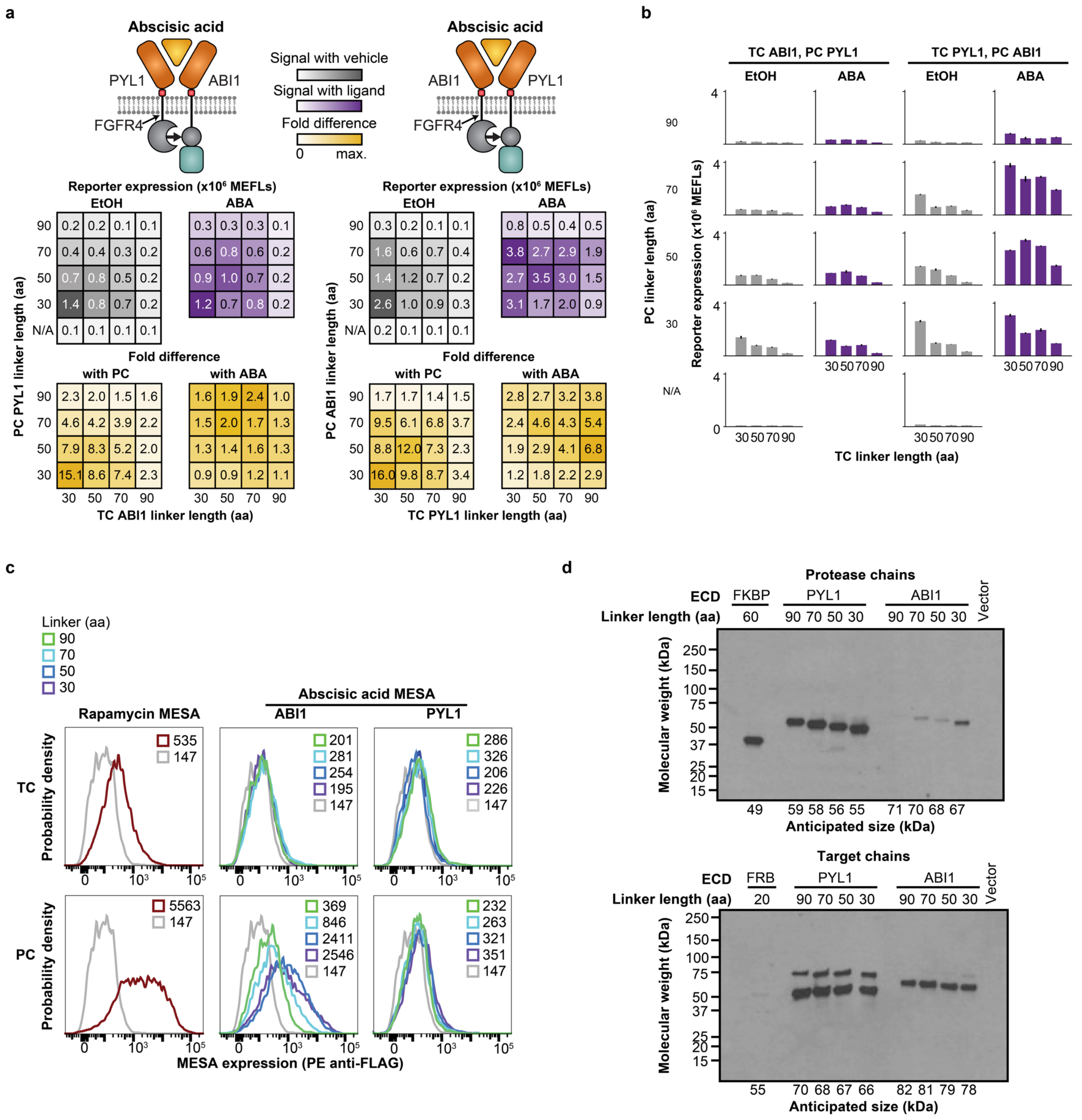


**Supplementary Fig. 23 Selection of an extracellular linker for the abscisic acid-sensing receptor system.** Abscisic acid-sensing MESA receptor variants containing different LBD-ICD configurations (fusion of PYL1 and ABI1 to the PC or TC) and extracellular linker lengths (30, 50, 70, 90 amino acids) were built and characterized. These receptors all contain FGFR4-TMD. (**a**,**b**) Functional assay for receptor configurations. Receptor plasmids were transfected at equal plasmid dose along with a reporter plasmid and an EBFP2 transfection control plasmid. Heatmaps and bar graphs depict the mean of three biological replicates of transfected (EBFP2+) cells treated with ligand (ABA) or vehicle only (EtOH). Fold difference was calculated by dividing the reporter signal with ligand treatment by reporter signal with vehicle-only treatment. Error bars depict S.E.M. (**c**) Surface staining with anti-FLAG-PE for each single chain configuration to assay for surface expression. Numbers indicate mean PE signal for each sample. EBFP2 (without any FLAG tag-containing component) is the negative control, and rapamycin-sensing MESA receptor containing FGFR4-TMD is a positive control. (**d**) Western blots for each single chain configuration to assay for whole-cell expression.

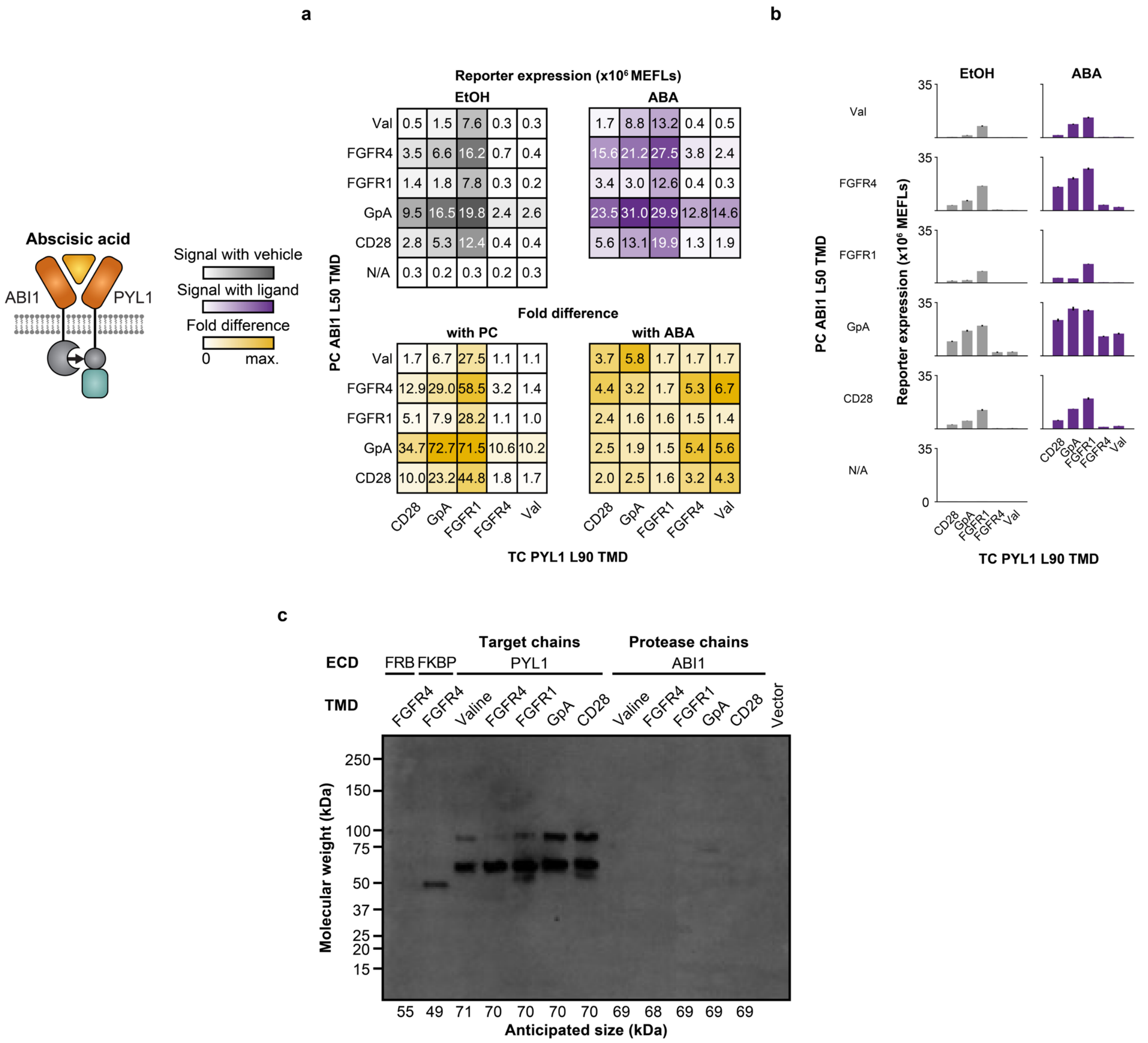


**Supplementary Fig. 24 Characterization of TMDs in the ABA-sensing receptor system.** Abscisic acid-sensing MESA receptor variants containing different TMDs (CD28, GpA, FGFR1, FGFR4, and Valine) were built and characterized. (**a,b**) Functional assay for receptor configurations. Receptor plasmids were transfected at equal plasmid dose along with a reporter plasmid and an EBFP2 transfection control plasmid. Some of the data shown here are also shown in **Fig. 6c**. Heatmaps and bar graphs depict the mean of three biological replicates of transfected (EBFP2+) cells treated with ligand (ABA) or vehicle only (EtOH). Fold difference was calculated by dividing reporter signal with ligand treatment by reporter signal with vehicle-only treatment. Error bars depict S.E.M. Outcomes from ANOVAs and Tukey's HSD tests are in **Supplementary Notes 2 and 3**. (**c**) Western blot for each single chain configuration to assay for whole-cell expression.

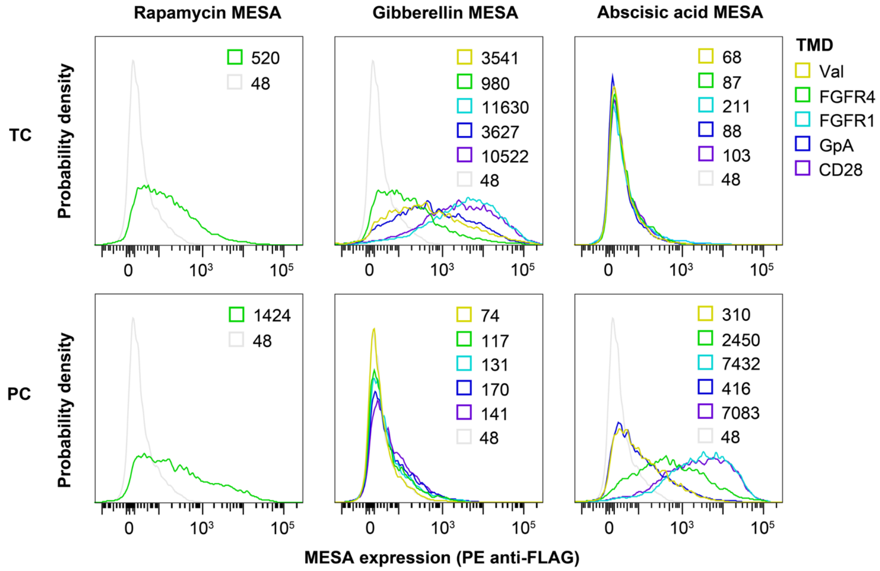


**Supplementary Fig. 25 Characterization of surface expression of abscisic acid-sensing and gibberellin-sensing receptor systems.** Surface staining with anti-FLAG-PE for each single chain containing each TMD for abscisic acid-sensing and gibberellin-sensing systems. Histograms are normalized to unit area. The numbers indicate mean PE signal for each sample. EBFP2 (without any FLAG tag-containing component) is the negative control, and a rapamycin-sensing MESA receptor containing FGFR4-TMD is a positive control.

a

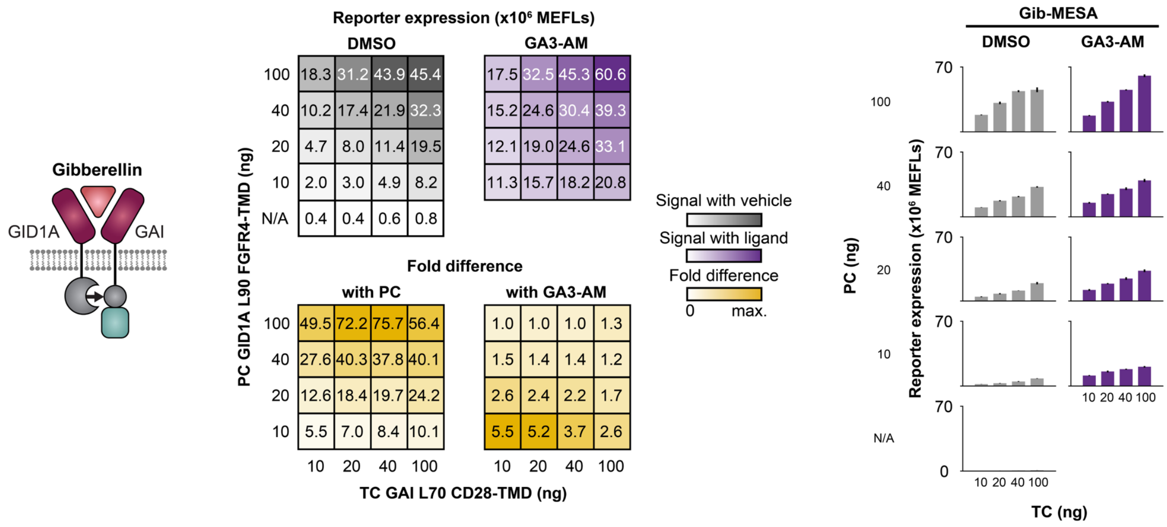


b

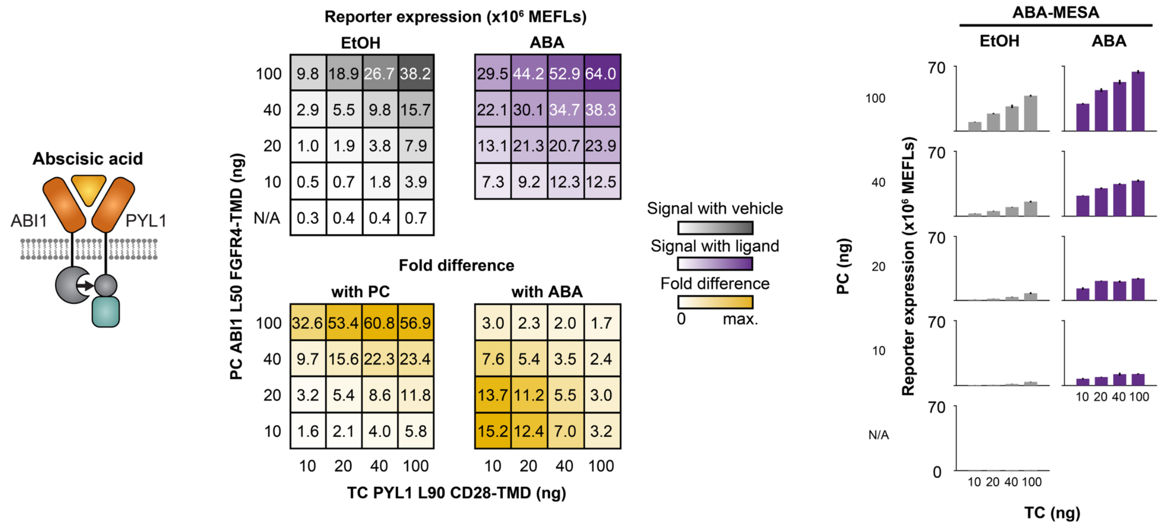


**Supplementary Fig. 26 Characterization of performance across plasmid doses for gibberellin-sensing and abscisic acid-sensing MESA systems.** Functional assays for gibberellin-sensing MESA receptor variants (TC GAI L70 CD28-TMD, PC GID1A L90 FGFR4-TMD) and abscisic acid-sensing MESA receptor variants (TC PYL1 L90 CD28-TMD, PC ABI1 L50 FGFR4-TMD) with varying plasmid dose combinations. Background and ligand-induced reporter expression generally increase with increasing TC and PC doses, and fold difference generally increases with decreasing TC and PC doses. Bars depict the mean reporter expression of three biological replicates of transfected (EBFP2+) cells treated with ligand (GA3-AM or ABA) or vehicle only (DMSO or EtOH). Error bars depict the S.E.M. Heatmaps depict reporter expression (upper) and PC-induced and ligand-induced fold difference (lower).

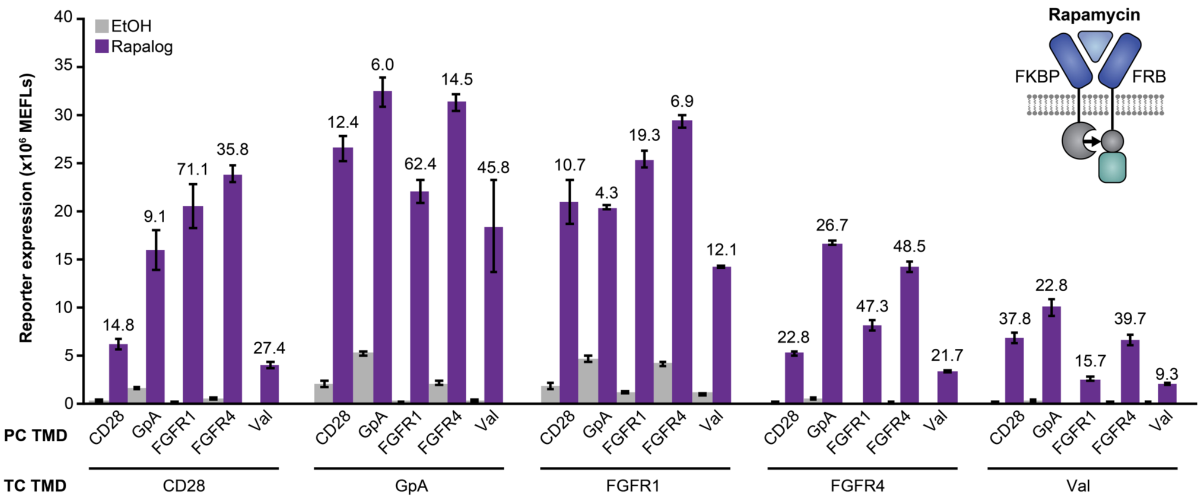


**Supplementary Fig. 27 Characterization of TMDs in the rapamycin-sensing receptor system.** Functional assay for rapamycin-sensing MESA receptor variants containing CD28, GpA, FGFR1, FGFR4, or Valine TMDs. Receptor plasmids were transfected at equal dose (not the protein expression-normalized doses used in **Fig. 2**) along with a reporter plasmid and an EBFP2 transfection control plasmid. Bars depict the mean reporter expression of three biological replicates of transfected (EBFP2+) cells treated with ligand (rapalog) or vehicle only (EtOH). The numbers above the purple bars indicate ligand-induced fold difference. The same data are in **Fig. 6a**. Error bars depict the S.E.M. Outcomes from ANOVAs and Tukey's HSD tests are in **Supplementary Notes 2 and 3**.

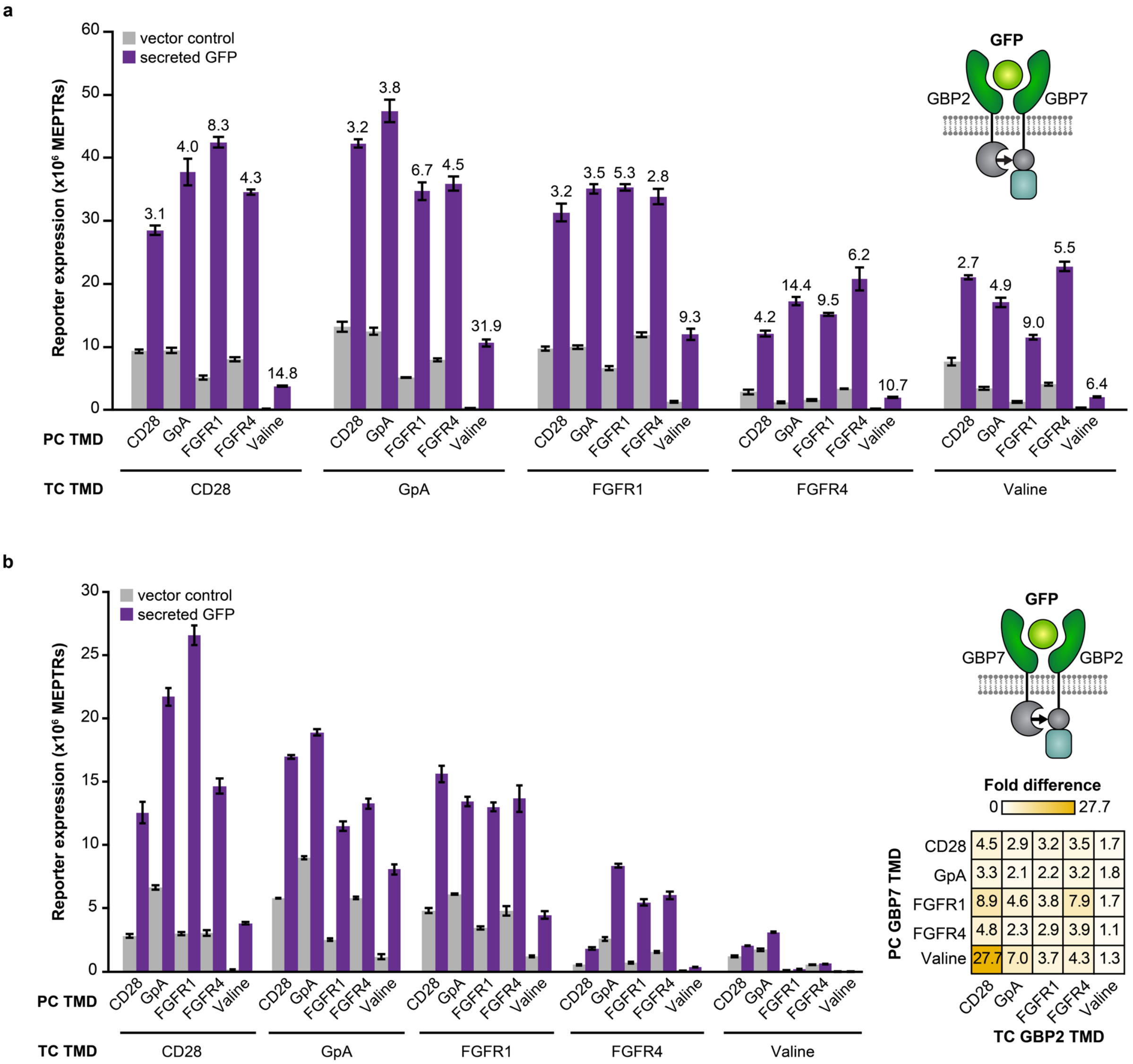


**Supplementary Fig. 28  Characterization of TMDs in the GFP-sensing receptor system.** Functional assay for sGFP-sensing MESA receptor variants containing CD28, GpA, FGFR1, FGFR4, or Valine TMDs. Receptor plasmids were transfected at equal plasmid dose along with a reporter plasmid and an EBFP2 transfection control plasmid. Bars depict the mean reporter expression of three biological replicates of transfected (EBFP2+) cells with ligand-encoding plasmid (sGFP) or vector control (pcDNA). (**a**) Bars depict the same data in **Fig. 6d**. The numbers above the purple bars indicate ligand-induced fold difference. (**b**) Bars depict functional assay data for the alternative LBD-ICD configuration in which GBP2 is on the TC and GBP7 is on the PC. Error bars depict the S.E.M. Outcomes from ANOVAs and Tukey's HSD tests are in **Supplementary Notes 2 and 3**.

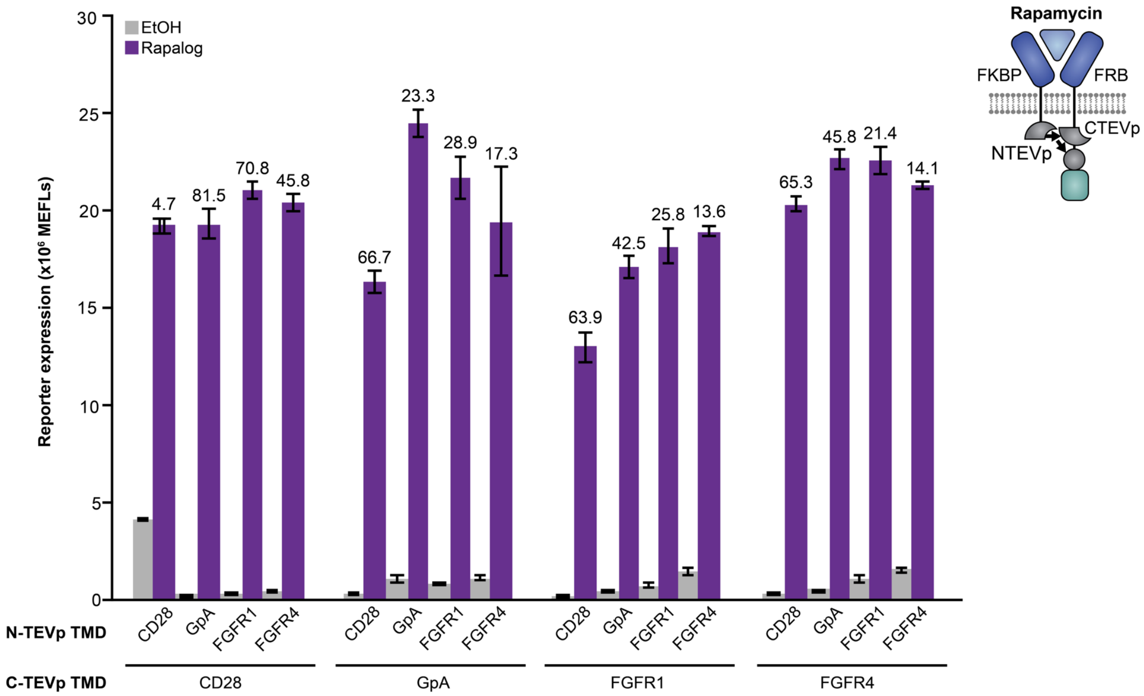


**Supplementary Fig. 29 Characterization of TMDs in the split TEVp rapamycin-sensing receptor system.** Functional assay for rapamycin-sensing split TEVp MESA receptor variants containing CD28, GpA, FGFR1, or FGFR4 TMDs. Receptor plasmids were transfected at equal plasmid dose along with a reporter plasmid and an EBFP2 transfection control plasmid. Bars depict the mean reporter expression of three biological replicates of transfected (EBFP2+) cells treated with ligand (rapalog) or vehicle only (EtOH). The numbers above the purple bars indicate ligand-induced fold difference. The same data are in **Fig. 6e**. Error bars depict the S.E.M. Outcomes from ANOVAs and Tukey's HSD tests are in **Supplementary Notes 2 and 3**.

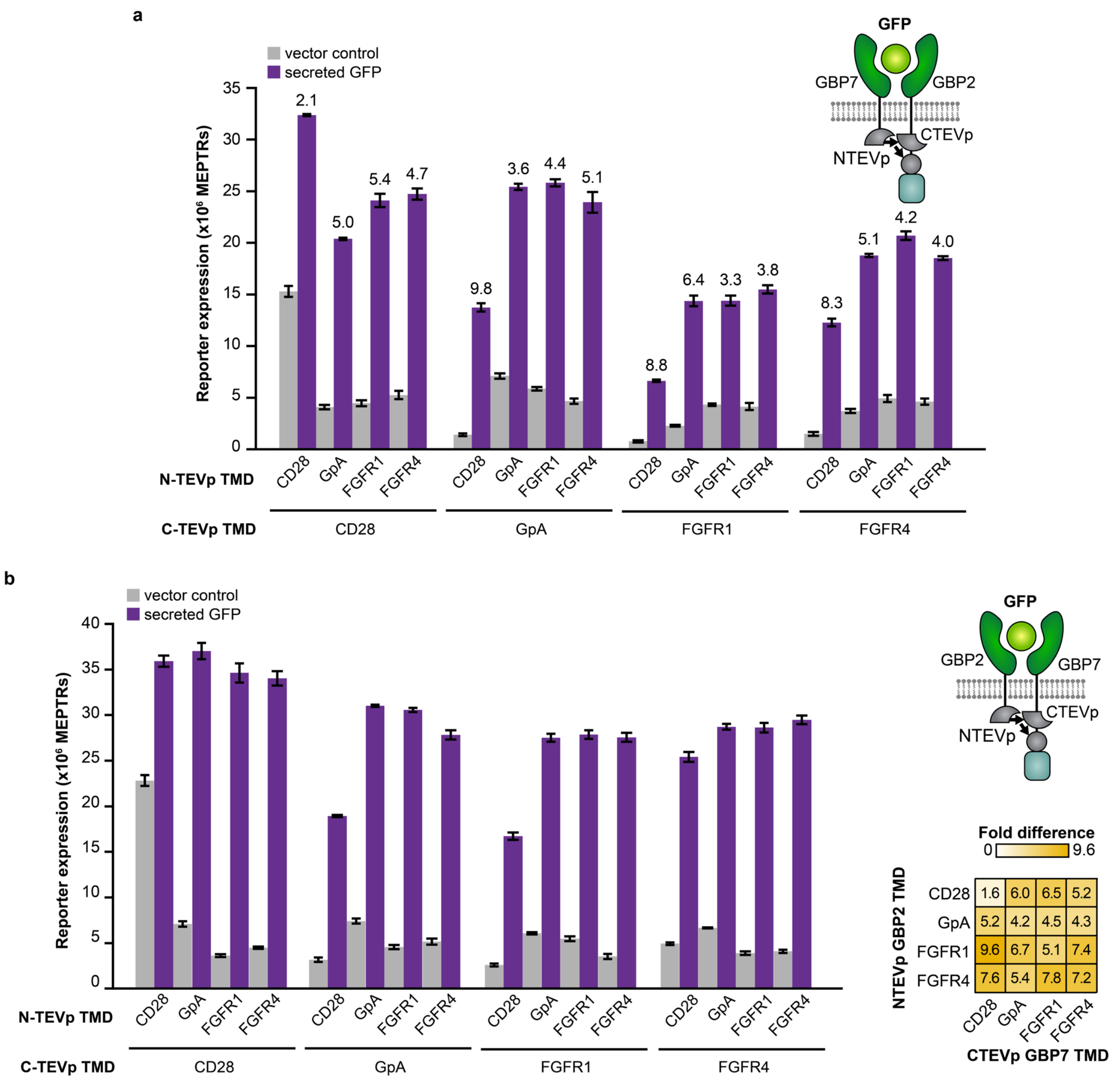


**Supplementary Fig. 30 Characterization of TMDs in the split TEVp sGFP-sensing receptor system.** Functional assay for GFP-sensing split TEVp MESA receptor variants containing CD28, GpA, FGFR1, or FGFR4 TMDs. Receptor plasmids were transfected at equal plasmid dose along with a reporter plasmid and an EBFP2 transfection control plasmid. Bars depict the mean reporter expression of three biological replicates of transfected (EBFP2+) cells with ligand-encoding plasmid (sGFP) or vector control (pcDNA). (**a**) Bars depict the same data in **Fig. 6f**. The numbers above the purple bars indicate ligand-induced fold difference. (**b**) Bars depict functional assay data for the alternative LBD-ICD configuration in which GBP2 is on the NTEVp chain and GBP7 is on the CTEVp chain. Error bars depict the S.E.M. Outcomes from ANOVAs and Tukey's HSD tests are in **Supplementary Notes 2 and 3**.

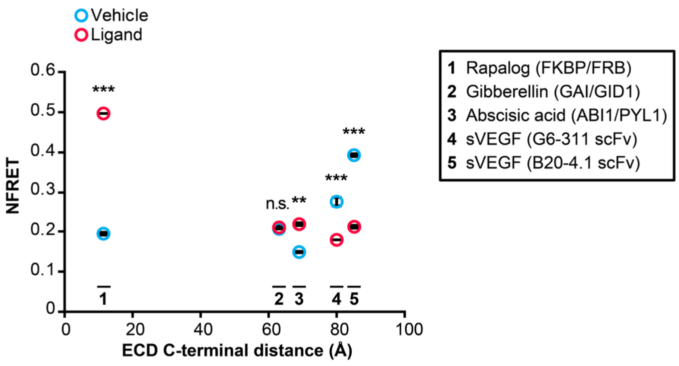


**Supplementary Fig. 31 Quantification of NFRET across ligand-sensing systems.** NFRET values for rapamycin, abscisic acid, gibberellin, and VEGF-sensing receptor systems underlying the fold difference values in **Fig. 7a**. The rapamycin-sensing and abscisic acid-sensing MESA FRET receptors exhibit a significant increase in NFRET upon ligand treatment (two-tailed Welch's *t*-test, $^{**}p < 0.01$, $^{***}p < 0.001$). VEGF-sensing MESA FRET receptors exhibit a significant decrease in NFRET when co-expressed with secreted VEGF (two-tailed Welch's *t*-test, $^{***}p < 0.001$). The gibberellin-sensing MESA FRET receptors do not exhibit a significant change in NFRET upon ligand treatment (two-tailed Welch's *t*-test, n.s. $p > 0.05$). Plotted points are the means of three biological replicates, and error bars depict S.E.M.

### SUPPLEMENTARY TABLES

#### Supplementary Table 1. Ligand concentrations for small molecule-sensing MESA systems

| MESA | Ligand | Vehicle (% in stock) | Stock concentration | Vehicle (% in culture) | Working concentration |
|---|---|---|---|---|---|
| Rapamycin | Rapamycin (Santa Cruz Biotechnology sc-3504) | 50% DMSO, 50% $H_2O$ | 0.1 mM | 0.05% DMSO | 0.1 µM |
| | Rapalog (Takara AP21967) | 100% EtOH | 0.1 mM | 0.1% EtOH | 0.1 µM |
| Gibberellin | GA3-AM (Tocris 5407) | 100% DMSO | 10 mM | 0.1% DMSO | 10 µM |
| Abscisic acid | ABA (GoldBio A-050) | 100% EtOH | 100 mM | 0.1% EtOH | 100 µM |

**Supplementary Table 2. Rapamycin-sensing MESA expression normalization doses**[1]

| Plasmid # | MESA components | Initial (S6a) | Round 1 (S6b) | Round 2 (S6c) | Round 3/Final (S6d, 2b) |
|---|---|---|---|---|---|
| pPD801 | PC CD28-TMD | 25 | 18 | 32.3 | 21.9 |
| pPD802 | PC GpA-TMD | 25 | 1 | 1.9 | 3.7 |
| pPD803 | PC FGFR1-TMD | 25 | 750 | 75.0 | 88.2[2] |
| pPD804 | PC FGFR2-TMD | 25 | 2 | 9.2 | 9.6 |
| pPD805 | PC FGFR3-TMD | 25 | 2 | 14.6 | 8.7 |
| pPD806 | PC FGFR4-TMD | 25 | 9 | 7.9 | 16.3 |
| pPD807 | PC FGFR-S-TMD | 25 | 3 | 6.9 | 6.5 |
| pPD808 | PC VEGFR1-TMD | 25 | 1 | 1.6 | 2.0 |
| pPD809 | PC EphA4-TMD | 25 | 4 | 9.2 | 9.6 |
| pPD810 | TC CD28-TMD | 25 | 25 | 14.8 | 28.5 |
| pPD811 | TC GpA-TMD | 25 | 9 | 25.0 | 25.0 |
| pPD812 | TC FGFR1-TMD | 25 | 50 | 19.1 | 13.6 |
| pPD813 | TC FGFR2-TMD | 25 | 15 | 34.7 | 37.6 |
| pPD814 | TC FGFR3-TMD | 25 | 24 | 16.2 | 22.0 |
| pPD815 | TC FGFR4-TMD | 25 | 181 | 13.5 | 24.9 |
| pPD816 | TC FGFR-S-TMD | 25 | 16 | 36.2 | 78.9 |
| pPD817 | TC VEGFR1-TMD | 25 | 12 | 38.5 | 20.5 |
| pPD818 | TC EphA4-TMD | 25 | 2 | 13.9 | 14.2 |
| pPD819 | PC Valine-TMD | 25 | 12 | 18.5 | 8.2 |
| pPD832 | TC Valine-TMD | 25 | 750 | 211.4 | 85.8 |

[1]Plasmid doses in this table are those used in a single well of a 24-well plate. As transfection doses scale with the volume of media, for the western blots shown in these figures (**Fig 2b**, **Supplementary Fig. 6**), a mass four times the doses shown in this table were used (i.e., 100 ng of pPD810 TC CD28-TMD).

[2]88.2 ng of plasmid (used per well of a 24-well plate) was used to calculate the plasmid dose for **Fig. 2b**. This resulted in multiple full-length PC bands (49 kDa and approx. 55 kDa, the latter of which we speculate is glycosylated). The 49 kDa band had a similar intensity as the other 49 kDa PC bands from MESA with other TMDs. To keep the total amount of full-length PC in the cells constant, we normalized the sum of the 49 and 55 kDa bands to the intensities of the other 49 kDa bands. This resulted in a dose of 33.5 ng (per well of a 24-well plate) of FGFR1-TMD PC used in functional assays (**Fig. 2c and 2d**).

**Supplementary Table 3. Instrument specifications for analytical flow cytometry to quantify reporter expression**

| Instrument | Fluorescent protein | Parameter/ Channel name | Excitation laser | Filter set |
|---|---|---|---|---|
| BD LSR Fortessa | EBFP2 | Pacific Blue | Violet, 405 nm | 450/50 |
| | EYFP | FITC | Blue, 488 nm | 505LP, 530/30 |
| | DsRed2 | PE-Texas Red | Light Green, 552 nm | 600LP, 610/20 |

**Supplementary Table 4. Instrument specifications for analytical flow cytometry to quantify surface expression**

| Instrument | Fluorescent protein | Parameter/ Channel name | Excitation laser | Filter set |
|---|---|---|---|---|
| BD LSR Fortessa | EBFP2 | Pacific Blue | Violet, 405 nm | 450/50 |
| | Phycoerythrin | PE | Light Green, 552 nm | 575/26 |
| | miRFP670 | APC | Red, 640 nm | 670/30 |
| | Allophycocyanin | APC | Red, 640 nm | 670/30 |

**Supplementary Table 5. Instrument specifications for confocal microscopy to quantify FRET**

| Instrument | Fluorescent protein | Excitation laser | Beam splitter | Sensor | Acquisition band-pass filter |
|---|---|---|---|---|---|
| Leica SP5 II Laser Scanning Confocal Microscope | mCerulean (D) | Argon 458 nm | 458/514 double dichroic | HyD* | 465–500 nm |
| | mVenus (A) | Argon 514 nm | | HyD* | 520–560 nm |
| | FRET | Argon 458 nm | | HyD* | 520–560 nm |

*Abbreviation: HyD, hybrid detector

### SUPPLEMENTARY NOTES

#### Supplementary Note 1

Below are the outcomes from one-way ANOVAs and Tukey's HSD tests. Null hypotheses were that there existed no effects of PC C-terminal peptide on the measured reporter expression, evaluated separately for vehicle-and ligand-treated conditions.

Functional signaling assay in **Fig. 1c, Supplementary Fig. 3e**

Vehicle treatment

- PC C-terminal peptide $p = 3.9\times10^{-13}$
  - All differences were significant (all $p < 0.05$) except for:
    - Comparisons between PCs that each contain a PRS-based peptide
    - PCs that contain the $AIP_Y$ or $AIP_A$ peptide and all of the PRS-based peptides
    - PC-AIP and PC-$AIP_K$ or PC-$AIP_M$
    - PC-$AIP_A$ and PC-$AIP_Y$
    - PC-$AIP_M$ and PC-$AIP_K$
  - Notably, included within the comparisons that are significantly different (all $p \ll 0.01$) are comparisons between each PC variant with a C-terminal peptide and the PC without a C-terminal peptide.

Ligand treatment

- PC C-terminal peptide $p = 7.8\times10^{-19}$
  - All differences were significant (all $p < 0.05$) except for:
    - Comparisons between PCs that each contain a PRS-based peptide
    - PC-AIP and PC-$AIP_K$

Below are the outcomes from one-way ANOVAs and Tukey's HSD tests. Null hypotheses were that there existed no effects of TMD on the measured reporter expression, evaluated separately for vehicle- and ligand-treated conditions.

Functional signaling assay in **Fig. 2c, Supplementary Fig. 7b**

Vehicle treatment

- TMD $p = 2.0\times10^{-9}$
  - All differences were not significant (all $p > 0.05$) except for comparisons between each TMD and the FGFR1 TMD ($p \ll 0.001$).

Ligand treatment

- TMD $p = 2.4\times10^{-18}$
  - All differences were significant (all $p < 0.05$) except for:
    - CD28 and Valine
    - Comparisons between EphA4 and FGFR2, FGFR3, FGFRS, and VEGFR1
    - Comparisons between FGFRS and FGFR2, FGFR3, and VEGFR1
    - Comparisons between FGFR2, FGFR3 and VEGFR1
- Notably, included within the comparisons that are significantly different (all $p \ll 0.001$) are comparisons between each TMD and CD28, excluding Valine.

**Supplementary Note 2**

Below are the outcomes from two-way ANOVAs and Tukey's HSD tests. Null hypotheses were that there existed no effects of PC C-terminal peptide or PC inner linker (PCIL), ligand treatment, or their interaction on the measured reporter expression.

Functional signaling assay in **Fig. 1c**

- PC C-terminal peptide $p = 1.6 \times 10^{-33}$
  - All differences were significant (all $p << 0.01$) except for AIP vs $AIP_K$ and all comparisons between PRS-containing samples (all $p > 0.9$).
- Treatment $p = 1.4 \times 10^{-37}$
  - The samples with no PC C-terminal peptide and with any AIP PC C-terminal peptide show a significant increase in reporter expression upon ligand treatment (all $p << 0.01$).
  - The samples with any PRS PC C-terminal peptide show no significant increase in reporter expression upon ligand treatment (all $p = 1$).
- Interaction between TMD and ligand treatment $p = 3.6 \times 10^{-31}$

Functional signaling assay in **Fig. 1f**

- PCIL $p = 1.3 \times 10^{-29}$
- Receptors with EN performed similarly to receptors with RKMK, RN, RR, and RRLI but significantly differently than all other receptors ($p < 0.05$). Treatment $p = 2.5 \times 10^{-34}$
  - All PCILs produced significant increases in reporter expression upon ligand treatment (all $p << 0.01$)
  - Compared to the base case EN PCIL:
    - RR(R), RN, and RRLI show no significant change to background signal ($p > 0.05$) and either no change ($p > 0.05$) or a significant decrease ($p < 0.05$) in ligand-induced signal.
    - RRR, RRRR, RKMK, and KMKS show a significant increase to background signal ($p << 0.01$) and either no change ($p > 0.05$) or a significant increase to ligand-induced signal ($p < 0.01$).
    - RR shows a significant increase to background signal ($p << 0.01$) and a significant decrease to ligand-induced signal ($p < 0.01$).
- Interaction between TMD and ligand treatment $p = 3.5 \times 10^{-11}$

Below are the outcomes from two-way ANOVAs and Tukey's HSD tests. Null hypotheses were that there existed no effects of TMD, ligand treatment, or their interaction on the measured reporter expression.

Functional signaling assay in **Fig. 2c**

- TMD $p = 1.7 \times 10^{-32}$
  - The following TMDs performed significantly differently than the CD28 TMD: FGFR1, FGFR2, FGFR3, FGFR4, FGFR-S, VEGFR1 (all $p << 0.01$).
  - The GpA ($p = 0.13$) and Valine ($p = 0.93$) TMDs did not perform significantly differently than the CD28 TMD.
  - No vehicle-only conditions were different than the CD28-TMD vehicle-only condition (all $p > 0.3$).
  - Of the ligand-treated conditions, only the Valine-TMD ligand ($p = 0.6$) condition was not significantly different from the CD28-TMD ligand-treated condition ($p << 0.01$ for all conditions except GpA-TMD where $p = 0.015$).
- Treatment $p = 1.8 \times 10^{-31}$
  - The following TMDs did show significantly different reporter expression upon ligand treatment: CD28, GpA, FGFR1, FGFR4, Valine (all $p << 0.01$).
  - The following TMDs did not show significantly different reporter expression upon ligand treatment: FGFR2, FGFR3, EphA4, VEGFR1 (all $p = 1$).
- Interaction between TMD and ligand treatment $p = 1.5 \times 10^{-30}$

Below are the outcomes from two-way ANOVAs and Tukey's HSD tests. Null hypotheses were that there existed no effects of TMD, ligand treatment, or their interaction on the measured NFRET values.

NFRET assay in **Fig. 4a**

- TMD $p = 5.2\times10^{-10}$
  - All differences were significant (all $p << 0.01$).
- Treatment $p = 1.6\times10^{-20}$
  - Vehicle vs. ligand was significant (all $p << 0.01$).
- Interaction between TMD and ligand treatment $p = 8.9\times10^{-6}$

NFRET assay in **Fig. 4b**

- TMD $p = 1.6\times10^{-32}$
  - All differences were significant (all $p << 0.01$).
- Ligand treatment dose $p = 3.3\times10^{-66}$
  - All differences were significant except for 50 vs. 100 ($p = 0.15$), 50 vs. 1000 ($p = 0.06$), 100 vs. 500 ($p = 0.99$), 100 vs. 1000 ($p = 1$), and 500 vs. 1000 ($p = 1$).
- Interaction between TMD and ligand treatment $p = 7.0\times10^{-12}$

Below are the outcomes from two-way ANOVAs and Tukey's HSD tests. Null hypotheses were that there existed no effects of position of the first dimerizing residue (from the inner linker) in the synthetic TMDs, ligand treatment, or their interaction on the measured reporter expression.

Functional signaling assay in **Fig. 5b**

- Position of first dimerizing residue $p = 1.9\times10^{-14}$
  - All differences were significant (all $p < 0.05$) except for:
    - Comparisons between positions 10-15
    - Positions 4 and 6
    - Positions 8 and 9
- Treatment $p = 2.3\times10^{-5}$
  - Only TMDs with the first dimerizing residue in positions 4-9 produced significant increases in reporter expression upon ligand treatment (all $p << 0.01$)
  - Moving the TMD from positions 4-5, 5-6, 6-7, 7-8, 9-10 resulted in significant differences in ligand-induced reporter expression (all $p << 0.01$)
- Interaction between TMD and ligand treatment $p = 1.0\times10^{-4}$

Below are the outcomes from two-way ANOVAs and Tukey's HSD tests. Null hypotheses were that there existed no effects of TC TMD, PC TMD, or their interaction on the measured reporter expression, evaluated separately for vehicle- and ligand-treated conditions.

Functional signaling assay in **Fig. 2d, Supplementary Fig. 8c**

Vehicle treatment

- TC TMD $p = 1.6\times10^{-37}$
  - All differences were significant (all $p << 0.01$) except for comparisons between FGFR4, Valine, and CD28.
- PC TMD $p = 3.8\times10^{-17}$
  - All differences were significant (all $p << 0.01$) except for:
    - Comparisons between FGFR1, Valine, and CD28
    - FGFR4 vs. CD28
- Interaction between TC TMD and PC TMD $p = 3.0\times10^{-20}$
- Variance explained by TC TMD $\omega^2 = 0.64$
- Variance explained by PC TMD $\omega^2 = 0.10$

- Variance explained by the interaction $\omega^2 = 0.22$

Ligand treatment

- TC TMD $p = 9.8\times10^{-53}$
  - All differences were significant (all $p << 0.01$).
- PC TMD $p = 1.9\times10^{-44}$
  - All differences were significant (all $p << 0.01$).
- Interaction between TC TMD and PC TMD $p = 6.9\times10^{-33}$
- Variance explained by TC TMD $\omega^2 = 0.58$
- Variance explained by PC TMD $\omega^2 = 0.27$
- Variance explained by the interaction $\omega^2 = 0.14$

Functional signaling assays in **Fig. 6a**

Vehicle treatment

- TC TMD $p = 1.7\times10^{-33}$
  - All differences were significant (all $p << 0.01$) except for Valine vs. FGFR4 ($p = 0.97$).
- PC TMD $p = 7.3\times10^{-29}$
  - All differences were significant (all $p << 0.01$) except for Valine vs. FGFR1 ($p = 0.99$).
- Interaction between TC TMD and PC TMD $p = 2.1\times10^{-22}$
- Variance explained by TC TMD $\omega^2 = 0.45$
- Variance explained by PC TMD $\omega^2 = 0.28$
- Variance explained by the interaction $\omega^2 = 0.24$

Ligand treatment

- TC TMD $p = 8.8\times10^{-30}$
  - All differences were significant (all $p << 0.01$).
- PC TMD $p = 2.1\times10^{-19}$
  - All differences were significant (FGFR1 vs. CD28 $p = 0.04$, all other $p << 0.01$) except for GpA vs. FGFR4 ($p = 0.16$).
- Interaction between TC TMD and PC TMD $p = 2.7\times10^{-9}$
- Variance explained by TC TMD $\omega^2 = 0.63$
- Variance explained by PC TMD $\omega^2 = 0.21$
- Variance explained by the interaction $\omega^2 = 0.10$

Functional signaling assays in **Fig. 6b**

Vehicle treatment

- TC TMD $p = 6.2\times10^{-47}$
  - All differences were significant (all $p << 0.01$) except for Valine vs. FGFR4 ($p = 0.61$).
- PC TMD $p = 2.7\times10^{-44}$
  - All differences were significant (Valine vs. FGFR1 $p = 0.02$, all other $p << 0.01$) except for FGFR4 vs. CD28 ($p = 0.26$).
- Interaction between TC TMD and PC TMD $p = 6.2\times10^{-33}$
- Variance explained by TC TMD $\omega^2 = 0.45$
- Variance explained by PC TMD $\omega^2 = 0.35$
- Variance explained by the interaction $\omega^2 = 0.19$

Ligand treatment

- TC TMD $p = 2.6\times10^{-51}$
  - All differences were significant (Valine vs. FGFR4 $p = 0.03$, all other $p << 0.01$).
- PC TMD $p = 5.9\times10^{-39}$
  - All differences were significant (all $p << 0.01$).
- Interaction between TC TMD and PC TMD $p = 7.0\times10^{-23}$
- Variance explained by TC TMD $\omega^2 = 0.70$
- Variance explained by PC TMD $\omega^2 = 0.22$
- Variance explained by the interaction $\omega^2 = 0.07$

Functional signaling assays in **Fig. 6c**

Vehicle treatment

- TC TMD $p = 7.9\times10^{-55}$
  - All differences were significant (all $p << 0.01$) except for Valine vs. FGFR4 ($p = 1$).
- PC TMD $p = 5.6\times10^{-46}$
  - All differences were significant (all $p << 0.01$) except for Valine vs. FGFR1 ($p = 0.41$).
- Interaction between TC TMD and PC TMD $p = 8.5\times10^{-33}$
- Variance explained by TC TMD $\omega^2 = 0.61$
- Variance explained by PC TMD $\omega^2 = 0.27$
- Variance explained by the interaction $\omega^2 = 0.12$

Ligand treatment

- TC TMD $p = 2.1\times10^{-50}$
  - All differences were significant (all $p << 0.01$) except for Valine vs. FGFR4 ($p = 0.97$).
- PC TMD $p = 3.3\times10^{-51}$
  - All differences were significant (all $p << 0.01$).
- Interaction between TC TMD and PC TMD $p = 3.8\times10^{-26}$
- Variance explained by TC TMD $\omega^2 = 0.44$
- Variance explained by PC TMD $\omega^2 = 0.48$
- Variance explained by the interaction $\omega^2 = 0.07$

Functional signaling assays in **Fig. 6d**

Vehicle treatment

- TC TMD $p = 7.2\times10^{-36}$
  - All differences were significant (all $p << 0.01$) except for GpA vs. FGFR1 ($p = 0.99$).
- PC TMD $p = 1.2\times10^{-39}$
  - All differences were significant (all $p << 0.01$) except for GpA vs. FGFR4 ($p = 0.78$).
- Interaction between TC TMD and PC TMD $p = 6.3\times10^{-23}$
- Variance explained by TC TMD $\omega^2 = 0.34$
- Variance explained by PC TMD $\omega^2 = 0.49$
- Variance explained by the interaction $\omega^2 = 0.15$

Ligand treatment

- TC TMD $p = 4.7\times10^{-39}$
  - All differences were significant (all $p << 0.01$) except for Valine vs. FGFR4 ($p = 0.14$) and FGFR1 vs. CD28 ($p = 1$).
- PC TMD $p = 7.1\times10^{-41}$
  - All differences were significant (all $p < 0.05$) except for GpA vs. FGFR4 ($p = 0.19$) and FGFR1 vs. CD28 ($p = 0.71$).
- Interaction between TC TMD and PC TMD $p = 8.7\times10^{-21}$
- Variance explained by TC TMD $\omega^2 = 0.40$
- Variance explained by PC TMD $\omega^2 = 0.48$
- Variance explained by the interaction $\omega^2 = 0.11$

Below are the outcomes from two-way ANOVAs and Tukey's HSD tests. Null hypotheses were that there existed no effects of CTEVp TMD, NTEVp TMC, or their interaction on the measured reporter expression in the presence of either ligand or vehicle.

Functional signaling assays in **Fig. 6e**

Vehicle treatment

- CTEVp TMD $p = 7.1\times10^{-9}$
  - All differences were significant (all $p << 0.01$) except for GpA vs. FGFR4 ($p = 0.08$).
- NTEVp TMD $p = 4.6\times10^{-11}$
  - All differences were significant except for GpA vs. FGFR1 ($p = 0.36$) and FGFR4 vs. FGFR1 ($p = 0.31$).
- Interaction between CTEVp TMD and NTEVp TMD $p = 1.8\times10^{-22}$

- Variance explained by CTEVp TMD $\omega^2 = 0.06$
- Variance explained by NTEVp TMD $\omega^2 = 0.09$
- Variance explained by the interaction $\omega^2 = 0.82$

Ligand treatment

- CTEVp TMD $p = 9.3\times10^{-8}$
  - All differences were significant (all $p \ll 0.01$) except for GpA vs. CD28 ($p = 0.86$), GpA vs. FGFR4 ($p = 0.24$), andFGFR4 vs. CD28 ($p = 0.051$).
- NTEVp TMD $p = 4.1\times10^{-6}$
  - All differences were significant (all $p \ll 0.01$) except for GpA vs. FGFR4 ($p = 0.54$), GpA vs. FGFR1 ($p = 1$), and FGFR4 vs. FGFR1 ($p = 0.56$).
- Interaction between CTEVp TMD and NTEVp TMD $p = 3.5\times10^{-3}$
- Variance explained by CTEVp TMD $\omega^2 = 0.35$
- Variance explained by NTEVp TMD $\omega^2 = 0.23$
- Variance explained by the interaction $\omega^2 = 0.13$

Functional signaling assays in **Fig. 6f**

Vehicle treatment

- CTEVp TMD $p = 3.0\times10^{-24}$
  - All differences were significant (all $p \ll 0.01$) except for GpA vs. FGFR4 ($p = 0.75$) and FGFR4 vs. FGFR1 ($p = 0.06$).
- NTEVp TMD $p = 8.1\times10^{-22}$
  - All differences were significant (all $p \ll 0.01$) except for FGFR4 vs. FGFR1 ($p = 0.98$).
- Interaction between CTEVp TMD and NTEVp TMD $p = 1.4\times10^{-29}$
- Variance explained by CTEVp TMD $\omega^2 = 0.20$
- Variance explained by NTEVp TMD $\omega^2 = 0.14$
- Variance explained by the interaction $\omega^2 = 0.65$

Ligand treatment

- CTEVp TMD $p = 9.5\times10^{-23}$
- All differences were significant (all $p \ll 0.01$) except for GpA vs. FGFR4 ($p = 0.08$)
- NTEVp TMD $p = 7.5\times10^{-18}$
- All differences were significant (all $p \ll 0.01$) except for FGFR4 vs. FGFR1 ($p = 0.30$) and GpA vs. FGFR1 ($p = 0.36$).
- Interaction between CTEVp TMD and NTEVp TMD $p = 4.7\times10^{-13}$
- Variance explained by CTEVp TMD $\omega^2 = 0.54$
- Variance explained by NTEVp TMD $\omega^2 = 0.25$
- Variance explained by the interaction $\omega^2 = 0.17$

**Supplementary Note 3**

Below are the outcomes from three-way ANOVAs and Tukey's HSD tests. Null hypotheses were that there existed no effects of TC TMD, PC TMD, ligand treatment, or their pairwise interactions on the measured NFRET values or reporter expression.

Functional signaling assays in **Fig. 2d**

- TC TMD $p = 2.5\times10^{-92}$
  - All differences were significant (all $p << 0.01$) except for Valine vs. FGFR4 ($p = 0.21$).
- PC TMD $p = 2.8\times10^{-75}$
  - All differences were significant (all $p << 0.01$) except for FGFR1 vs. CD28 ($p = 0.06$), FGFR4 vs. FGFR1 ($p = 0.83$), and Valine vs. GpA ($p = 0.10$).
- Ligand treatment $p = 2.6\times10^{-124}$
  - Vehicle vs. ligand was significant for all TMD pairs (all $p << 0.01$).
- Interaction between TC TMD and PC TMD $p = 8.9\times10^{-56}$
- Interaction between TC TMD and ligand treatment $p = 8.6\times10^{-89}$
- Interaction between PC TMD and ligand treatment $p = 5.0\times10^{-73}$
- Interaction between TC TMD, PC TMD, and ligand treatment $p = 4.2\times10^{-54}$

Functional assays in **Supplementary Fig. 8a-b**

- TC TMD $p = 2.3\times10^{-169}$
- PC TMD $p = 1.9\times10^{-93}$
- Ligand $p = 2.1\times10^{-137}$
  - Significant ligand-induced signaling was observed for all receptor pairs that included a CD28-TMD on the TC (all $p << 0.01$).
  - Significant ligand-induced signaling was not observed for any receptor pairs that included an EphA4, FGFR2, or FGFR3-TMD on the TC (all $p > 0.15$).
  - Significant ligand-induced signaling was observed for receptor pairs that included a VEGFR1-TMD on the TC and a either a CD28, GpA, FGFR1, FGFR3, FGFR4, or Valine-TMD on the PC (all $p << 0.02$).
  - Significant ligand-induced signaling was not observed for all receptor pairs that included a VEGFR1-TMD on the TC and either a FGFR2, EphA4, or VEGFR1-TMD on the PC (all $p > 0.5$).
- Interaction between TC TMD and PC TMD $p = 3.2\times10^{-115}$
- Interaction between TC TMD and ligand treatment $p = 1.7\times10^{-167}$
- Interaction between PC TMD and ligand treatment $p = 6.6\times10^{-92}$
- Interaction between TC TMD, PC TMD, and ligand treatment $p = 6.5\times10^{-114}$

NFRET assays in **Fig. 4c**

- FRB chain TMD $p = 1.0\times10^{-12}$
  - All differences were significant (all $p << 0.01$).
- FKBP chain TMD $p = 3.9\times10^{-15}$
  - All differences were significant (all $p << 0.01$).
- Ligand treatment $p = 6.0\times10^{-59}$
  - Vehicle vs. ligand was significant for all TMD pairs (all $p << 0.01$).
- Interaction between FRB chain TMD and FKBP chain TMD $p = 5.2\times10^{-29}$
- Interaction between FRB chain TMD and ligand treatment $p = 5.5\times10^{-11}$
- Interaction between FKBP chain TMD and ligand treatment $p = 9.4\times10^{-11}$
- Interaction between FRB chain TMD, FKBP chain TMD, and ligand treatment $p = 1.1\times10^{-10}$

NFRET assays in **Supplementary Fig. 11a**

- FRB chain TMD $p = 3.5\times10^{-16}$
  - All differences were significant (GpA vs. FGFR4 $p = 0.02$, all other $p << 0.01$).
- FKBP chain TMD $p = 5.2\times10^{-20}$
  - All differences were significant except for GpA vs. CD28 ($p = 0.10$).

- Ligand treatment $p = 1.2\times10^{-59}$
  - Vehicle vs. ligand was significant for all TMD pairs (all $p << 0.01$).
- Interaction between FRB chain TMD and FKBP chain TMD $p = 8.8\times10^{-32}$
- Interaction between FRB chain TMD and ligand treatment $p = 4.9\times10^{-7}$
- Interaction between FKBP chain TMD and ligand treatment $p = 5.9\times10^{-9}$
- Interaction between FRB chain TMD, FKBP chain TMD, and ligand treatment $p = 6.0\times10^{-13}$

Functional signaling assays in **Fig. 6a**
- TC TMD $p = 2.1\times10^{-50}$
  - All differences were significant (all $p << 0.01$).
- PC TMD $p = 3.4\times10^{-31}$
  - All differences were significant (all $p << 0.01$) except for FGFR1 vs. CD28 ($p = 0.14$). and GpA vs. FGFR4 ($p = 0.83$).
- Ligand treatment $p = 2.2\times10^{-74}$
  - Vehicle vs. ligand was significant for all pairs (TC CD28-TMD with PC-CD28 TMD $p = 0.03$, all other $p << 0.01$) except for TC FGFR4-TMD with PC CD28-TMD ($p = 0.17$), TC Valine-TMD with PC FGFR1-TMD ($p = 1$), TC CD28-TMD with PC Valine-TMD ($p = 0.73$), TC FGFR4-TMD with PC Valine-TMD ($p = 0.96$), TC Valine-TMD with PC Valine-TMD ($p = 1$).
- Interaction between TC TMD and PC TMD $p = 2.0\times10^{-13}$
- Interaction between TC TMD and ligand treatment $p = 3.3\times10^{-41}$
- Interaction between PC TMD and ligand treatment $p = 2.0\times10^{-23}$
- Interaction between TC TMD, PC TMD, and ligand treatment $p = 2.0\times10^{-12}$

Functional signaling assays in **Fig. 6b**
- TC TMD $p = 9.5\times10^{-99}$
  - All differences were significant (all $p << 0.01$).
- PC TMD $p = 6.5\times10^{-81}$
  - All differences were significant (all $p << 0.01$).
- Ligand treatment $p = 3.8\times10^{-66}$
  - Vehicle vs. ligand was significant for all pairs (all $p << 0.01$) except for TC FGFR4-TMD with PC CD28-TMD ($p = 0.97$), TC Valine-TMD with PC CD28-TMD ($p = 1$), TC CD28-TMD with PC FGFR1-TMD ($p = 0.84$), TC FGFR4-TMD with PC FGFR1-TMD ($p = 1$), TC GpA-TMD with PC FGFR1-TMD ($p = 0.68$), TC Valine-TMD with PC FGFR1-TMD ($p = 1$), TC Valine-TMD with PC FGFR4-TMD ($p = 1$), TC CD28-TMD with PC Valine-TMD ($p = 1$), TC FGFR4-TMD with PC Valine-TMD ($p = 1$), TC Valine-TMD with PC Valine-TMD ($p = 1$).
- Interaction between TC TMD and PC TMD $p = 4.3\times10^{-53}$
- Interaction between TC TMD and ligand treatment $p = 5.4\times10^{-50}$
- Interaction between PC TMD and ligand treatment $p = 1.3\times10^{-25}$
- Interaction between TC TMD, PC TMD, and ligand treatment $p = 2.3\times10^{-27}$

Functional signaling assays in **Fig. 6c**
- TC TMD $p = 1.5\times10^{-102}$
  - All differences were significant (all $p << 0.01$) except for Val vs. FGFR4 ($p = 0.99$).
- PC TMD $p = 1.6\times10^{-97}$
  - All differences were significant (all $p << 0.01$) except for Val vs. FGFR1 ($p = 0.16$).
- Ligand treatment $p = 7.2\times10^{-79}$
  - Vehicle vs. ligand was significant for all pairs (all $p << 0.01$) except for TC FGFR4-TMD with PC CD28-TMD ($p = 1$), TC Valine-TMD with PC CD28-TMD ($p = 0.75$), TC CD28-TMD with PC FGFR1-TMD ($p = 0.10$), TC FGFR4-TMD with PC FGFR1-TMD ($p = 1$), TC GpA-TMD with PC FGFR1-TMD ($p = 0.98$), TC Valine-TMD with PC FGFR1-TMD ($p = 1$), TC Valine-TMD with PC FGFR4-TMD ($p = 0.08$), TC CD28-TMD with PC Valine-TMD ($p = 0.95$), TC FGFR4-TMD with PC Valine-TMD ($p = 1$), TC Valine-TMD with PC Valine-TMD ($p = 1$).

- Interaction between TC TMD and PC TMD $p = 4.7\times10^{-56}$
- Interaction between TC TMD and ligand treatment $p = 3.3\times10^{-41}$
- Interaction between PC TMD and ligand treatment $p = 2.4\times10^{-59}$
- Interaction between TC TMD, PC TMD, and ligand treatment $p = 9.2\times10^{-27}$

Functional signaling assays in **Fig. 6d**

- TC TMD $p = 4.2\times10^{-72}$
    - All differences were significant (all $p$ << 0.01) except for FGFR1 vs. CD28 ($p = 0.12$).
- PC TMD $p = 1.0\times10^{-75}$
    - All differences were significant (all $p$ << 0.01) except for FGFR4 vs. CD28 ($p = 0.49$) and GpA vs. FGFR4 ($p = 0.10$).
- Ligand treatment $p = 2.2\times10^{-98}$
    - Vehicle vs. ligand was significant for all pairs (all $p$ << 0.01) except for TC CD28-TMD with PC Valine-TMD ($p = 0.31$), TC FGFR4-TMD with PC Valine-TMD ($p = 1$), TC Valine-TMD with PC Valine-TMD ($p = 1$)
- Interaction between TC TMD and PC TMD $p = 1.7\times10^{-39}$
- Interaction between TC TMD and ligand treatment $p = 7.3\times10^{-49}$
- Interaction between PC TMD and ligand treatment $p = 1.7\times10^{-53}$
- Interaction between TC TMD, PC TMD, and ligand treatment $p = 9.1\times10^{-23}$

Functional signaling assays in **Supplementary Fig. 30b**

- TC TMD $p = 1.8\times10^{-88}$
    - All differences were significant (all $p$ << 0.01) except for GpA vs. CD28 ($p = 0.65$).
- PC TMD $p = 1.3\times10^{-70}$
    - All differences were significant (all $p$ << 0.01) except for FGFR1 vs. CD28 ($p = 0.48$), FGFR4 vs. CD28 ($p = 1$), and FGFR4 vs. FGFR1 ($p = 0.40$).
- Ligand treatment $p = 1.5\times10^{-88}$
    - Vehicle vs. ligand was significant for all pairs (all $p$ << 0.01) except for TC FGFR4-TMD with PC CD28-TMD ($p = 0.75$), TC Valine-TMD with PC CD28-TMD ($p = 1$), TC Valine-TMD with PC FGFR1-TMD ($p = 1$), TC Valine-TMD with PC FGFR4-TMD ($p = 1$), TC Valine-TMD with PC GpA-TMD ($p = 0.61$), TC FGFR4-TMD with PC Valine-TMD ($p = 1$), TC Valine-TMD with PC Valine-TMD ($p = 1$).
- Interaction between TC TMD and PC TMD $p = 1.9\times10^{-54}$
- Interaction between TC TMD and ligand treatment $p = 2.4\times10^{-67}$
- Interaction between PC TMD and ligand treatment $p = 2.7\times10^{-40}$
- Interaction between TC TMD, PC TMD, and ligand treatment $p = 2.3\times10^{-40}$

Below are the outcomes from three-way ANOVAs and Tukey's HSD tests. Null hypotheses were that there existed no effects of NTEVp TMD, CTEVp TMD, ligand treatment, or their pairwise interactions on the measured reporter expression.

Functional signaling assays in **Fig. 6e**

- CTEVp TMD $p = 3.0\times10^{-10}$
  - All differences were significant (all $p << 0.01$) except for FGFR4 vs. CD28 ($p = 0.20$), GpA vs. CD28 ($p = 1$), and GpA vs. FGFR4 ($p = 0.20$).
- NTEVp TMD $p = 1.2\times10^{-5}$
  - All differences were significant (all $p << 0.01$) except for FGFR4 vs. FGFR1 ($p = 0.91$), GpA vs. FGFR1 ($p = 1$), GpA vs. FGFR4 ($p = 0.97$).
- Ligand treatment $p = 1.2\times10^{-66}$
  - Vehicle vs. ligand was significant for all TMD pairs (all $p << 0.01$).
- Interaction between CTEVp TMD and NTEVp TMD $p = 2.1\times10^{-7}$
- Interaction between CTEVp TMD and ligand treatment $p = 4.2\times10^{-9}$
- Interaction between NTEVp TMD and ligand treatment $p = 5.7\times10^{-9}$
- Interaction between CTEVp TMD, NTEVp TMD, and ligand treatment $p = 1.0\times10^{-2}$

Functional signaling assays in **Fig. 6f**

- CTEVp TMD $p = 1.8\times10^{-44}$
  - All differences were significant (all $p << 0.01$) except for GpA vs. FGFR4 ($p = 0.27$).
- NTEVp TMD $p = 3.3\times10^{-17}$
  - All differences were significant (all $p << 0.01$) except for FGFR4 vs. FGFR1 ($p = 0.31$).
- Ligand treatment $p = 3.9\times10^{-83}$
  - Vehicle vs. ligand was significant for all TMD pairs (all $p << 0.01$).
- Interaction between CTEVp TMD and NTEVp TMD $p = 7.9\times10^{-40}$
- Interaction between CTEVp TMD and ligand treatment $p = 1.0\times10^{-17}$
- Interaction between NTEVp TMD and ligand treatment $p = 9.8\times10^{-35}$
- Interaction between CTEVp TMD, NTEVp TMD, and ligand treatment $p = 2.6\times10^{-17}$

Functional signaling assays in **Supplementary Fig. 28b**

- CTEVp TMD $p = 1.5\times10^{-50}$
  - All differences were significant (all $p << 0.01$).
- NTEVp TMD $p = 7.0\times10^{-21}$
  - All differences were significant (all $p << 0.01$) except for FGFR4 vs. FGFR1 ($p = 0.14$).
- Ligand treatment $p = 1.5\times10^{-75}$
  - Vehicle vs. ligand was significant for all TMD pairs (all $p << 0.01$).
- Interaction between CTEVp TMD and NTEVp TMD $p = 1.4\times10^{-46}$
- Interaction between CTEVp TMD and ligand treatment $p = 3.4\times10^{-33}$
- Interaction between NTEVp TMD and ligand treatment $p = 9.8\times10^{-21}$
- Interaction between CTEVp TMD, NTEVp TMD, and ligand treatment $p = 8.5\times10^{-9}$

**Supplementary Note 4**

Below are the outcomes two-tailed Welch's *t*-tests followed by BH procedure. Null hypotheses were that there existed no effect of TMD pairing on NFRET fold difference within each harvest method group. TMD pairings are annotated as follows:

A = FRB CD28-TMD mV/FKBP CD28-TMD mC
B = FRB FGFR4-TMD mV/FKBP FGFR4-TMD mC
C = FRB CD28-TMD mV/FKBP FGFR4-TMD mC
D = FRB CD28-TMD mC/FKBP FGFR4-TMD mV
E = FRB FGFR4-TMD mV/FKBP CD28-TMD mC
F = FRB FGFR4-TMD mC/FKBP CD28-TMD mV

NFRET assays in **Supplementary Fig. 14b (bottom)**

FACS buffer

- A vs B $p = 0.0255$
- A vs C $p = 0.0044$
- A vs D $p = 0.0011$
- A vs E $p = 0.4067$ (not significant, n.s.)
- A vs F $p = 0.0007$
- B vs C $p = 0.6731$ (n.s.)
- B vs D $p = 0.0946$ (n.s.)
- B vs E $p = 0.0125$
- B vs F $p = 0.2977$ (n.s.)
- C vs D $p = 0.1167$ (n.s.)
- C vs E $p = 0.0063$
- C vs F $p = 0.4302$ (n.s.)
- D vs E $p = 0.0015$
- D vs F $p = 0.1695$ (n.s.)
- E vs F $p = 0.0062$

Trypsin

- A vs B $p = 0.0059$
- A vs C $p = 0.0002$
- A vs D $p = 0.0006$
- A vs E $p = 0.0050$
- A vs F $p = 0.0101$
- B vs C $p = 0.3644$ (n.s.)
- B vs D $p = 0.5538$ (n.s.)
- B vs E $p = 0.0116$
- B vs F $p = 1.0000$ (n.s.)
- C vs D $p = 0.6535$ (n.s.)
- C vs E $p = 0.0006$
- C vs F $p = 0.4717$ (n.s.)
- D vs E $p = 0.0086$
- D vs F $p = 0.6311$ (n.s.)
- E vs F $p = 0.0201$

### Supplementary Note 5

Below is a list of acronyms used in this study:

#### MESA features and domains

| | |
|---|---|
| MESA | Modular extracellular sensor architecture |
| TC | Target chain |
| PC | Protease chain |
| ECD | Extracellular domain (comprises ligand binding domain and extracellular linker) |
| ICD | Intracellular domain (comprises protease or transcription factor and inner linker) |
| TMD | Transmembrane domain |
| TEVp | Tobacco etch virus protease |
| NTEVp | N-terminal protein component of split TEVp |
| CTEVp | C-terminal protein component split TEVp |
| PRS | Protease recognition sequence (for TEVp) |
| AIP | Autoinhibitory peptide (for TEVp) |
| IL | Inner linker |
| PCIL | Protease chain inner linker |
| TF | Transcription factor |

#### Transmembrane domains

| | |
|---|---|
| CD28 | Cluster of differentiation 28 |
| EphA4 | Ephrin type-A receptor 4 |
| FGFR | Fibroblast growth factor receptor (1–4) |
| FGFR-Syn | Fibroblast growth factor receptor synthetic (a consensus sequence of 1–4) |
| GpA | Glycophorin A |
| VEGFR1 | Vascular endothelial growth factor 1 receptor |

#### Ligand binding domains

| | | |
|---|---|---|
| FKBP | FK506-Binding protein | binder of rapamycin/rapalog |
| FRB | FKBP rapamycin binding | binder of rapamycin/rapalog |
| GID1 | Gibberellin insensitive dwarf 1 | binder of gibberellin |
| GAI | Gibberellin insensitive | binder of gibberellin |
| ABI1 | Abscisic acid-insensitive1 | binder of abscisic acid |
| PYL1 | Pyrabactin like protein 1 | binder of abscisic acid |
| GBP | GFP-binding protein | binder of GFP, nanobody |
| scFv | Single chain variable fragment | a type of binding protein |
| G6-311 | scFv against VEGF | binder of VEGF, previously termed “V1” |
| B20-4.1 | scFv against VEGF | binder of VEGF, previously termed “V2” |

#### Ligands

| | |
|---|---|
| EtOH | Ethanol |
| DMSO | Dimethyl sulfoxide |
| Rapalog, Rapa | Rapamycin analog, here specifically Takara AP21967 |
| GA3 | Gibberellin |
| GA3-AM | Gibberellin analog, cell-permeable |
| ABA | Abscisic acid |
| GFP | Green fluorescent protein |
| sGFP | Secreted green fluorescent protein |
| VEGF | Vascular endothelial growth factor |
| sVEGF | secreted VEGF |

**Fluorescent proteins and other reporter proteins**

| | |
|---|---|
| EBFP2 | Enhanced blue fluorescent protein 2 |
| EYFP | Enhanced yellow fluorescent protein |
| mC | mCerulean |
| miRFP670 | monomeric infrared fluorescent protein (with 670 nm emission) |
| mV | mVenus |
| NanoLuc | NanoLuciferase |

**Cellular components**

| | |
|---|---|
| AKT | Protein kinase B |
| CAR | Chimeric antigen receptor |
| ERK | Extracellular signal-regulated kinases |
| GPCR | G protein-coupled receptor |
| JAK | Janus kinase |
| MAPK | Mitogen-activated protein kinases/ |
| NFAT | Nuclear factor of activated T-cells |
| PI3K | Phosphoinositide 3-kinase |
| PLCG | Phospholipase C |
| RTK | Receptor tyrosine kinase |
| STAT | Signal transducer and activator of transcription |
| synNotch | synthetic Notch |

**Reagents**

| | |
|---|---|
| BSA | Bovine serum albumin |
| DMEM | Dulbecco's Modified Eagle Medium |
| DNA | Deoxyribonucleic acid |
| EDTA | Ethylenediaminetetraacetic acid |
| FBS | Fetal bovine serum |
| HEPES | 4-(2-hydroxyethyl)-1-piperazineethanesulfonic acid |
| IgG | Immunoglobulin G |
| LB | Luria-Bertani |
| PBS | Phosphate-buffered saline |
| PVDF | Polyvinylidene fluoride |
| RCP | Rainbow calibration particles |
| RIPA | Radioimmunoprecipitation assay buffer |
| RNA | Ribonucleic acid |
| RNase | Ribonuclease that degrades RNA |
| SDS | Sodium dodecyl sulfate |
| TBS | Tris-buffered saline |
| TBST | Tris-buffered saline with Tween |
| TE | Tris EDTA buffer |
| URCP | Ultra-rainbow calibration particles |

**Other terms**

| | |
|---|---|
| ANOVA | Analysis of variance |
| APC | Allophycocyanin |
| AU | Arbitrary units |
| BCA | Bicinchoninic acid assay |
| ECL | Enhanced chemiluminescence |
| FACS | Fluorescence activated cell sorting |
| FRET | Förster resonance energy transfer |
| FSC-A | Area in the Forward Scatter channel |
| FSC-H | Height in the Forward Scatter channel |
| HSD | Tukey's honest significance test |
| HyD | Hybrid detector |
| MEFLs | Molecules of Equivalent Fluorescein |

| | |
|---|---|
| MEPTRs | Molecules of Equivalent PE-Texas Red |
| MFI | Mean fluorescence intensity (in arbitrary units) |
| NA | Numerical aperture |
| NEB | New England Biolabs |
| NFRET | Normalized FRET signal |
| PDB | Protein Data Bank |
| PE | Phycoerythrin |
| RLU | Relative luciferase units |
| S.E.M. | Standard error of the mean |
| SSC-A | Area in the Side Scatter channel |

### REFERENCES CITED IN THIS DOCUMENT


1. Clackson, T. *et al.* (1998) Redesigning an FKBP-ligand interface to generate chemical dimerizers with novel specificity. *Proc Natl Acad Sci U S A,* 95, 10437-10442.

2. Lomize, A.L., Pogozheva, I.D. (2017) TMDOCK: An Energy-Based Method for Modeling alpha-Helical Dimers in Membranes. *J Mol Biol,* 429, 390-398.

3. Stepanenko, O.V. *et al.* (2008) Understanding the role of Arg96 in structure and stability of green fluorescent protein. *Proteins,* 73, 539-551.

4. Kapust, R.B. *et al.* (2001) Tobacco etch virus protease: mechanism of autolysis and rational design of stable mutants with wild-type catalytic proficiency. *Protein Eng,* 14, 993–1000.
